# scASK: A novel ensemble framework for classifying cell types based on single-cell RNA-seq data

**DOI:** 10.1101/2020.06.07.138271

**Authors:** Bo Liu, Fang-Xiang Wu, Xiufen Zou

## Abstract

The Human Cell Atlas (HCA) is a large project that aims to identify all cell types in the human body. The dimension reduction and clustering for identification of cell types from single-cell RNA-sequencing (scRNA-seq) data have become foundational approaches to HCA. The major challenges of current computational analyses are of poor performance on large scale data and sensitive to initial data. We present a new ensemble framework called Adaptive Slice KNNs (scASK) to address the challenges for analysing scRNA-seq data with high dimensionality. scASK consists of three innovational modules, called DAS (Data Adaptive Slicing), MCS (Meta Classifiers Selecting) and EMS (Ensemble Mode Switching), respectively, which facilitate scASK to approximate a bias-variance tradeoff beyond classification. Thirteen real scRNA-seq datasets are used to evaluate the performance of scASK. Compared with five popular classification algorithms, our experimental results indicate that scASK achieves the best accuracy and robustness among all competing methods. In conclusion, adaptive slicing is an effective structural reduction procedure, and meanwhile scASK provides novel and robust ensemble framework especially for classifying cell types based on scRNA-seq data. scASK is publically available at https://github.com/liubo2358/scASKcmd.

## INTRODUCTION

The cell is the fundamental unit of living organisms, which is the key to study human biology and diseases (1). The Human Cell Atlas (HCA) project, with the ultimate goal of profiling all human cell types, has huge potential to transform many aspects of fundamental biology and clinical practice (2, 3), and thus attracts more and more attention from the scientific community. In just a few years, the ever-increasing scale of single-cell datasets (4, 5) and the emergence of new technologies and methods (6–8) have pushed single-cell transcriptome to a new level. Identification of subgroups (e.g., cell types or functional cell states) from single-cell data, as the most fundamental question (9, 10) for whole downstream analyses, has became the hot research topic and main driving force in single-cell transcriptome. However, the identification of cell types across datasets is confronting with both data and methodological challenges (3).

As mentioned above, popular methods for dimension reduction in single-cell transcriptome, like principal component analysis (PCA) and t-distributed stochastic neighbour embedding (t-SNE), often suffer from the high dimensionality and high variability in data. Although PCA and t-SNE have been widely used as de facto gold standards for analysing or visualizing single-cell data, these algorithms assume that the underlying data are drawn from a Gaussian or a t-distribution, which does not always hold for scRNA-seq data (11). The discrepancy between the assumed and actual distribution fundamentally limits the accuracy of the resulting predictions (12). In practice, PCA which relies on singular value decomposition (SVD) is always computationally expensive, the traditional PCA computing out all singular values *m × n* of matrix has time complexity of O(*min*{*mn*^2^, *m*^2^*n*^2^}) (13). The computational complexity makes it infeasible to apply to very large-scale datasets, for example, when the sample size is more than one million. Moreover, the exact SVD is an algorithm which is difficult to parallelize with high efficiency, due to its matrix factorization nature (14). t-SNE is another popular, but more complicated method for dimension reduction and is particularly well suited for the visualization of high-dimensional data (15). The t-SNE algorithm doesn’t always produce similar output on successive runs, for example, and there are additional hyperparameters (such as “perplexity”) related to the optimization process, which makes it tricky to interpret. In other words, t-SNE plots can sometimes be mysterious or misleading (16), it is a valuable tool in generating hypotheses and understanding, but does not produce conclusive evidence. Non-negative matrix factorization (NMF) is another matrix decomposition technique widely studied (17–19) in computational biology for dimension reduction (or feature extraction). Benefiting from the nonnegative constraint, the results of NMF are usually easier to interpret than those of SVD. However, the existence, uniqueness, effectiveness of NMF solutions still need to be further studied (20–22).

Paralleling with the dimension reduction, several clustering methods have been developed for identification of novel cell types, such as SNN-cliq (23), RaceID (24), pcaReduce (25), SC3 (26) and SIMLR (27). Most of these clustering methods are based on metrics comparing intra- and inter-cluster similarity without references to external information. As unsupervised learning, the validation for clustering is very difficult in the absence of prior knowledge (28). For example, given a dataset, each clustering method will always find some partitioning, no matter whether the structure exists or not (29). Mathematically, clustering is essentially a heuristic algorithm to detect the subpopulation structure of existing data, different methods usually lead to different clusters, and even for the same method, the choice of the number of clusters K or the initial states of the same set of data may affect the final results. From an algorithmic perspective, clustering is a feasible approach to identify novel cell types from existing data, but is not an ideal way to apply known labels to new data — if only one type of cells in a sample, clustering can do nothing. In contrast, for many samples there exists a hierarchy of cell-types and cell-states, and they may all be of interest (30). Based on the analysis above, as more and more known cell types have been located within the HCA. Therefore, it is appealing to develop more reliable methods to achieve high accuracy of cell type identification for new samples beyond clustering and dimension reduction.

Generally, a better approach of cell type identification for gene expression data with the label information is to use machine learning techniques based on supervised learning, such as classification. In the past few years, numerous classification methods from single-cell data have been proposed (31–35), which were focusing mostly on specific tissues (e.g., bone marrow and neurons) or specific phenomenon of data (e.g., dropout and cross-dataset). Several classification algorithms were employed by the methods above, such as Naive Bayes (NB), Decision Trees (DT), Support Vector Machine (SVM) and k Nearest Neighbours (kNN). Regardless of underlying data distribution and tuning parameters, all of these classifiers can make effective classifications on specific datasets. However, high dimensionality and high variability as intrinsic characteristics of single-cell data are always obstacles for all computational techniques. Classification of cell types from single-cell data with label information, which guarantees the high accuracy as well as the high robustness on most of datasets, is a very challenging problem in the current study. To the best of our knowledge, there is no generic method available that performs well for both the objectives on scRNA-seq data.

Due to the nature of scRNA-seq data, there are some challenges in single-cell data analysis. The first notorious characteristic of single-cell data is high dimensionality. In a typical single-cell dataset, the number of cells or genes can easily reach or exceed 10^4^, while new experimental techniques have raised the upper limit of cell counts to 10^6^ and are still growing. The second notorious characteristic of single-cell data is high sparsity. In some single-cell datasets, the proportion of zeroes is as high as 80%-90%. Unfortunately, even in non-zero data, there are a lot of high levels of technical noise and confounding factors. Based on the above facts, we need to fully consider the intrinsic characteristics of single-cell data when developing new methods. Specifically, how to fully extract the classification information hidden in high-dimensional data, while at the same time reducing the sensitivity of the algorithm to missing values and noise as much as possible.

In this study, we present a novel ensemble classification framework called scASK based on adaptive data slicing, which is the first generic ensemble classification framework especially for classifying cell types based on scRNA-seq data with high dimensionality. Compared with traditional individual or ensemble classifiers, the most remarkable advantage of scASK is that it applies the known cell-type labels to new samples with high accuracy and high robustness in a near-real time manner.

## MATERIAL AND METHODS

### The scASK framework

One important target for scRNA-seq analysis is the identification of cell types from different samples, where each cell-type consists of cells with the similar gene expression patterns (36). In this paper, we defined the similarity of gene expression patterns as the differential expression of cells and its structural similarity at different levels of gene expression values. Computational techniques focus on extracting, measuring and evaluating these cell-to-cell similarities in scRNA-seq data to construct the core functions of scASK. The framework of scASK is shown in Figure 1, which contains the following three modules:

**Figure 1.**
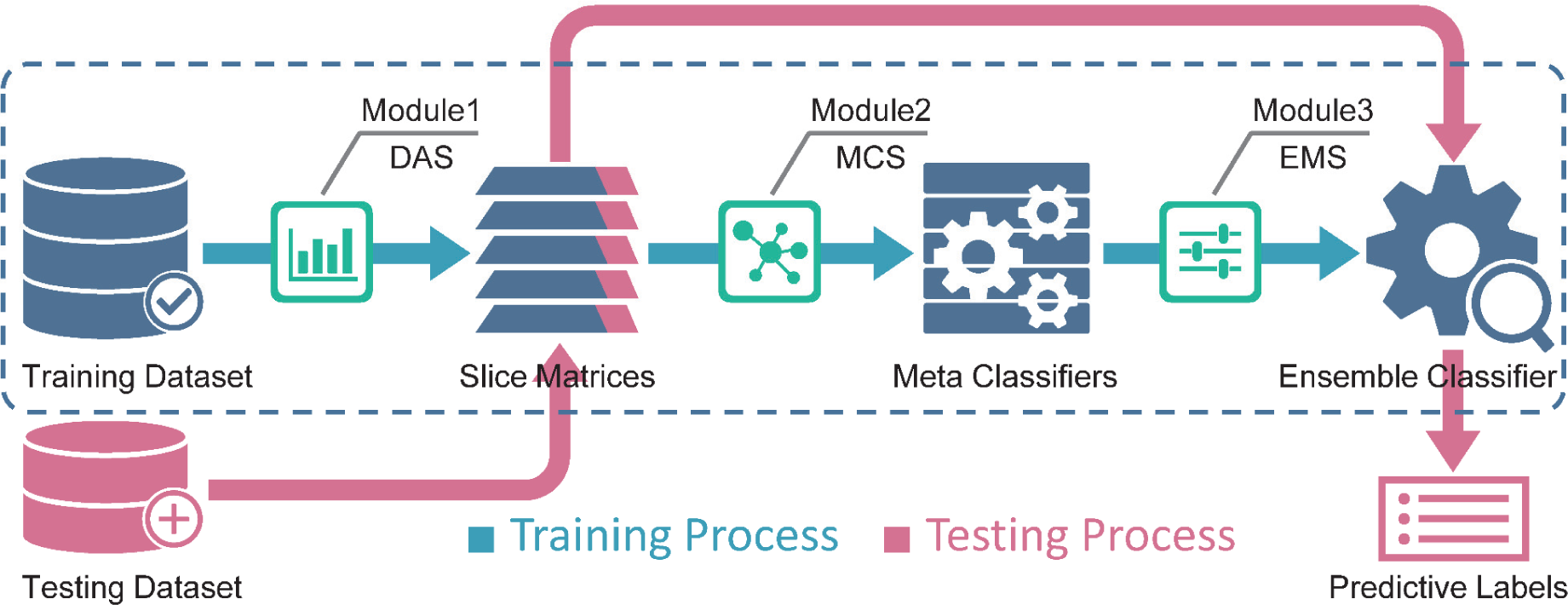
The framework of the proposed scASK. It consists of three modules, i.e., Data Adaptive Slicing (DAS), Meta Classifiers Selecting (MCS) and Ensemble Mode Switching (EMS), respectively. The first module DAS uses novel adaptive slicing procedure to extract latent classification features. The second module MCS selects three kinds of efficient distance measures Correlation, Jaccard and Cosine to construct competitive meta classifiers for diversity and complementarity. The third module EMS applies a new switching strategy for ensemble classification with dual operating mode to enhance the accuracy and the robustness of the final prediction. For evaluating the real performance of the ensemble classifier, we have reserved a portion of raw dataset as “testing dataset” independent of the training process, then calculate the overall accuracy by comparing predictive labels with true labels.

Module 1: Simplify the computational complexity of high-dimensional scRNA-seq data with novel adaptive slicing procedure for sufficiently extracting latent classification features.

Module 2: Select kNN with three kinds of efficient distance measures Correlation, Jaccard and Cosine to construct competitive meta classifiers for diversity and complementarity.

Module 3: Apply a new switching strategy for ensemble classification with dual operating mode to enhance the accuracy and the robustness of the final prediction.

### Module 1: Data Adaptive Slicing (DAS)

To sufficiently extract latent classification features from scRNA-seq data, we execute the module DAS on raw gene expression matrix. Then we obtain a series of binary matrices called slice matrices that are selected for the subsequent ensemble learning. The core function and basic principle of module 1 are shown in Figure 2.

**Figure 2.**
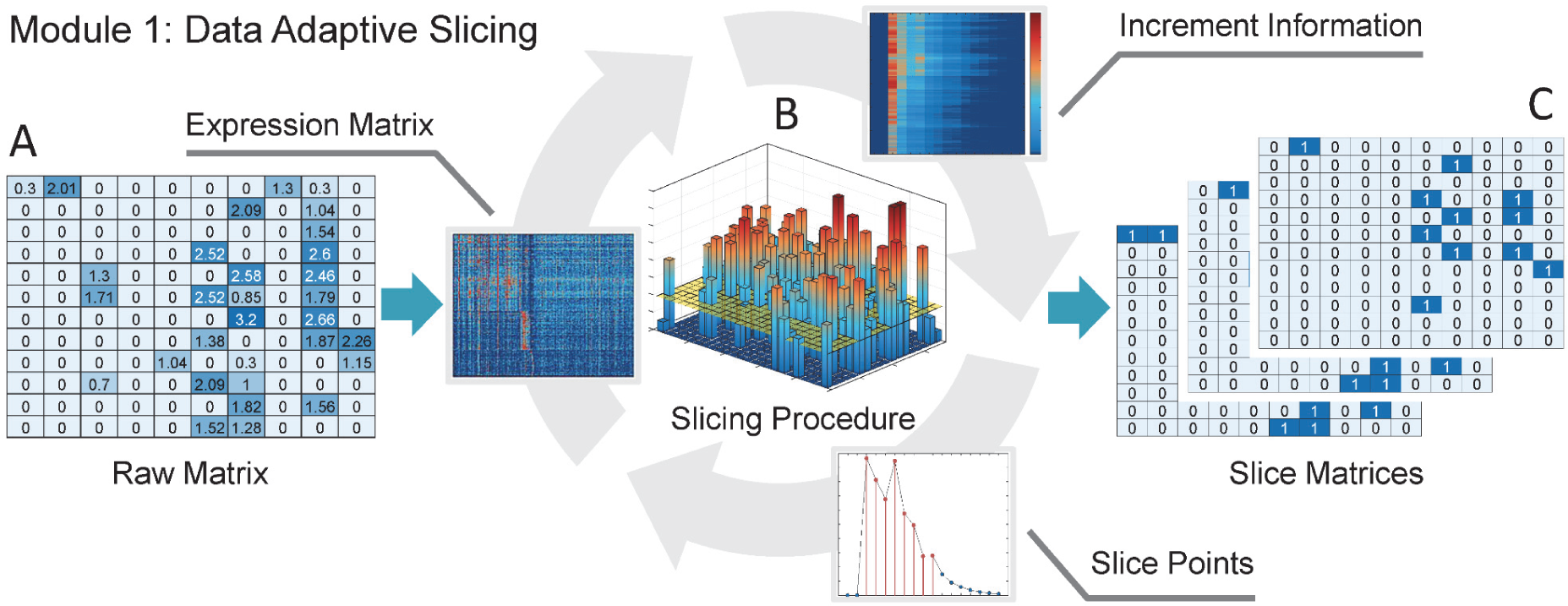
Module 1 of the scASK: Data Adaptive Slicing. (**A**) The raw matrix of scRNA-seq data with rows representing cells and columns representing genes, which measures the distribution of expression levels for each gene across a population of cells. (**B**) scASK stretches the data matrix along Z axis to three-dimensional space according to the range of gene expression values, and then slices the raw matrix to many slice matrices on selected slice points with the aid of quantifying the cumulation of binary switching in expression state between all slice matrices. (**C**) A series of binary slice matrices are obtained after executing the Data Adaptive Slicing. The core function of module 1 is deriving more informative slice matrices from raw matrix, which mainly depends upon the design of increment index *SCRIstd*.

The slicing procedure is designed as the threshold operation:

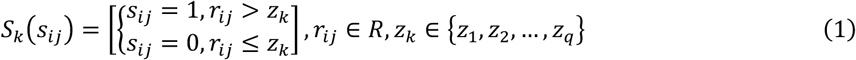

where R is the raw matrix of gene expression counts that has been normalized and transformed to proper range (27, 37). *S*_*k*_ is the *kth* slice matrix of the raw scRNA-seq data, which keeps the structure of differential expression at value *z*_*k*_ with binary manner. {*z*_1_, *z*_2_, …, *z*_*q*_} is a set of slice points between the minimum and maximum values of *R*, which are in ascending order for slicing procedure. This slicing procedure is used to extract necessary classification features from scRNA-seq data for machine learning.

By analyzing the histogram distribution of raw gene expression matrix, we can obtain the following results: (i) Different slice matrices at different slice points are derived, while these matrices keep different information of differential expression for cells. (ii) Informative slice points are selected. To quantify the information of switching in expression states, we defined the row increment index named *SCRIstd* as

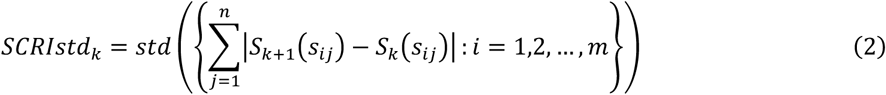

where *m* represents the maximum number of rows, and *n* represents the maximum number of columns. The term “*std*” is used here to refer to the process of calculating standard deviation of all row increments. If the calculated increment index *SCRIstd*_*k*+1_ > *SCRIstd*_*k*+2_ from slice points *z*_*k*_, *z*_*k*+1_, *z*_*k*+2_, which indicates that the slice matrix *S*_*k*+1_ contains more expression state switching increment information than *S*_*k*+2_. We calculate all the increment index *SCRIstd* between slice matrices and arrange them in descending order, then select the slice points corresponding to the first *p* increment indices to obtain the candidate set of slice points {*z*_(1)_, *z*_(2)_, …, *z*_(*p*)_}. Note that in scASK, we added 0 as the first slice point for the technical requirements, and the actual set of candidate slice points is {0, *z*_(1)_, *z*_(2)_,…, *z*_(*p*)_}.

### Module 2: Meta Classifiers Selecting (MCS)

The kNN classifier is a commonly used supervised machine learning algorithm, it is particularly well suited for multi-modal classes as its classification decision is based on a small neighbourhood of similar objects. As such, even if the target class consists of cells whose types have different characteristics for gene expression patterns, it can still lead to good classification accuracy (38).

scASK adopts Pearson’s correlation coefficient, Jaccard similarity and Cosine similarity as the default distance measures, where these three approaches could keep both diversity and complementarity in extracting the classification features from scRNA-seq data. Specifically, Pearson’s correlation coefficient measures the increasing and decreasing trend of gene expression data between the cells, Jaccard similarity measures the degree of overlap between intercellular binary expression patterns, and Cosine similarity calculates the vector angle between the cells using gene expression data as features. The exact formula definition of these three distance measures for scRNA-seq data can be found in Supplementary Materials Formulas S1 to S3.

As we know, kNN with three distance measures can produce 3 (*p* + 1) classifiers on (*p* + 1) slice matrices. Not all of trained classifiers could carry out adequate classification, some of classifiers may be strong, others may be weak. A common practice is to calculate the accuracy on training data and testing data for every classifier, then compare and select the more competitive classifiers (called meta classifiers). The core function and basic principle of module 2 are shown in Figure 3.

**Figure 3.**
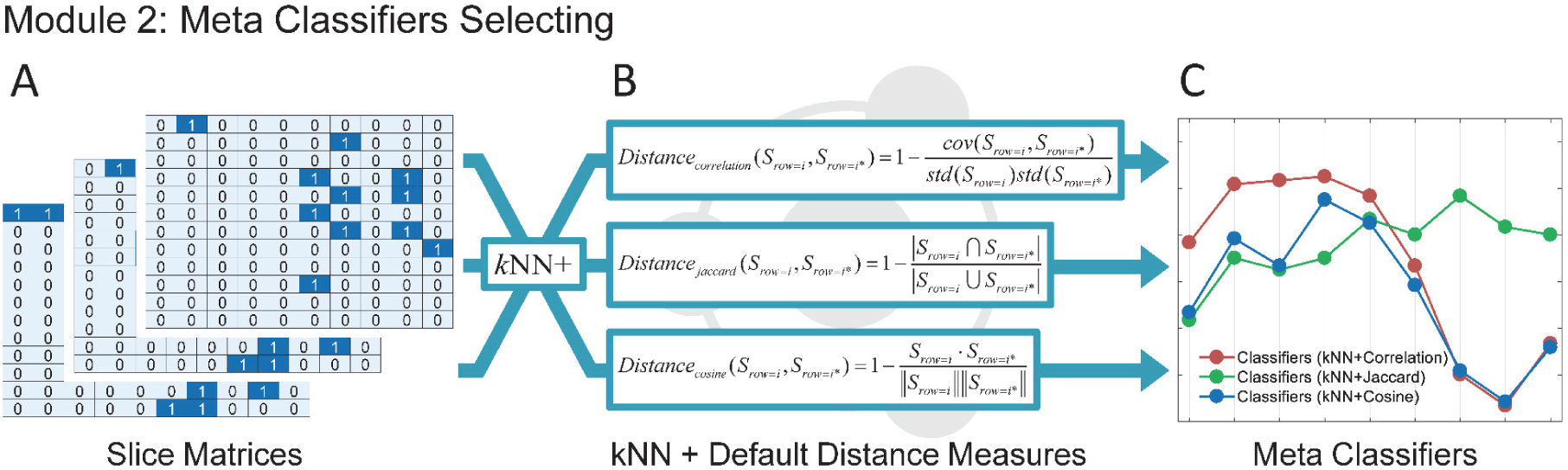
Module 2 of the scASK: Meta Classifiers Selecting. (**A**) The series of binary slice matrices. (**B**) scASK adopts Pearson’s correlation coefficient, Jaccard similarity and Cosine similarity as the default distance measures, where these three approaches could keep both diversity and complementarity in extracting the classification features from scRNA-seq data. (**C**) Three types of meta classifiers could be obtained on every slice point (training on every slice matrix). The core function of module 2 is selecting more appropriate fundamental algorithm and distance measures for binary slice matrices, which guarantees that all trained meta classifiers are competitive enough for subsequent ensemble classification.

### Module 3: Ensemble Mode Switching (EMS)

In many metrics used for evaluating the performance of a classification model, one important metric is the accuracy on testing data. In real scenarios, the accuracy on testing data is often inconsistent with the accuracy on training data, the essential reason of this phenomenon in machine learning is a high bias problem or a high variance problem, i.e., an under-fitting problem or an over-fitting problem (39). Both of problems could cause the significant inconsistency. Therefore, in scASK, we propose a weighted accuracy named *SMARwit* to evaluate and select out the meta classifiers with better performance, which is designed as follows:

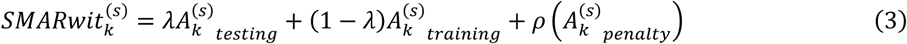

where *λ* is a predefined weight coefficient,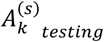 represents the accuracy of the *sth* meta classifier on testing data from the *kth* slice matrix, and 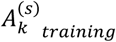 represents the accuracy of the *sth* meta classifier on training data from the *kth* slice matrix. The strength function *ρ* and the penalty term 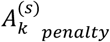 are designed as follows:

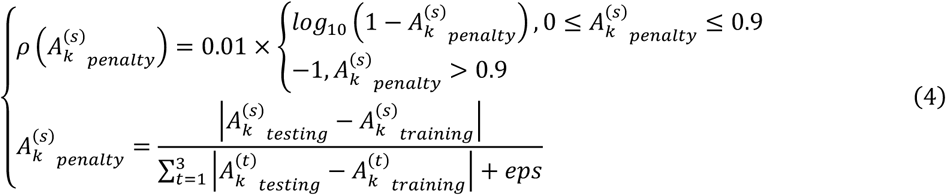

where 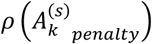 is a piecewise and decreasing function based on logarithm, which controls the strength of the penalty term affecting the weighted accuracy, and the penalty term 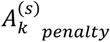 is used for measuring the consistency of 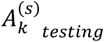 and 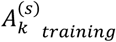, where the differences between testing accuracy and training accuracy for classifiers on each slice matrix are mapped to the proportional values in a same scale. It is important to note here that the term “*eps*” represents a floating-point number (is 2.2204e-16 in Matlab) small enough to keep the denominator away from zero. Generally, the value of *SMARwit* is mainly adjusted by *λ* (set 0.8 as default), and is slightly adjusted by the 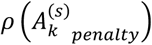 (range from −0.01 to 0). If the value of *SMARwit* is close to 1, it means that the meta classifier is more robust.

The generalization ability, which characterizes how well the results learned from a given training dataset can be applied to a new dataset, is the most central concept in machine learning. Researchers have devoted tremendous efforts to the pursuit of techniques that could lead to a learning system with a strong generalization ability. One of the most successful paradigms is ensemble learning (40). In scASK, we developed a new switching strategy especially for slice matrices, which is distinct from the traditional strategies for ensemble classification (i.e., bagging, boosting and stacking).

Now we can associate the steps established previously within a complete process. Suppose that a set of (*p* + 1) candidate slice points was selected out by adaptive slice algorithm in module 1, then (*p* + 1) slice matrices would be obtained accordingly as the training dataset. Three types of meta classifiers were constructed in module 2, would totally produce 3(*p* + 1) classifiers on all slice matrices. Finally, the last step of scASK is to decide how the candidate classifiers can be integrated in the ensemble.

In order to simplify expressions, we first define a special function named sort which sorts the given accuracy set in descending order and selects out the element of specified order number as

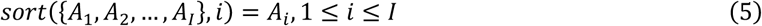

Then, two operating modes have been developed respectively. One mode named *SMESrws*, which is designed as follows:

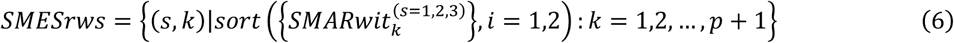

where (*s, k*) represents the index location of classifier (i.e., the *sth* meta classifier from the *kth* slice matrix) that satisfies the given condition, 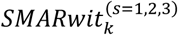 represents the set of every weighted accuracy from the *kth* slice matrix. After that the index locations of optimal and suboptimal classifiers are derived by the aid of the *sort* function. The procedure of *SMESrws* arranges classifiers on each slice matrix, which generates the index table of meta classifiers for the final ensemble classifier.

Another mode named *SMESabs*, which is designed as follows:

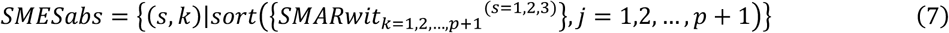

where (*s, k*) represents the index location of classifier (i.e., the *sth* meta classifier from the *kth* slice matrix) that satisfies the given condition, *SMARwit*_*k*=1,2, …,*p*+1_ ^(*s*=1,2, 3)^ represents the set of every weighted accuracy from all slice matrices. After that the index locations of the top (*p* + 1) best classifiers are derived by the aid of the *sort* function. The procedure of *SMESabs* arranges classifiers on all slice matrices, which generates the index table of meta classifiers for the final ensemble classifier.

To summarize, *SMESrws* selects out two meta classifiers from each slice matrix for the final ensemble with the simple voting scheme. The process of the selection performed by *SMESrws* is called “*railway switching*” (RWS). *SMESabs* selects out (*p* + 1) meta classifiers from all slice matrices for the final ensemble with the simple voting scheme. The process of the selection performed by *SMESabs* is called “*absolute best switching*” (ABS). Briefly, the RWS mode is a local optimal solution, while the ABS mode is a global optimal solution. For better accuracy and robustness, scASK switches the ensemble strategy between two modes manually according to the practice and experience. The core function and basic principle of module 3 are shown in Figure 4.

**Figure 4.**
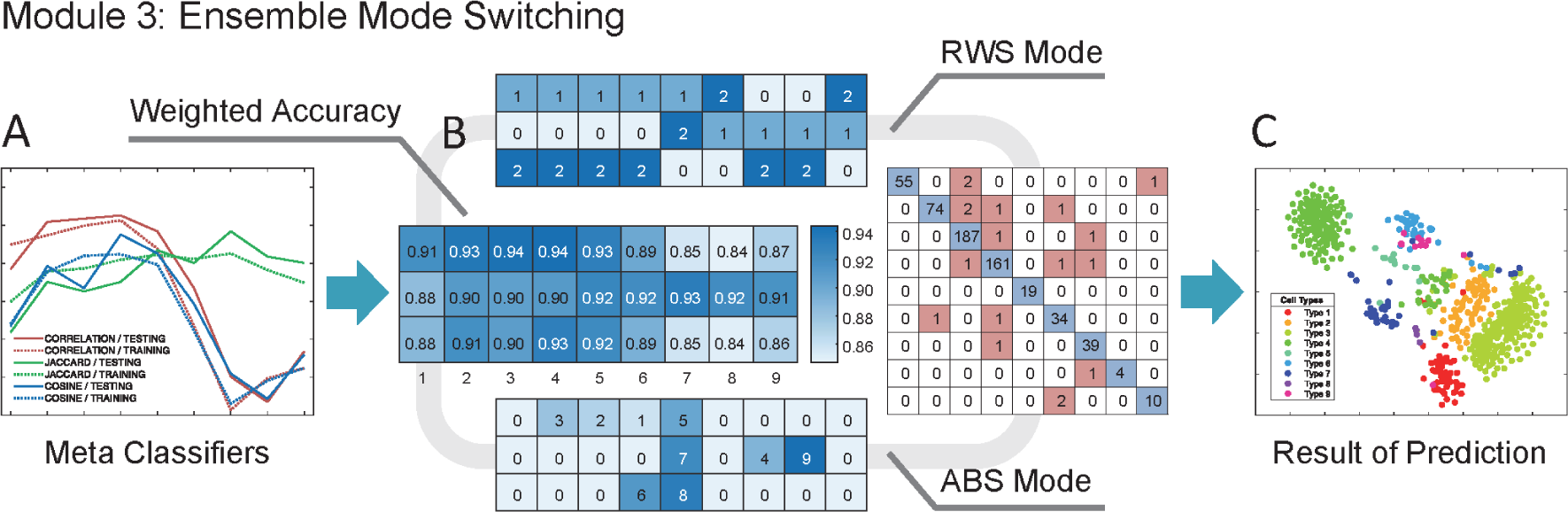
Module 3 of the scASK: Ensemble Mode Switching. (**A**) The accuracy of meta classifiers is calculated on every slice point. The solid line represents accuracy on testing data, and the dotted line represents accuracy on training data. Different colours represent different distance measures adopted by kNN. (**B**) scASK sorts all of meta classifiers with the weighted accuracy (implementing by the *SMARwit*), and then integrates them for final ensemble with two modes: the RWS mode and the ABS mode. We denote the dual operating mode on slice points for ensemble as “switching strategy”, which enhances the accuracy and the robustness of the final ensemble classifier. (**C**) scASK running with switching strategy (implementing by the *SMESrws* and *SMESabs*) can achieve better prediction for classifying cell types based on scRNA-seq data. The core function of module 3 is switching meta classifiers for ensemble according to the weighted accuracy.

## DATASETS AND RUNNING PROCEDURES

To verify the classification performance of scASK on real single-cell datasets, we used five single-cell datasets from Wang et al., 2017 (27) and eight single-cell datasets from Park and Zhao, 2018 (36). All these datasets were preprocessed and transformed into appropriate ranges (e.g.,*log*_1O_ (*x* + 1)). All cells have accurate and reliable cell-type labels with sample dimensions from 80 to 3005, the gene dimensions from 959 to 17772. The proportion of zeros is from 27.87% to 78.10%. These data have typical characteristics of high dimensionality and high sparsity. The summary of all datasets used in this paper is shown in Table 1.

**Table 1.** Summary of the thirteen real single-cell datasets

| Dataset | Number of Cells | Number of Genes | Populations | Sparsity |
| --- | --- | --- | --- | --- |
| Buettner | 182 | 8989 | 3 | 37.94% |
| Kolod | 704 | 10685 | 3 | 27.87% |
| Pollen | 249 | 14805 | 11 | 51.00% |
| Usoskin | 622 | 17772 | 4 | 78.10% |
| Zeisel | 3005 | 4412 | 9 | 46.01% |
| Buettner* | 182 | 8989 | 3 | 36.29% |
| Deng | 135 | 12548 | 7 | 31.85% |
| Ginhoux | 251 | 11834 | 3 | 66.54% |
| Pollen* | 249 | 14805 | 11 | 50.79% |
| Tasic | 1727 | 5832 | 48 | 32.71% |
| Ting | 114 | 14405 | 5 | 52.17% |
| Treutlin | 80 | 959 | 5 | 51.65% |
| Zeisel* | 3005 | 4412 | 48 | 46.01% |

Note: The superscript \* means the dataset from the same origin literature with different data preprocessing. Datasets 1 to 5 are available from the Supplementary data of Wang et al., 2017, and datasets 6 to 13 are available from the Supplementary data of Park and Zhao, 2018, where the descriptions for datasets with more details can be found respectively.

In machine learning, we always build a model based on the training set (a subset of a raw dataset), and evaluate the model based on the test set (the remaining subset of the raw dataset). Because the test set is completely independent of the training process, so the prediction accuracy on the test set is regarded as the most significant evaluation metric of the model. The cross validation is a popular resampling procedure which is also used to evaluate the performance of the model on new data. This procedure has a single parameter called *k*-fold that refers to the number of pieces that a given dataset is to be split into. There is no hard and fast rule for the choice of *k*: the larger value of *k* causes the less bias and more variance, while the smaller value of *k* yields more bias and less variance (39). It is noteworthy that, the procedure of cross validation is also adopted in scASK for evaluating the performance of meta classifiers during training process.

Next we present the running procedures of scASK for cell type classification on single-cell datasets, and explain some key parameters and give empirical values. The running procedures include six steps. On each step, one or more computing tasks are executed by running specific functions sequentially. The complete codes and output results for each single-cell dataset can be found in Supplementary Materials.

**Step 1**. Input the dataset and run the **slicematrix** function to initially determine the starting point, end point and interval of the slice by plotting the raw data histogram. The slice interval needs to cover all occurrences of “peaks” and “valleys” in the histogram. In theory, the smaller the slice interval is, the easier it is to capture the classification features hidden in the slice matrices. In practice, this value is determined by the equilibrium between storage cost and calculation duration. (the empirical value: *q*=30∼60).

**Step 2**. Run the **slicediffer** function to calculate the increment index *SCRIstd* between the slice matrices at all slicepoints, and select the first (*p* + 1) slice points to form the candidate set of slice points. (the empirical value: (*p* + 1) =6, 9, 15, 21, 36).

**Step 3**. Run the **slicemethod** function to train the kNN classifiers with Pearson’s correlation coefficient, Jaccard similarity and Cosine similarity on the (*p* + 1) slice matrices. The total number of 3 (*p* + 1) classifiers form the candidate set of meta classifiers. (the empirical value: *k*=1 or 5, distance weighted by “inverse”).

**Step 4**. Run the **sliceweight** function to calculate the accuracy of the test set and the training set for each classifier, then calculate the corresponding weighted accuracy according to the *SMARwit*.

**Step 5**. Run the **sliceswitch** function in *SMESrw*s and *SMESabs* modes respectively to implement the meta classifiers ensemble with the switching strategy. As a matter of experience, *SMESrws* is more conservative than *SMESabs* on the prediction.

**Step 6**. Run the **sliceprerws** function and the **slicepreabs** function respectively to verify the prediction accuracy of the test set of scASK under the two ensemble strategies. The classification results are given in the form of confusion matrix, and the cell-type label and support degree of each sample are given at the same time.

Finally, we provide the empirical parameters and the command-line format for scASK on Buettner dataset as a running example:

Demo: running procedures for scASK on Buettner dataset

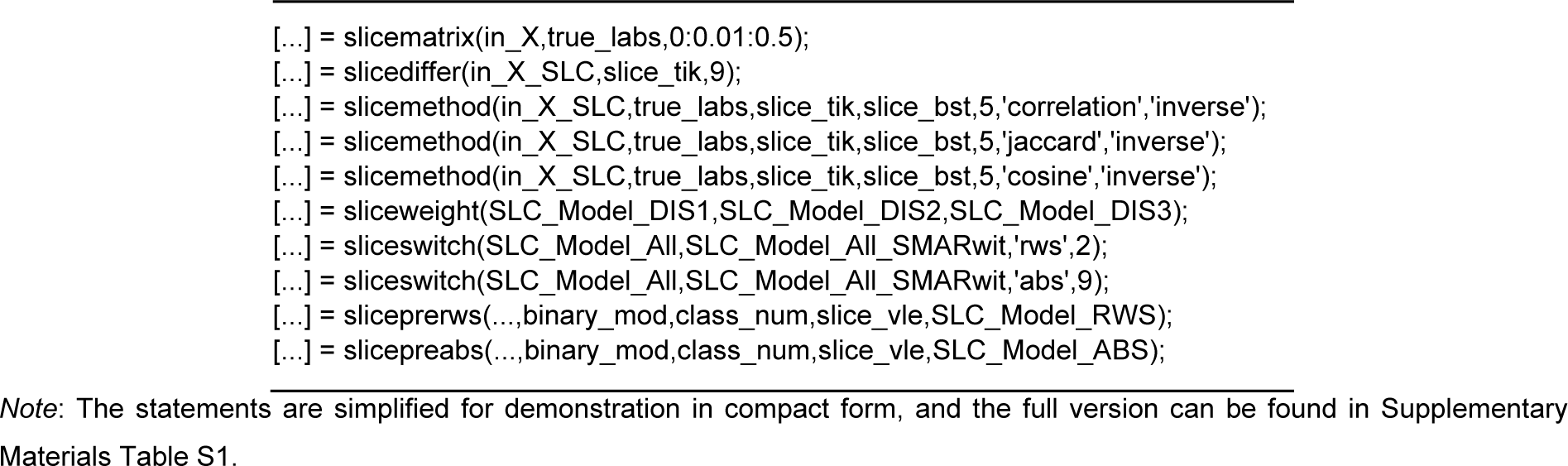

## RESULT

To evaluate the effectiveness and generality of our new ensemble framework for classifying cell types based on scRNA-seq data, we have implemented scASK on thirteen real single-cell datasets listed in Table 1. All of computational results and figures during the running process of scASK on every dataset have been collected in Supplementary Materials Tables S1 to S17 and Figures S1 to S187.

In this section, firstly we demonstrate the core functions of scASK on three real single-cell datasets: Buettner, Pollen and Zeisel. These datasets, from small to large scale, are typical kind of scRNA-seq data. Then we evaluate the performance of scASK by comparing with five baseline methods on all datasets listed in Table 1. The parameters for scASK on one specific dataset are dependent of data experiments. Therefore, the given empirical parameters are only for result reproducibility, and may change in different scenarios.

### Performance of scASK on three real datasets

To evaluate the performance of our proposed scASK, we first introduce three real single-cell datasets: Buettner, Pollen and Zeisel (36).

1. Buettner: In this dataset, the transcriptional profiles of 182 embryonic stem cells (ESCs) had been staged for cell-cycle phases (G1, S, and G2M) based on sorting of the Hoechst 33342-stained cell area of a flow cytometry (FACS) distribution (41). The cells were grouped into three stages of the cell cycle, and they were validated using the gold-standard Hoechst staining. scASK achieves the classification accuracy of 100% on Buettner dataset, and the key parameters and summary results are shown in Figure 5A.
2. Pollen: it consists of the transcriptional profiles of 249 single cells from 11 populations using microuidics, including neural cells and blood cells (42). The 11 clusters in the dataset were from different sources (CRL-2338, CRL-2339, K562, BJ, HL60, hiPSC, Keratinocyte, Fetal cortex (GW21+3), Fetal cortex (GW21), Fetal cortex(GW16), and NPC) that are expected to show robust differences in gene expression. scASK achieves the classification accuracy of 98% on Pollen dataset, and the key parameters and summary results are shown in Figure 5B.
3. Zeisel: it consists of the transcriptional profiles of 3005 cells from the mouse cortex and hippocampus. Zeisel et al., 2015 (30) found 9 or 48 molecularly distinct subclasses identified by hierarchical biclustering and validated by gene markers. scASK achieves the classification accuracy of 97% on Zeisel dataset (9 classes), and the key parameters and summary results are shown in Figure 5C.

### Comparisons of scASK with five baseline methods for scRNA-seq Data

To further demonstrate the effectiveness of the proposed scASK, we compare scASK with five representative classification algorithms including Complex Tree, Quadratic SVM, Medium KNN, Bagged Tree and Boosted Tree in terms of accuracy and robustness. All approaches are run on a laptop computer with Intel Core i5-6200U @ 2.30GHz and 8 GB of RAMs and five other algorithms are implemented by using their corresponding functions provided by Matlab 2017a.

**Figure 5.**
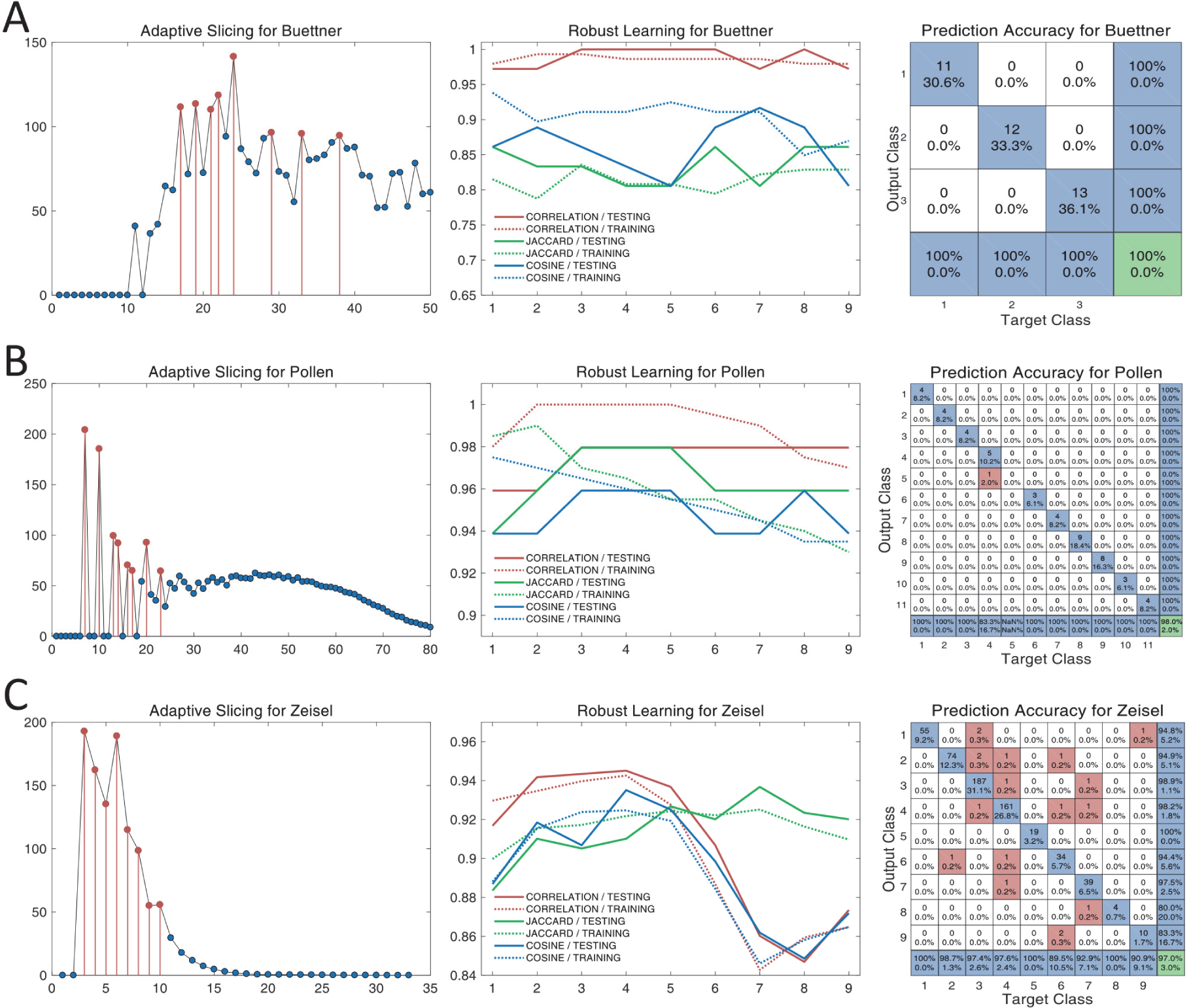
Performance of scASK on three real datasets Buettner, Pollen and Zeisel. (A) Result for Buettner dataset. 9 slice points (0.17,0.19,0.21,0.22,0.24,0.29,0.33, 0.38, and 0) were selected out with the aid of adaptive slicing procedure; 27 meta classifiers were constructed by kNN with 3 kinds of efficient distance measures (Correlation, Jaccard and Cosine); After ensemble of these meta classifiers, scASK (both of RWS mode and ABS mode) achieved the classification accuracy of 100% on the Buettner dataset (test set). (B) Result for Pollen dataset. 9 slice points (0.35,0.5,0.65,0.70,0.80,0.85,1,1.15, and 0) were selected out with the aid of adaptive slicing procedure; 27 meta classifiers were constructed by kNN with 3 kinds of efficient distance measures (Correlation, Jaccard and Cosine); After ensemble of these meta classifiers, scASK (both of RWS mode and ABS mode) achieved the classification accuracy of 98% on the Pollen dataset (test set). (C) Result for Zeisel dataset (9 classes). 9 slice points (0.9,1.2,1.5,1.8,2.1,2.4,2.7,3, and 0) were selected out with the aid of adaptive slicing procedure; 27 meta classifiers were constructed by kNN with 3 kinds of efficient distance measures (Correlation, Jaccard and Cosine); After ensemble of these meta classifiers, scASK (RWS mode) achieved the classification accuracy of 97% on the Zeisel dataset (test set).

On the one hand, we compare the prediction accuracy of all methods. The prediction accuracy on test sets is not the only metric for evaluating the performance of a classification model, but it is most important for measuring the classification rate which ranges from 0 to 100%, and a high value means that the algorithm accurately discovers cell types from new samples (43). For classifying cell types based on scRNA-seq data, three competing algorithms (Bagged Tree, Quadratic SVM and scASK) have been selected out because of their outstanding performance. From Figure 6 we can observe that scASK achieves the best prediction accuracy on all datasets.

**Figure 6.**
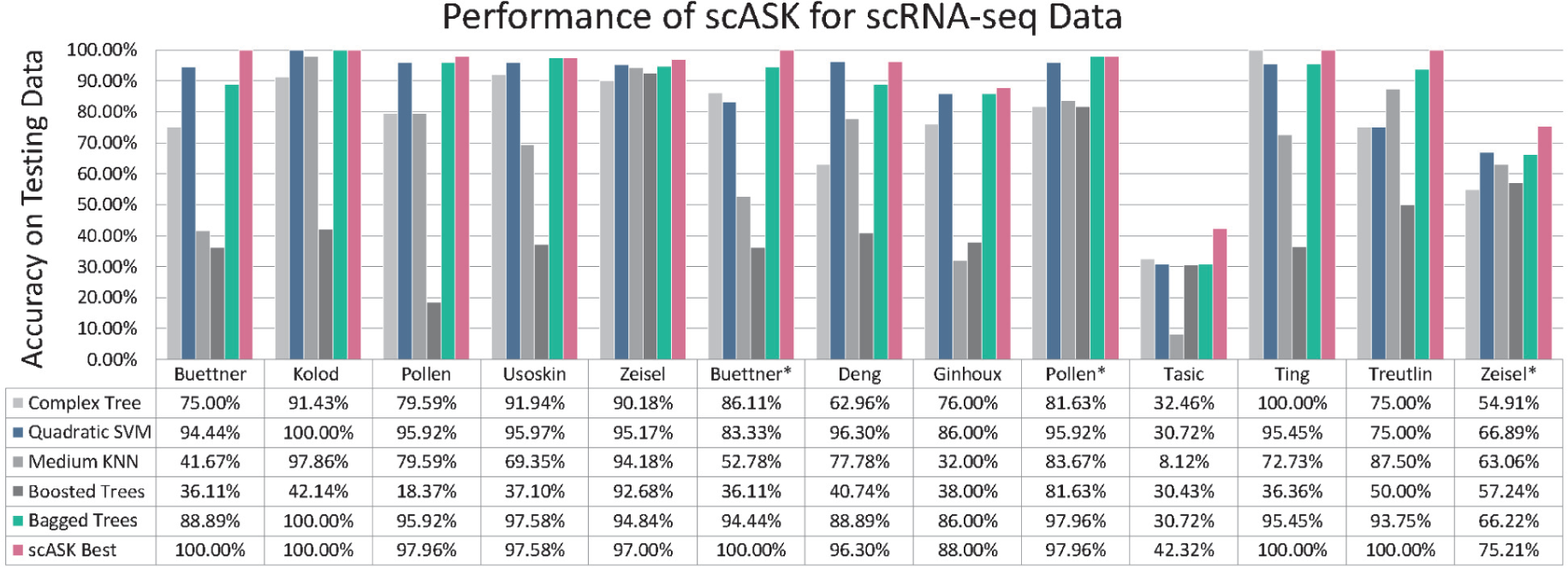
Comparisons of scASK in accuracy with five other baseline methods for scRNA-seq Data. The term “scASK Best” represents the optimal mode between *SMESrws* and *SMESabs*. scASK keeps the highest prediction accuracy on thirteen real scRNA-seq datasets.

On the other hand, we evaluate the robustness of these algorithms on real single-cell datasets. One important characteristic of scRNA-seq data is the “dropout” phenomenon where a gene is observed in one cell but undetected in another cell (44). Based on this consideration, we simulate missing values by randomly replacing the non-zero elements of the original data with zeroes for a certain proportion (10%) on thirteen real scRNA-seq datasets. The variations of prediction accuracy before and after the simulations are used for measuring the robustness of the algorithms. As shown in Figure 7, scASK keeps the best robustness under the strong disturbances during the simulations across all datasets.

**Figure 7.**
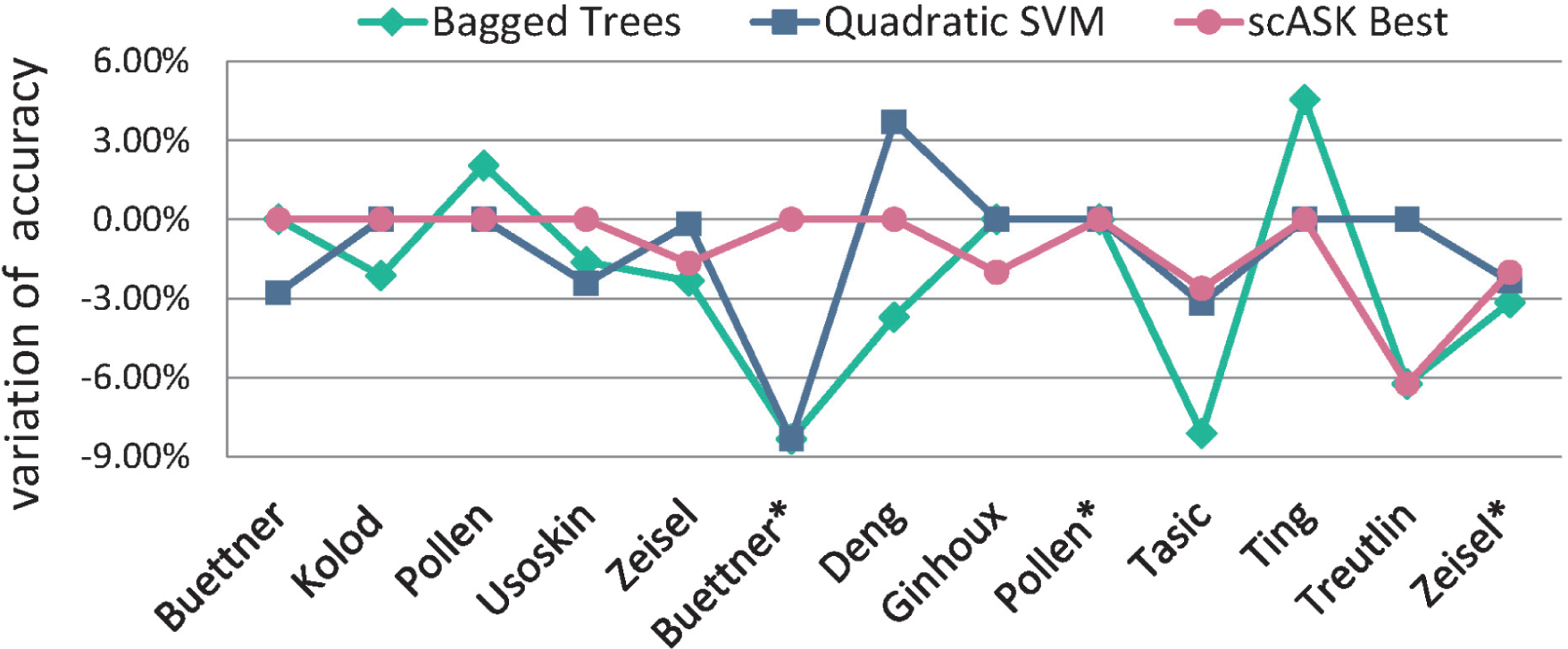
Comparisons of scASK in robustness with two other methods Bagged Trees and Quadratic SVM. The green, blue and red colours are used to represent Bagged Trees, Quadratic SVM and the scASK, respectively. The smaller variation of accuracy on datasets indicate that the method has stronger robustness.

### Software package

We develop a graphical user interface (GUI) software package for implementing scASK in Matlab 2019a (shown in Figure 8), which can significantly simplify the analysis and the process of parameter selection. The software is publicly available at https://github.com/liubo2358/scASKapp. The description documentation and quick tutorials of this software package are presented online in the same repository.

**Figure 8.**
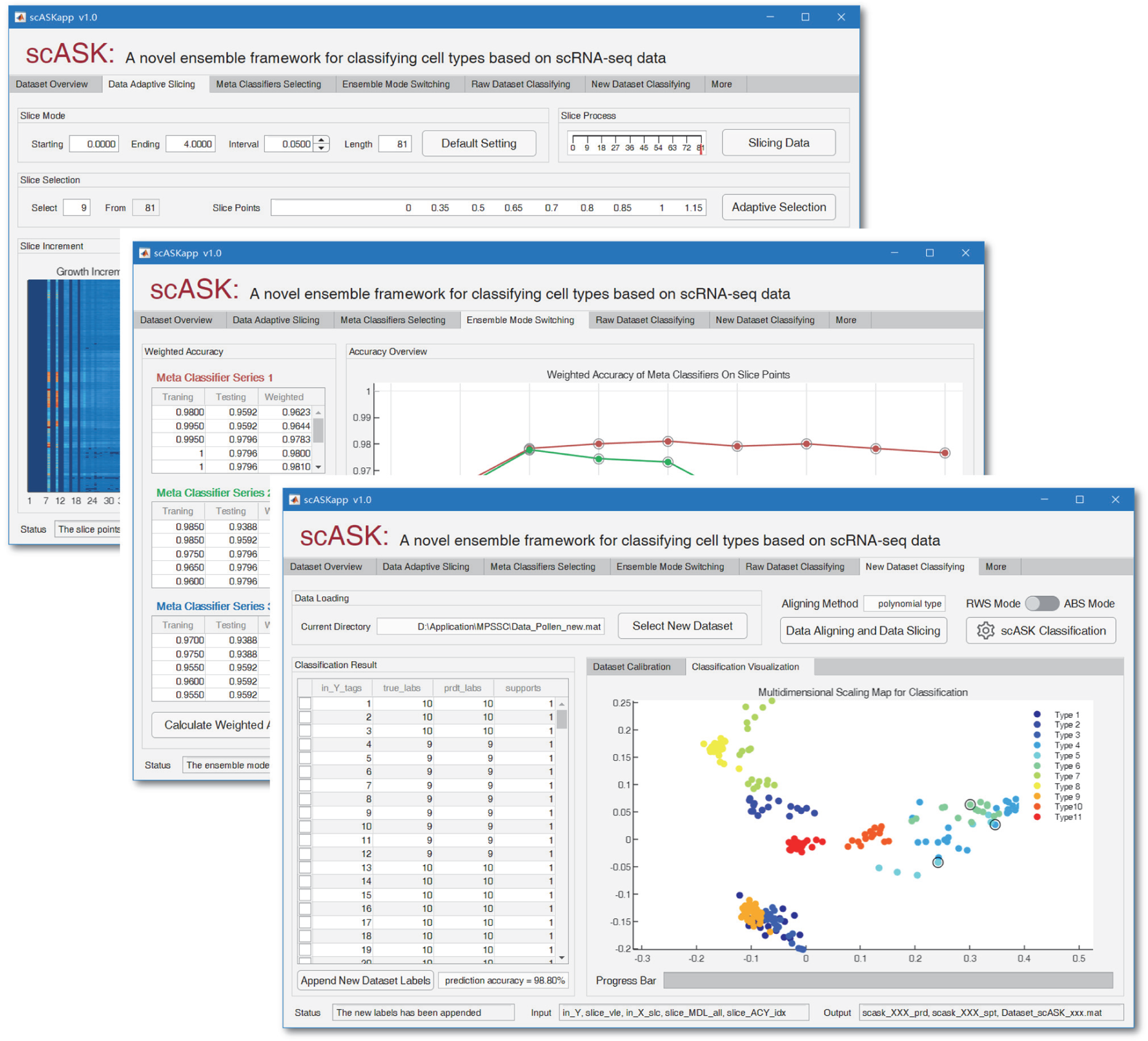
Screenshots of scASKapp. Shown here are the Data Adaptive Slicing tab, Ensemble Mode Switching tab and New Dataset Classifying tab.

## DISCUSSION

Generally, if we regard the Human Cell Atlas as the dictionary of information on all cell types in human body, scASK provides a fast, accurate and reliable method for querying this dictionary. The main goal of scASK is to address the challenges of high dimensionality and high variability in single-cell data. Although, there is no universally best classification algorithm for all datasets, we emphasize that scASK has the potential to be extended to the most matching classification algorithm for classifying cell types based on scRNA-seq data, and even for classifying cancer types based on gene expression data or DNA methylation data. In this section, we discuss some key details and further functions of scASK.

First of all, the slicing procedure used by scASK is not intended to replace PCA (or t-SNE), but rather provides a choice that is parallel to existing mainstream dimension reduction methods for high dimensional single-cell data. On the one hand, the exact SVD of very large matrix is still a difficult computational problem, essentially because matrix decomposition is a series of related operations which are hard to parallelize. Since the slicing procedure is a linearization process, the classifier training based on each slice matrix is completely independent, so it can be performed without interference at the same time. This feature makes scASK more easily for parallelization. On the other hand, the matrix decomposition (such as PCA) of the existing data during the data preprocessing step may cause some troubles for classification on future data. It is unrealistic to require subsequent samples to participate in the matrix decomposition of initial samples in the training phase, because the matrix decomposition is equivalent to the feature reconstruction. The lack of the process could finally lead to the failure of classification of cell types across datasets. For example, the Pollen* dataset cannot be directly imported into the classifiers trained from the Pollen dataset with PCA for classification, although they are essentially the same data with different preprocessing. If we align the Pollen* dataset with the Pollen dataset after appropriate scale-transformation, the classifiers trained from the Pollen dataset by scASK can directly identify cell types for Pollen* dataset. Actually, the data alignment across datasets is an important topic of our future work (for some basic implementation, see the tutorials of scASKapp).

Secondly, scASK does not exclude PCA, in fact, they can cooperate very well. Let’s take the Macosk dataset as an example, the dimension of raw data is very large (6418×12822). In the process of implementing scASK, the slicing procedure takes up so much memory that exceeds the capacity of our laptop. Back to square one, once we reduce the dimension to 6148×100 by the aid of PCA, and append a non-negative processing of matrix elements, then scASK can also achieve a good enough classification accuracy of 90.34% on test set (for running codes and output results, see Supplementary Materials Table S11 and Figure S111 to S121). Next, it is worth mentioning that Pearson’s correlation coefficient, Jaccard similarity and Cosine similarity as the default distance measures of scASK is recommended but not necessary. In fact, we find that the accuracy of scASK using only the Pearson’s correlation coefficient is obviously higher in some datasets (such as Usoskin data and Tasic data, see Supplementary Materials Table S5 and Figure S45 to S55, Table S14 and Figure S144 to S154). It is indicated that Pearson’s correlation coefficient as distance measure always performs well in scASK for classifying cell types based on scRNA-seq data.

Finally, the significant advantages of scASK in implementing cell type classification for single-cell data is its high accuracy and high robustness. The adaptive slicing and switching strategy all play crucial role in scASK, the former can be regarded as a new structural reduction procedure which is distinct from the popular procedure for high-dimensional data (e.g., PCA, t-SNE and NMF), and the latter can be regarded as a new ensemble strategy which is distinct from the traditional strategies for classification (i.e., bagging, boosting and stacking). scASK runs on slice matrices with the switching strategy, which not only helps confirm the results, but also enhances the reliability of the classification. Both of technologies above can be extended to a wider range beyond this work, including medical diagnosis, data analysis, machine learning and so on.

## Supporting information

Supplementary Demonstration

## ACKNOWLEDGEMENTS

We thank xxx and xxx for helpful discussions of this manuscript.

