## Supplementary Materials for "scASK: A novel ensemble framework for classifying cell types based on single-cell RNA-seq data"

### This PDF file includes:

#### 1. Supplementary Formulas

Correlation coefficient, Jacquard and Cosine distance formulas (Formulas S1 to S3)

#### 2. Supplementary Parameters and Results

The running parameters for fourteen scRNA-seq datasets (Tables S1 to S17)

The output figures for fourteen scRNA-seq datasets (Figures S1 to S187)

### 1. Supplementary Formulas

1.1 Correlation coefficient measures the distance between cells via

$$Distance_{correlation}(S_{row=i}, S_{row=i*}) = 1 - \frac{cov(S_{row=i}, S_{row=i*})}{std(S_{row=i})std(S_{row=i*})} = 1 - \frac{\sum\left(S_{row=i} - \overline{S_{row=i}}\right)\left(S_{row=i*} - \overline{S_{row=i*}}\right)}{\sqrt{\sum_{j=1}^{n}(S_{row=i,col=j} - \overline{S_{row=i}})^2}\sqrt{\sum_{j=1}^{n}(S_{row=i*,col=j} - \overline{S_{row=i*}})^2}}, \quad \text{(S1)}$$

where $S_{row=i}$ and $S_{row=i*}$ are specific rows of slice matrix $S$, which represent two distinct cell samples. The quotient of $cov(S_{row=i}, S_{row=i*})$ divided by $std(S_{row=i})std(S_{row=i*})$ represents the consistency for variation trend of gene expression data between cells. The better consistency indicates the higher degree of similarity.

1.2 Jacquard similarity measures the distance between cells via

$$Distance_{jaccard}(S_{row=i}, S_{row=i*}) = 1 - \frac{|S_{row=i} \cap S_{row=i*}|}{|S_{row=i} \cup S_{row=i*}|} = \frac{\sum_{j=1}^{n}(S_{row=i,col=j} - S_{row=i*,col=j})^2}{\sum_{j=1}^{n}\left((S_{row=i,col=j})^2 + (S_{row=i*,col=j})^2\right) - \sum_{j=1}^{n}(S_{row=i,col=j}S_{row=i*,col=j})}, \quad \text{(S2)}$$

where the quotient of $S_{row=i} \cap S_{row=i*}$ divided by $S_{row=i} \cup S_{row=i*}$ represents the overlap degree of binary expression patterns between cells. The more overlap indicates the higher degree of similarity.

1.3 Cosine similarity measures the distance between cells via

$$Distance_{cosine}(S_{row=i}, S_{row=i*}) = 1 - \frac{S_{row=i} \cdot S_{row=i*}}{\|S_{row=i}\|\|S_{row=i*}\|} = 1 - \frac{\sum S_{row=i}S_{row=i*}}{\sqrt{\sum_{j=1}^{n}(S_{row=i,col=j})^2}\sqrt{\sum_{j=1}^{n}(S_{row=i*,col=j})^2}}, \quad \text{(S3)}$$

where the quotient of $S_{row=i} \cdot S_{row=i*}$ divided by $\|S_{row=i}\|\|S_{row=i*}\|$ represents cosine value of the angle between vectors(cells characterized by gene expression data). The closer cosine value is to 1, the more similar between cells for comparison.

### 2. Supplementary Parameters and Results

#### 2.1 Guide for running scASK from the command line:

Step 1: Unzip all of the files in scASKcmd.rar to the initial working folder of Matlab

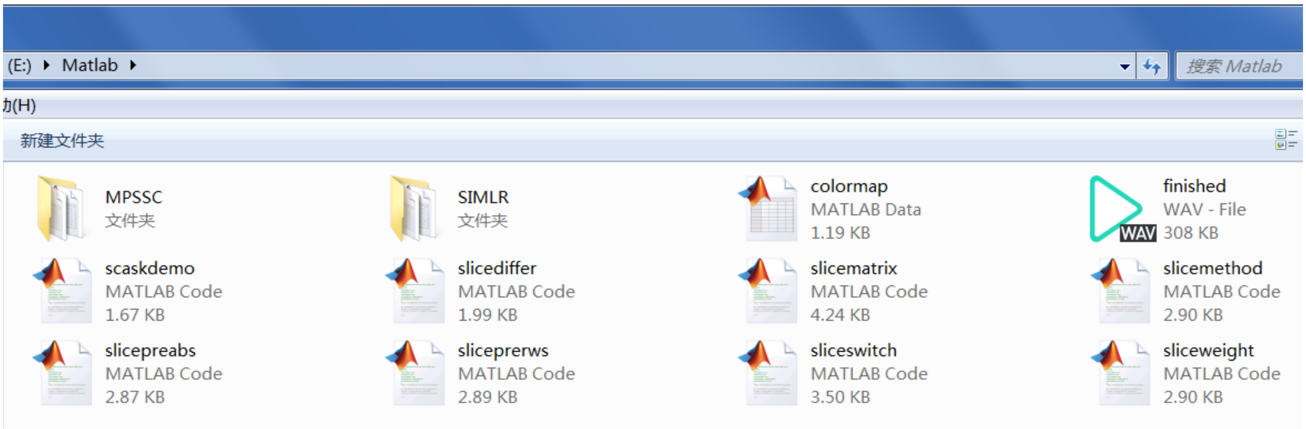


Step 2: Download example datasets, then copy the mat-files to initial working folder as well

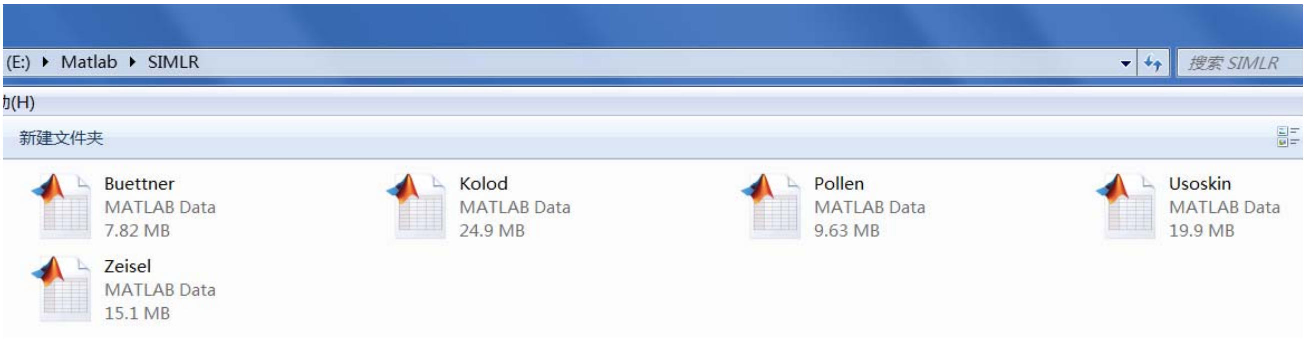


Step 3: Enter the following statements in Command Window, scASK will be launching soon

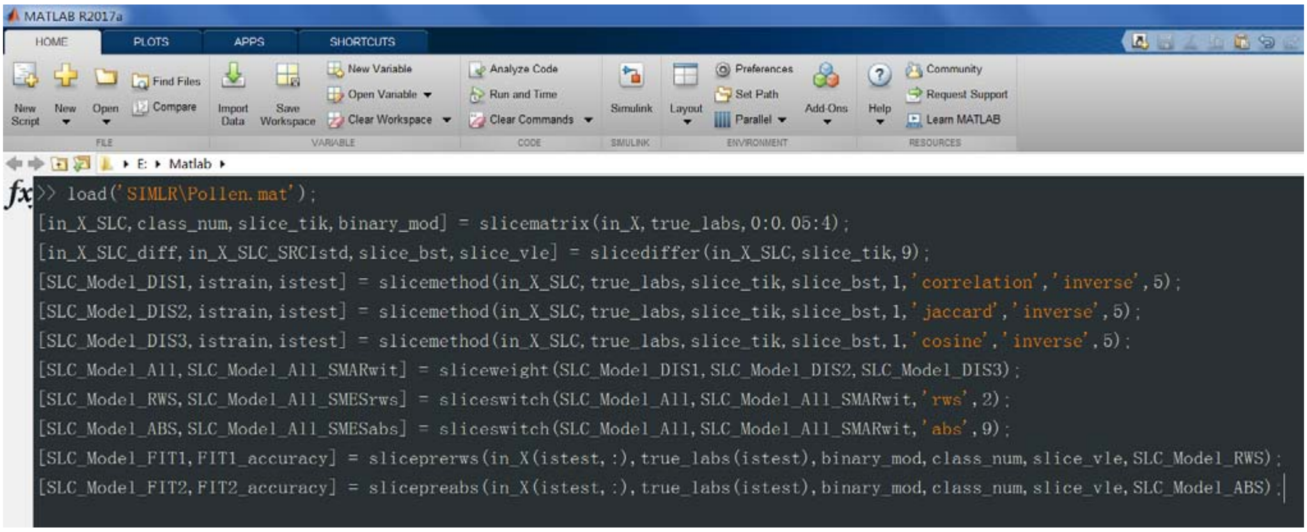


#### 2.2 The download URLs for example datasets:

SIMLR Dataset https://github.com/BatzoglouLabSU/SIMLR/tree/SIMLR/MATLAB/data
MPSSC Dataset https://github.com/ishspsy/project/tree/master/MPSSC/Data

#### 2.3 The running parameters and output figures for *Buettner* dataset

Table S1: The running parameters for *Buettner* dataset.

```
load('SIMLR\Buettner.mat');
[in_X_SLC,class_num,slice_tik,binary_mod] = slicematrix(in_X,true_labs,0:0.01:0.5);
[in_X_SLC_diff,in_X_SLC_SRCIstd,slice_bst,slice_vle] = slicedaffer(in_X_SLC,slice_tik,9);
[SLC_Model_DIS1,istrain,istest] = slicemethod(in_X_SLC,true_labs,slice_tik,slice_bst,5,'correlation','inverse',5);
[SLC_Model_DIS2,istrain,istest] = slicemethod(in_X_SLC,true_labs,slice_tik,slice_bst,5,'jaccard','inverse',5);
[SLC_Model_DIS3,istrain,istest] = slicemethod(in_X_SLC,true_labs,slice_tik,slice_bst,5,'cosine','inverse',5);
[SLC_Model_All,SLC_Model_All_SMARwit] = sliceweight(SLC_Model_DIS1,SLC_Model_DIS2,SLC_Model_DIS3);
[SLC_Model_RWS,SLC_Model_All_SMESrws] = sliceswitch(SLC_Model_All,SLC_Model_All_SMARwit,'rws',2);
[SLC_Model_ABS,SLC_Model_All_SMESabs] = sliceswitch(SLC_Model_All,SLC_Model_All_SMARwit,'abs',9);
[SLC_Model_FIT1,FIT1_accuracy] = sliceprerws(in_X(istest,:),true_labs(istest),binary_mod,class_num,slice_vle,SLC_Model_RWS);
[SLC_Model_FIT2,FIT2_accuracy] = slicepreabs(in_X(istest,:),true_labs(istest),binary_mod,class_num,slice_vle, SLC_Model_ABS);
```

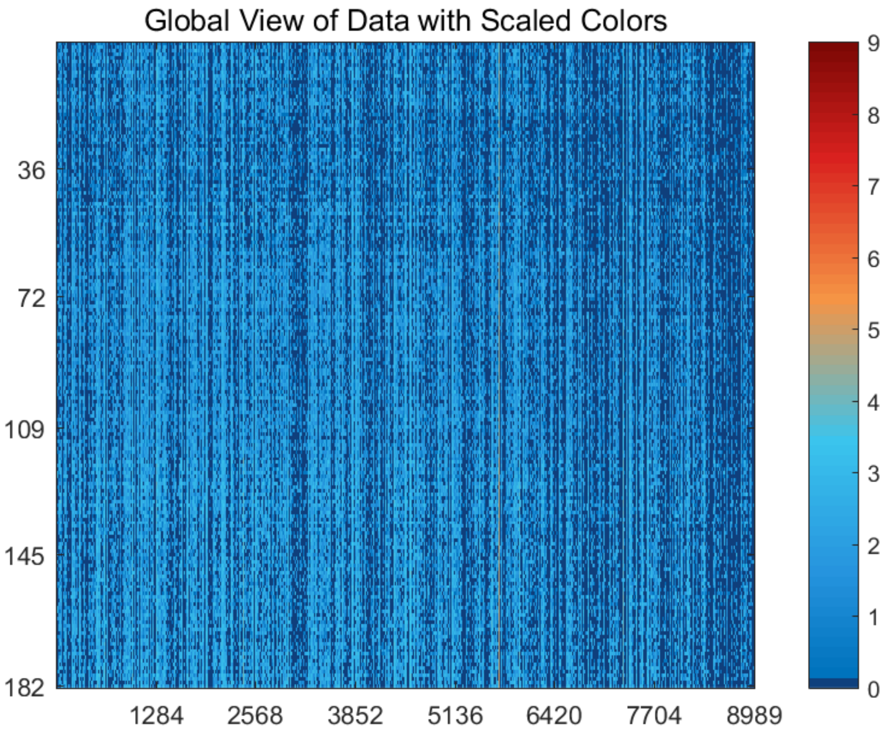


Figure S1: The global view of *Buettner* dataset with scaled colors.

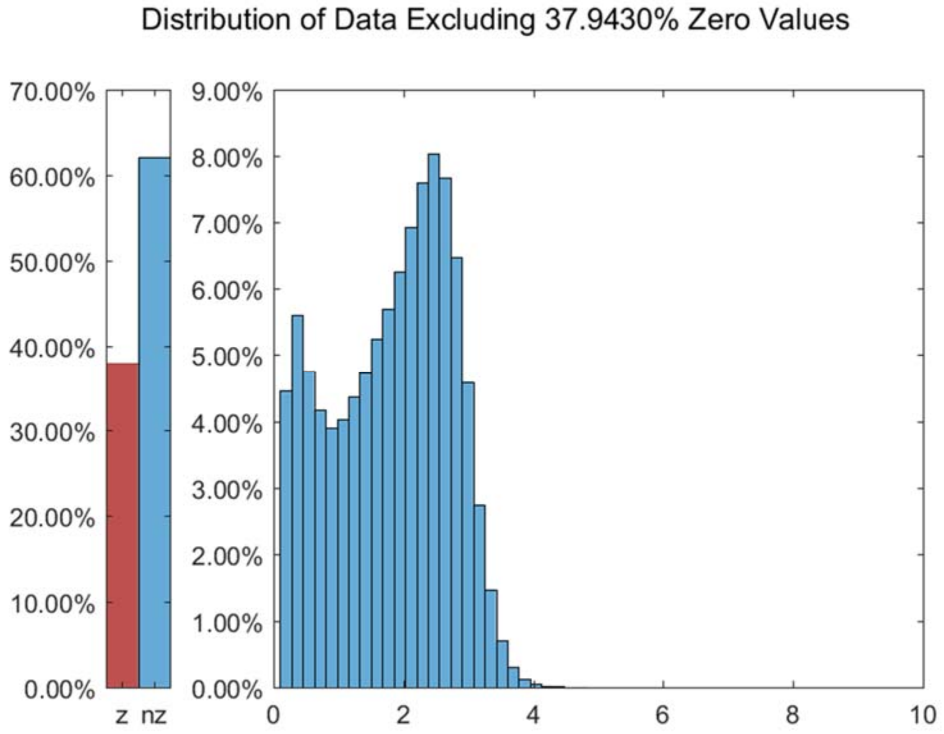


Figure S2: The distribution of *Buettner* dataset excluding all zero values.

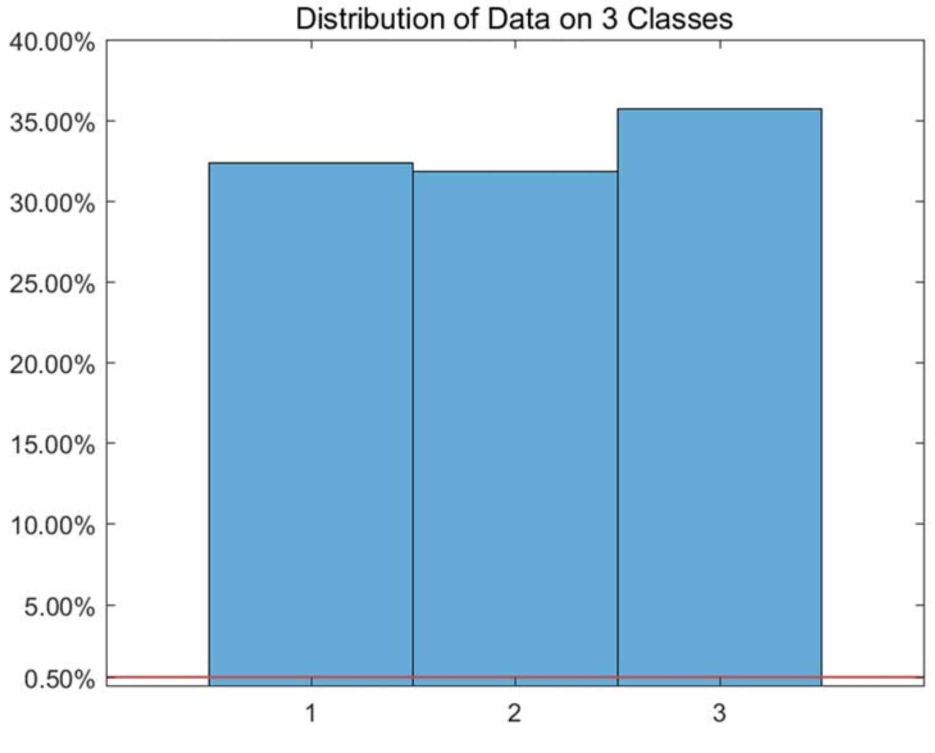


Figure S3: The distribution of *Buettner* dataset on every classes.

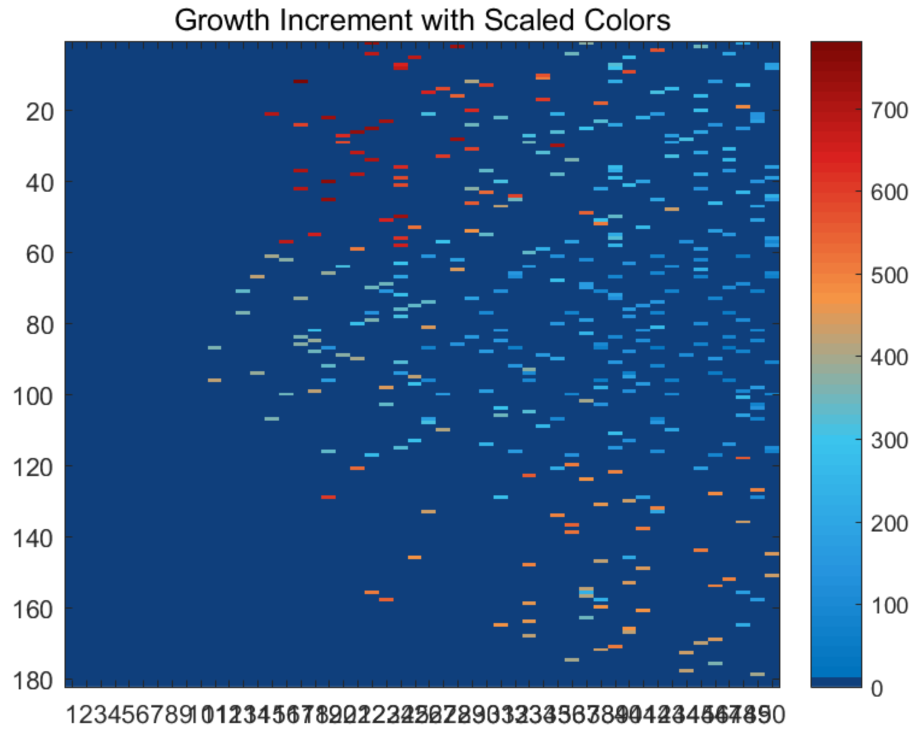


Figure S4: The growth increment with scaled colors for *Buettner* dataset.

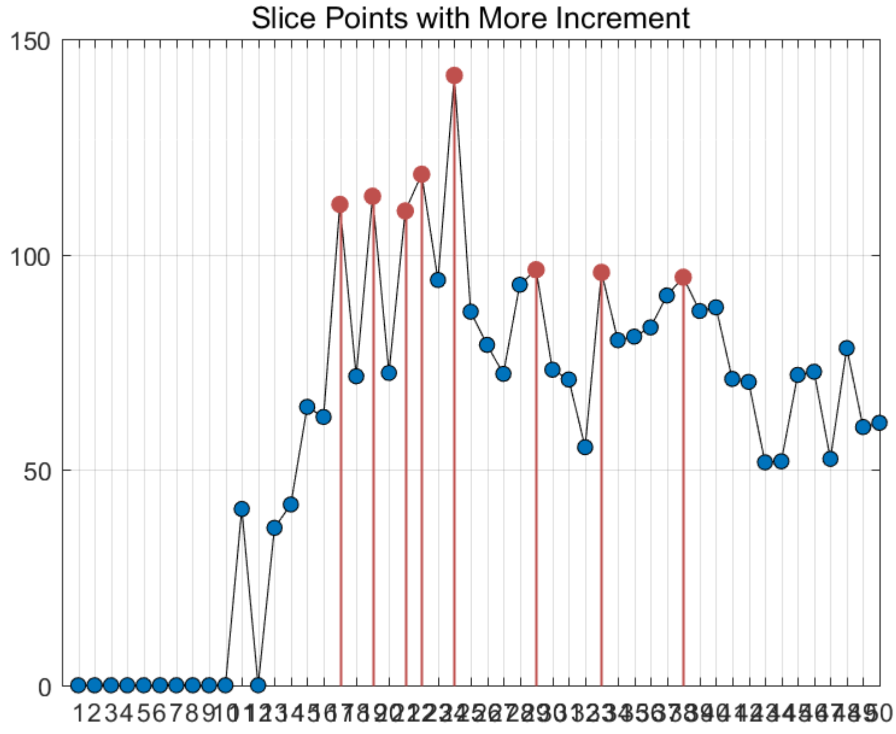


Figure S5: The slice points with more increment for *Buettner* dataset.

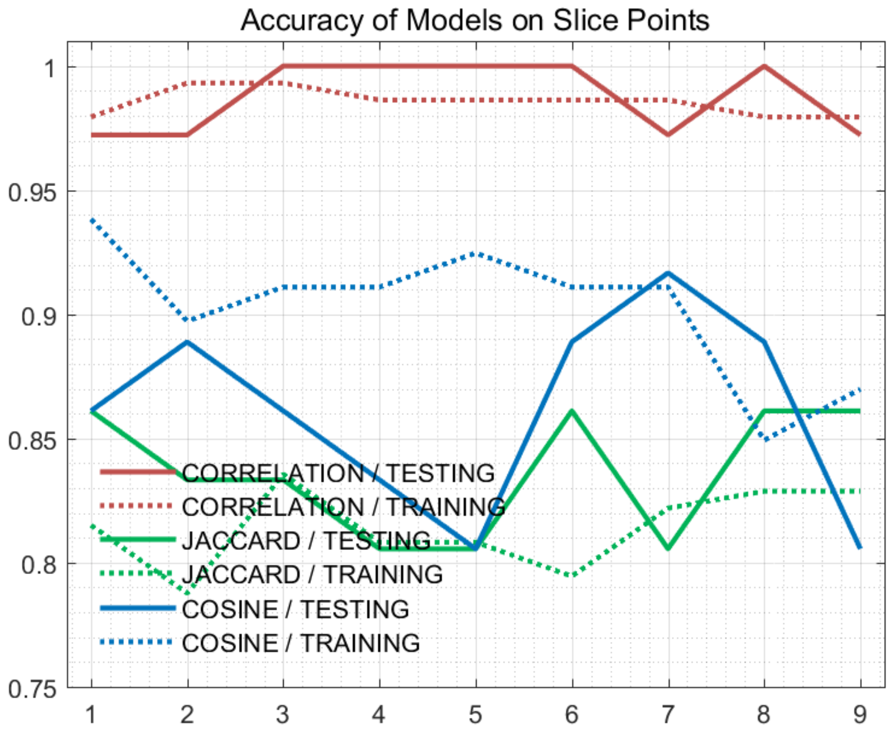


Figure S6: The accuracy of models on every slice points for *Buettner* dataset.

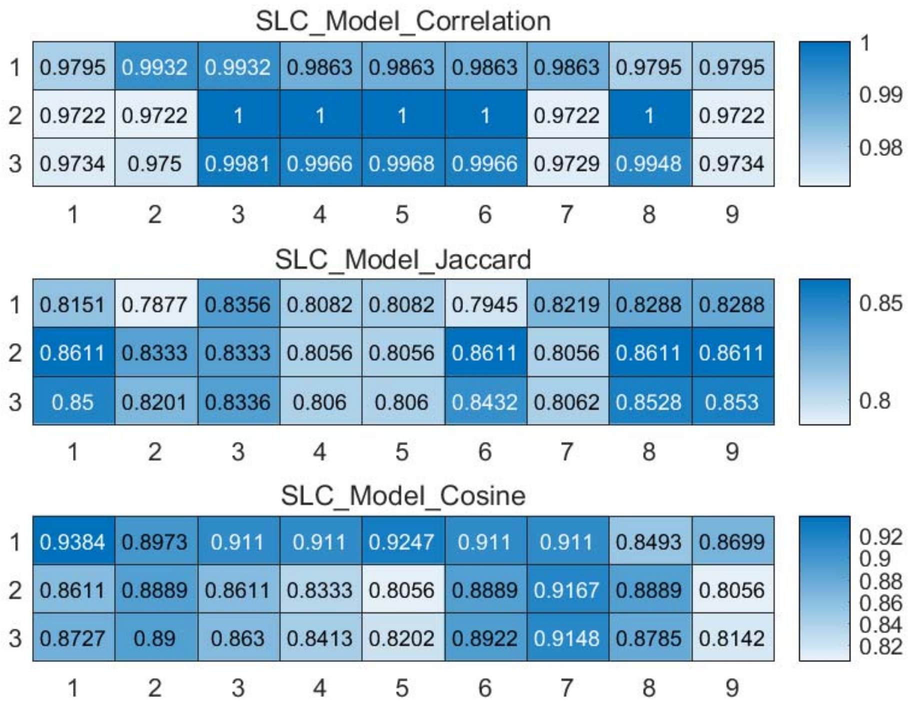


Figure S7: The weighted accuracy on every slice points for *Buettner* dataset.

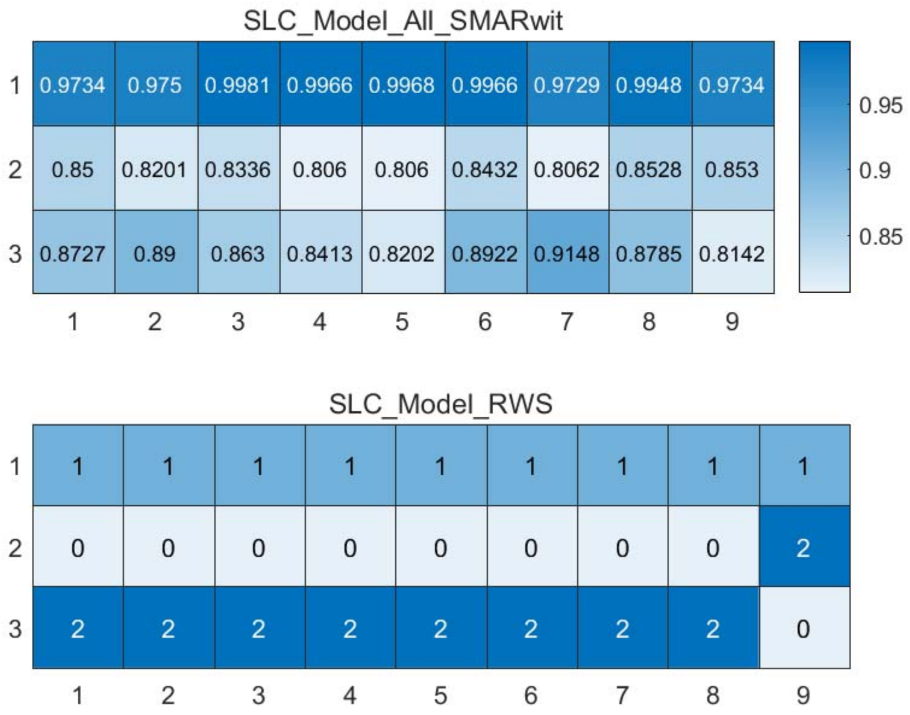


Figure S8: The RWS mode of meta classifiers for *Buettner* dataset.

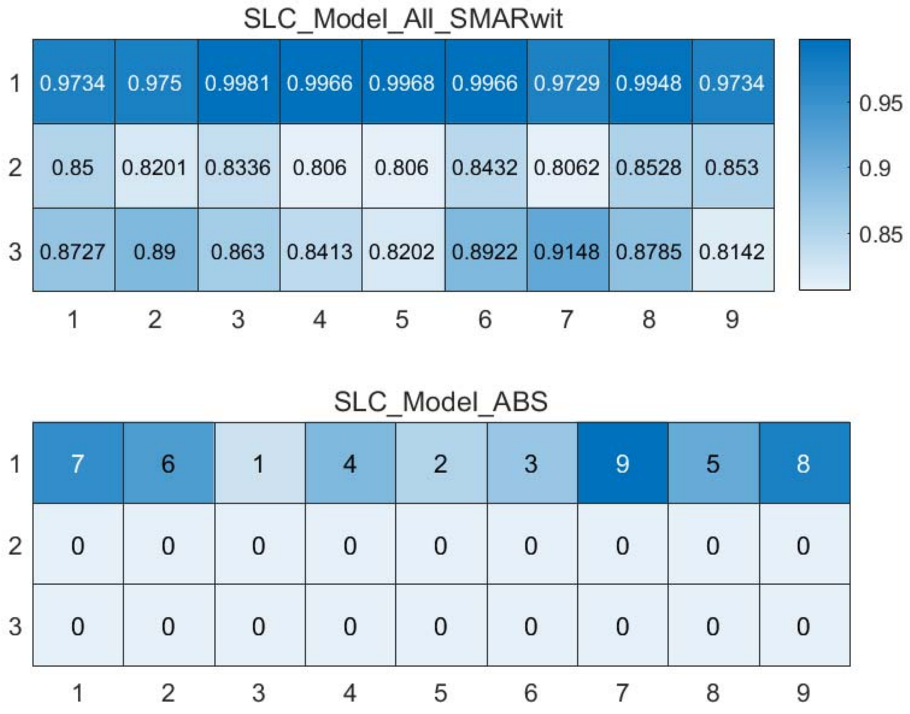


Figure S9: The ABS mode of meta classifiers for *Buettner* dataset.

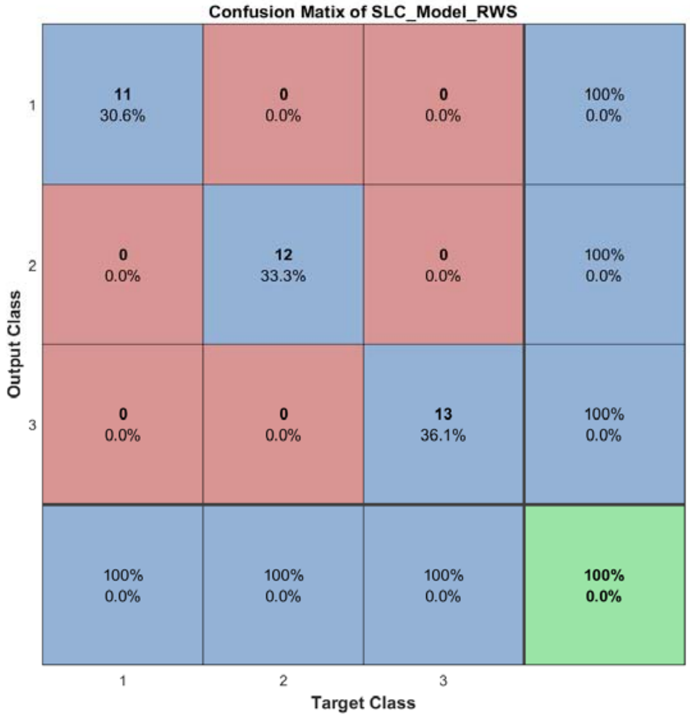


Figure S10: The confusion matrix of ensemble classifier with RWS mode for *Buettner* dataset.

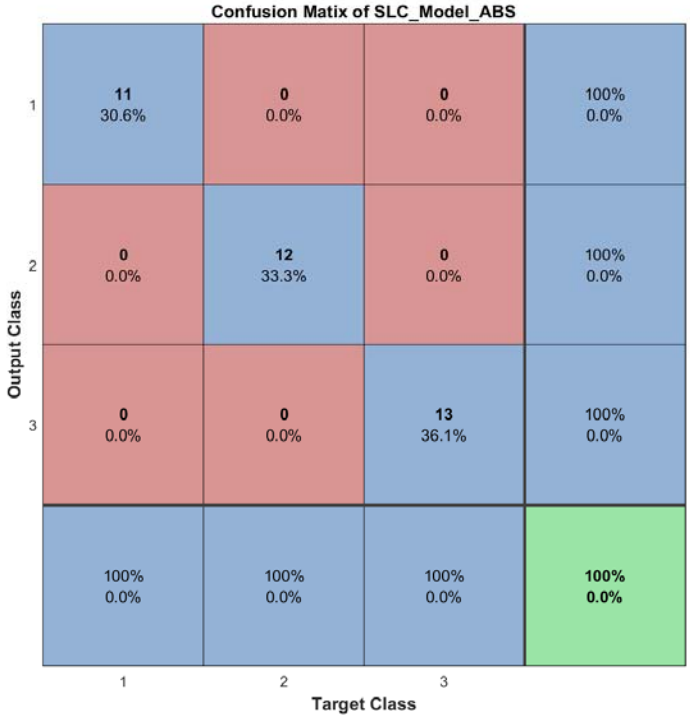


Figure S11: The confusion matrix of ensemble classifier with ABS mode for *Buettner* dataset.

#### 2.4 The running parameters and output figures for *Kolod* dataset

Table S2: The running parameters for *Kolod* dataset.

```
load('SIMLR\Kolod.mat');
[in_X_SLC,class_num,slice_tik,binary_mod] = slicematrix(in_X,true_labs,0:0.03:2);
[in_X_SLC_diff,in_X_SLC_SRCIstd,slice_bst,slice_vle] = slicediffer(in_X_SLC,slice_tik,9);
[SLC_Model_DIS1,istrain,istest] = slicemethod(in_X_SLC,true_labs,slice_tik,slice_bst,5,'correlation','inverse',5);
[SLC_Model_DIS2,istrain,istest] = slicemethod(in_X_SLC,true_labs,slice_tik,slice_bst,5,'jaccard','inverse',5);
[SLC_Model_DIS3,istrain,istest] = slicemethod(in_X_SLC,true_labs,slice_tik,slice_bst,5,'cosine','inverse',5);
[SLC_Model_All,SLC_Model_All_SMARwit] = sliceweight(SLC_Model_DIS1,SLC_Model_DIS2,SLC_Model_DIS3);
[SLC_Model_RWS,SLC_Model_All_SMESrws] = sliceswitch(SLC_Model_All,SLC_Model_All_SMARwit,'rws',2);
[SLC_Model_ABS,SLC_Model_All_SMESabs] = sliceswitch(SLC_Model_All,SLC_Model_All_SMARwit,'abs',6);
[SLC_Model_FIT1,FIT1_accuracy] = sliceprerws(in_X(istest,:),true_labs(istest),binary_mod,class_num,slice_vle,SLC_Model_RWS);
[SLC_Model_FIT2,FIT2_accuracy] = slicepreabs(in_X(istest,:),true_labs(istest),binary_mod,class_num,slice_vle,SLC_Model_ABS);
```

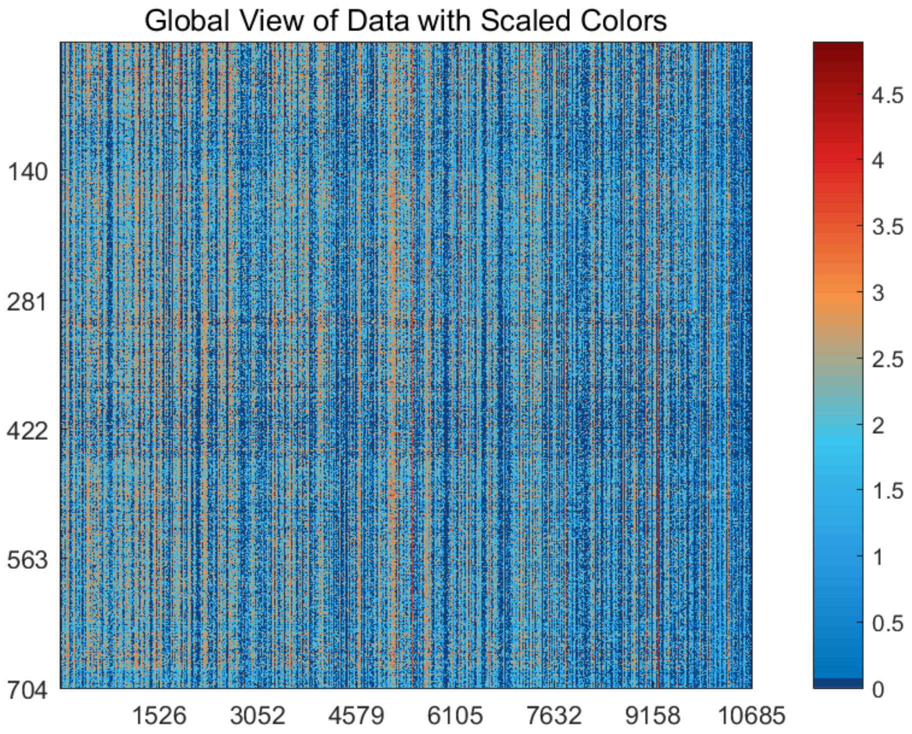


Figure S12: The global view of *Kolod* dataset with scaled colors.

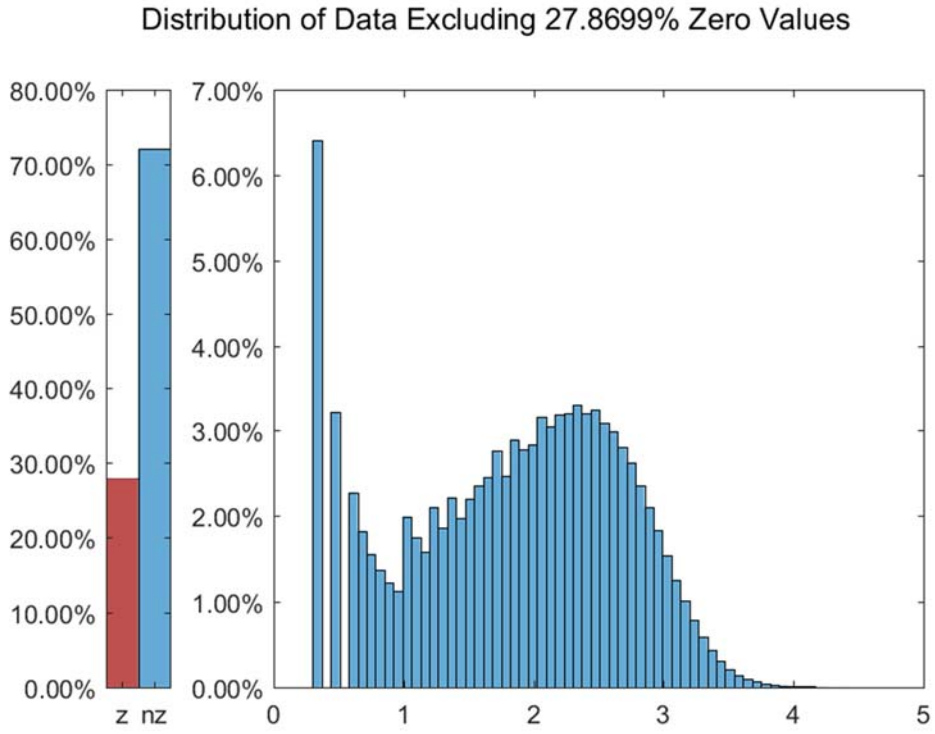


Figure S13: The distribution of *Kolod* dataset excluding all zero values.

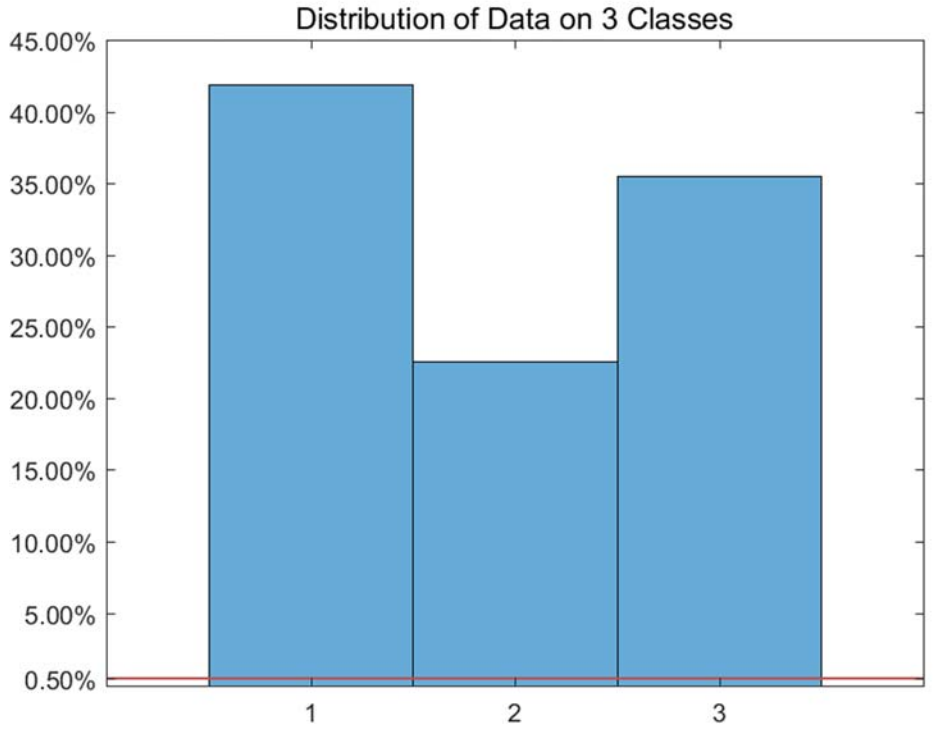


Figure S14: The distribution of *Kolod* dataset on every classes.

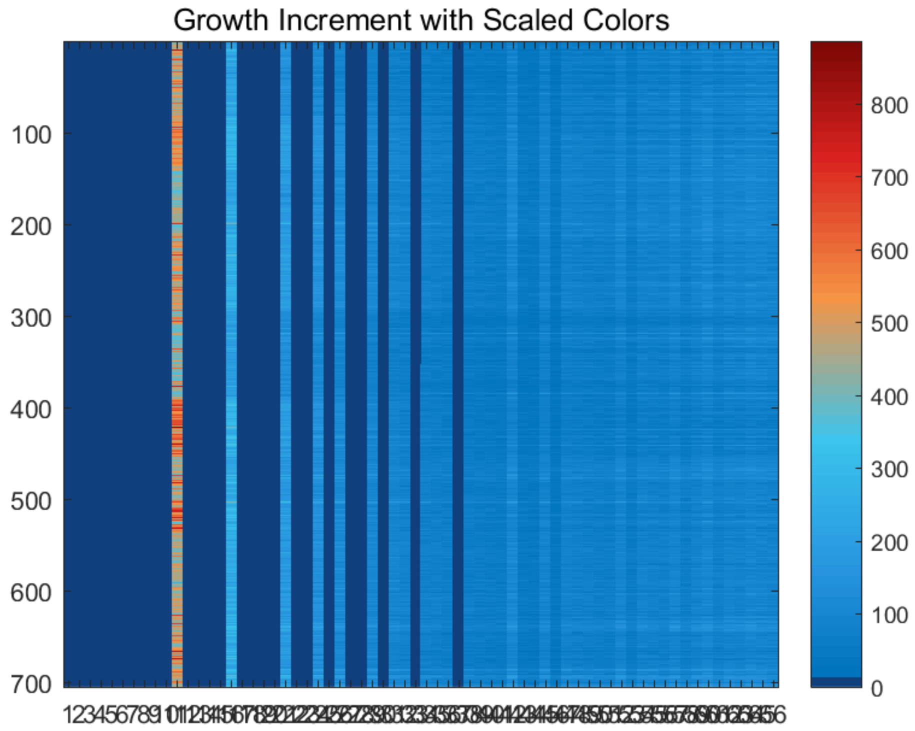


Figure S15: The growth increment with scaled colors for *Kolod* dataset.

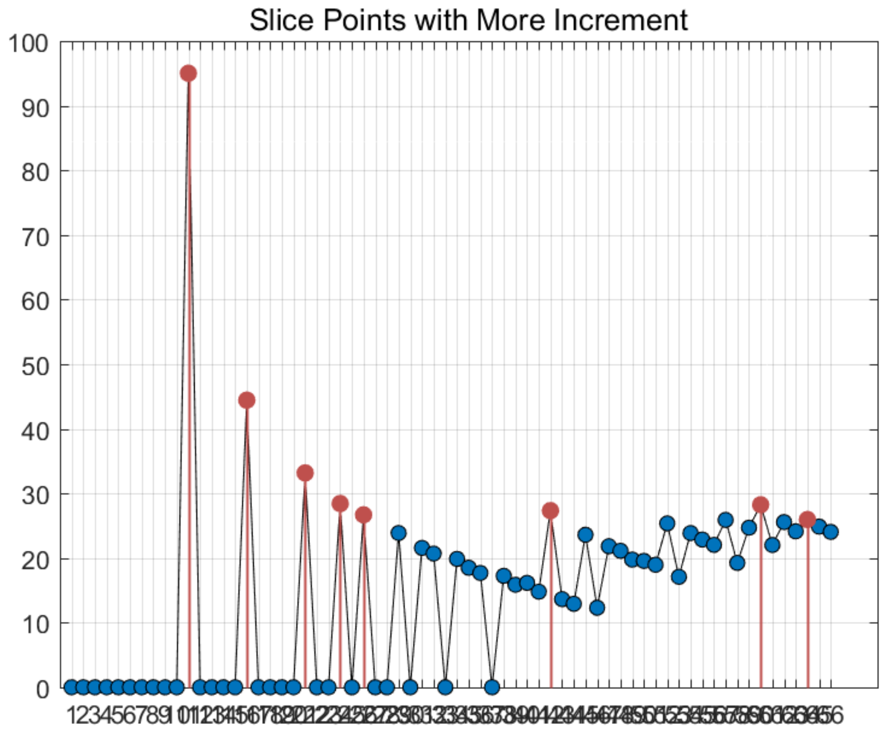


Figure S16: The slice points with more increment for *Kolod* dataset.

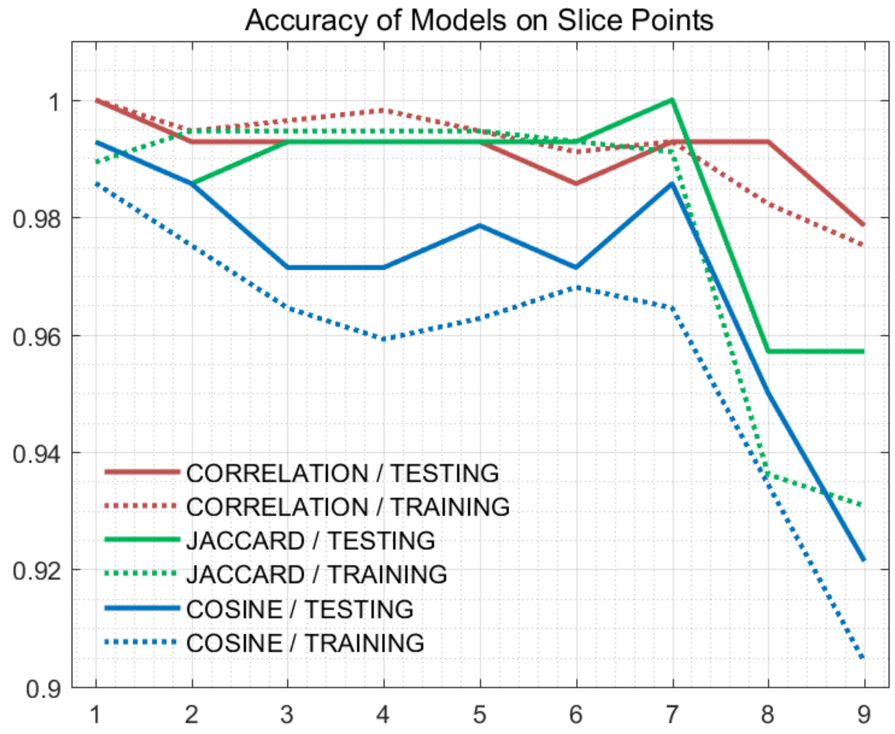


Figure S17: The accuracy of models on every slice points for *Kolod* dataset.

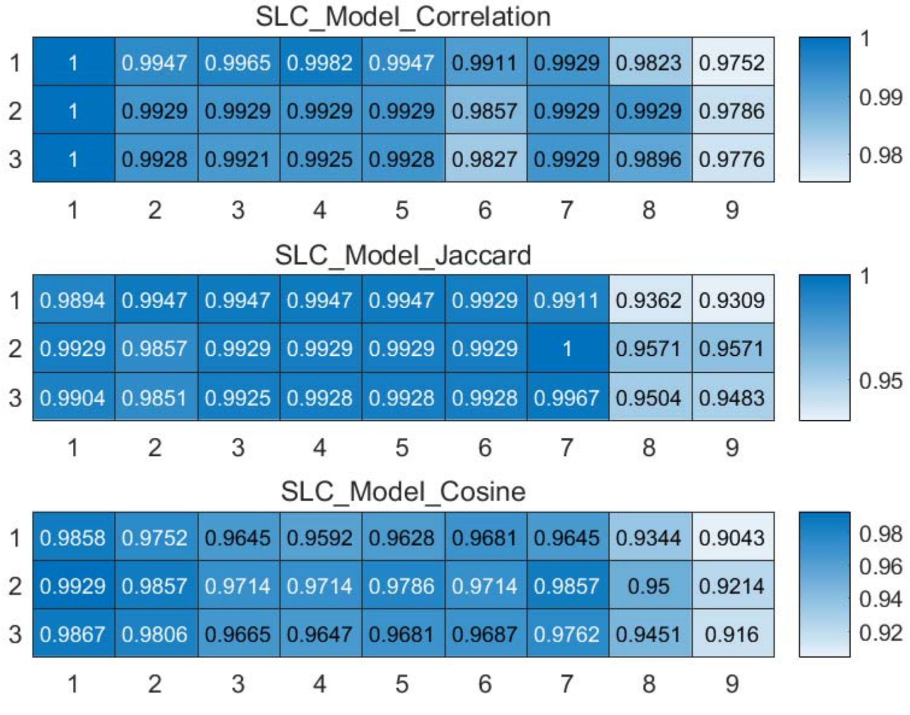


Figure S18: The weighted accuracy on every slice points for *Kolod* dataset.

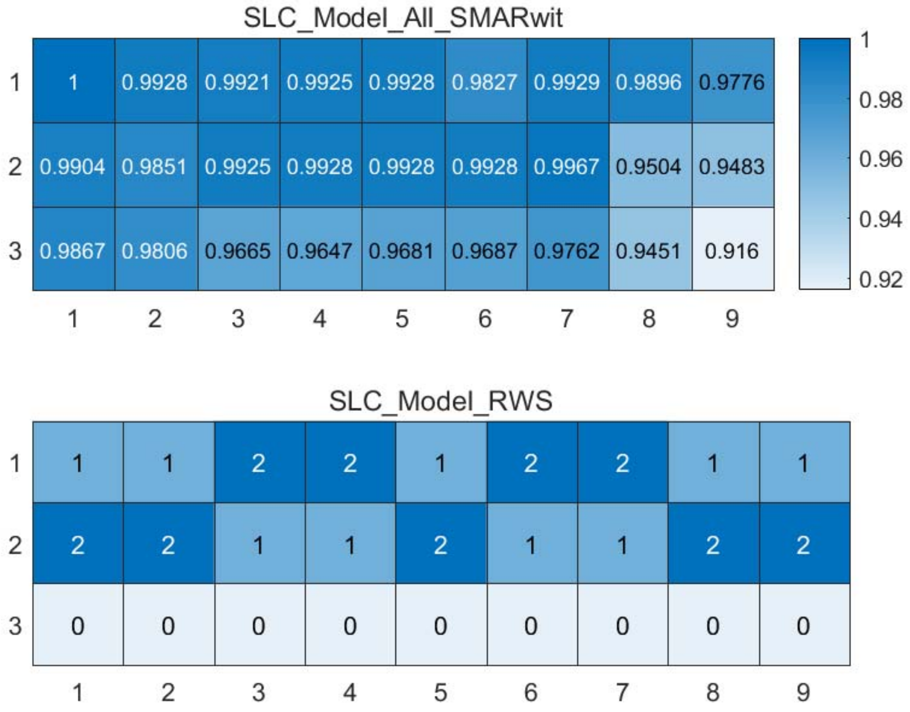


Figure S19: The RWS mode of meta classifiers for *Kolod* dataset.

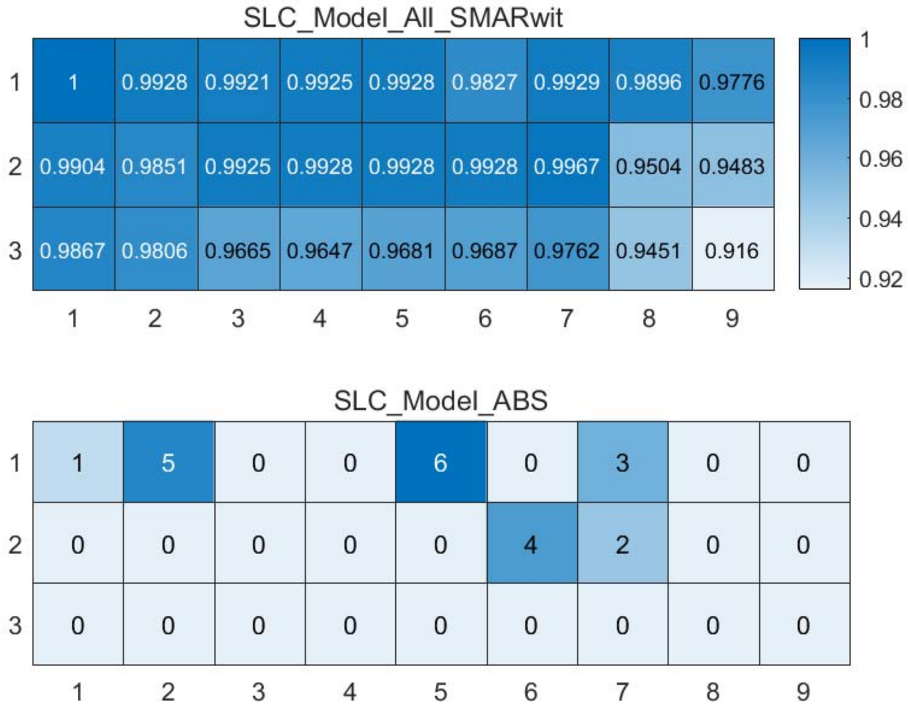


Figure S20: The ABS mode of meta classifiers for *Kolod* dataset.

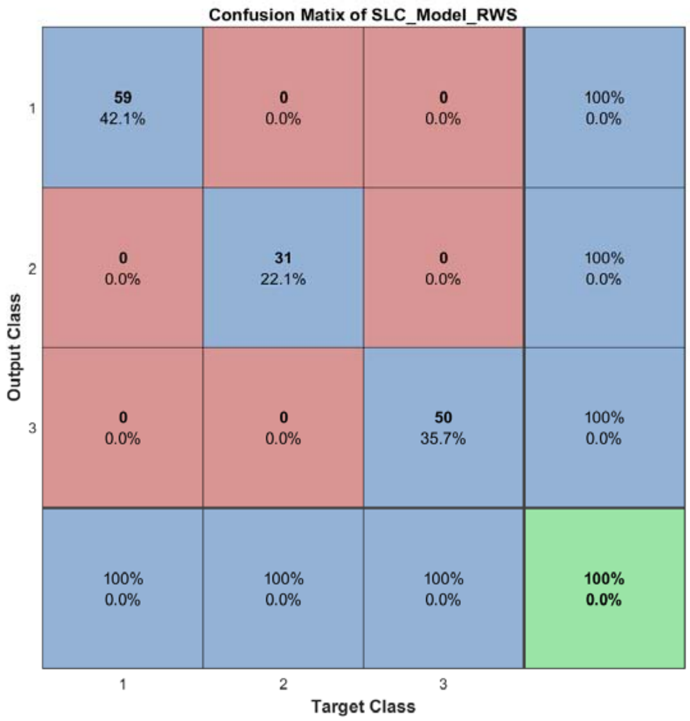


Figure S21: The confusion matrix of ensemble classifier with RWS mode for *Kolod* dataset.

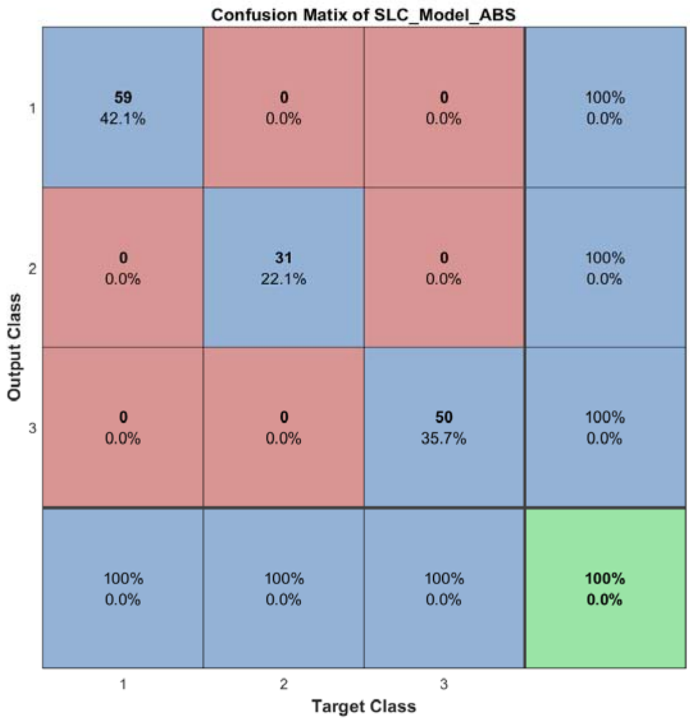


Figure S22: The confusion matrix of ensemble classifier with ABS mode for *Kolod* dataset.

#### 2.5 The running parameters and output figures for *Pollen* dataset

Table S3: The running parameters for *Pollen* dataset.

```
load('SIMLR\Pollen.mat');
[in_X_SLC,class_num,slice_tik,binary_mod] = slicematrix(in_X,true_labs,0:0.05:4);
[in_X_SLC_diff,in_X_SLC_SRCIstd,slice_bst,slice_vle] = slicediffer(in_X_SLC,slice_tik,9);
[SLC_Model_DIS1,istrain,istest] = slicemethod(in_X_SLC,true_labs,slice_tik,slice_bst,1,'correlation','inverse',5);
[SLC_Model_DIS2,istrain,istest] = slicemethod(in_X_SLC,true_labs,slice_tik,slice_bst,1,'jaccard','inverse',5);
[SLC_Model_DIS3,istrain,istest] = slicemethod(in_X_SLC,true_labs,slice_tik,slice_bst,1,'cosine','inverse',5);
[SLC_Model_All,SLC_Model_All_SMARwit] = sliceweight(SLC_Model_DIS1,SLC_Model_DIS2,SLC_Model_DIS3);
[SLC_Model_RWS,SLC_Model_All_SMESrws] = sliceswitch(SLC_Model_All,SLC_Model_All_SMARwit,'rws',2);
[SLC_Model_ABS,SLC_Model_All_SMESabs] = sliceswitch(SLC_Model_All,SLC_Model_All_SMARwit,'abs',9);
[SLC_Model_FIT1,FIT1_accuracy] = sliceprerws(in_X(istest,:),true_labs(istest),binary_mod,class_num,slice_vle,SLC_Model_RWS);
[SLC_Model_FIT2,FIT2_accuracy] = slicepreabs(in_X(istest,:),true_labs(istest),binary_mod,class_num,slice_vle,SLC_Model_ABS);
```

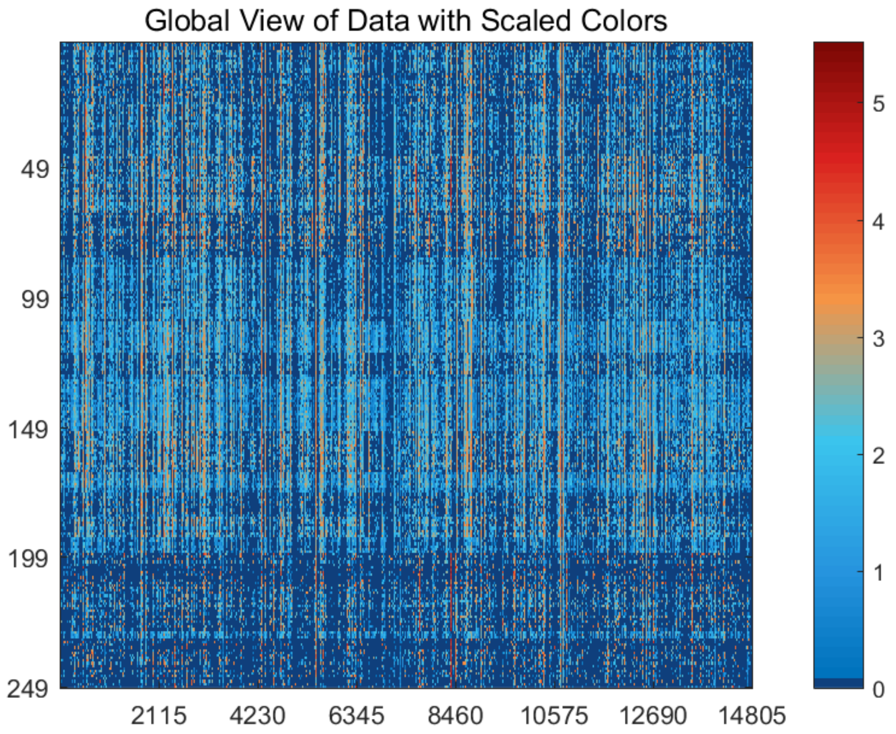


Figure S23: The global view of *Pollen* dataset with scaled colors.

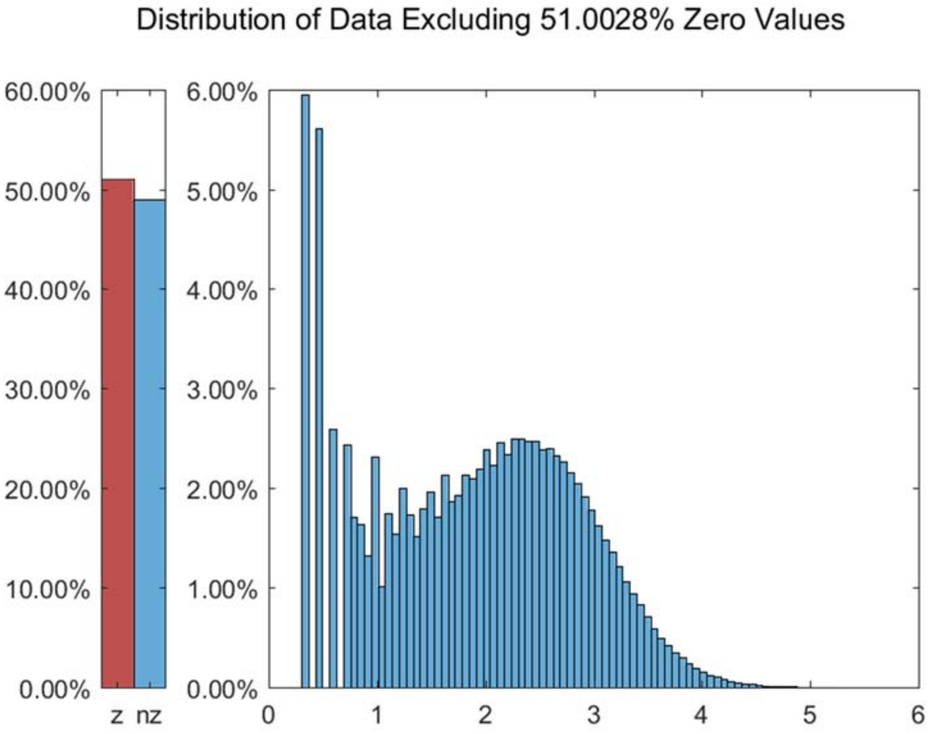


Figure S24: The distribution of *Pollen* dataset excluding all zero values.

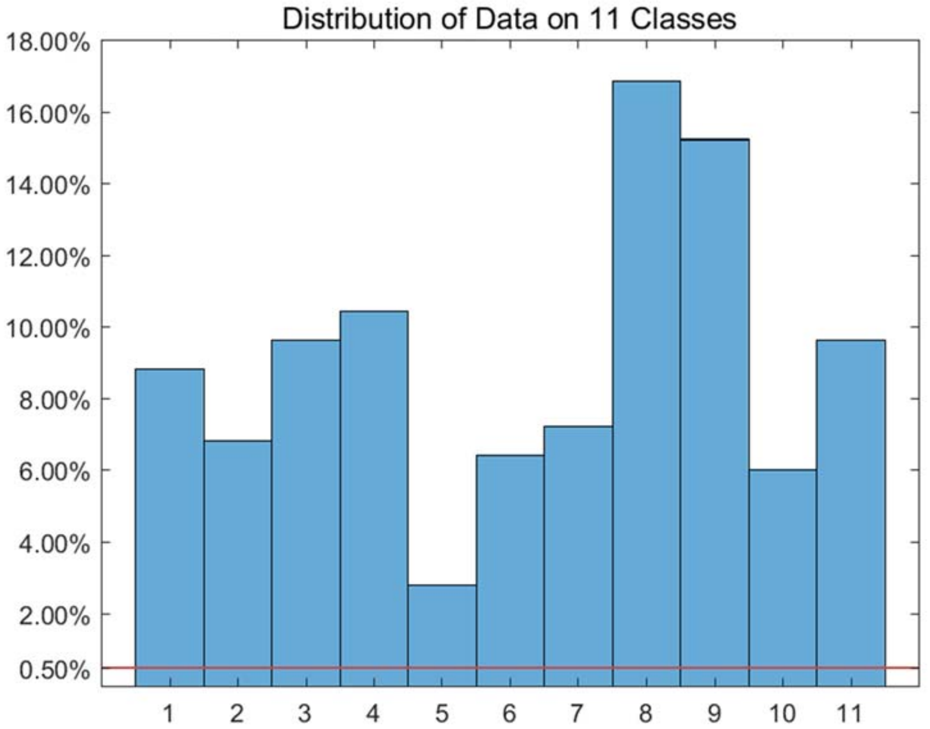


Figure S25: The distribution of *Pollen* dataset on every classes.

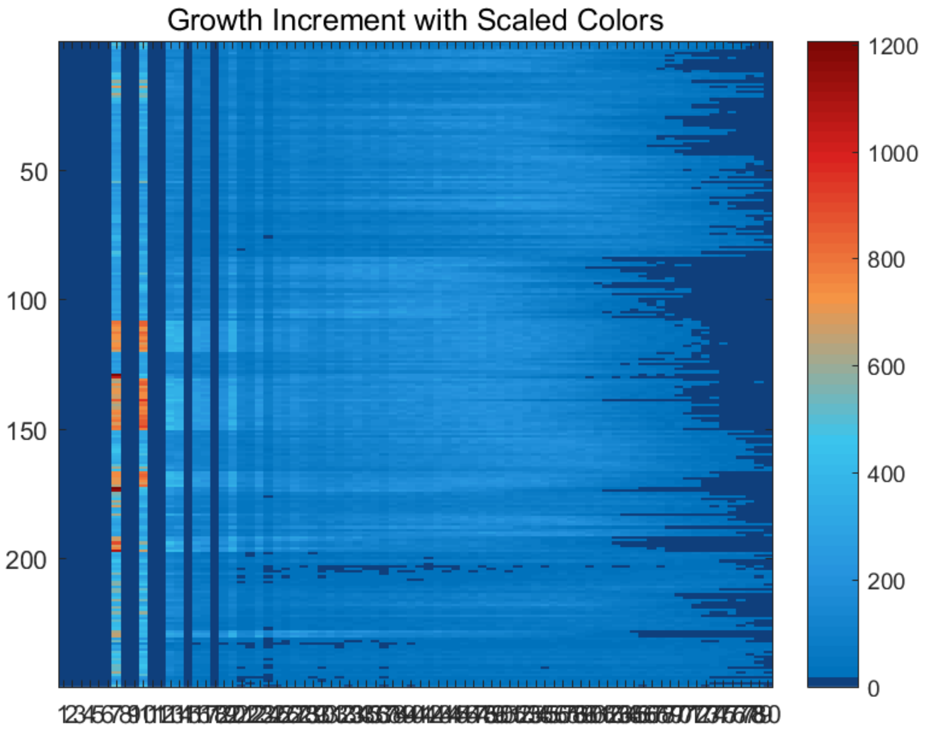


Figure S26: The growth increment with scaled colors for *Pollen* dataset.

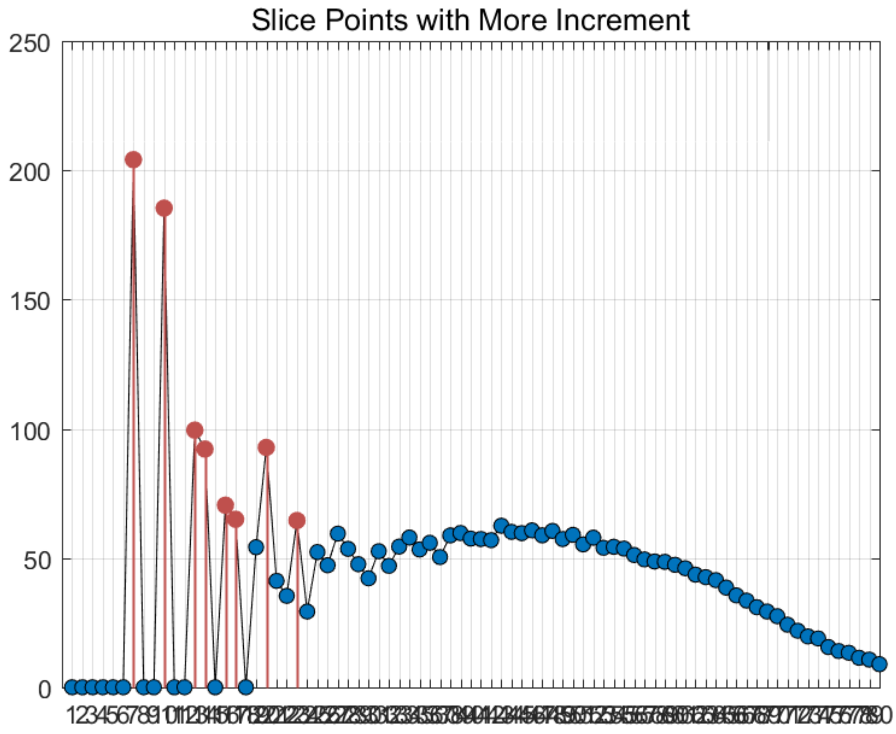


Figure S27: The slice points with more increment for *Pollen* dataset.

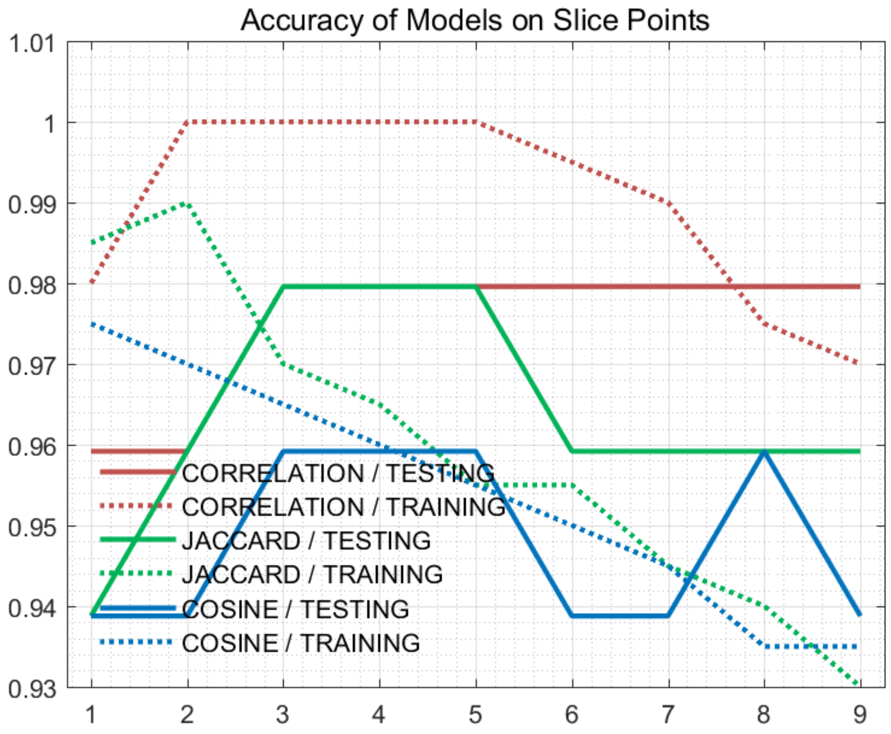


Figure S28: The accuracy of models on every slice points for *Pollen* dataset.

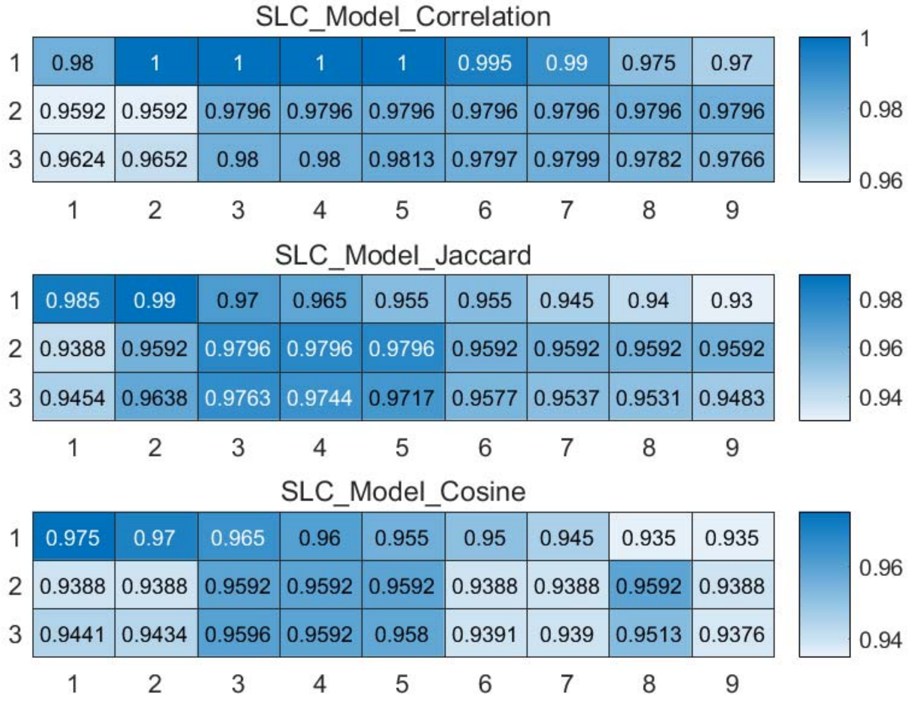


Figure S29: The weighted accuracy on every slice points for *Pollen* dataset.

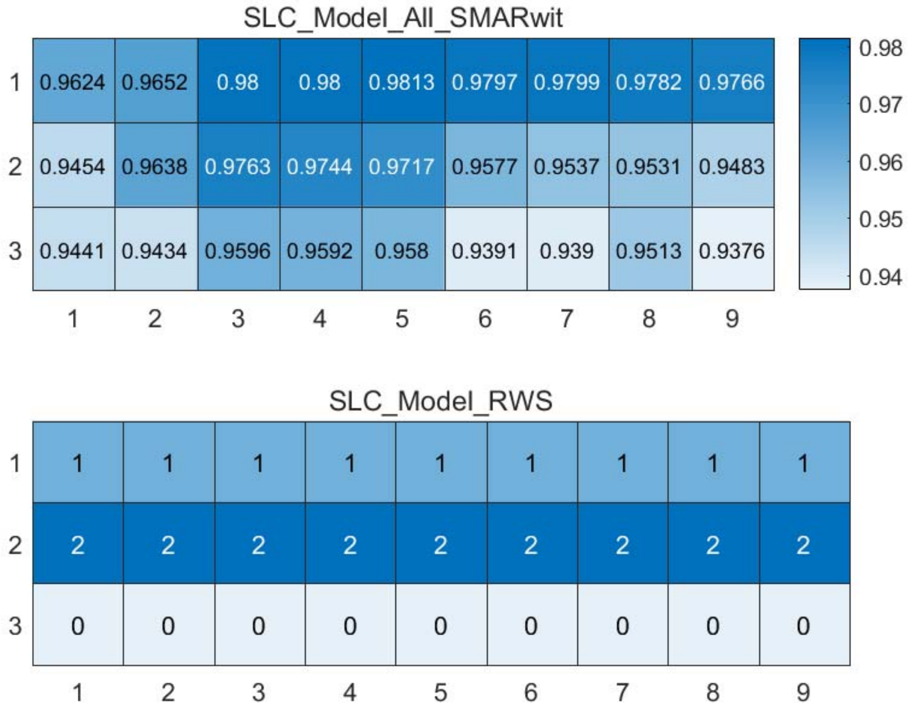


Figure S30: The RWS mode of meta classifiers for *Pollen* dataset.

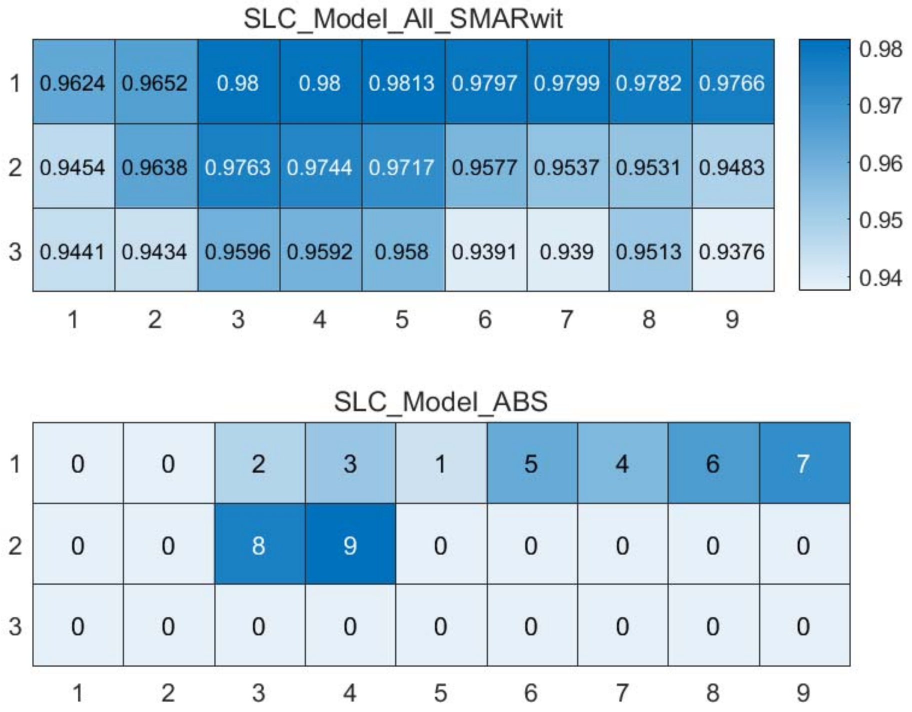


Figure S31: The ABS mode of meta classifiers for *Pollen* dataset.

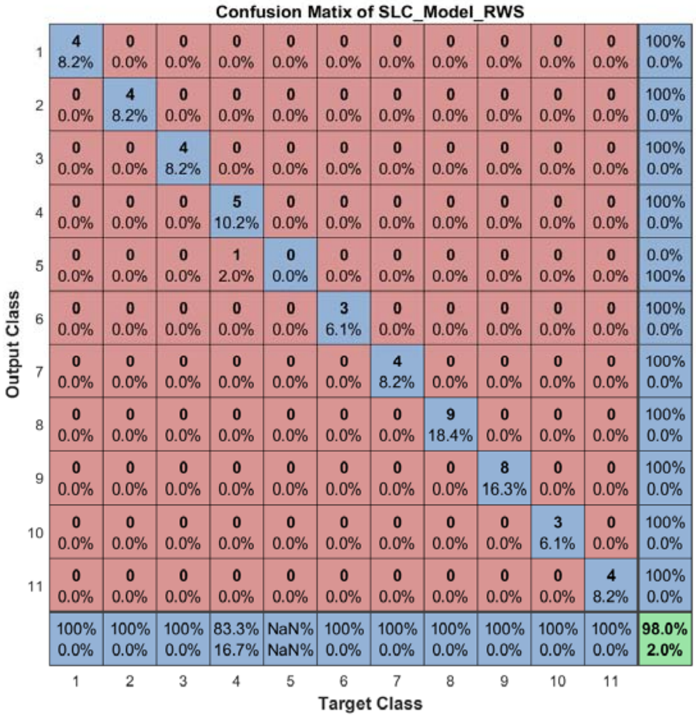


Figure S32: The confusion matrix of ensemble classifier with RWS mode for *Pollen* dataset.

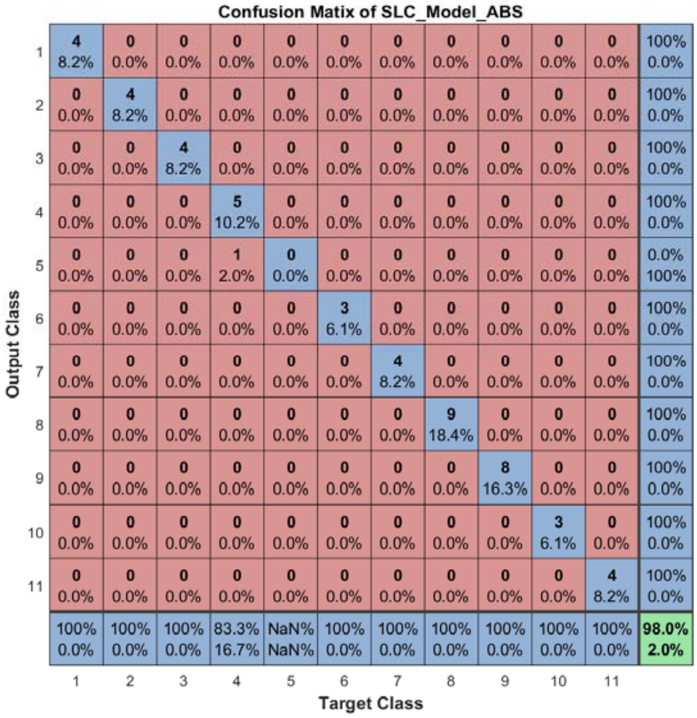


Figure S33: The confusion matrix of ensemble classifier with ABS mode for *Pollen* dataset.

#### 2.6 The running parameters and output figures for *Usoskin* dataset

Table S4: The running parameters for *Usoskin* dataset.

```
load('SIMLR\Usoskin.mat'); in_X = 0.3*in_X;
[in_X_SLC,class_num,slice_tik,binary_mod] = slicematrix(in_X,true_labs,0:0.05:2.6);
[in_X_SLC_diff,in_X_SLC_SRCIstd,slice_bst,slice_vle] = slicediffer(in_X_SLC,slice_tik,21);
[SLC_Model_DIS1,istrain,istest] = slicemethod(in_X_SLC,true_labs,slice_tik,slice_bst,5,'correlation','inverse',5);
[SLC_Model_DIS2,istrain,istest] = slicemethod(in_X_SLC,true_labs,slice_tik,slice_bst,5,'jaccard','inverse',5);
[SLC_Model_DIS3,istrain,istest] = slicemethod(in_X_SLC,true_labs,slice_tik,slice_bst,5,'cosine','inverse',5);
[SLC_Model_All,SLC_Model_All_SMARwit] = sliceweight(SLC_Model_DIS1,SLC_Model_DIS2,SLC_Model_DIS3);
[SLC_Model_RWS,SLC_Model_All_SMESrws] = sliceswitch(SLC_Model_All,SLC_Model_All_SMARwit,'rws',2);
[SLC_Model_ABS,SLC_Model_All_SMESabs] = sliceswitch(SLC_Model_All,SLC_Model_All_SMARwit,'abs',6);
[SLC_Model_FIT1,FIT1_accuracy] = sliceprerws(in_X(istest,:),true_labs(istest),binary_mod,class_num,slice_vle,SLC_Model_RWS);
[SLC_Model_FIT2,FIT2_accuracy] = slicepreabs(in_X(istest,:),true_labs(istest),binary_mod,class_num,slice_vle,SLC_Model_ABS);
```

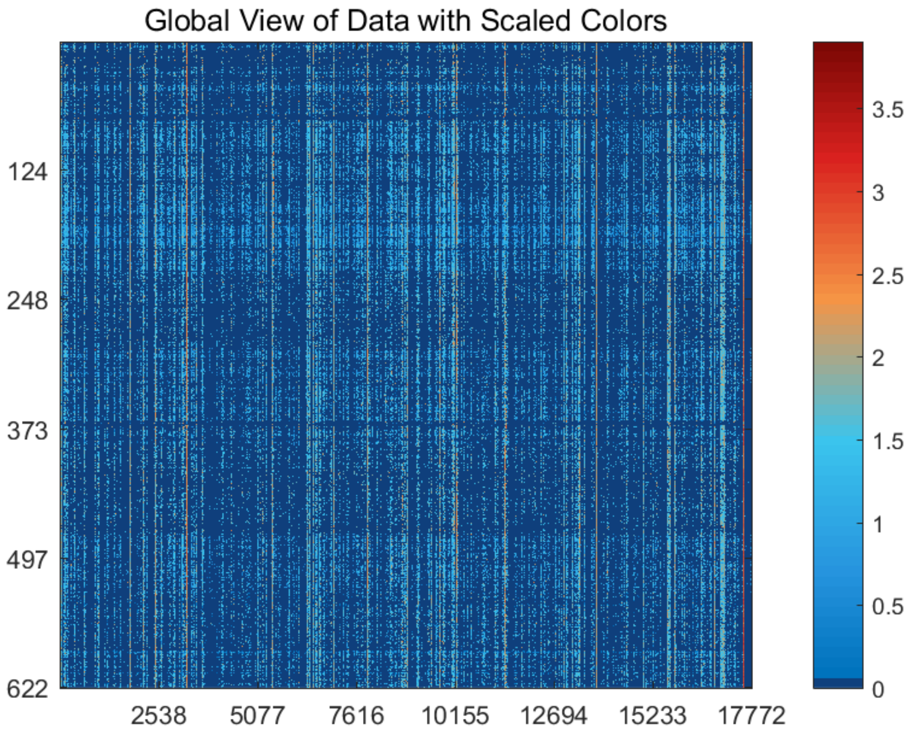


Figure S34: The global view of *Usoskin* dataset with scaled colors.

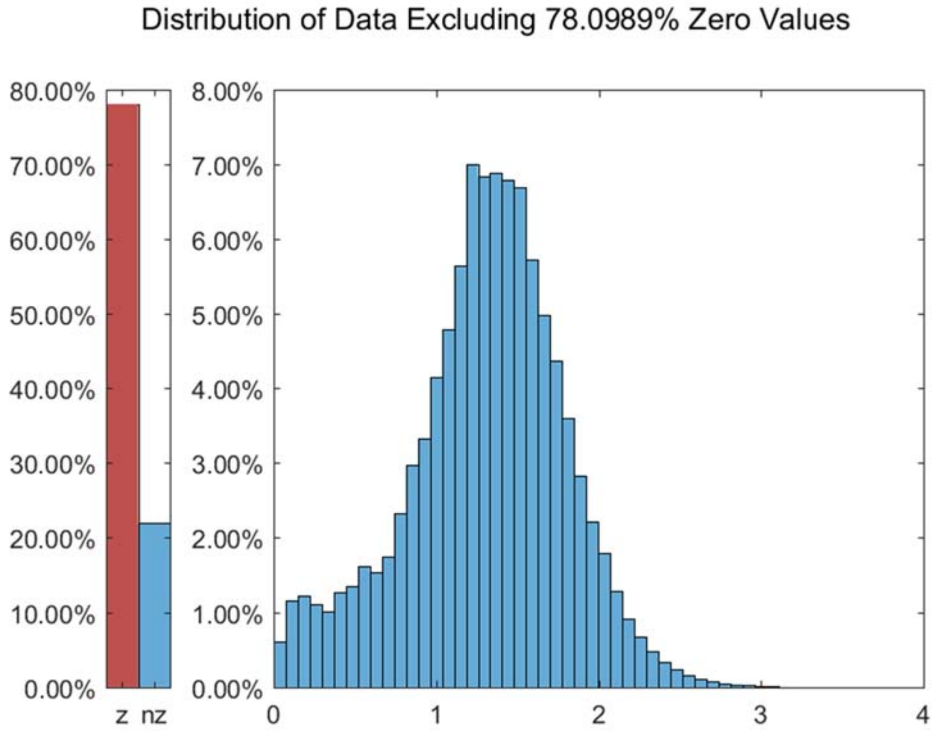


Figure S35: The distribution of *Usoskin* dataset excluding all zero values.

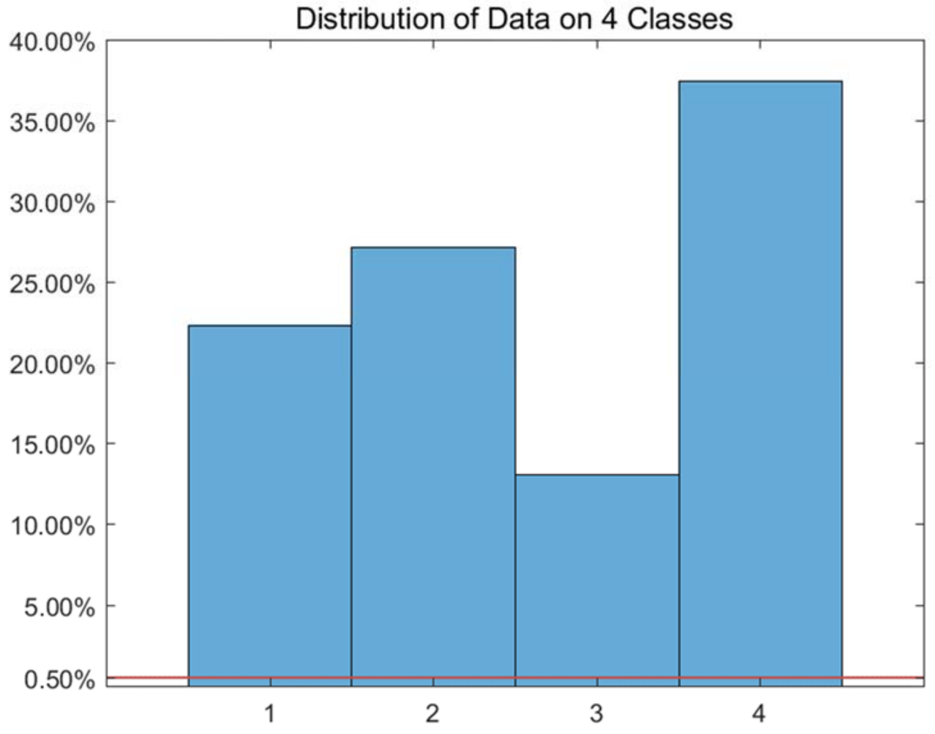


Figure S36: The distribution of *Usoskin* dataset on every classes.

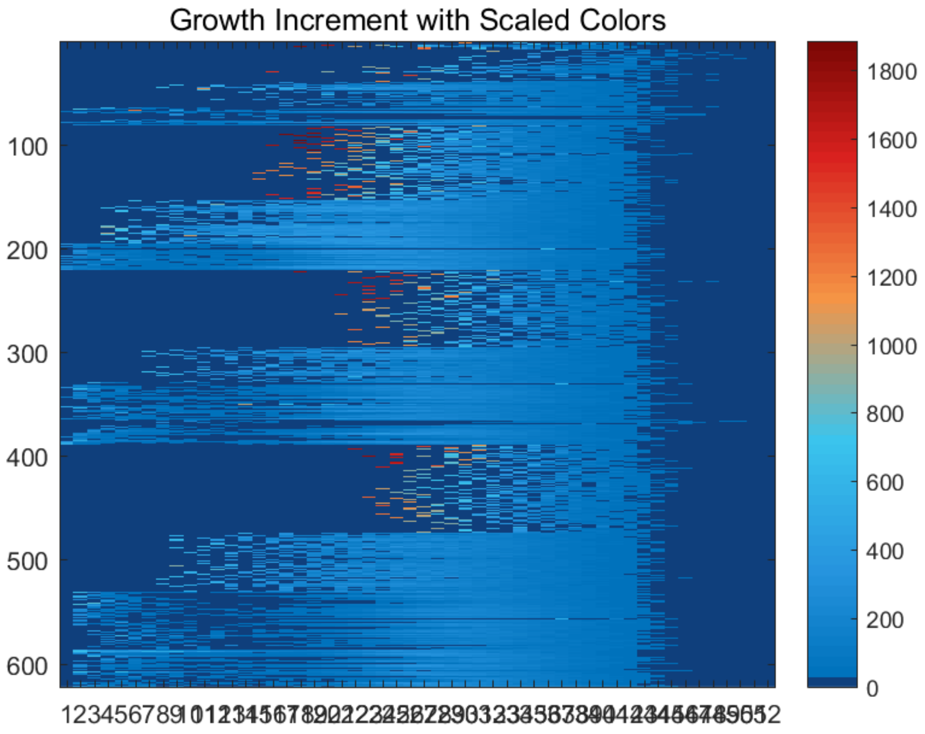


Figure S37: The growth increment with scaled colors for *Usoskin* dataset.

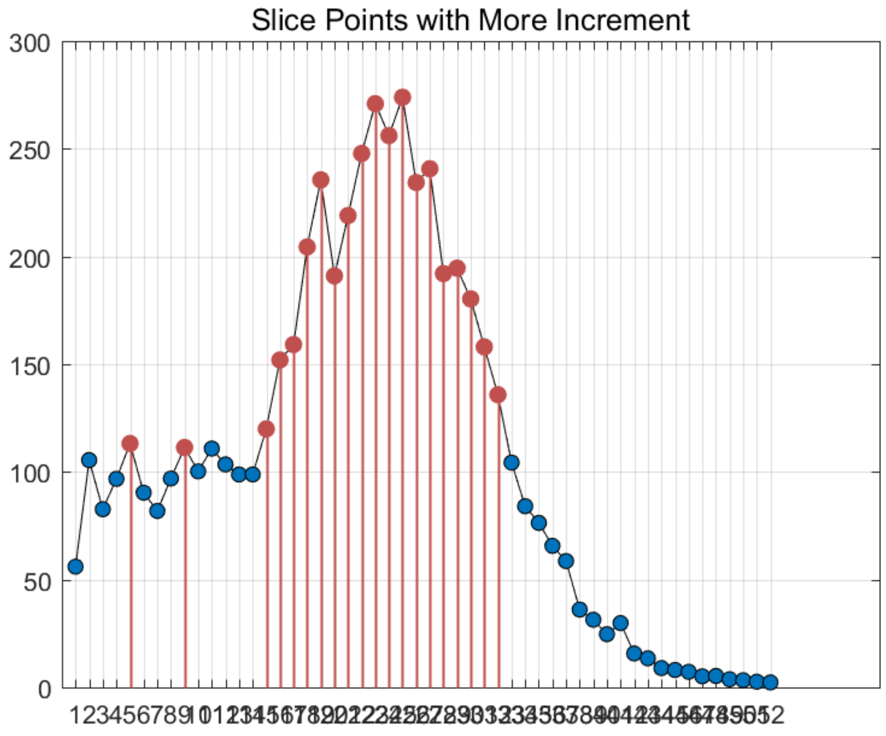


Figure S38: The slice points with more increment for *Usoskin* dataset.

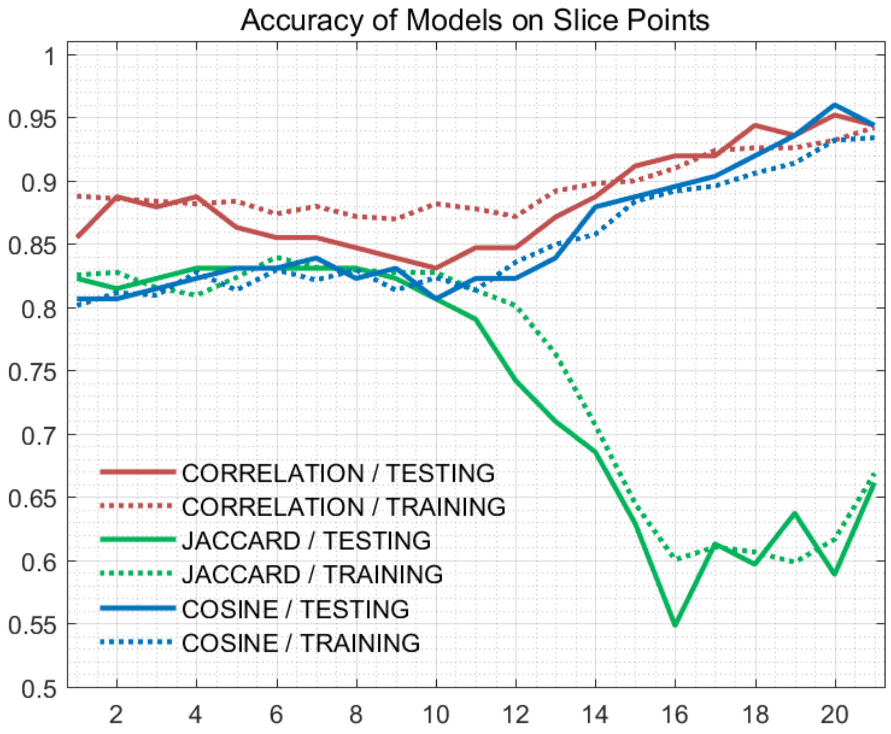


Figure S39: The accuracy of models on every slice points for *Usoskin* dataset.

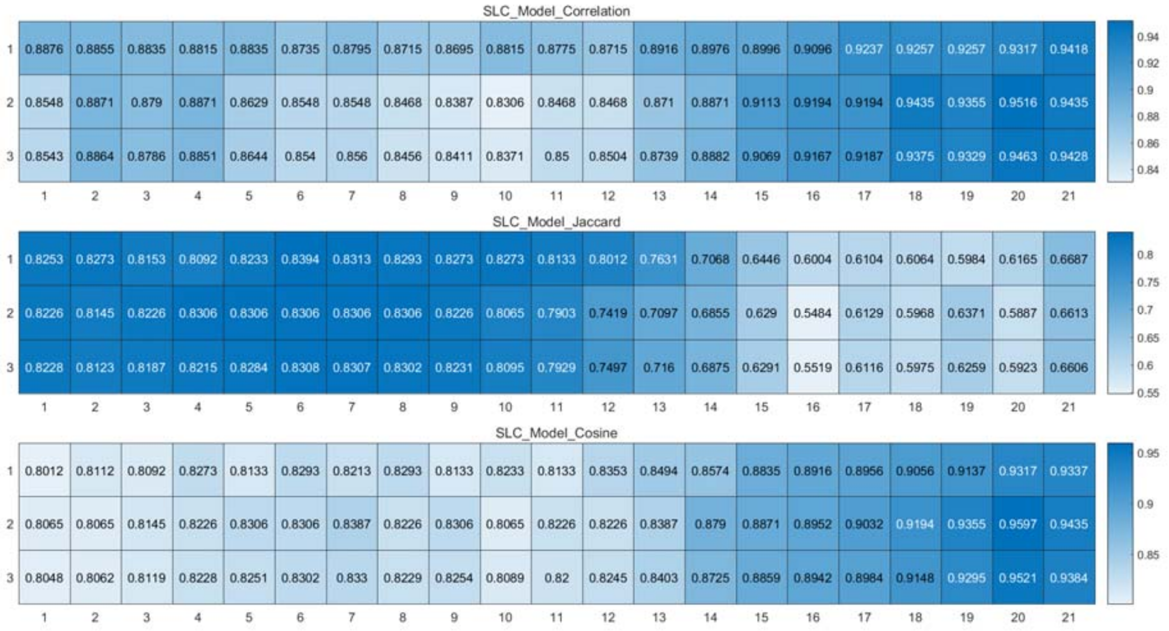


Figure S40: The weighted accuracy on every slice points for *Usoskin* dataset.

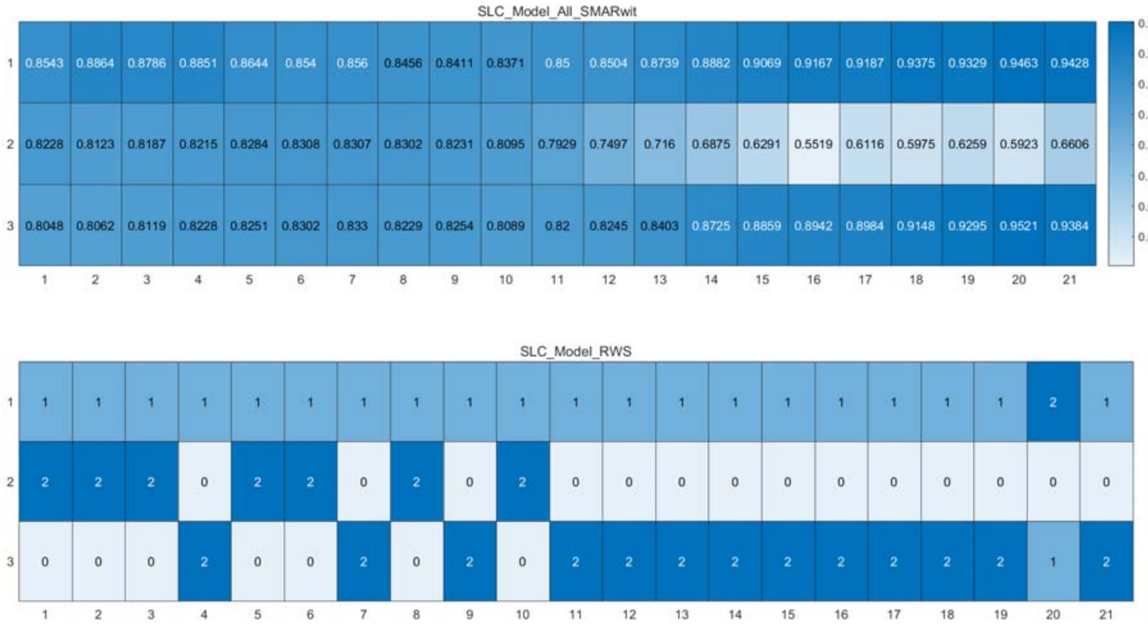


Figure S41: The RWS mode of meta classifiers for *Usoskin* dataset.

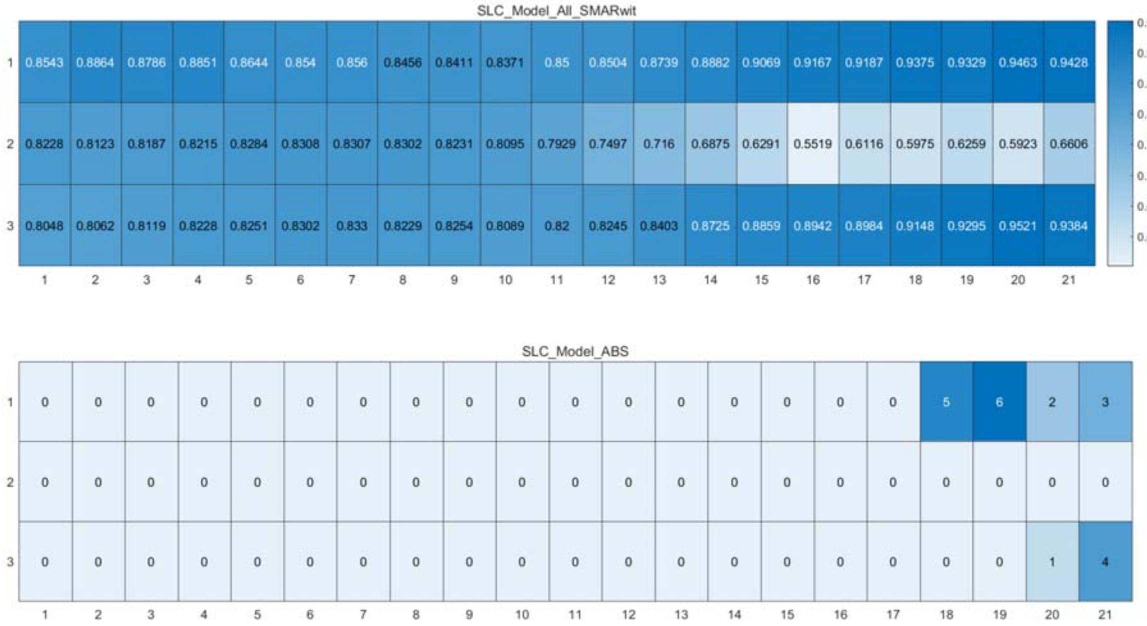


Figure S42: The ABS mode of meta classifiers for *Usoskin* dataset.

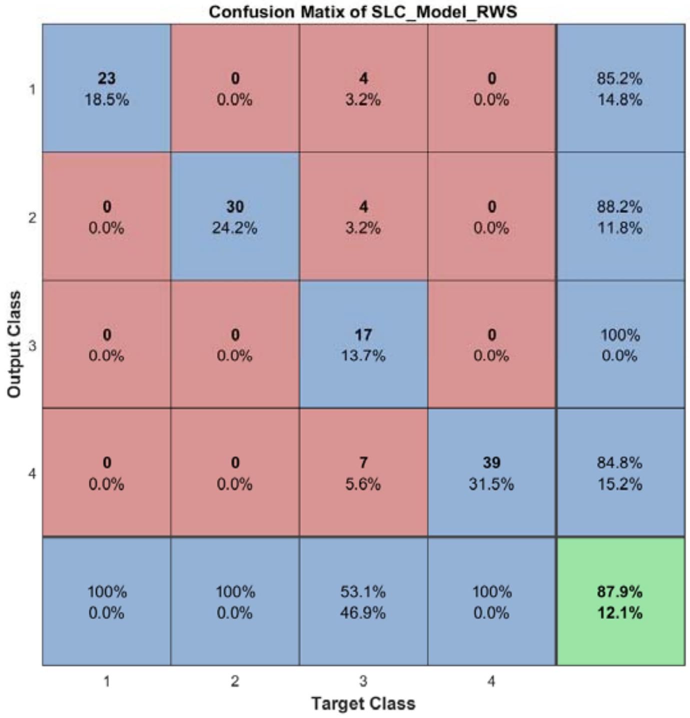


Figure S43: The confusion matrix of ensemble classifier with RWS mode for *Usoskin* dataset.

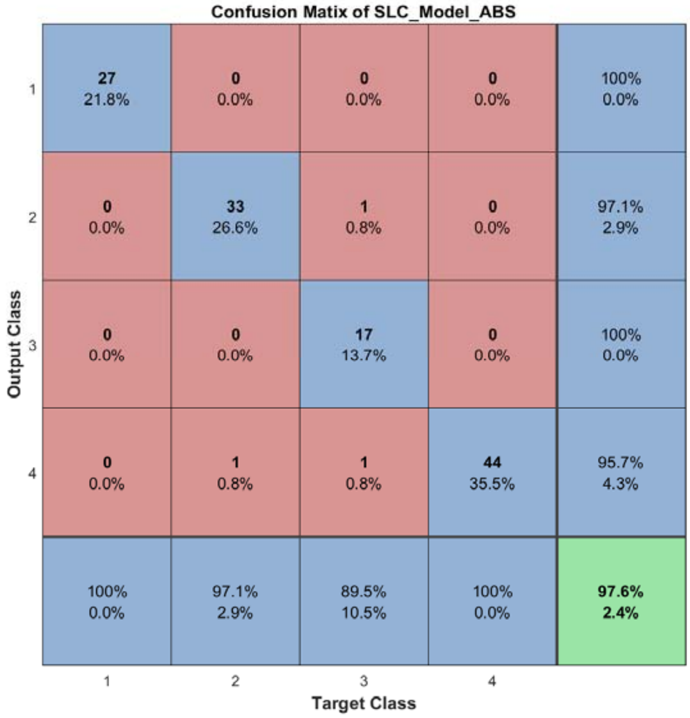


Figure S44: The confusion matrix of ensemble classifier with ABS mode for *Usoskin* dataset.

#### 2.7 The running parameters and output figures for *Usoskin* dataset (all corr)

Table S5: The running parameters for *Usoskin* dataset.

```
load('SIMLR\Usoskin.mat'); in_X = 0.3*in_X;
[in_X_SLC,class_num,slice_tik,binary_mod] = slicematrix(in_X,true_labs,1.4:0.01:1.8);
[in_X_SLC_diff,in_X_SLC_SRCIstd,slice_bst,slice_vle] = slicediffer(in_X_SLC,slice_tik,27);
[SLC_Model_DIS1,istrain,istest] = slicemethod(in_X_SLC,true_labs,slice_tik,slice_bst,6,'correlation','inverse',5);
[SLC_Model_DIS2,istrain,istest] = slicemethod(in_X_SLC,true_labs,slice_tik,slice_bst,9,'correlation','inverse',5);
[SLC_Model_DIS3,istrain,istest] = slicemethod(in_X_SLC,true_labs,slice_tik,slice_bst,12,'correlation','inverse',5);
[SLC_Model_All,SLC_Model_All_SMARwit] = sliceweight(SLC_Model_DIS1,SLC_Model_DIS2,SLC_Model_DIS3);
[SLC_Model_RWS,SLC_Model_All_SMESrws] = sliceswitch(SLC_Model_All,SLC_Model_All_SMARwit,'rws',2);
[SLC_Model_ABS,SLC_Model_All_SMESabs] = sliceswitch(SLC_Model_All,SLC_Model_All_SMARwit,'abs',9);
[SLC_Model_FIT1,FIT1_accuracy] = sliceprerws(in_X(istest,:),true_labs(istest),binary_mod,class_num,slice_vle,SLC_Model_RWS);
[SLC_Model_FIT2,FIT2_accuracy] = slicepreabs(in_X(istest,:),true_labs(istest),binary_mod,class_num,slice_vle,SLC_Model_ABS);
```

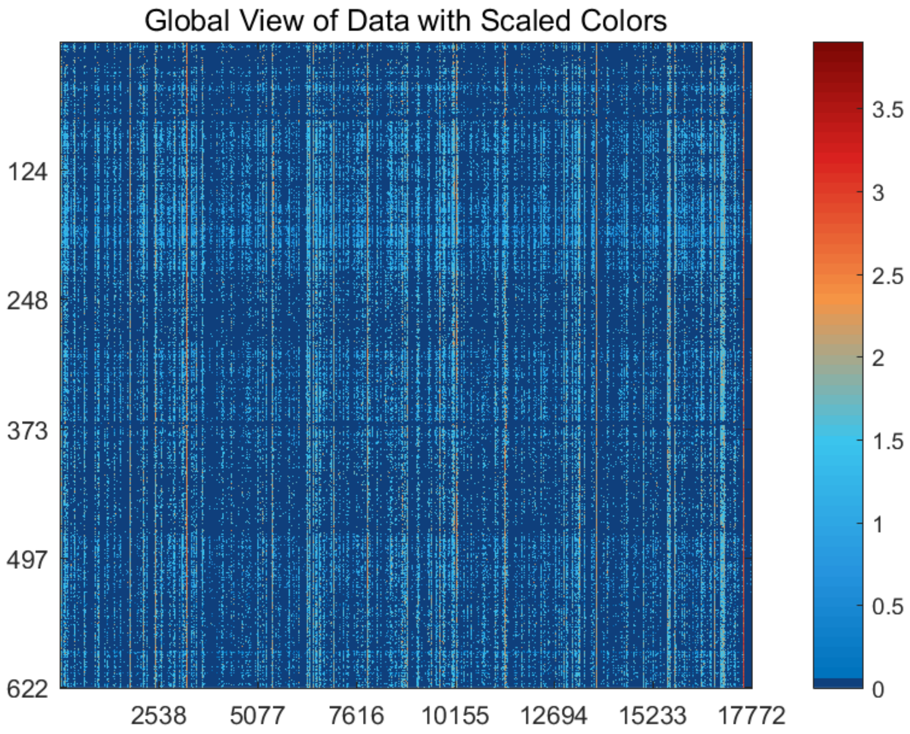


Figure S45: The global view of *Usoskin* dataset with scaled colors.

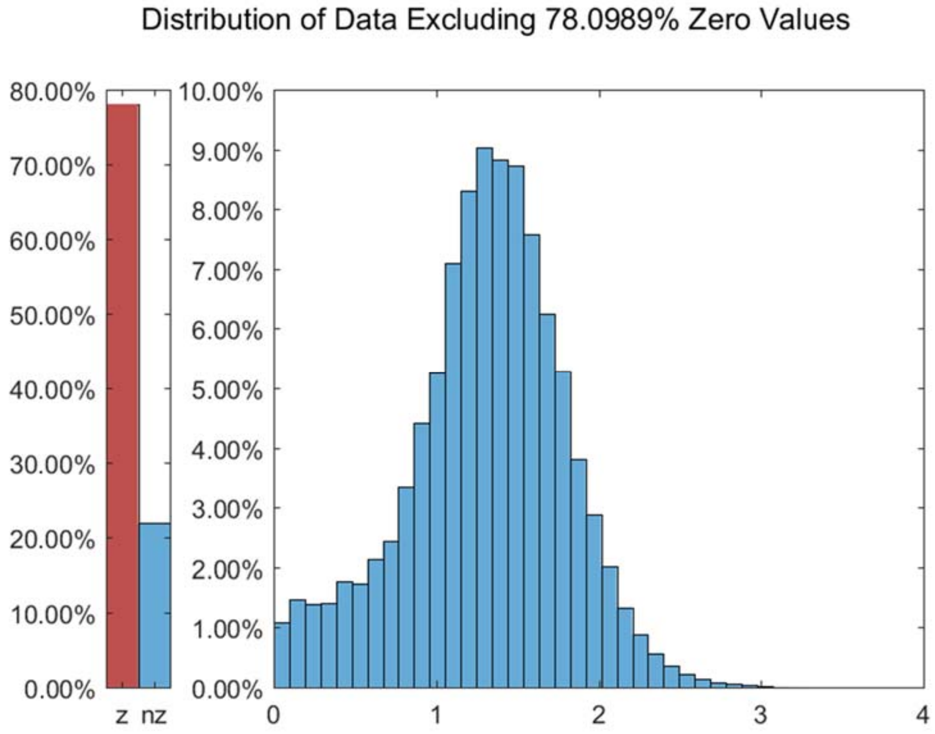


Figure S46: The distribution of *Usoskin* dataset excluding all zero values.

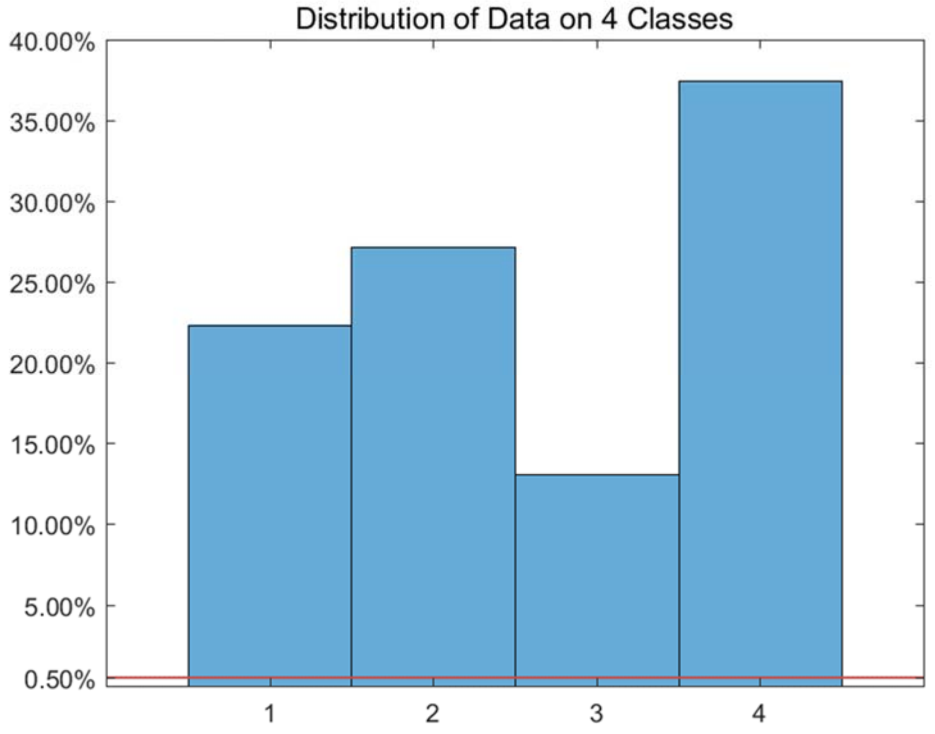


Figure S47: The distribution of *Usoskin* dataset on every classes.

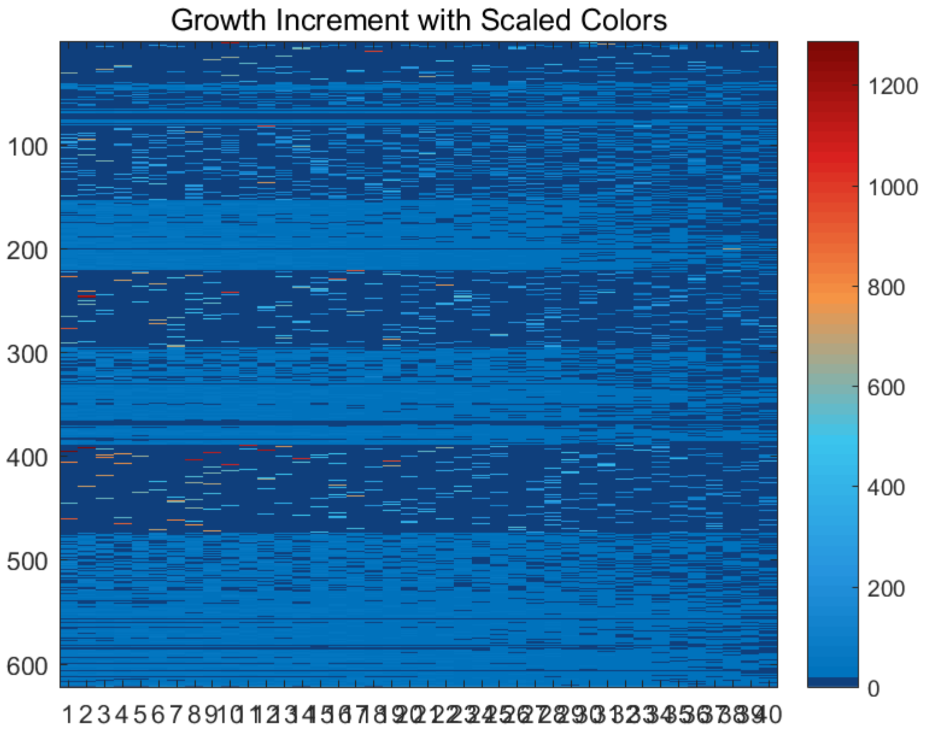


Figure S48: The growth increment with scaled colors for *Usoskin* dataset.

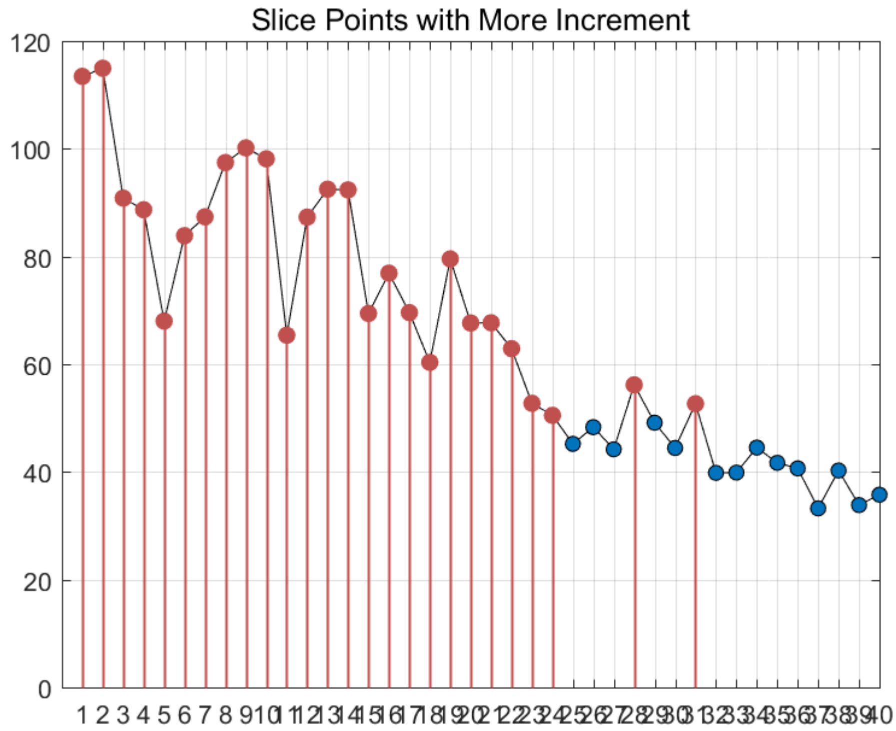


Figure S49: The slice points with more increment for *Usoskin* dataset.

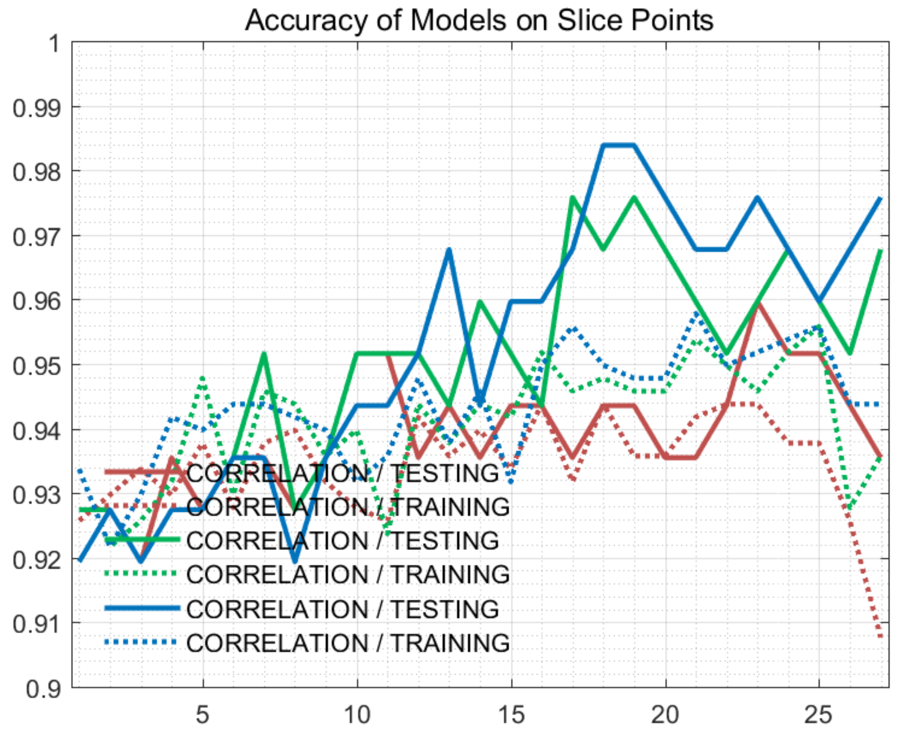


Figure S50: The accuracy of models on every slice points for *Usoskin* dataset.

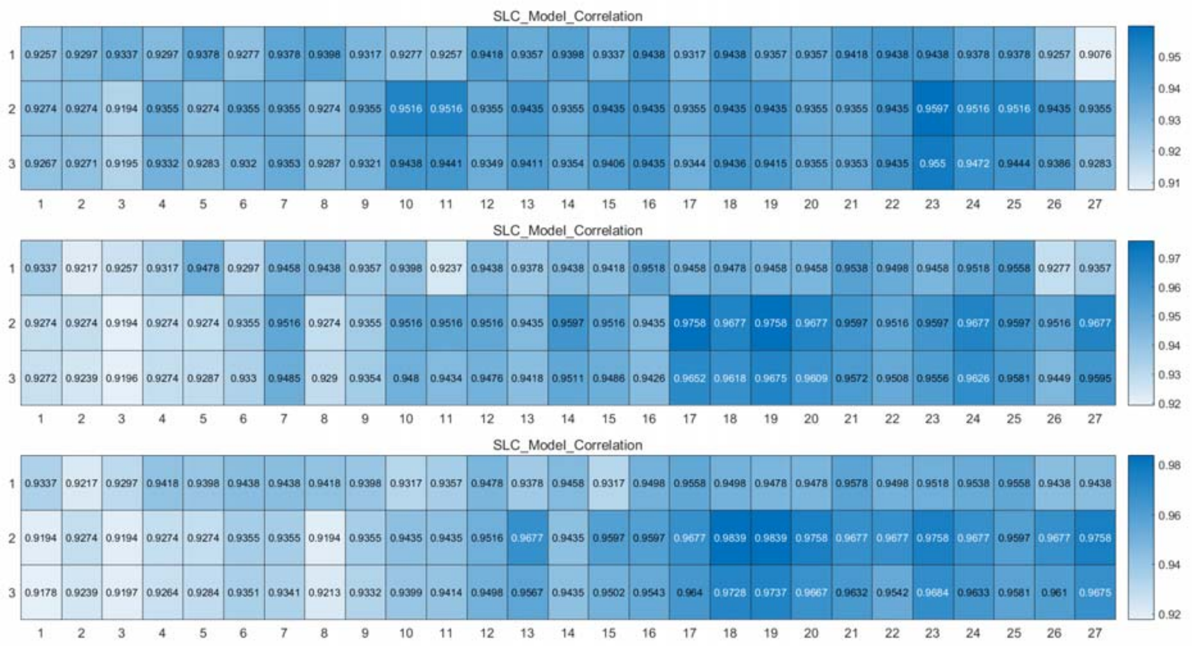


Figure S51: The weighted accuracy on every slice points for *Usoskin* dataset.

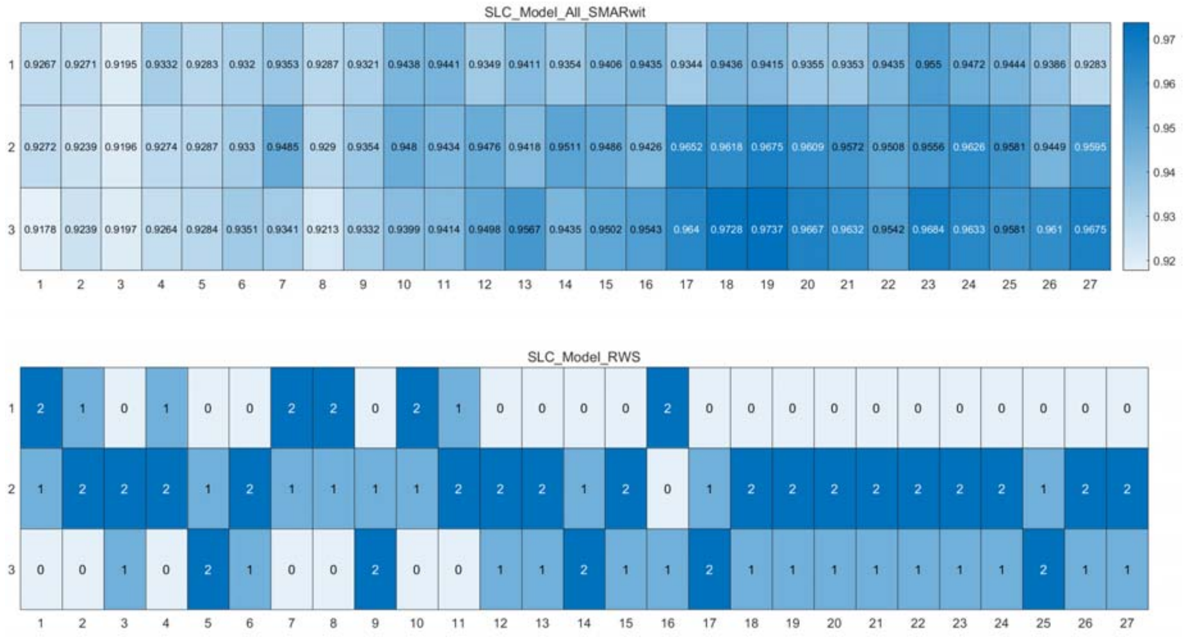


Figure S52: The RWS mode of meta classifiers for *Usoskin* dataset.

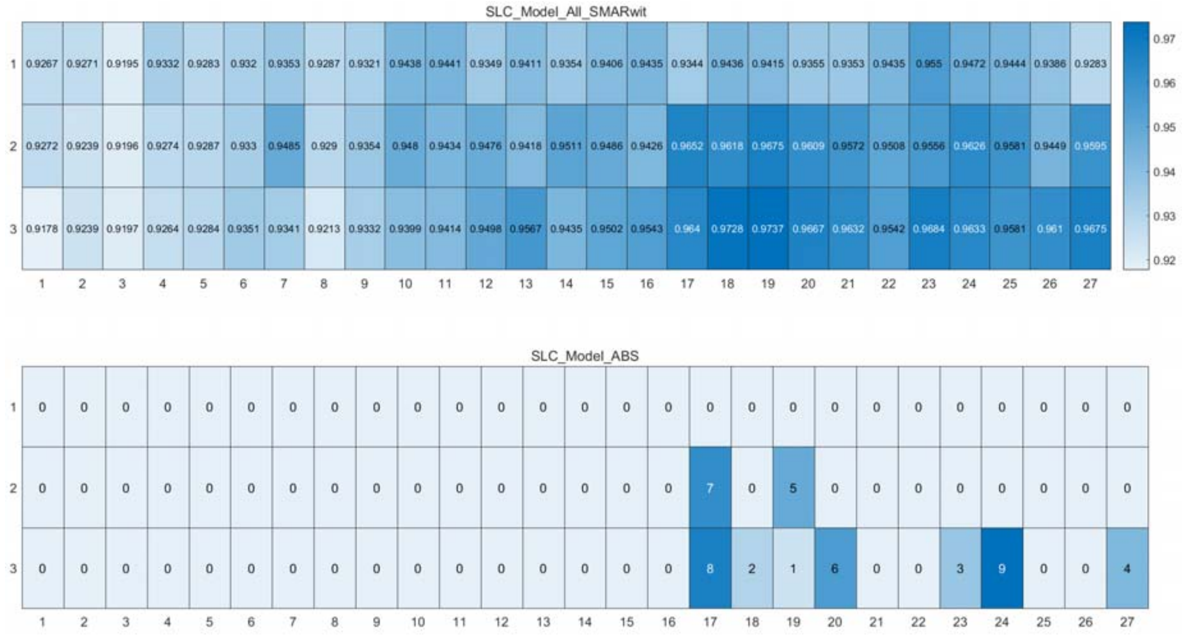


Figure S53: The ABS mode of meta classifiers for *Usoskin* dataset.

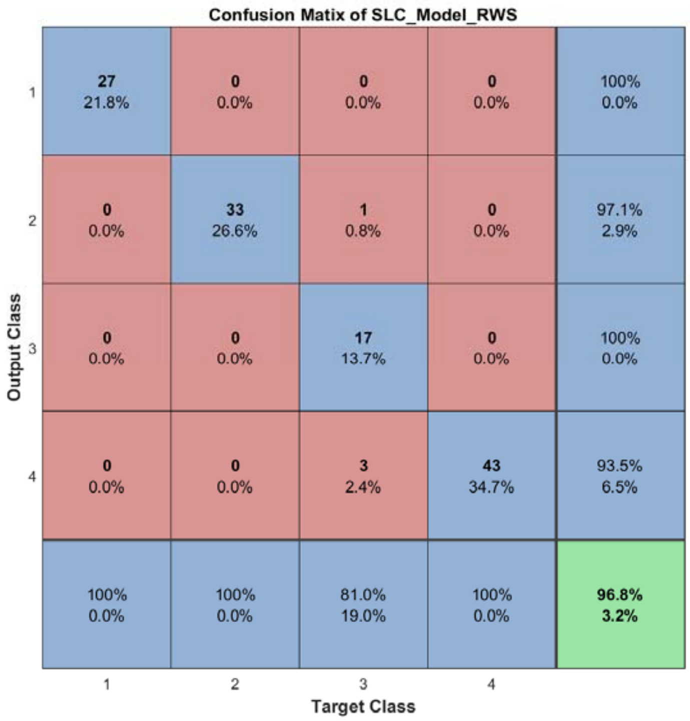


Figure S54: The confusion matrix of ensemble classifier with RWS mode for *Usoskin* dataset.

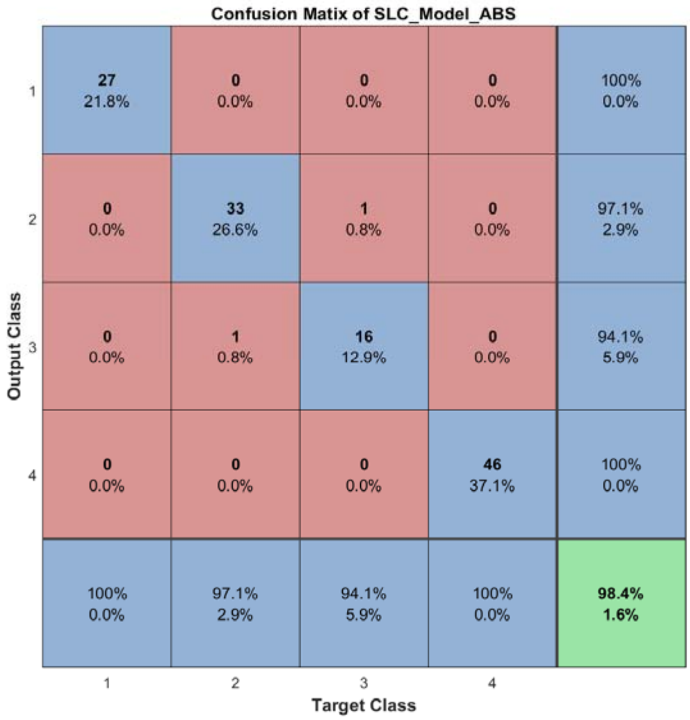


Figure S55: The confusion matrix of ensemble classifier with ABS mode for *Usoskin* dataset.

#### 2.8 The running parameters and output figures for *Zeisel* dataset (9 classes)

Table S6: The running parameters for *Zeisel* dataset.

```
load('SIMLR\Zeisel.mat'); in_X = full(in_X); in_X = log(1+in_X); clear label2;
[in_X_SLC,class_num,slice_tik,binary_mod] = slicematrix(in_X,true_labs,0:0.3:5.1);
[in_X_SLC_diff,in_X_SLC_SRCIstd,slice_bst,slice_vle] = slicediffer(in_X_SLC,slice_tik,9);
[SLC_Model_DIS1,istrain,istest] = slicemethod(in_X_SLC,true_labs,slice_tik,slice_bst,1,'correlation','inverse',5);
[SLC_Model_DIS2,istrain,istest] = slicemethod(in_X_SLC,true_labs,slice_tik,slice_bst,1,'jaccard','inverse',5);
[SLC_Model_DIS3,istrain,istest] = slicemethod(in_X_SLC,true_labs,slice_tik,slice_bst,1,'cosine','inverse',5);
[SLC_Model_All,SLC_Model_All_SMARwit] = sliceweight(SLC_Model_DIS1,SLC_Model_DIS2,SLC_Model_DIS3);
[SLC_Model_RWS,SLC_Model_All_SMESrws] = sliceswitch(SLC_Model_All,SLC_Model_All_SMARwit,'rws',2);
[SLC_Model_ABS,SLC_Model_All_SMESabs] = sliceswitch(SLC_Model_All,SLC_Model_All_SMARwit,'abs',9);
[SLC_Model_FIT1,FIT1_accuracy] = sliceprerws(in_X(istest,:),true_labs(istest),binary_mod,class_num,slice_vle,SLC_Model_RWS);
[SLC_Model_FIT2,FIT2_accuracy] = slicepreabs(in_X(istest,:),true_labs(istest),binary_mod,class_num,slice_vle,SLC_Model_ABS);
```

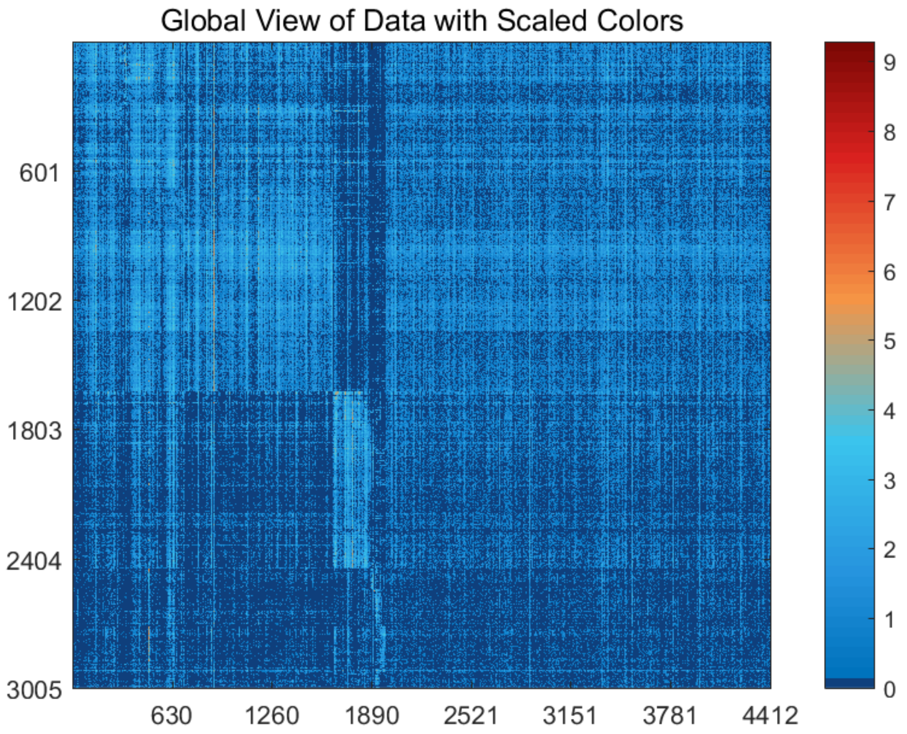


Figure S56: The global view of *Zeisel* dataset with scaled colors.

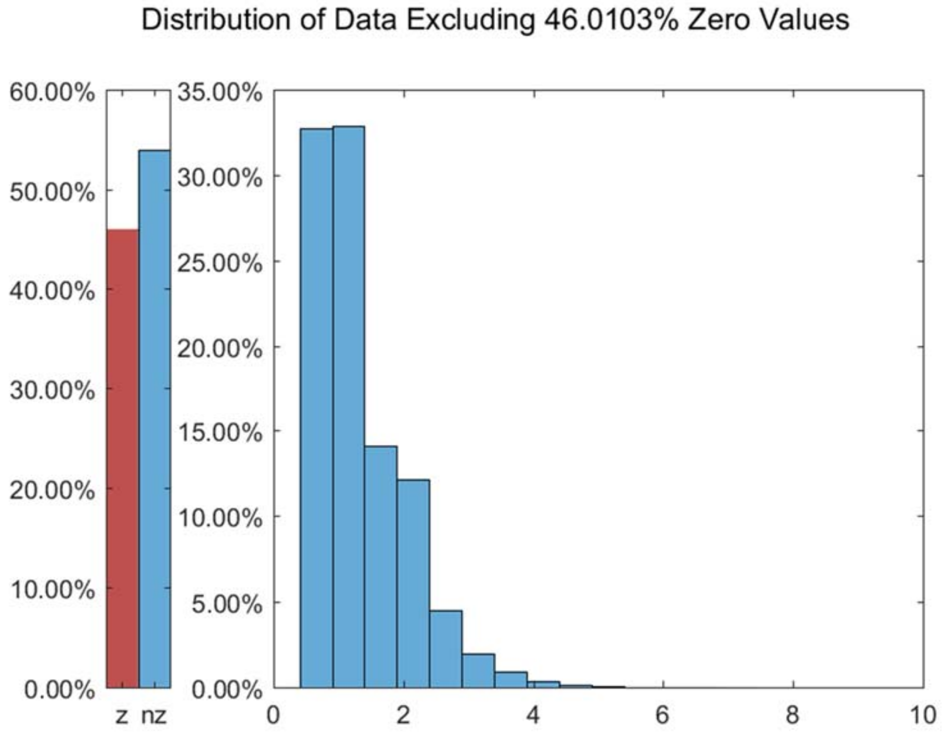


Figure S57: The distribution of *Zeisel* dataset excluding all zero values.

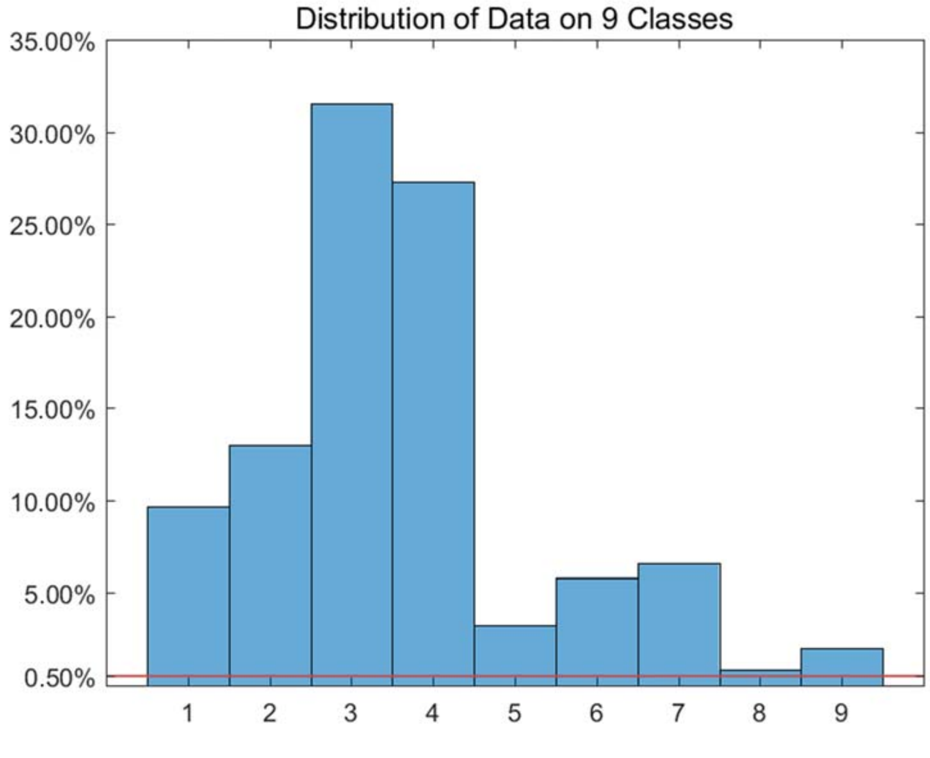


Figure S58: The distribution of *Zeisel* dataset on every classes.

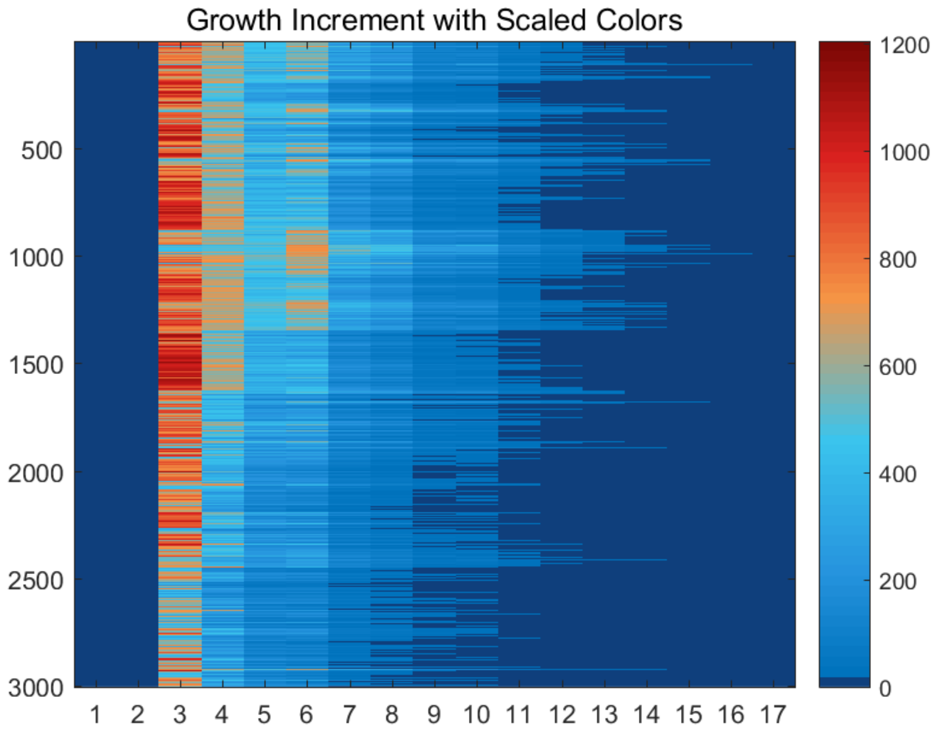


Figure S59: The growth increment with scaled colors for *Zeisel* dataset.

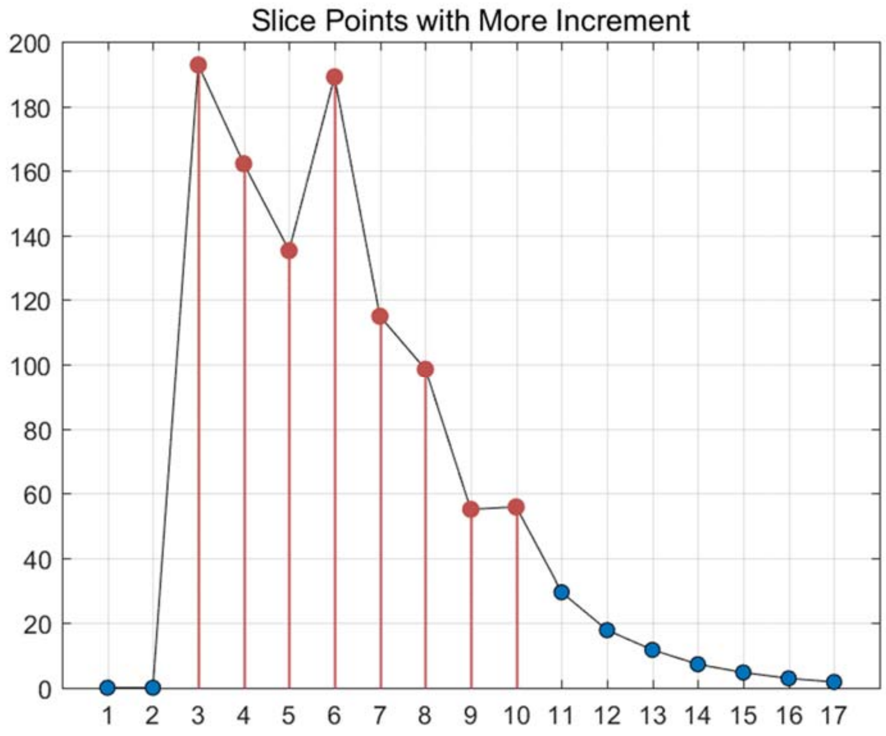


Figure S60: The slice points with more increment for *Zeisel* dataset.

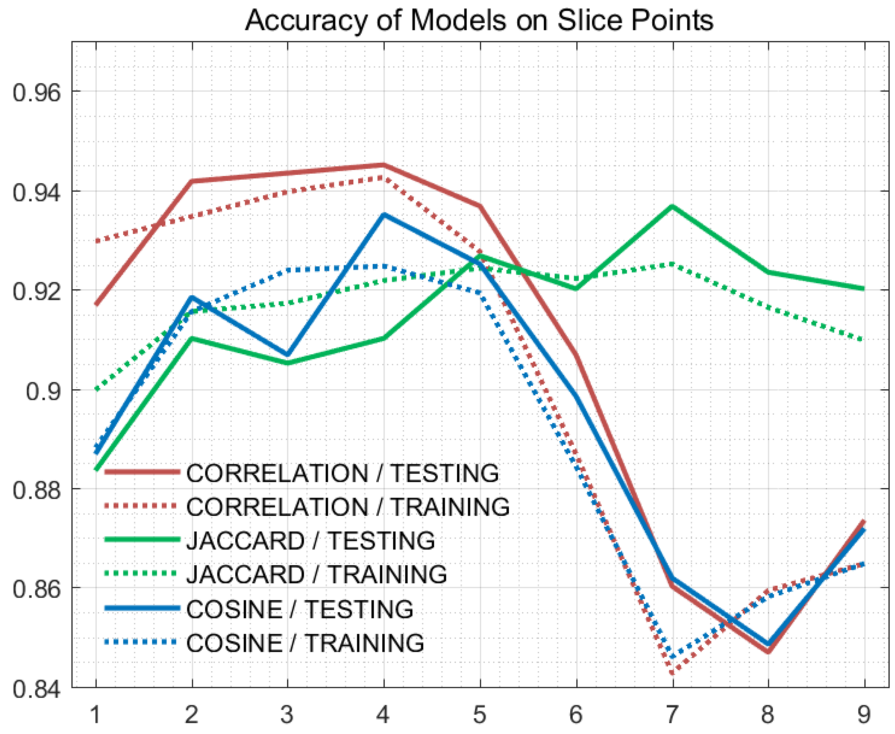


Figure S61: The accuracy of models on every slice points for *Zeisel* dataset.

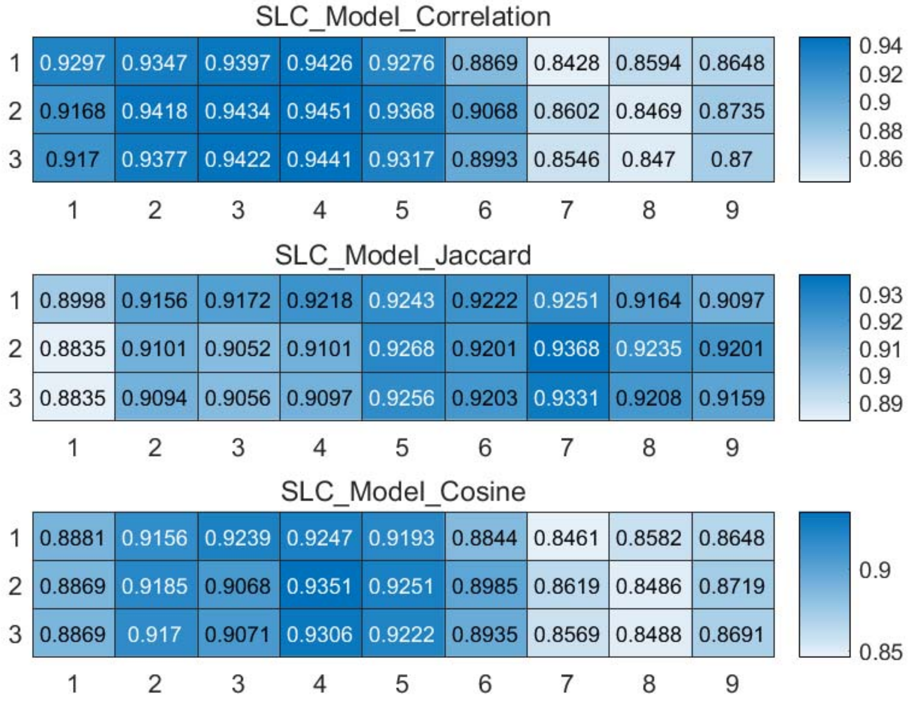


Figure S62: The weighted accuracy on every slice points for *Zeisel* dataset.

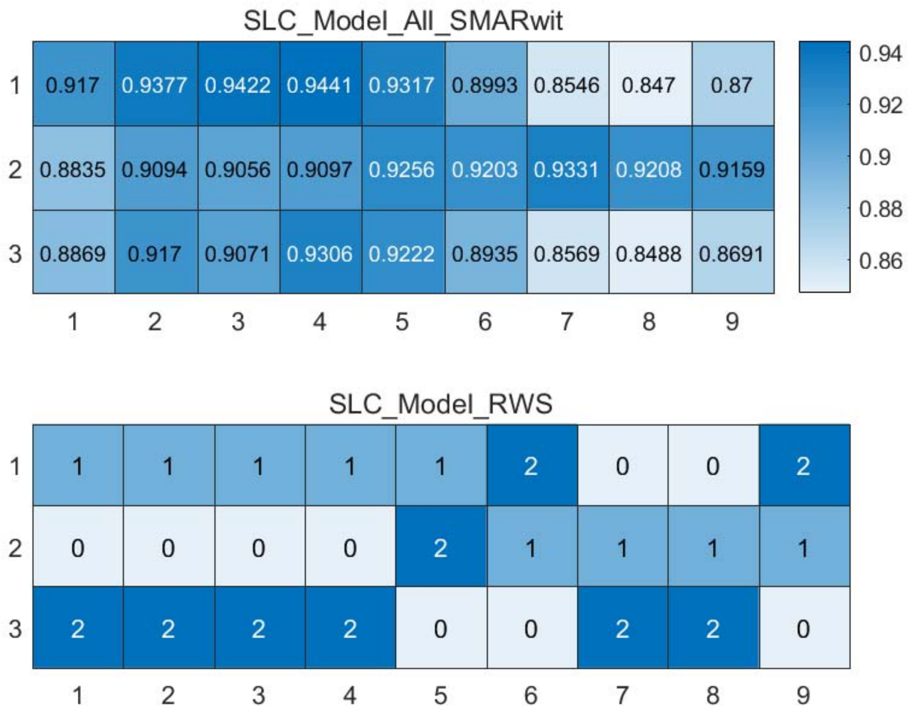


Figure S63: The RWS mode of meta classifiers for *Zeisel* dataset.

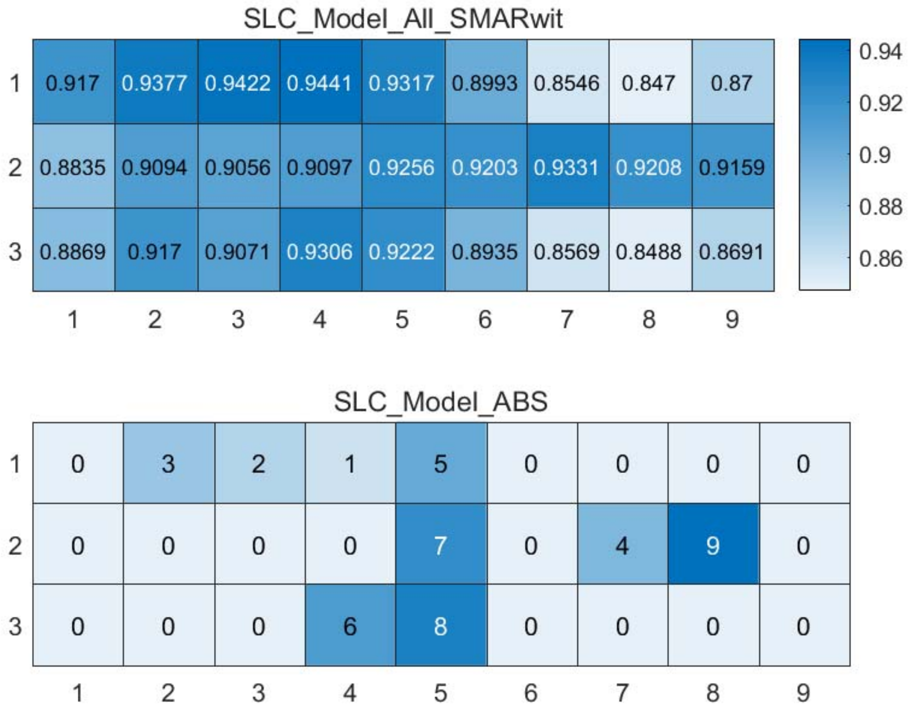


Figure S64: The ABS mode of meta classifiers for *Zeisel* dataset.

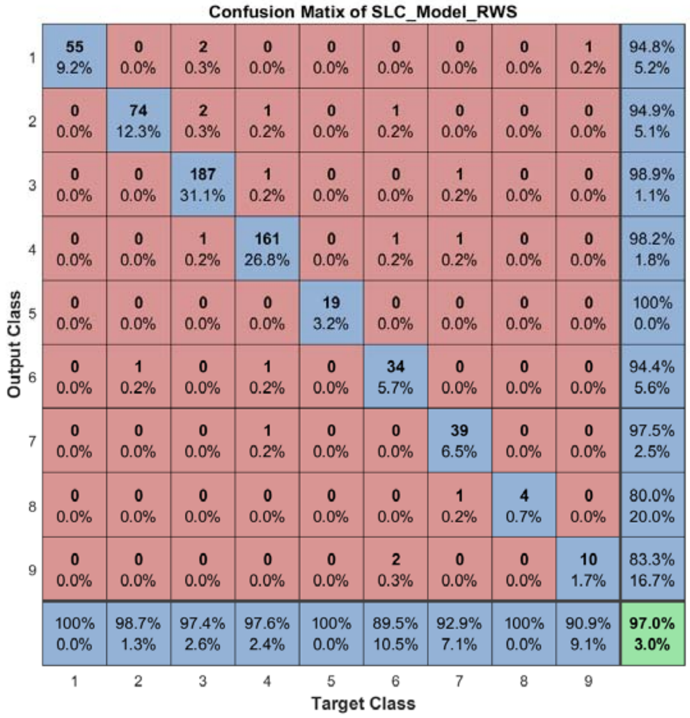


Figure S65: The confusion matrix of ensemble classifier with RWS mode for *Zeisel* dataset.

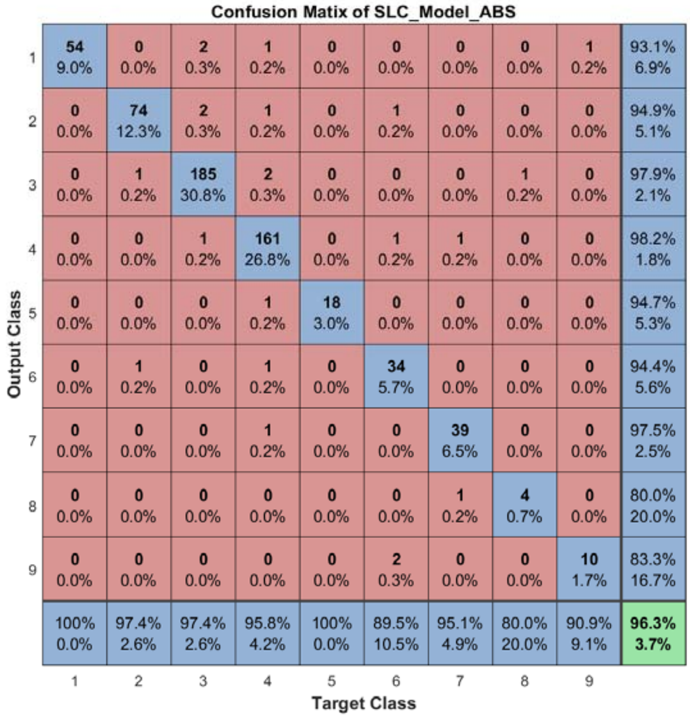


Figure S66: The confusion matrix of ensemble classifier with ABS mode for *Zeisel* dataset.

#### 2.9 The running parameters and output figures for *Zeisel* dataset (48 classes)

Table S7: The running parameters for *Zeisel* dataset.

```
load('SIMLR\Zeisel.mat'); in_X = full(in_X); in_X = log(1+in_X); true_labs = label2'; clear label2;
[in_X_SLC,class_num,slice_tik,binary_mod] = slicematrix(in_X,true_labs,0:0.3:5.1);
[in_X_SLC_diff,in_X_SLC_SRCIstd,slice_bst,slice_vle] = slicediffer(in_X_SLC,slice_tik,9);
[SLC_Model_DIS1,istrain,istest] = slicemethod(in_X_SLC,true_labs,slice_tik,slice_bst,1,'correlation','inverse',3);
[SLC_Model_DIS2,istrain,istest] = slicemethod(in_X_SLC,true_labs,slice_tik,slice_bst,1,'jaccard','inverse',3);
[SLC_Model_DIS3,istrain,istest] = slicemethod(in_X_SLC,true_labs,slice_tik,slice_bst,1,'cosine','inverse',3);
[SLC_Model_All,SLC_Model_All_SMARwit] = sliceweight(SLC_Model_DIS1,SLC_Model_DIS2,SLC_Model_DIS3);
[SLC_Model_RWS,SLC_Model_All_SMESrws] = sliceswitch(SLC_Model_All,SLC_Model_All_SMARwit,'rws',2);
[SLC_Model_ABS,SLC_Model_All_SMESabs] = sliceswitch(SLC_Model_All,SLC_Model_All_SMARwit,'abs',9);
[SLC_Model_FIT1,FIT1_accuracy] = sliceprerws(in_X(istest,:),true_labs(istest),binary_mod,class_num,slice_vle,SLC_Model_RWS);
[SLC_Model_FIT2,FIT2_accuracy] = slicepreabs(in_X(istest,:),true_labs(istest),binary_mod,class_num,slice_vle,SLC_Model_ABS);
```

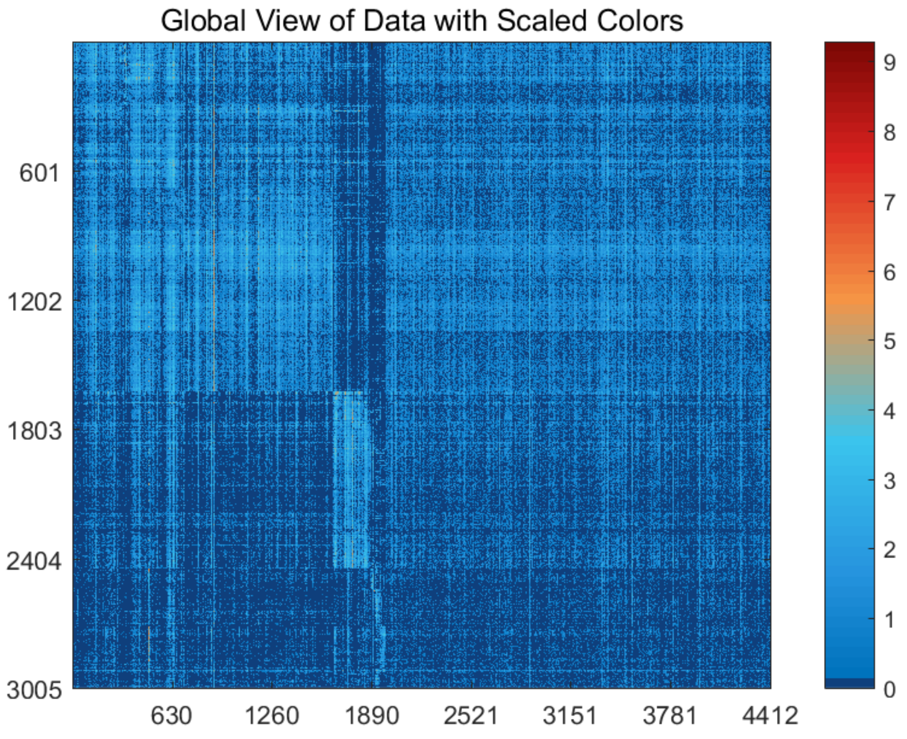


Figure S67: The global view of *Zeisel* dataset with scaled colors.

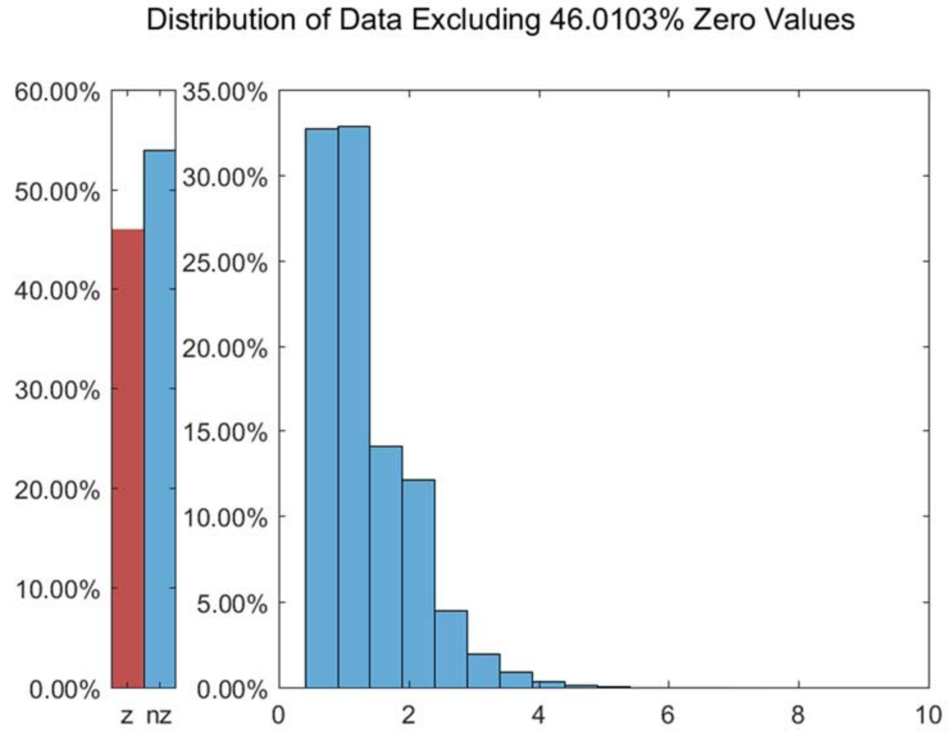


Figure S68: The distribution of *Zeisel* dataset excluding all zero values.

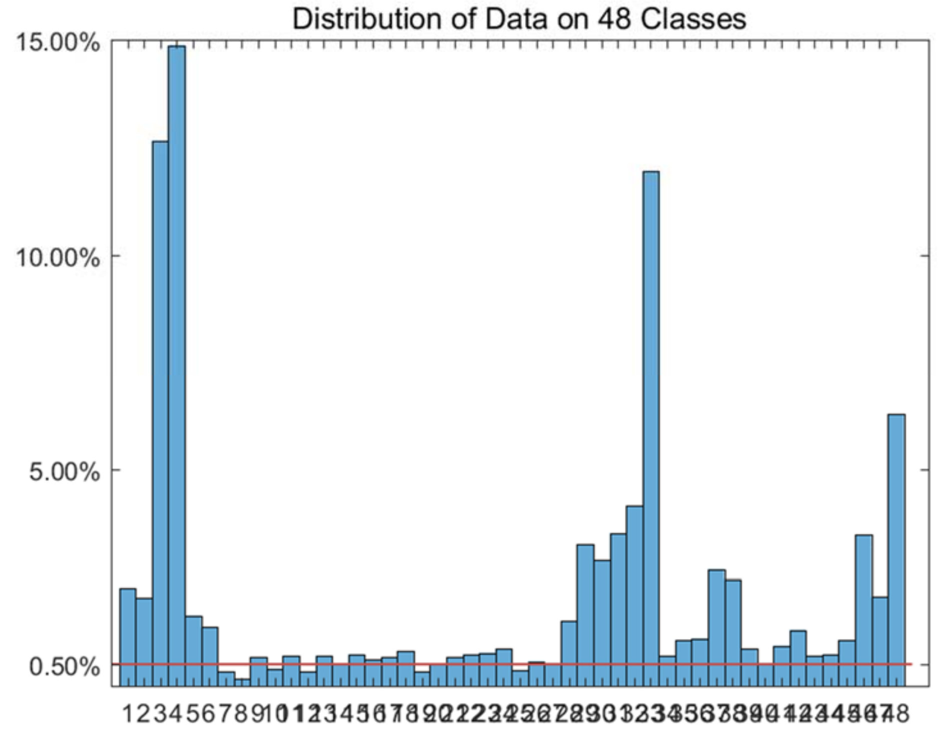


Figure S69: The distribution of *Zeisel* dataset on every classes.

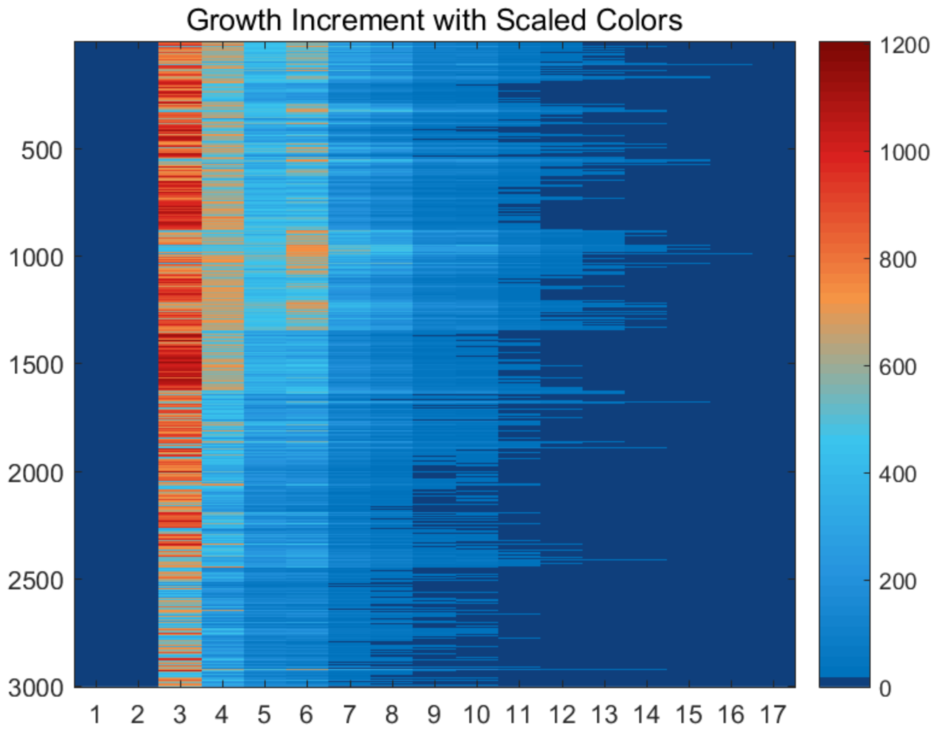


Figure S70: The growth increment with scaled colors for *Zeisel* dataset.

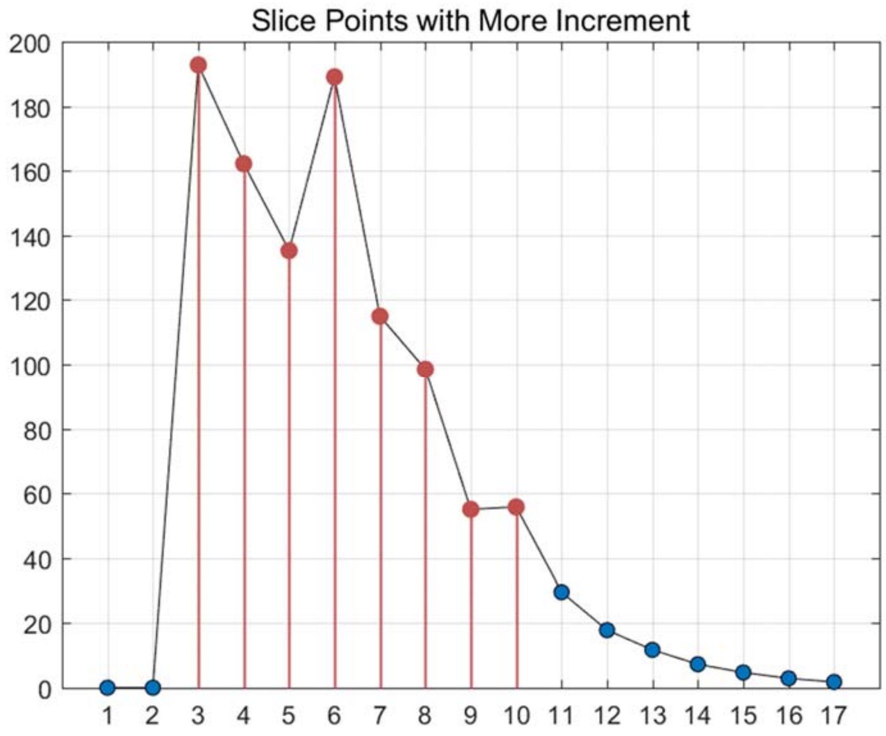


Figure S71: The slice points with more increment for *Zeisel* dataset.

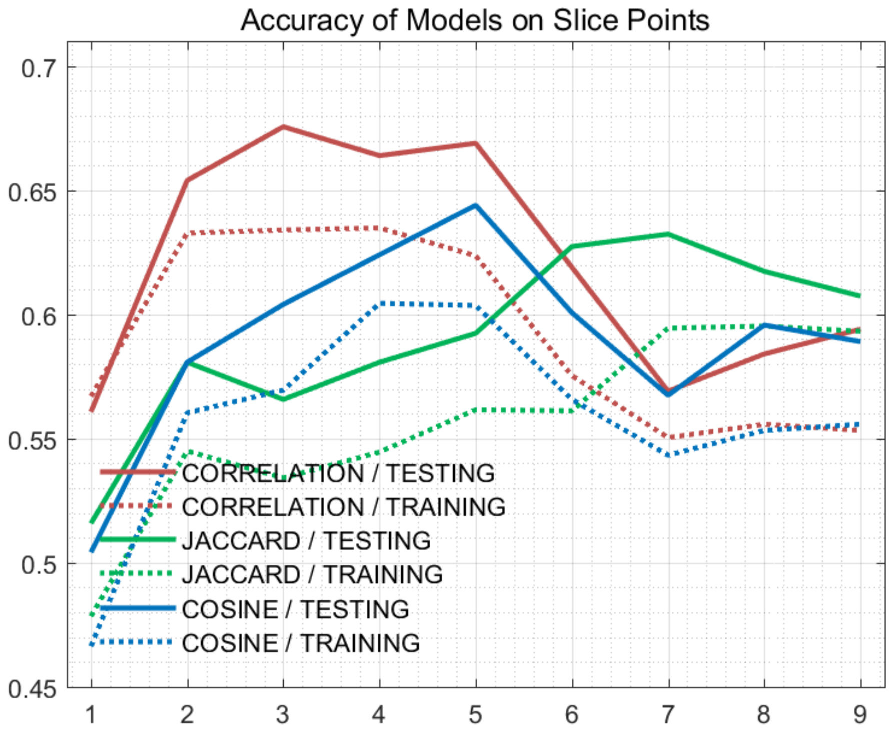


Figure S72: The accuracy of models on every slice points for *Zeisel* dataset.

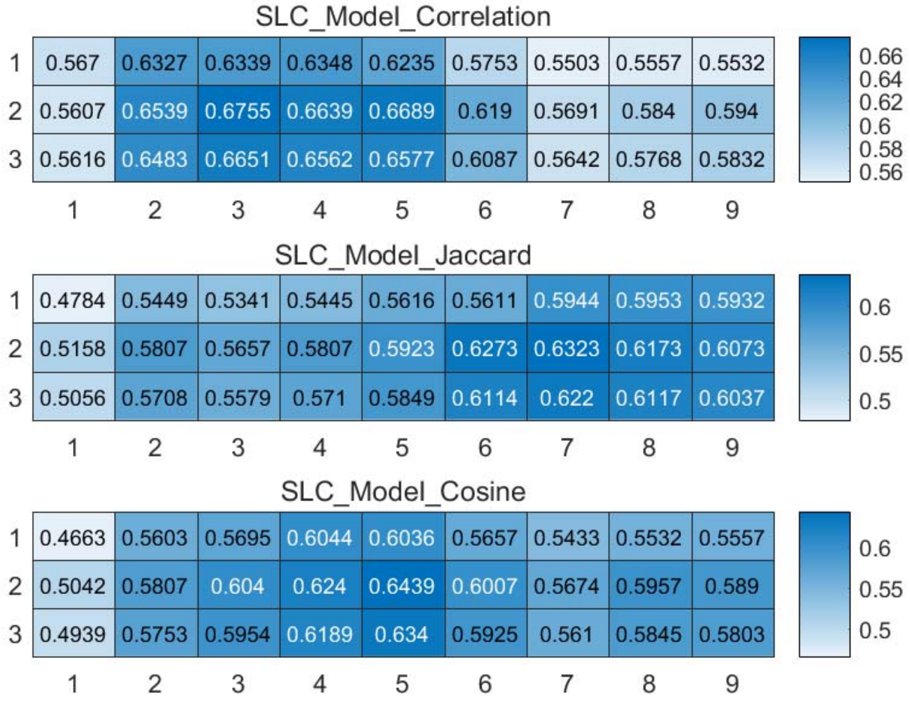


Figure S73: The weighted accuracy on every slice points for *Zeisel* dataset.

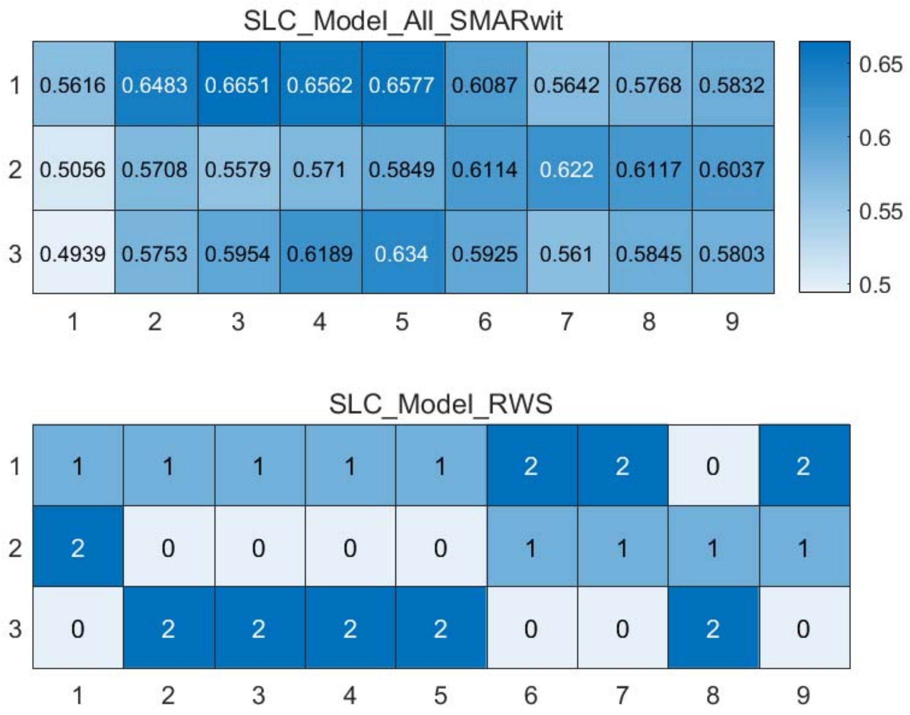


Figure S74: The RWS mode of meta classifiers for *Zeisel* dataset.

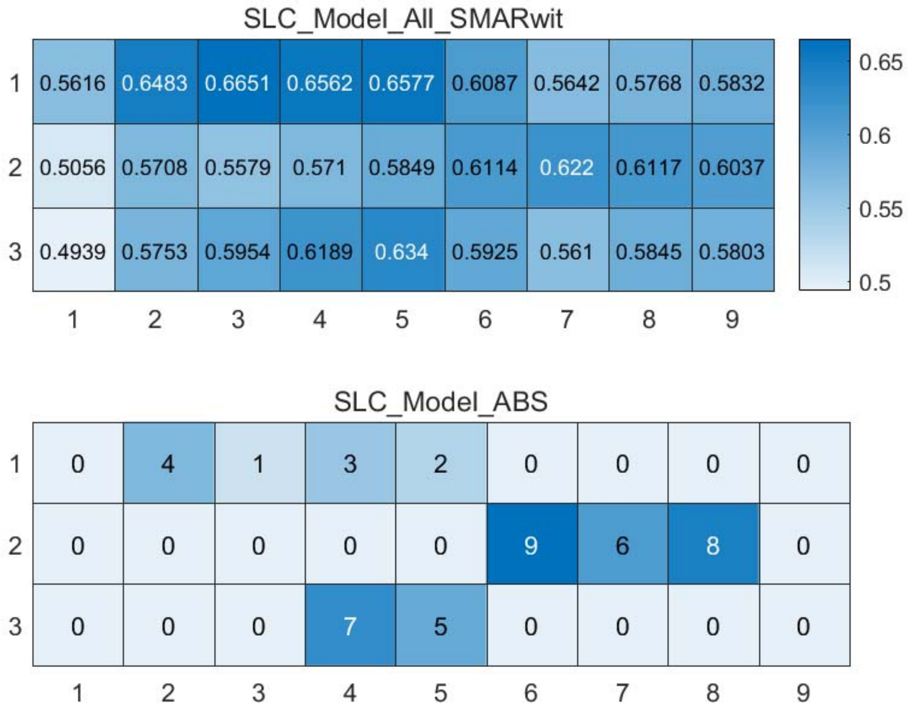


Figure S75: The ABS mode of meta classifiers for *Zeisel* dataset.

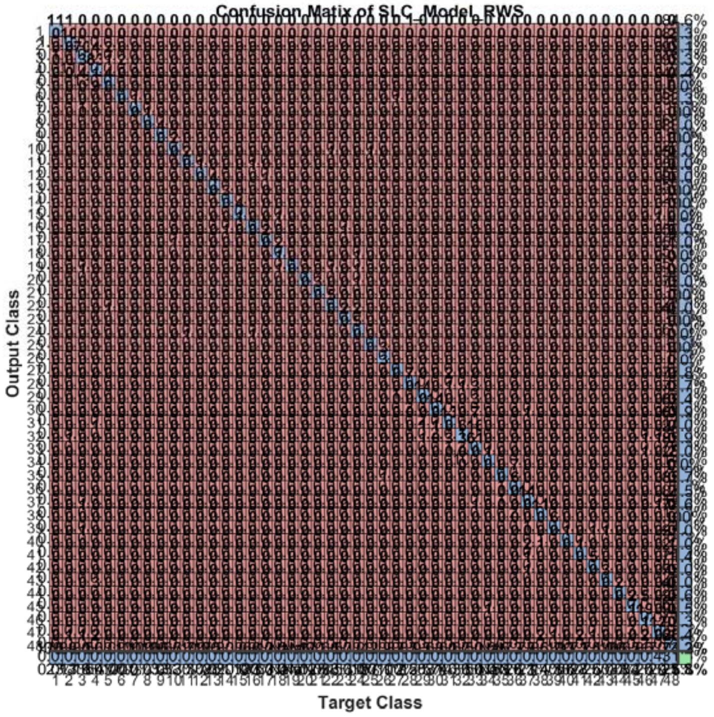


Figure S76: The confusion matrix of ensemble classifier with RWS mode for *Zeisel* dataset. (*Note*: the confusion matrix is too large to display completely in one figure)

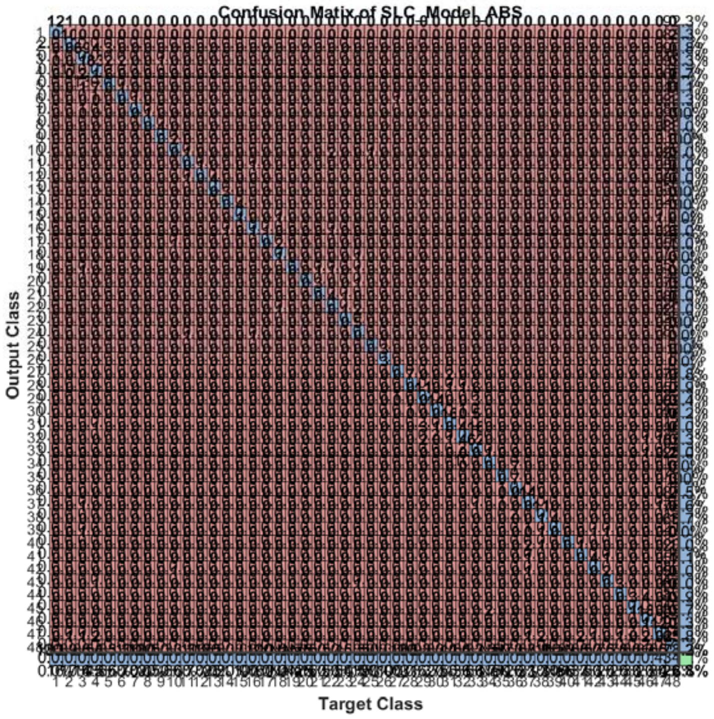


Figure S77: The confusion matrix of ensemble classifier with ABS mode for *Zeisel* dataset. (*Note*: the confusion matrix is too large to display completely in one figure)

#### 2.10 The running parameters and output figures for *Data_Buettner* dataset

Table S8: The running parameters for *Data_Buettner* dataset.

```
load('MPSSC\Data_Buettner.mat'); in_X = 0.2*in_X;
[in_X_SLC,class_num,slice_tik,binary_mod] = slicematrix(in_X,true_labs,0:0.1:2.5);
[in_X_SLC_diff,in_X_SLC_SRCIstd,slice_bst,slice_vle] = slicediffer(in_X_SLC,slice_tik,9);
[SLC_Model_DIS1,istrain,istest] = slicemethod(in_X_SLC,true_labs,slice_tik,slice_bst,5,'correlation','inverse',5);
[SLC_Model_DIS2,istrain,istest] = slicemethod(in_X_SLC,true_labs,slice_tik,slice_bst,5,'jaccard','inverse',5);
[SLC_Model_DIS3,istrain,istest] = slicemethod(in_X_SLC,true_labs,slice_tik,slice_bst,5,'cosine','inverse',5);
[SLC_Model_All,SLC_Model_All_SMARwit] = sliceweight(SLC_Model_DIS1,SLC_Model_DIS2,SLC_Model_DIS3);
[SLC_Model_RWS,SLC_Model_All_SMESrws] = sliceswitch(SLC_Model_All,SLC_Model_All_SMARwit,'rws',2);
[SLC_Model_ABS,SLC_Model_All_SMESabs] = sliceswitch(SLC_Model_All,SLC_Model_All_SMARwit,'abs',9);
[SLC_Model_FIT1,FIT1_accuracy] = sliceprerws(in_X(istest,:),true_labs(istest),binary_mod,class_num,slice_vle,SLC_Model_RWS);
[SLC_Model_FIT2,FIT2_accuracy] = slicepreabs(in_X(istest,:),true_labs(istest),binary_mod,class_num,slice_vle,SLC_Model_ABS);
```

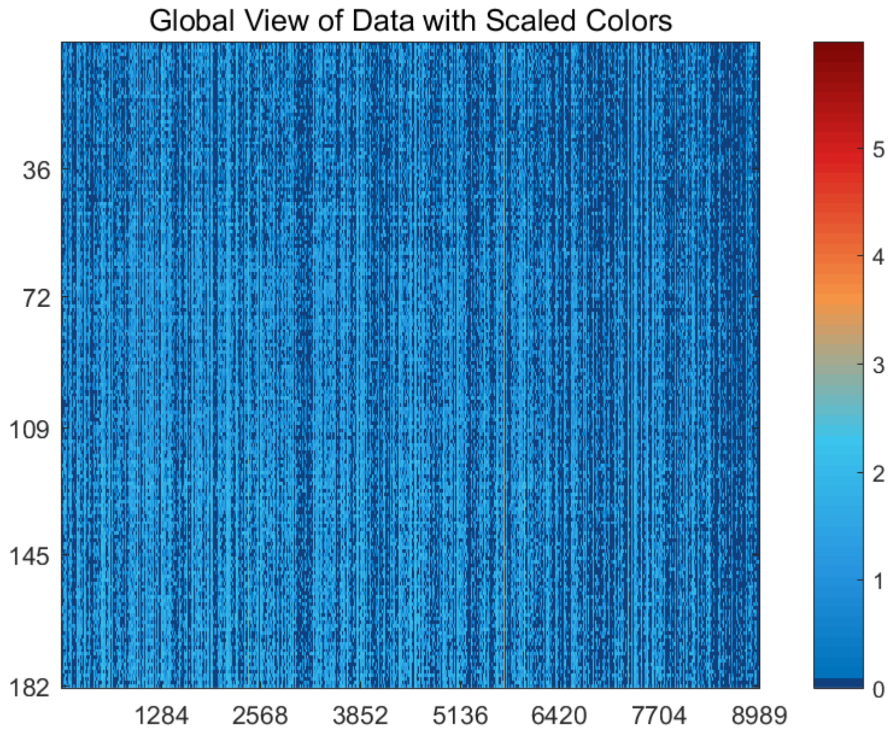


Figure S78: The global view of *Data_Buettner* dataset with scaled colors.

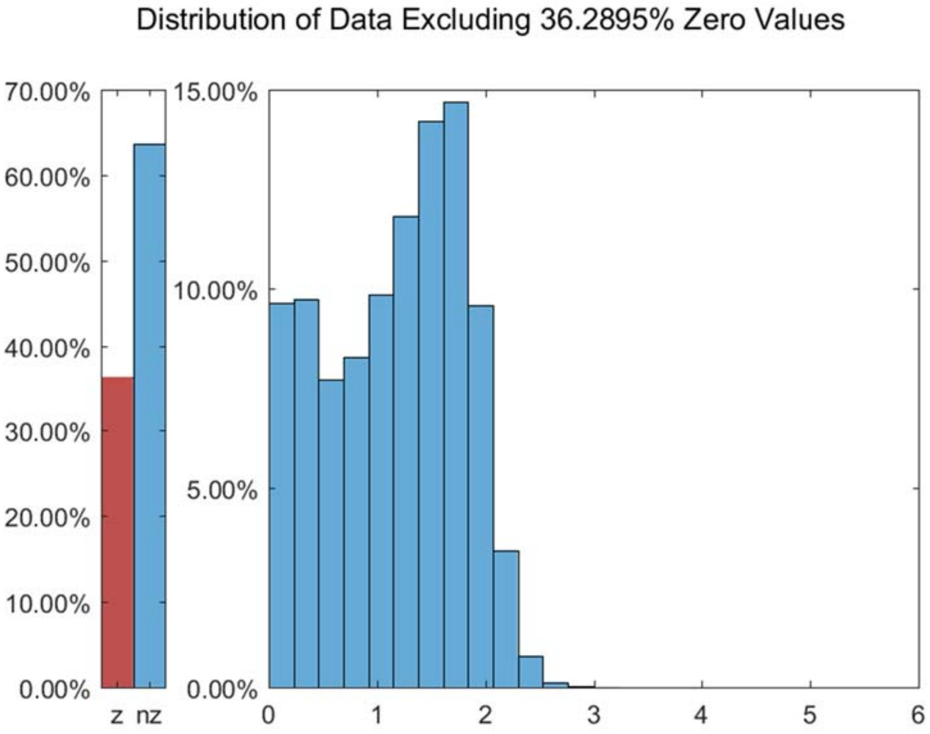


Figure S79: The distribution of *Data_Buettner* dataset excluding all zero values.

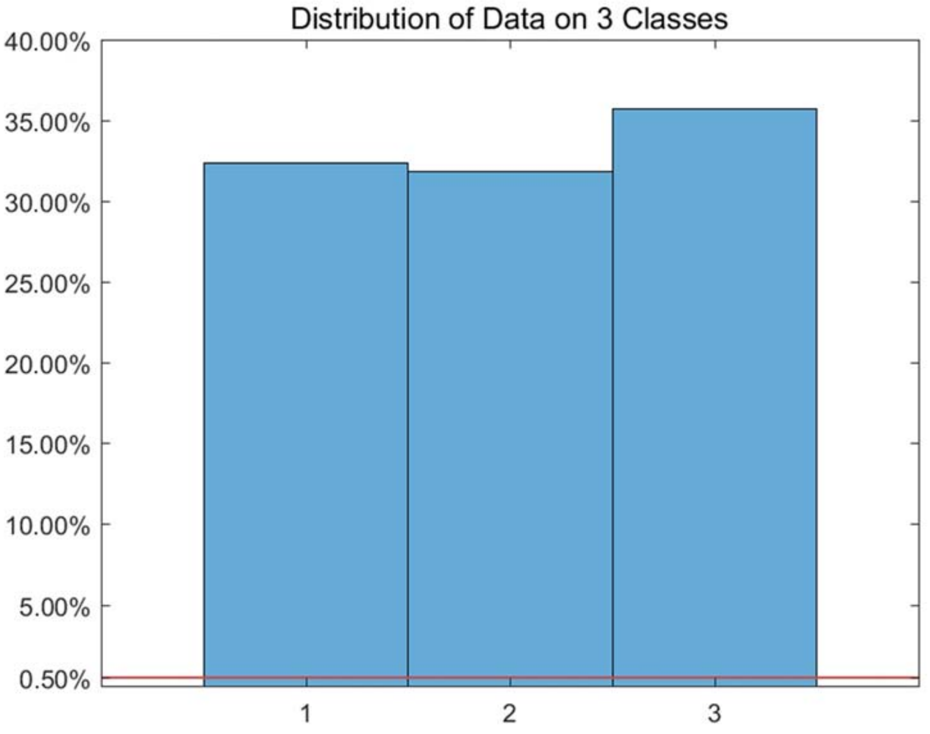


Figure S80: The distribution of *Data_Buettner* dataset on every classes.

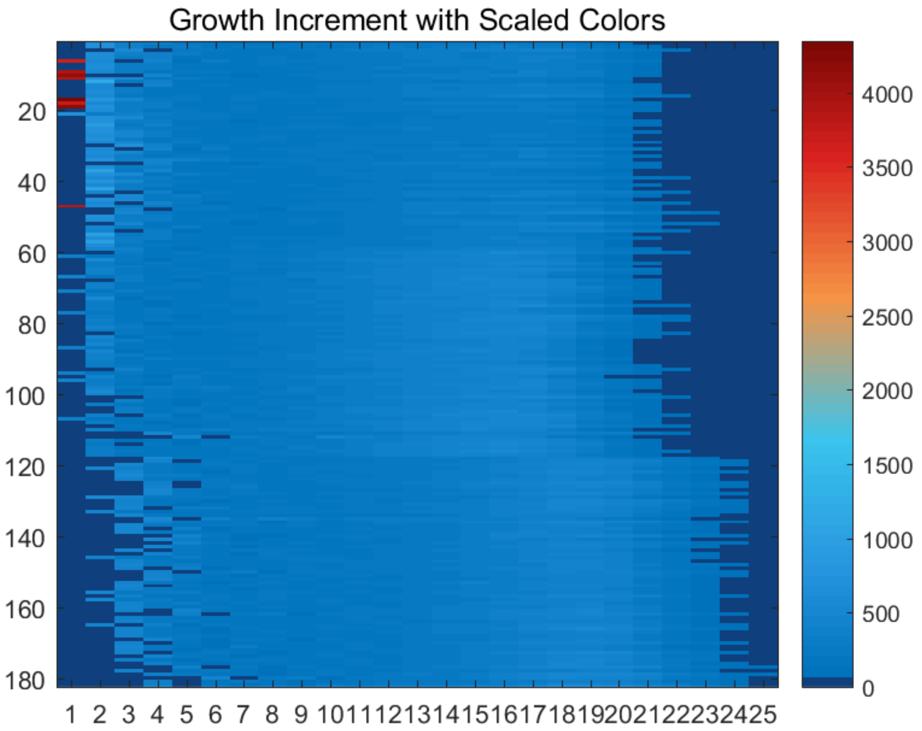


Figure S81: The growth increment with scaled colors for *Data_Buettner* dataset.

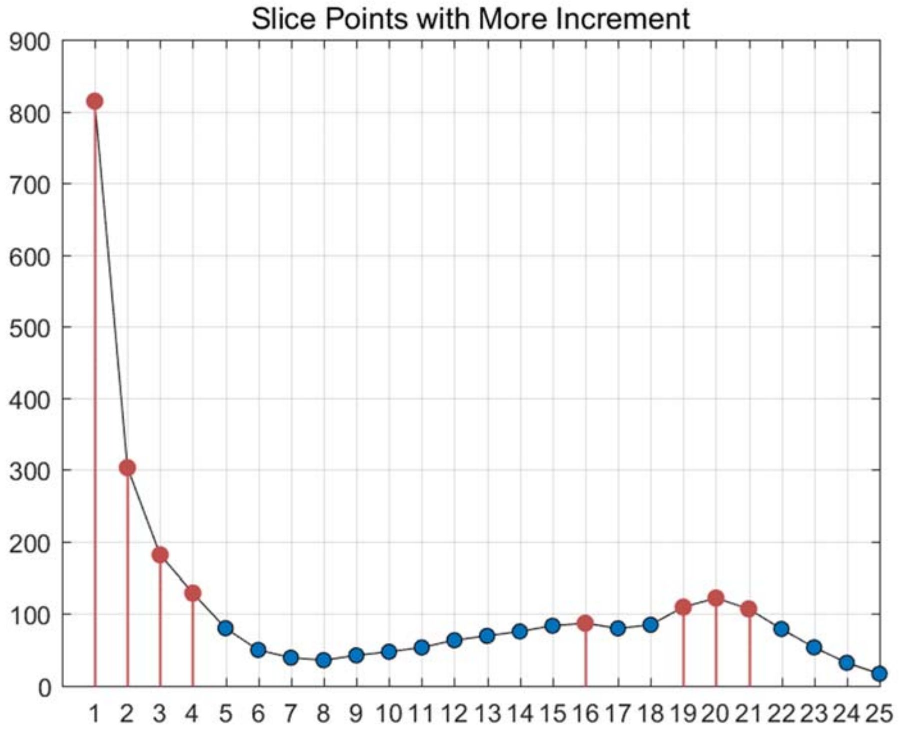


Figure S82: The slice points with more increment for *Data_Buettner* dataset.

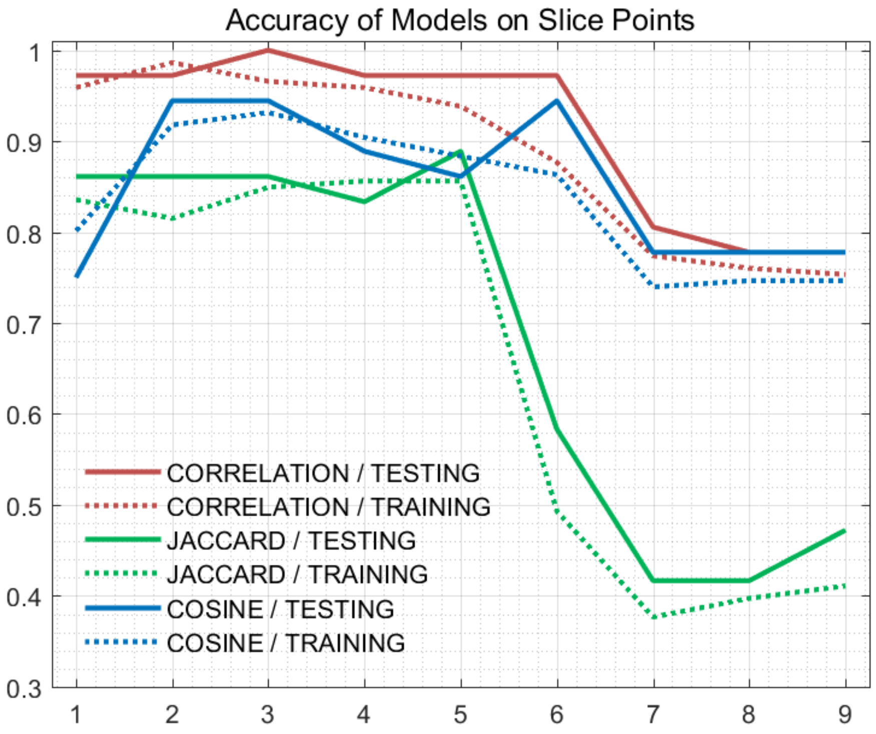


Figure S83: The accuracy of models on every slice points for *Data_Buettner* dataset.

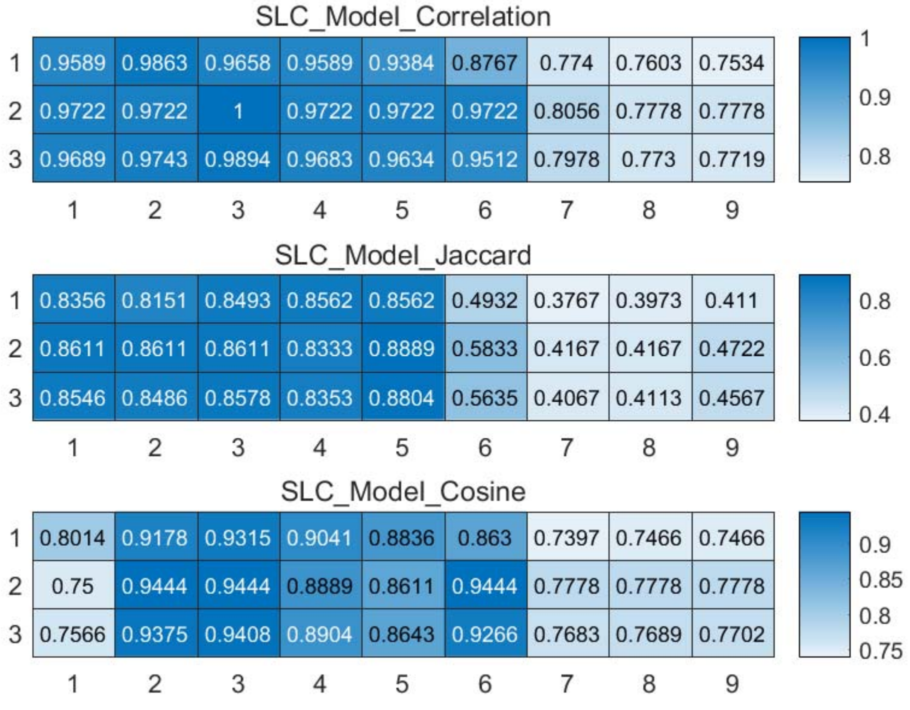


Figure S84: The weighted accuracy on every slice points for *Data_Buettner* dataset.

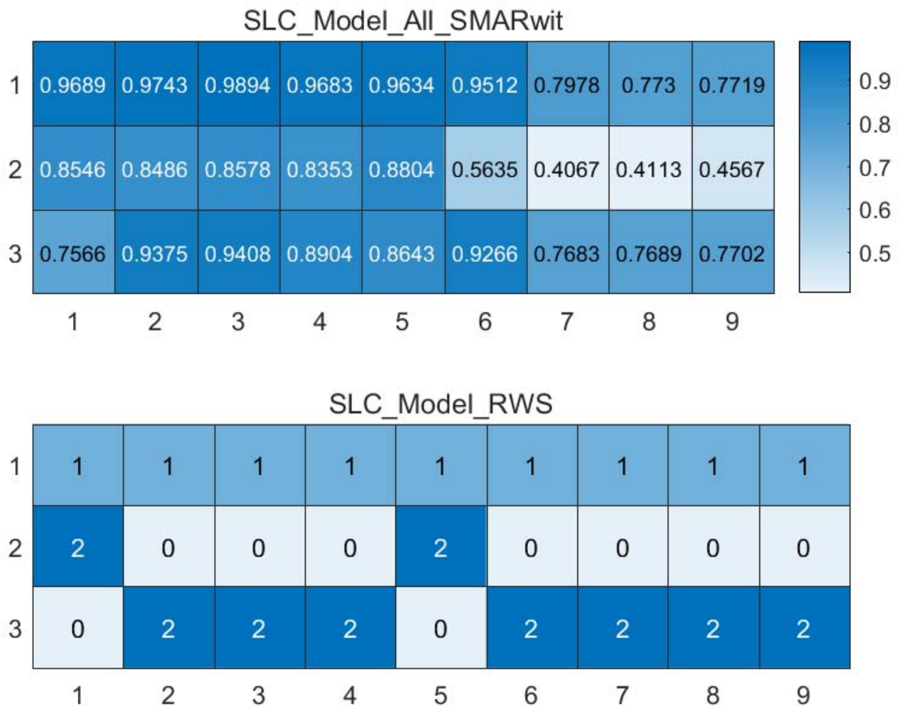


Figure S85: The RWS mode of meta classifiers for *Data_Buettner* dataset.

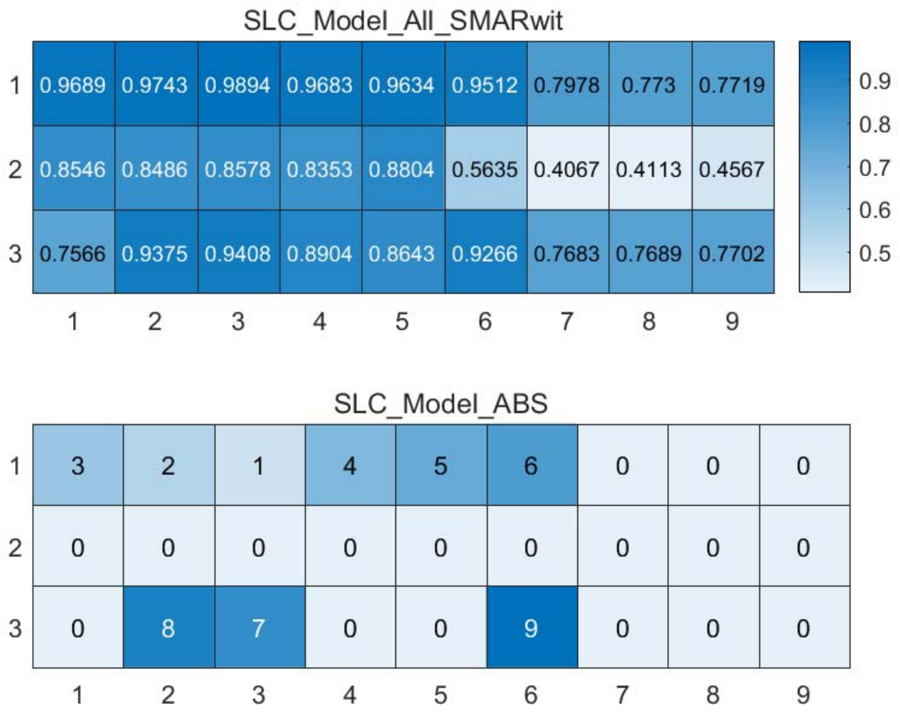


Figure S86: The ABS mode of meta classifiers for *Data_Buettner* dataset.

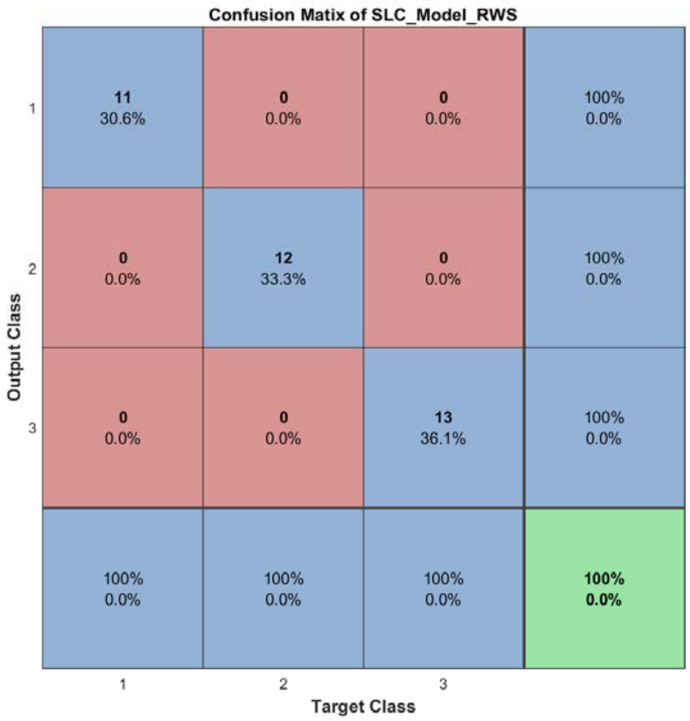


Figure S87: The confusion matrix of ensemble classifier with RWS mode for *Data_Buettner* dataset.

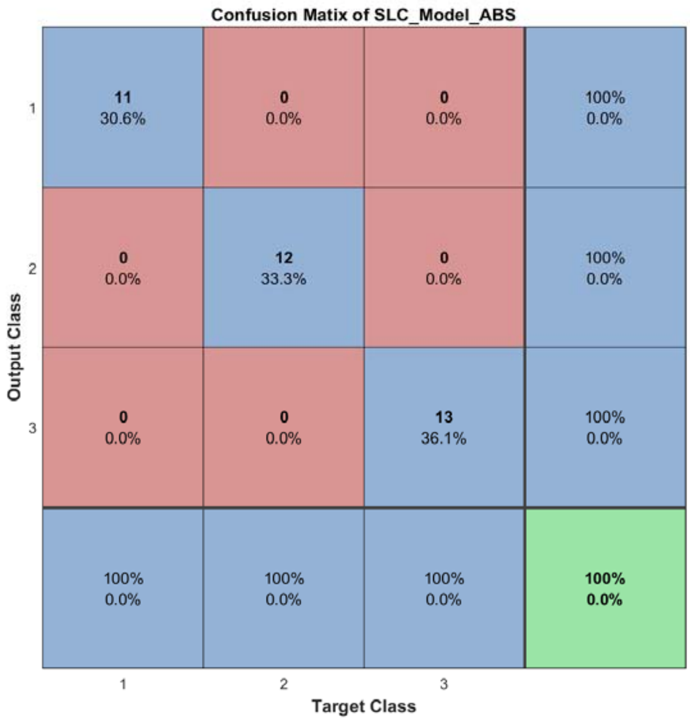


Figure S88: The confusion matrix of ensemble classifier with ABS mode for *Data_Buettner* dataset.

#### 2.11 The running parameters and output figures for *Data_Deng* dataset

Table S9: The running parameters for *Data_Deng* dataset.

```
load('MPSSC\Data_Deng.mat'); in_X = in_X*0.4;
[in_X_SLC,class_num,slice_tik,binary_mod] = slicematrix(in_X,true_labs,0:0.1:4.3);
[in_X_SLC_diff,in_X_SLC_SRCIstd,slice_bst,slice_vle] = slicediffer(in_X_SLC,slice_tik,9);
[SLC_Model_DIS1,istrain,istest] = slicemethod(in_X_SLC,true_labs,slice_tik,slice_bst,5,'correlation','inverse',3);
[SLC_Model_DIS2,istrain,istest] = slicemethod(in_X_SLC,true_labs,slice_tik,slice_bst,5,'jaccard','inverse',3);
[SLC_Model_DIS3,istrain,istest] = slicemethod(in_X_SLC,true_labs,slice_tik,slice_bst,5,'cosine','inverse',3);
[SLC_Model_All,SLC_Model_All_SMARwit] = sliceweight(SLC_Model_DIS1,SLC_Model_DIS2,SLC_Model_DIS3);
[SLC_Model_RWS,SLC_Model_All_SMESrws] = sliceswitch(SLC_Model_All,SLC_Model_All_SMARwit,'rws',2);
[SLC_Model_ABS,SLC_Model_All_SMESabs] = sliceswitch(SLC_Model_All,SLC_Model_All_SMARwit,'abs',9);
[SLC_Model_FIT1,FIT1_accuracy] = sliceprerws(in_X(istest,:),true_labs(istest),binary_mod,class_num,slice_vle,SLC_Model_RWS);
[SLC_Model_FIT2,FIT2_accuracy] = slicepreabs(in_X(istest,:),true_labs(istest),binary_mod,class_num,slice_vle,SLC_Model_ABS);
```

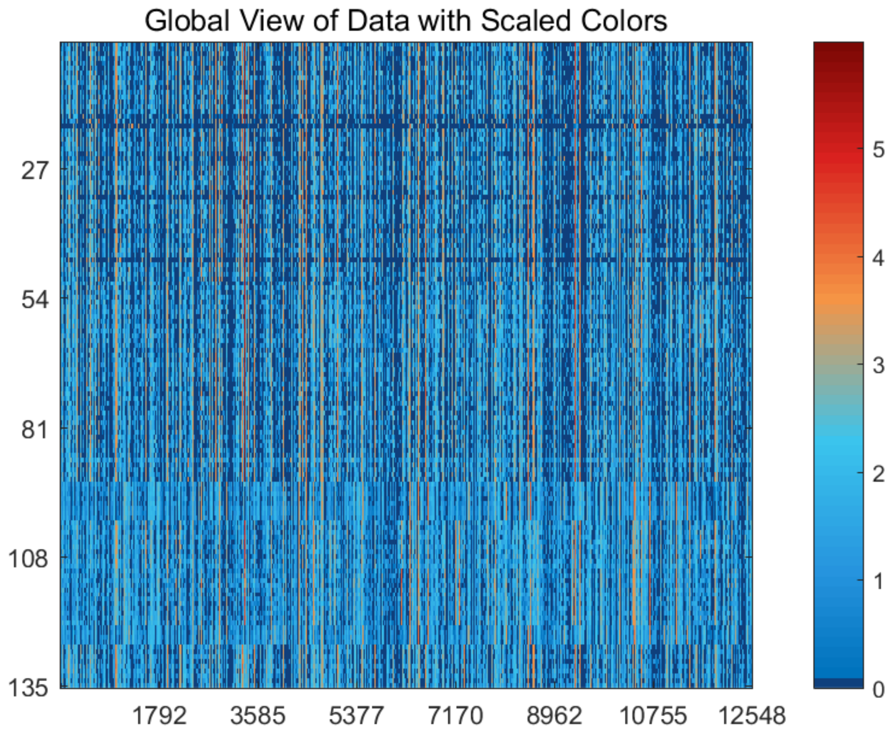


Figure S89: The global view of *Data_Deng* dataset with scaled colors.

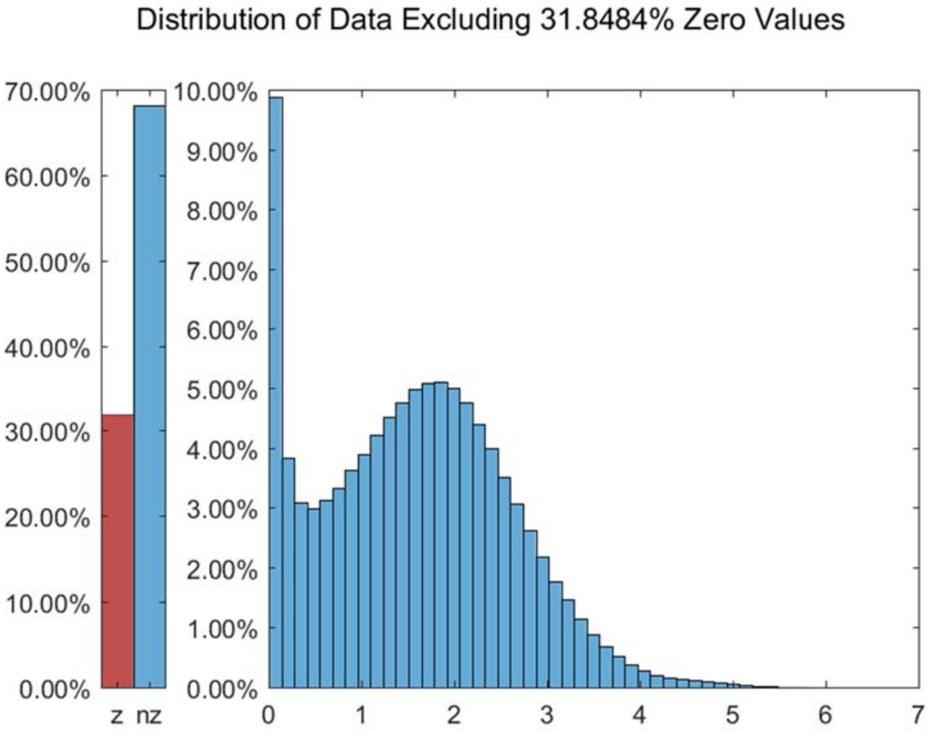


Figure S90: The distribution of *Data_Deng* dataset excluding all zero values.

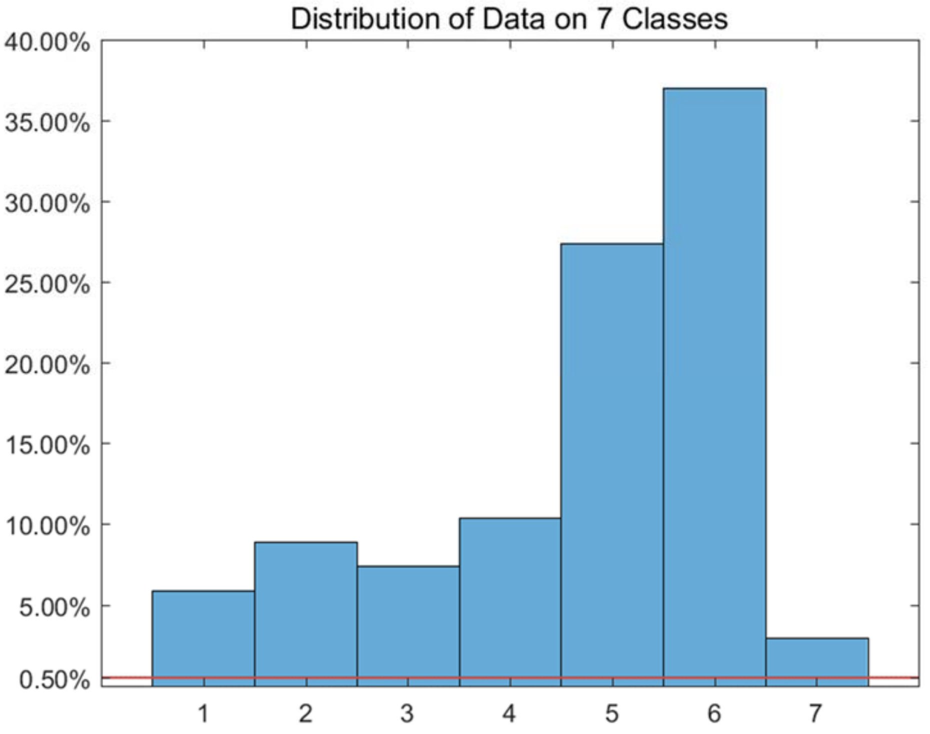


Figure S91: The distribution of *Data_Deng* dataset on every classes.

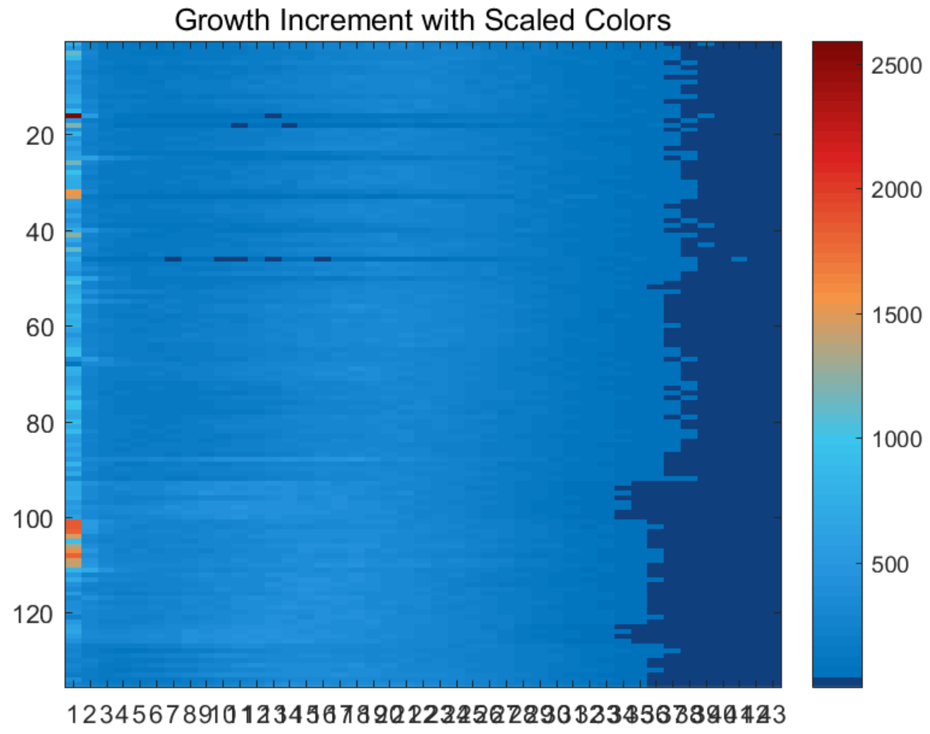


Figure S92: The growth increment with scaled colors for *Data_Deng* dataset.

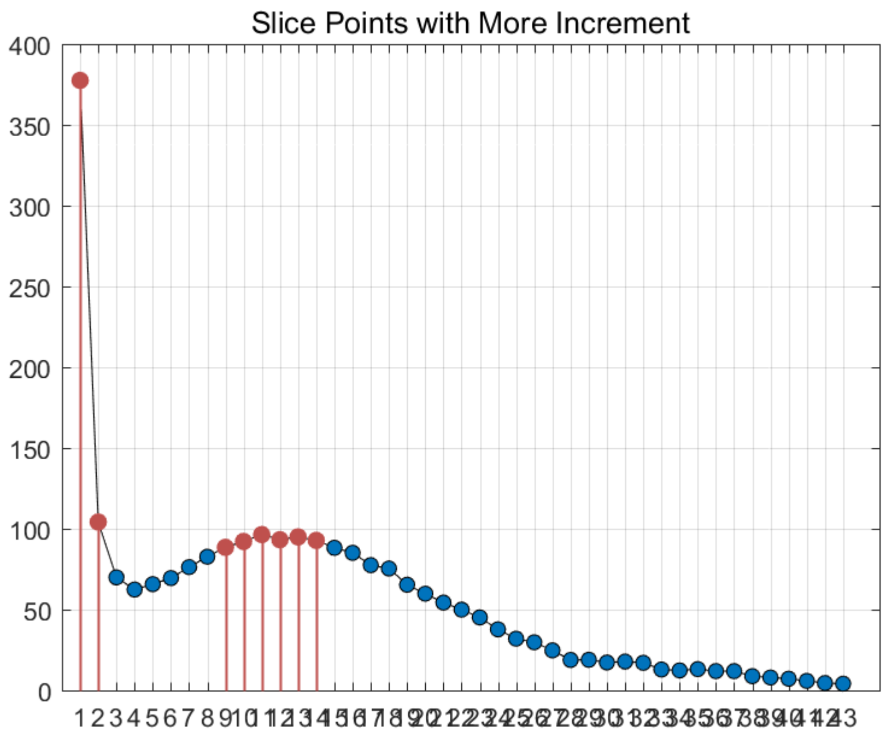


Figure S93: The slice points with more increment for *Data_Deng* dataset.

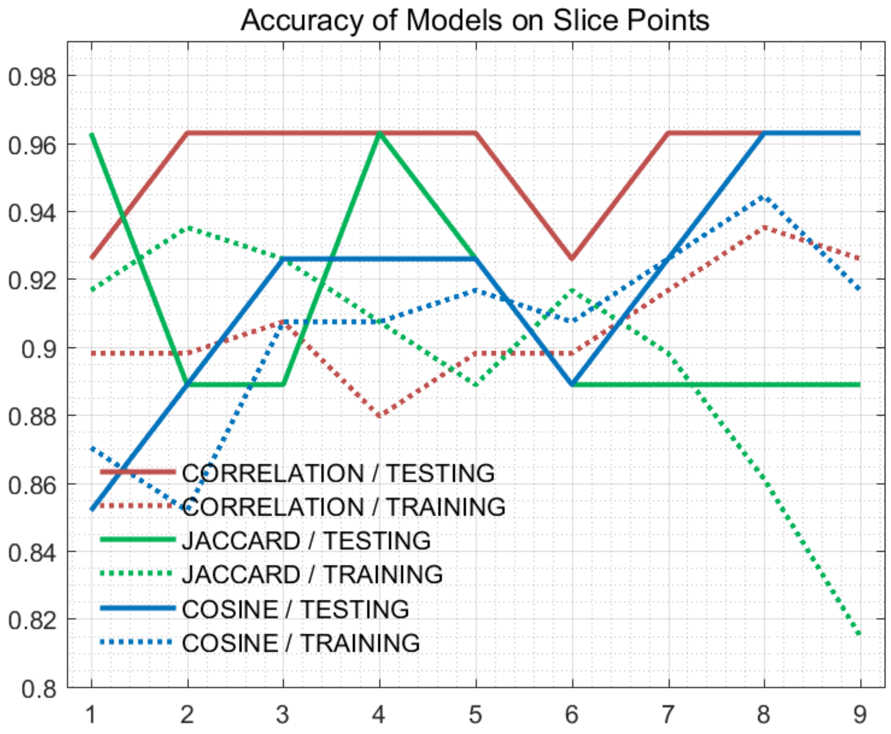


Figure S94: The accuracy of models on every slice points for *Data_Deng* dataset.

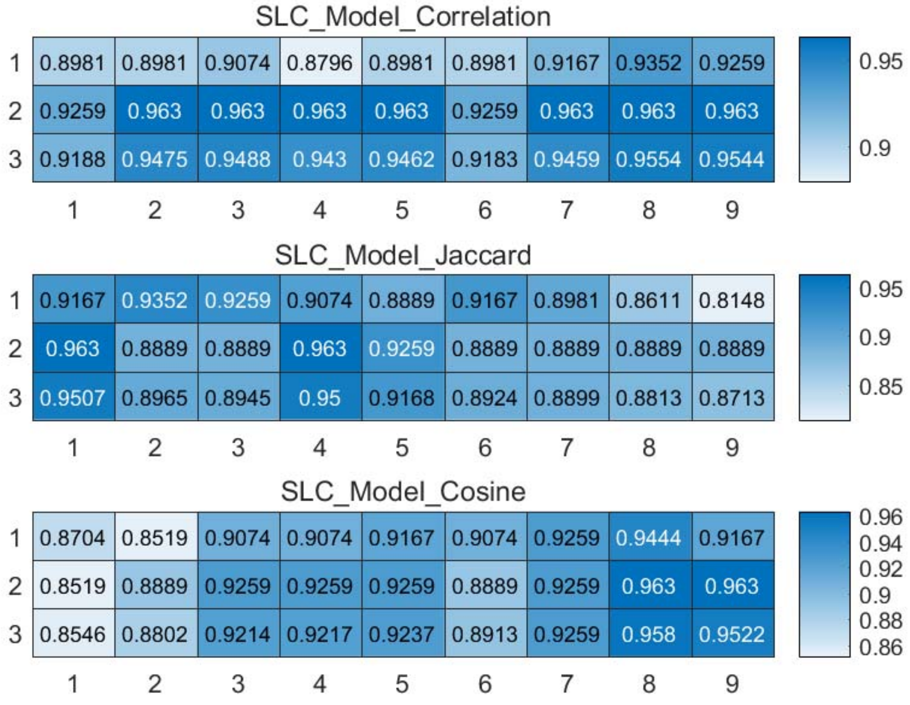


Figure S95: The weighted accuracy on every slice points for *Data_Deng* dataset.

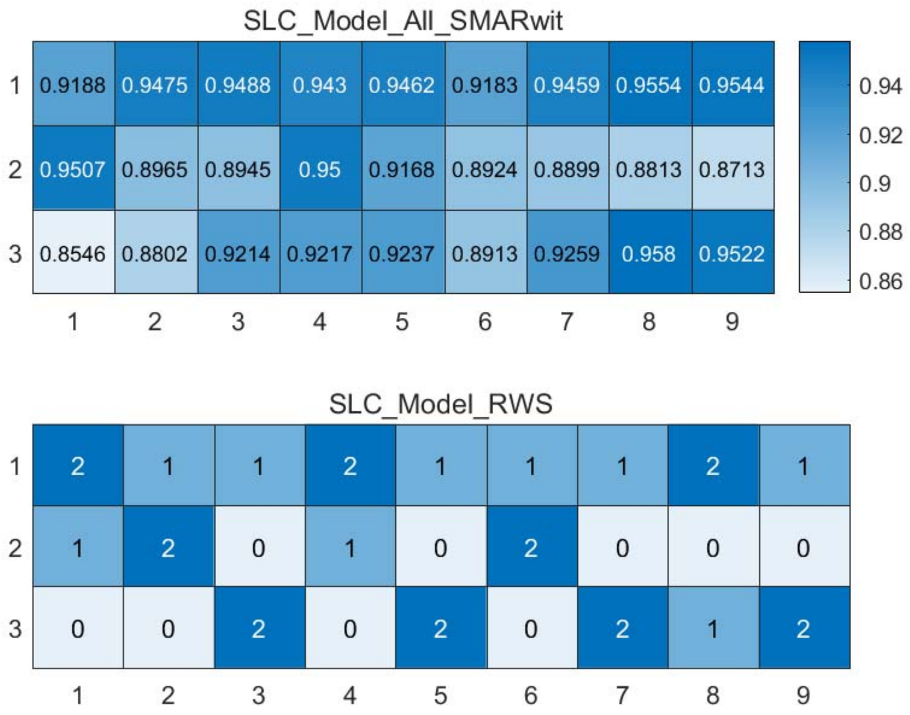


Figure S96: The RWS mode of meta classifiers for *Data_Deng* dataset.

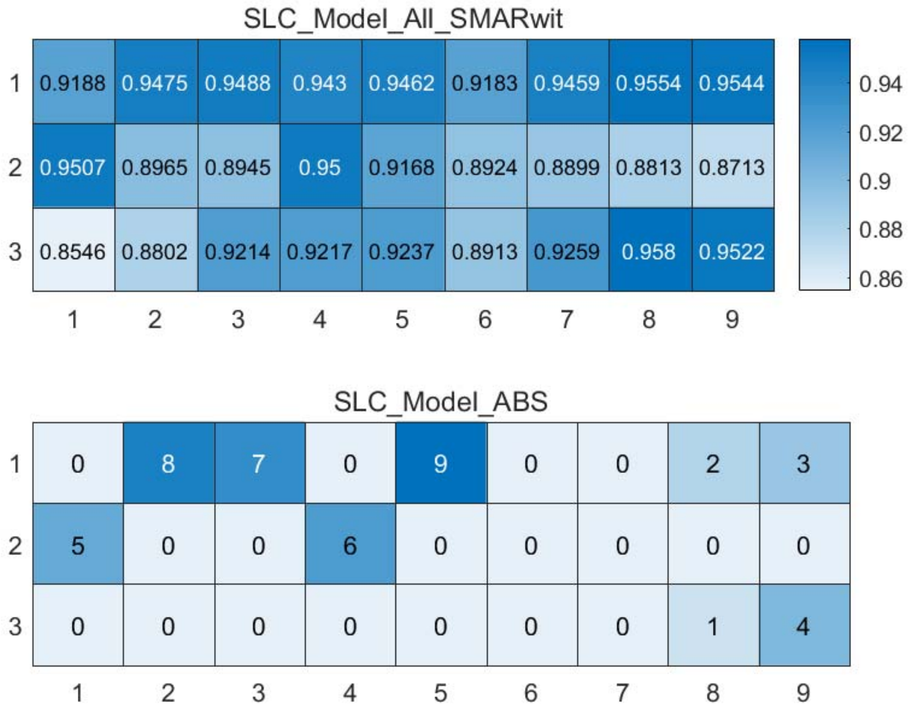


Figure S97: The ABS mode of meta classifiers for *Data_Deng* dataset.

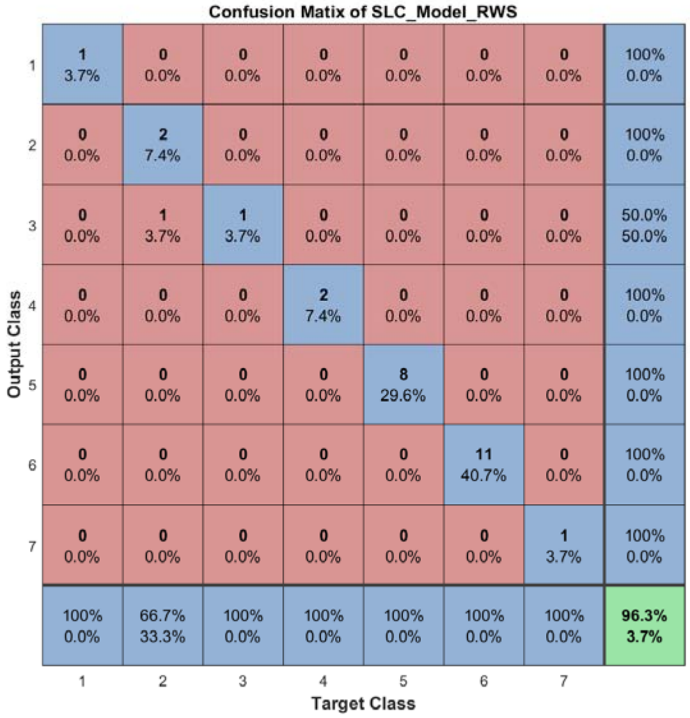


Figure S98: The confusion matrix of ensemble classifier with RWS mode for *Data_Deng* dataset.

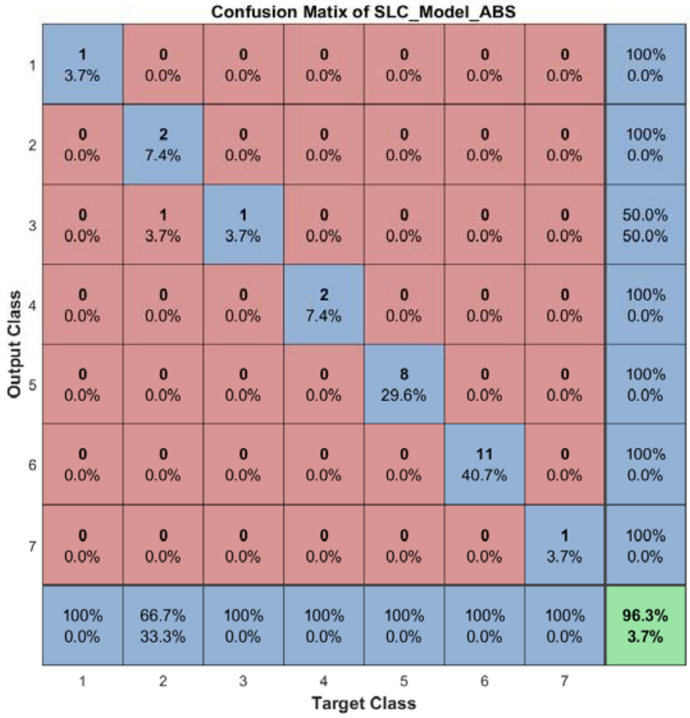


Figure S99: The confusion matrix of ensemble classifier with ABS mode for *Data_Deng* dataset.

#### 2.12 The running parameters and output figures for *Data_Ginhoux* dataset

Table S10: The running parameters for *Data_Ginhoux* dataset.

```
load('MPSSC\Data_Ginhoux.mat'); in_X = 0.3*in_X;
[in_X_SLC,class_num,slice_tik,binary_mod] = slicematrix(in_X,true_labs,linspace(0,1.6,100));
[in_X_SLC_diff,in_X_SLC_SRCIstd,slice_bst,slice_vle] = slicediffer(in_X_SLC,slice_tik,21);
[SLC_Model_DIS1,istrain,istest] = slicemethod(in_X_SLC,true_labs,slice_tik,slice_bst,15,'correlation','inverse',5);
[SLC_Model_DIS2,istrain,istest] = slicemethod(in_X_SLC,true_labs,slice_tik,slice_bst,15,'jaccard','inverse',5);
[SLC_Model_DIS3,istrain,istest] = slicemethod(in_X_SLC,true_labs,slice_tik,slice_bst,15,'cosine','inverse',5);
[SLC_Model_All,SLC_Model_All_SMARwit] = sliceweight(SLC_Model_DIS1,SLC_Model_DIS2,SLC_Model_DIS3);
[SLC_Model_RWS,SLC_Model_All_SMESrws] = sliceswitch(SLC_Model_All,SLC_Model_All_SMARwit,'rws',2);
[SLC_Model_ABS,SLC_Model_All_SMESabs] = sliceswitch(SLC_Model_All,SLC_Model_All_SMARwit,'abs',9);
[SLC_Model_FIT1,FIT1_accuracy] = sliceprerws(in_X(istest,:),true_labs(istest),binary_mod,class_num,slice_vle,SLC_Model_RWS);
[SLC_Model_FIT2,FIT2_accuracy] = slicepreabs(in_X(istest,:),true_labs(istest),binary_mod,class_num,slice_vle,SLC_Model_ABS);
```

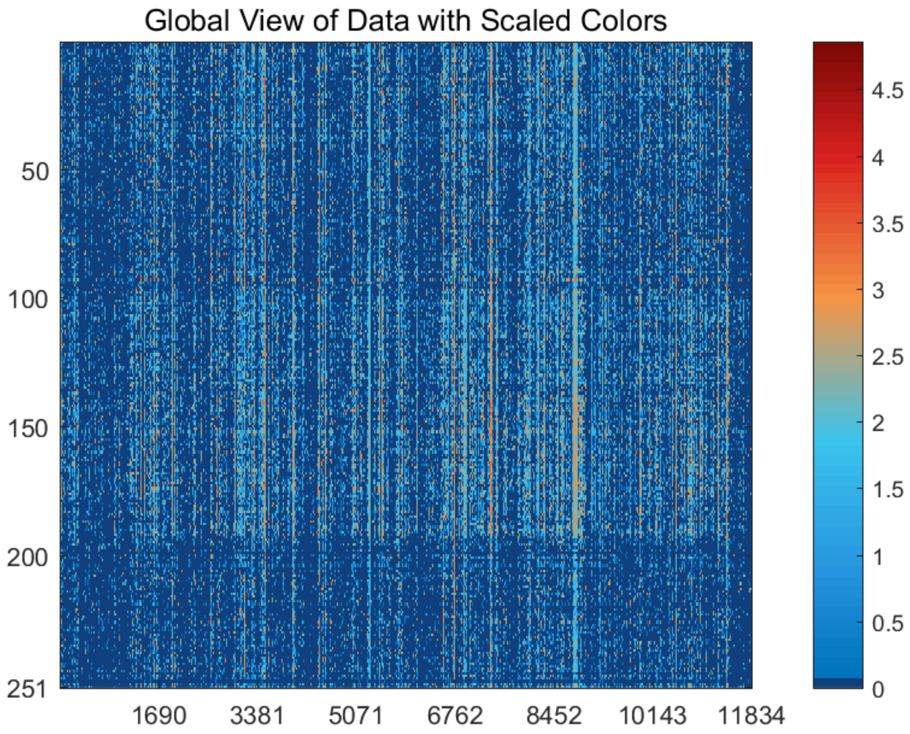


Figure S100: The global view of *Data_Ginhoux* dataset with scaled colors.

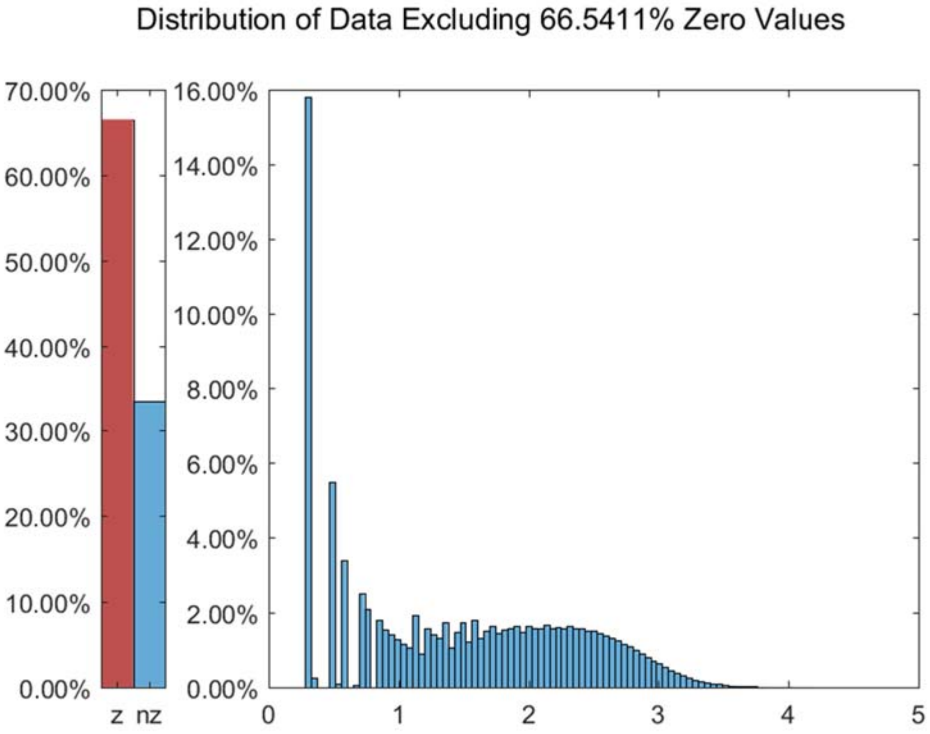


Figure S101: The distribution of *Data_Ginhoux* dataset excluding all zero values.

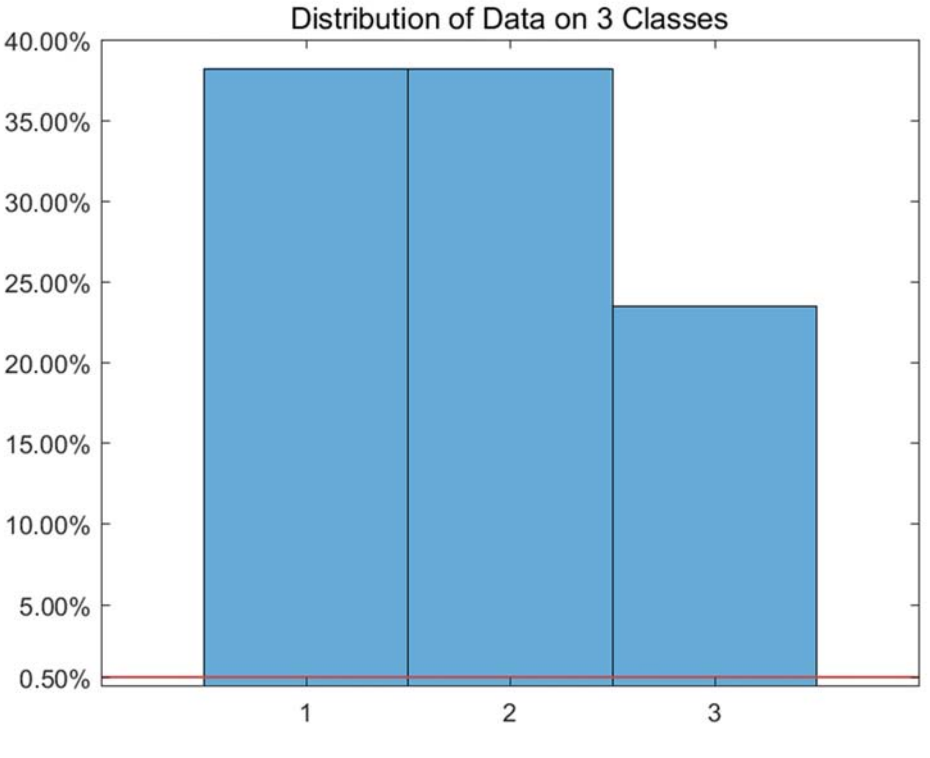


Figure S102: The distribution of *Data_Ginhoux* dataset on every classes.

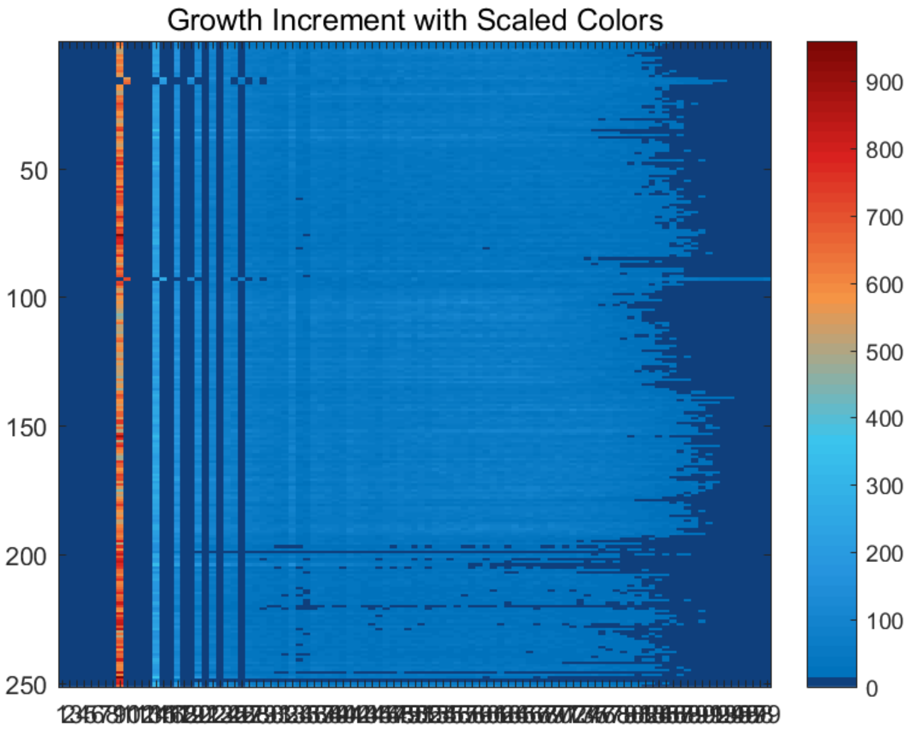


Figure S103: The growth increment with scaled colors for *Data_Ginhoux* dataset.

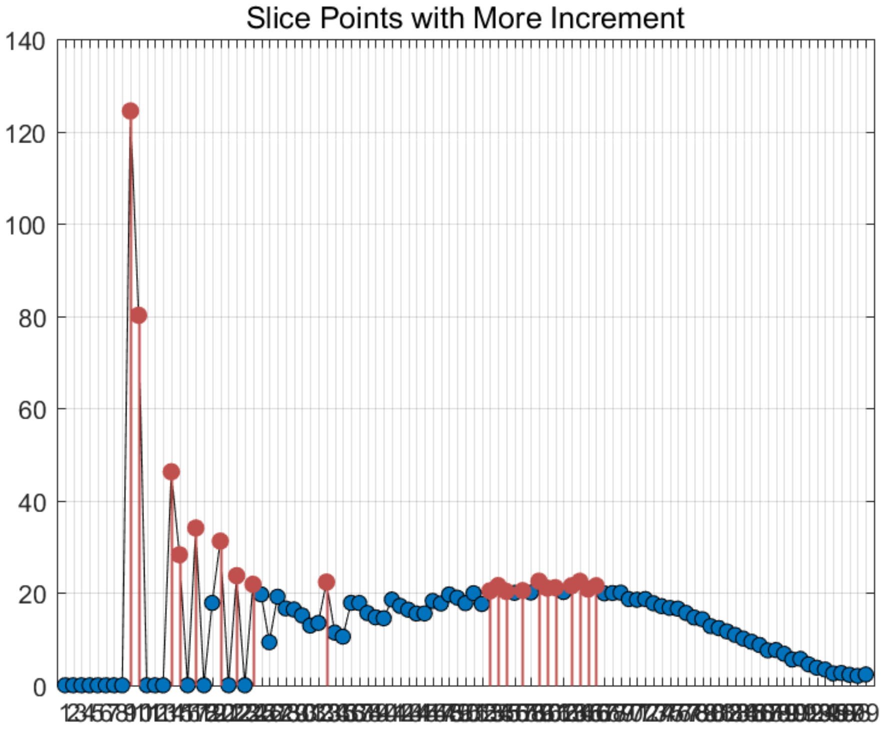


Figure S104: The slice points with more increment for *Data_Ginhoux* dataset.

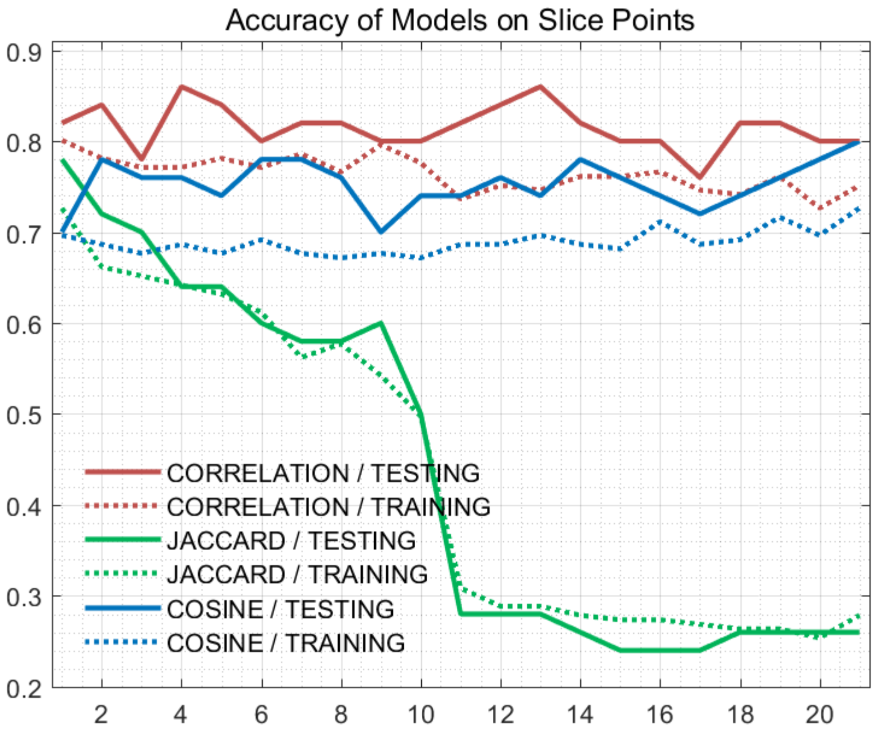


Figure S105: The accuracy of models on every slice points for *Data_Ginhoux* dataset.

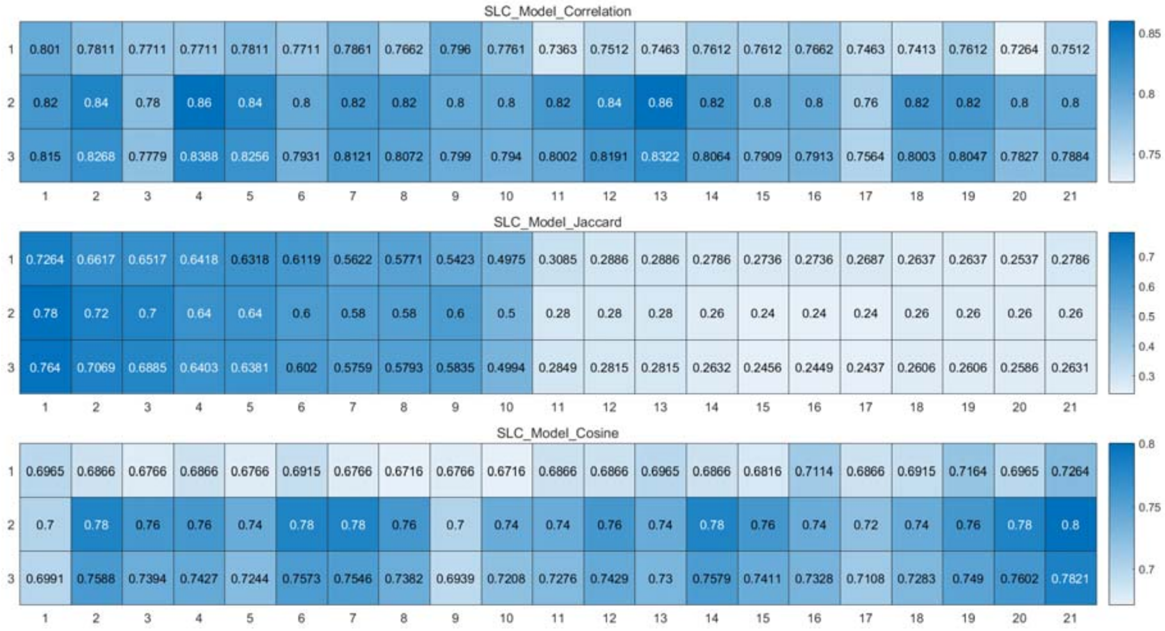


Figure S106: The weighted accuracy on every slice points for *Data_Ginhoux* dataset.

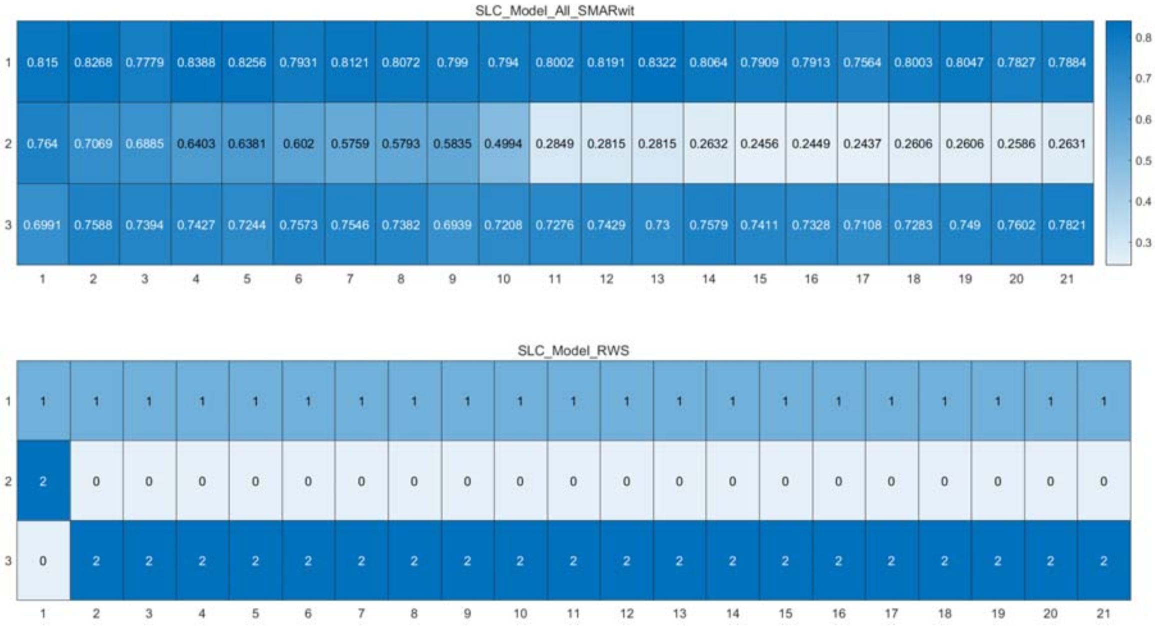


Figure S107: The RWS mode of meta classifiers for *Data_Ginhoux* dataset.

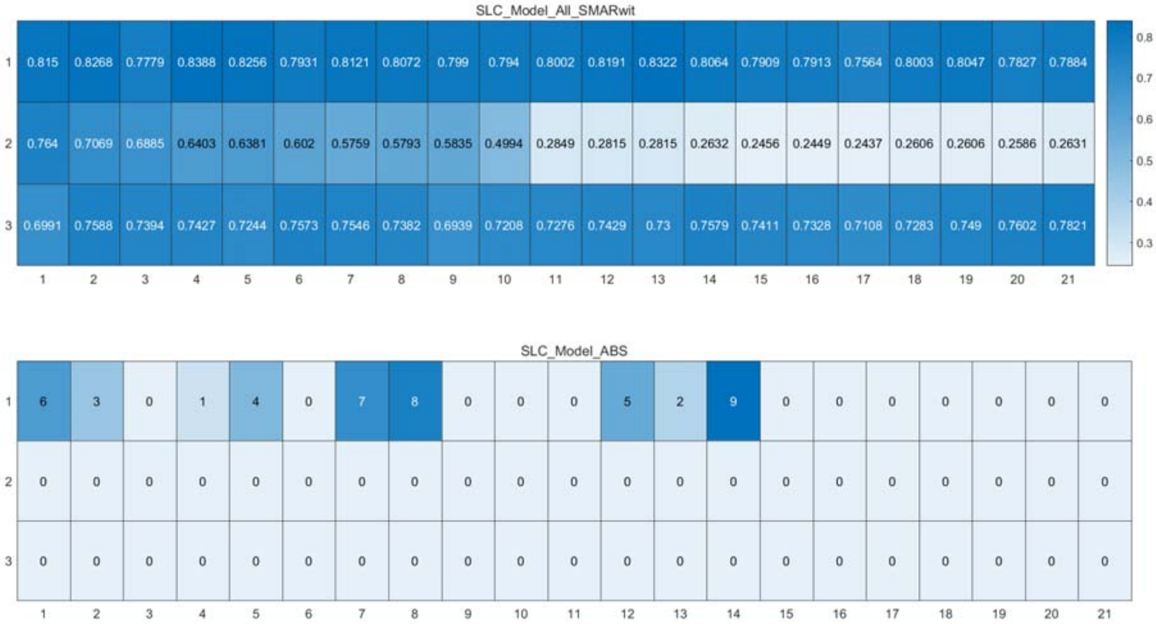


Figure S108: The ABS mode of meta classifiers for *Data_Ginhoux* dataset.

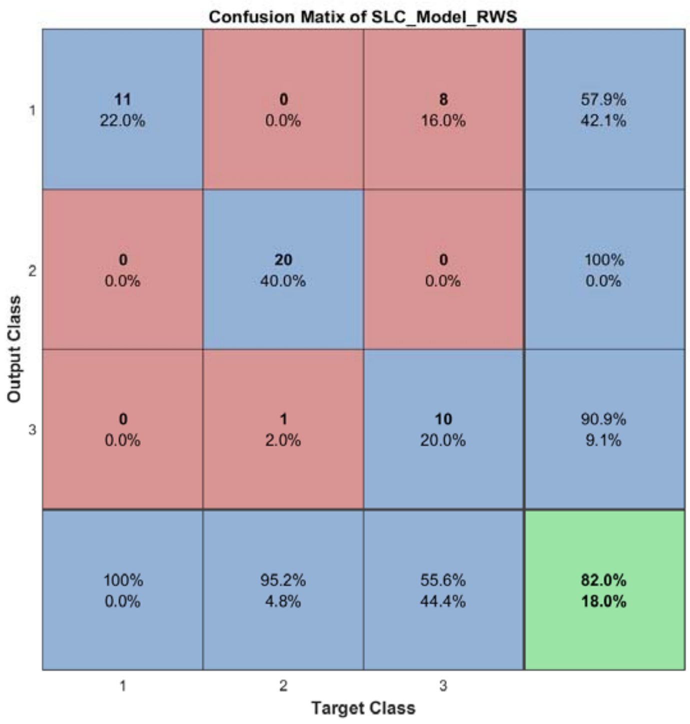


Figure S109: The confusion matrix of ensemble classifier with RWS mode for *Data_Ginhoux* dataset.

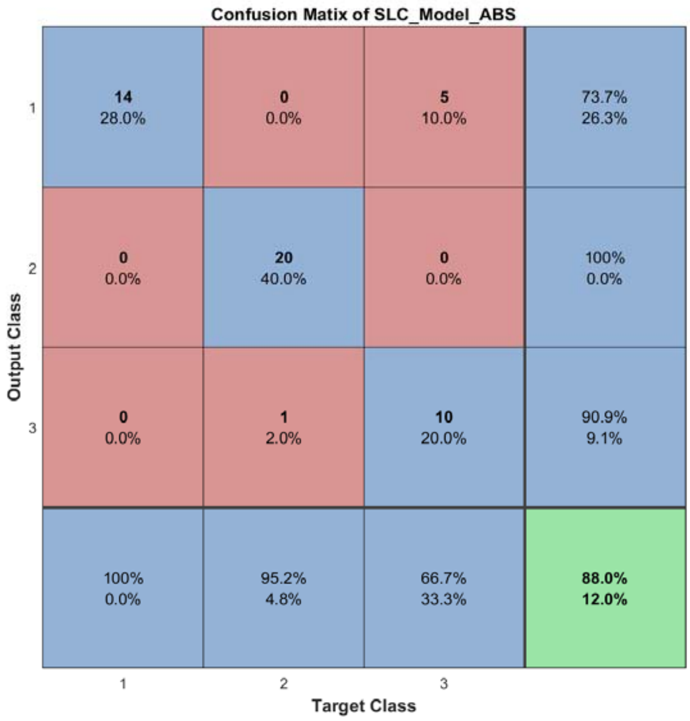


Figure S110: The confusion matrix of ensemble classifier with ABS mode for *Data_Ginhoux* dataset.

#### 2.13 The running parameters and output figures for *Data_Macosko* dataset (with pca)

Table S11: The running parameters for *Data_Macosko* dataset.

```
load('MPSSC\Data_Macosko.mat'); [coeff,score,latent,~,explained] = pca(in_X); in_X_pca = in_X*coeff(:,1:100);
in_X = in_X_pca; in_X = in_X-min(min(in_X)); in_X = log(in_X+1);
[in_X_SLC,class_num,slice_tik,binary_mod] = slicematrix(in_X,true_labs,linspace(3.6,4.6,60));
[in_X_SLC_diff,in_X_SLC_SRCIstd,slice_bst,slice_vle] = slicediffer(in_X_SLC,slice_tik,9);
[SLC_Model_DIS1,istrain,istest] = slicemethod(in_X_SLC,true_labs,slice_tik,slice_bst,1,'correlation','inverse',3);
[SLC_Model_DIS2,istrain,istest] = slicemethod(in_X_SLC,true_labs,slice_tik,slice_bst,1,'jaccard','inverse',3);
[SLC_Model_DIS3,istrain,istest] = slicemethod(in_X_SLC,true_labs,slice_tik,slice_bst,1,'cosine','inverse',3);
[SLC_Model_All,SLC_Model_All_SMARwit] = sliceweight(SLC_Model_DIS1,SLC_Model_DIS2,SLC_Model_DIS3);
[SLC_Model_RWS,SLC_Model_All_SMESrws] = sliceswitch(SLC_Model_All,SLC_Model_All_SMARwit,'rws',2);
[SLC_Model_ABS,SLC_Model_All_SMESabs] = sliceswitch(SLC_Model_All,SLC_Model_All_SMARwit,'abs',9);
[SLC_Model_FIT1,FIT1_accuracy] = sliceprerws(in_X(istest,:),true_labs(istest),binary_mod,class_num,slice_vle,SLC_Model_RWS);
[SLC_Model_FIT2,FIT2_accuracy] = slicepreabs(in_X(istest,:),true_labs(istest),binary_mod,class_num,slice_vle,SLC_Model_ABS);
```

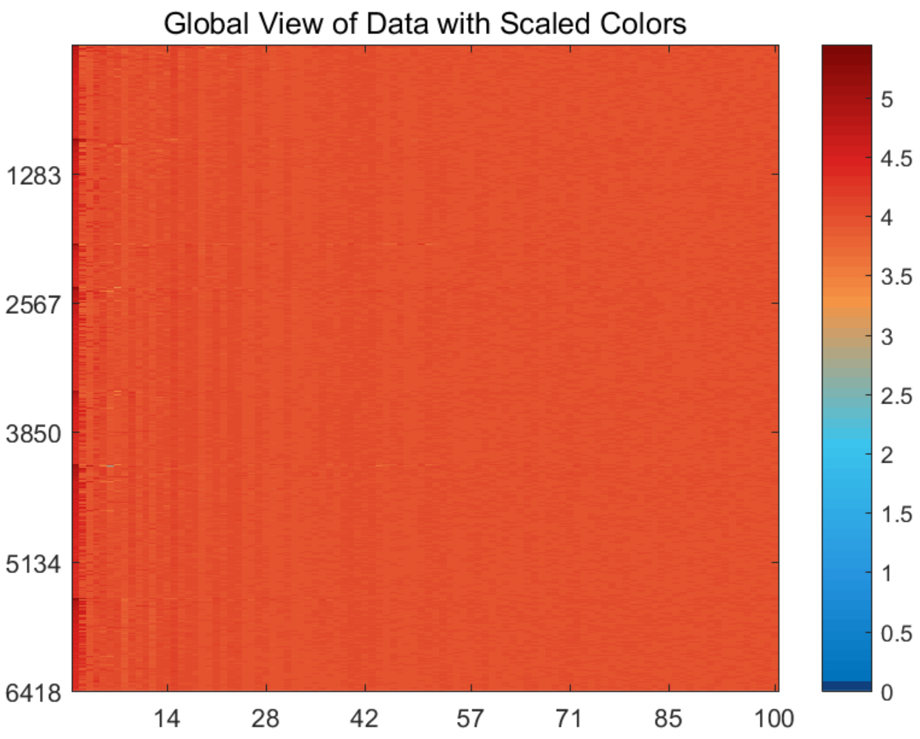


Figure S111: The global view of *Data_Macosko* dataset with scaled colors.

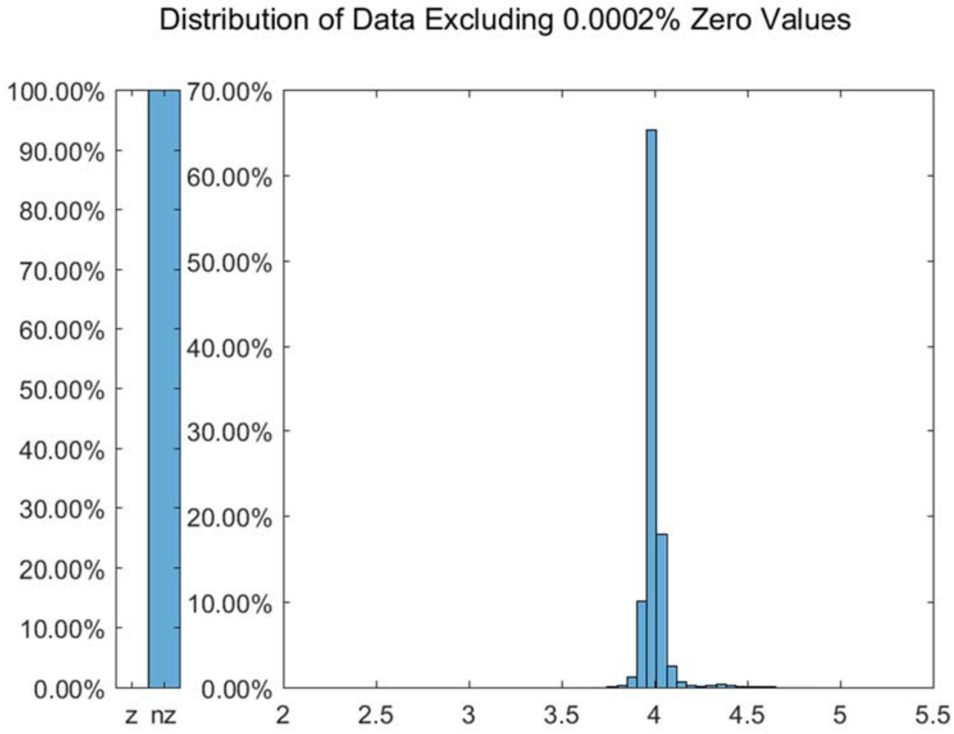


Figure S112: The distribution of *Data_Macosko* dataset excluding all zero values.

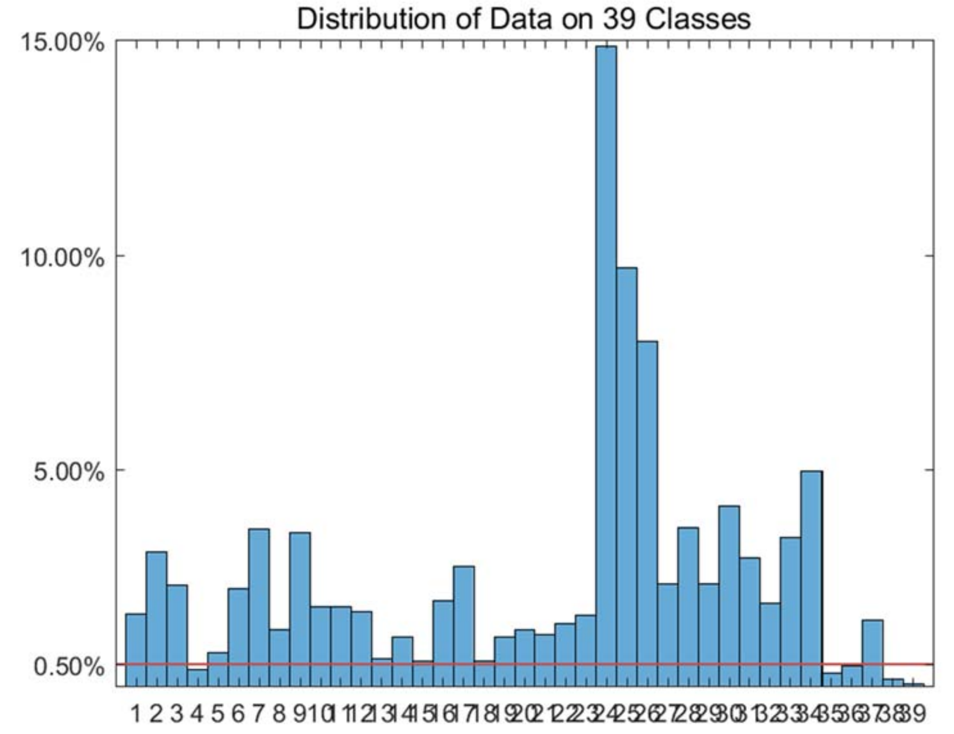


Figure S113: The distribution of *Data_Macosko* dataset on every classes.

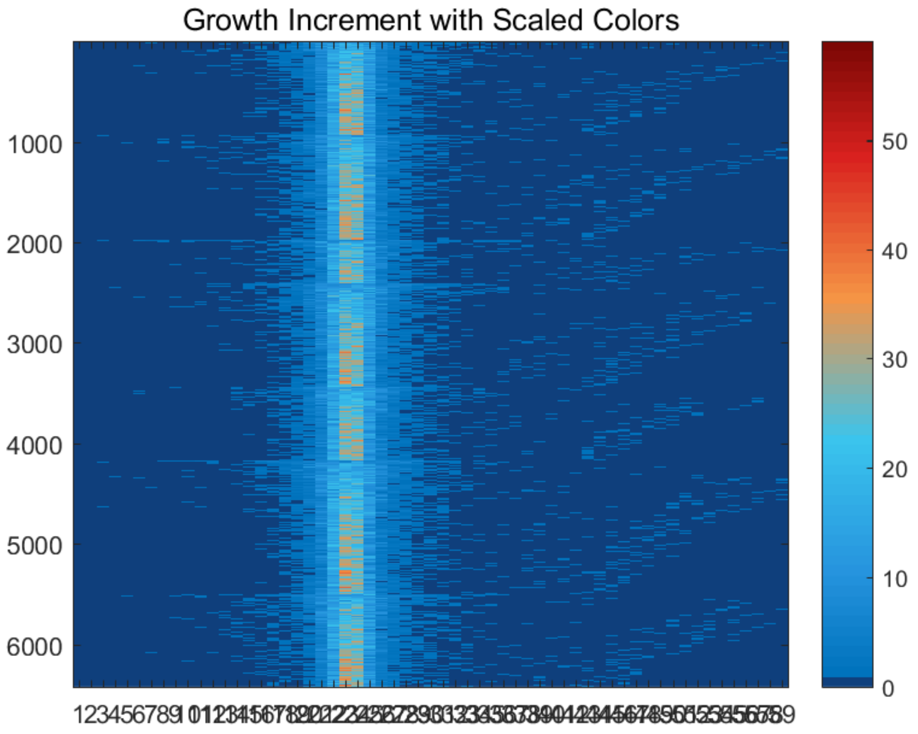


Figure S114: The growth increment with scaled colors for *Data_Macosko* dataset.

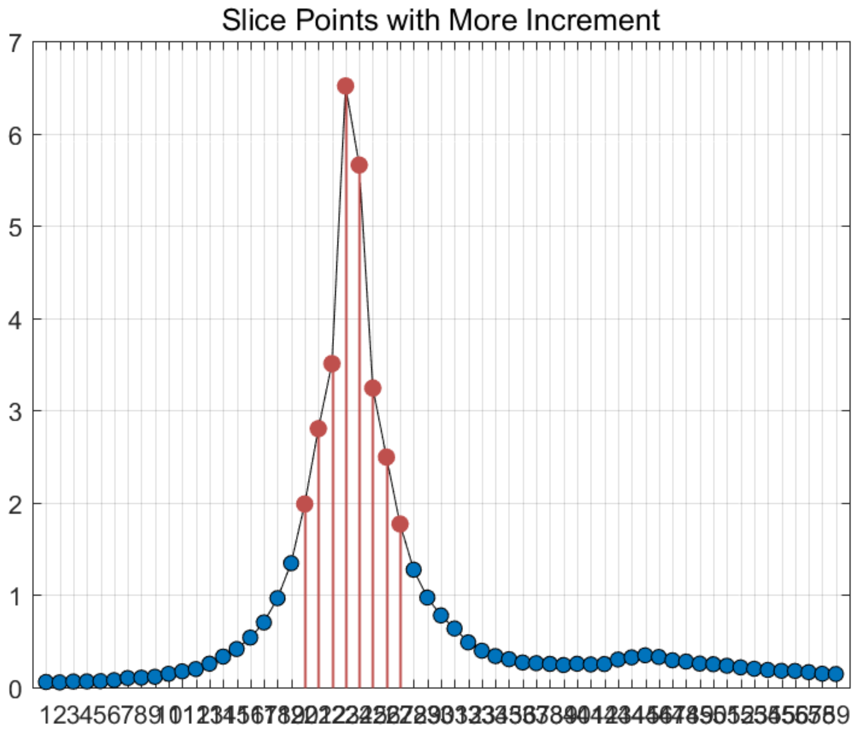


Figure S115: The slice points with more increment for *Data_Macosko* dataset.

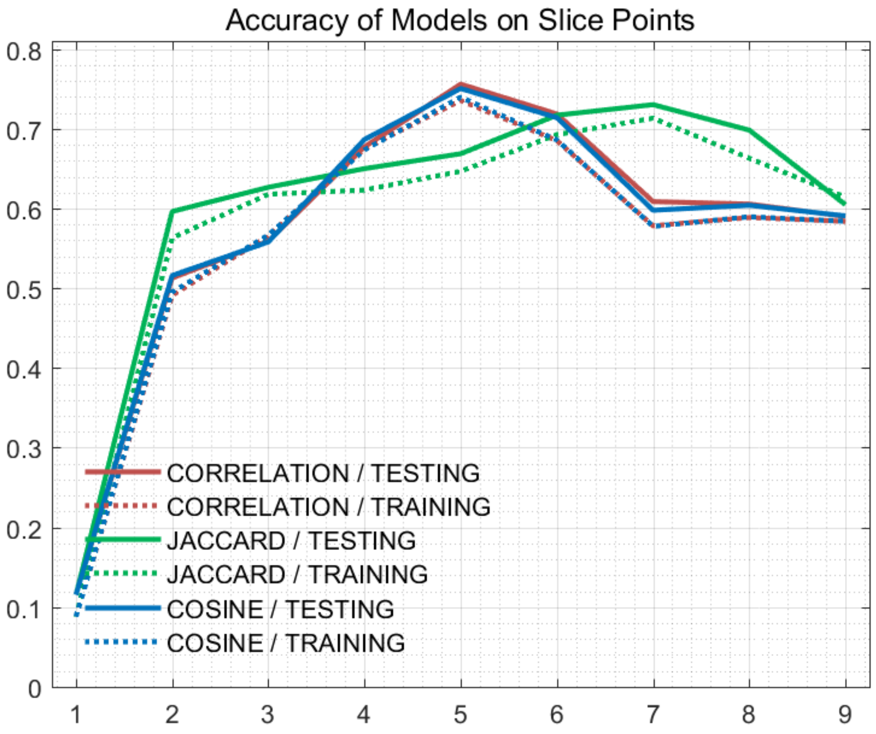


Figure S116: The accuracy of models on every slice points for *Data_Macosko* dataset.

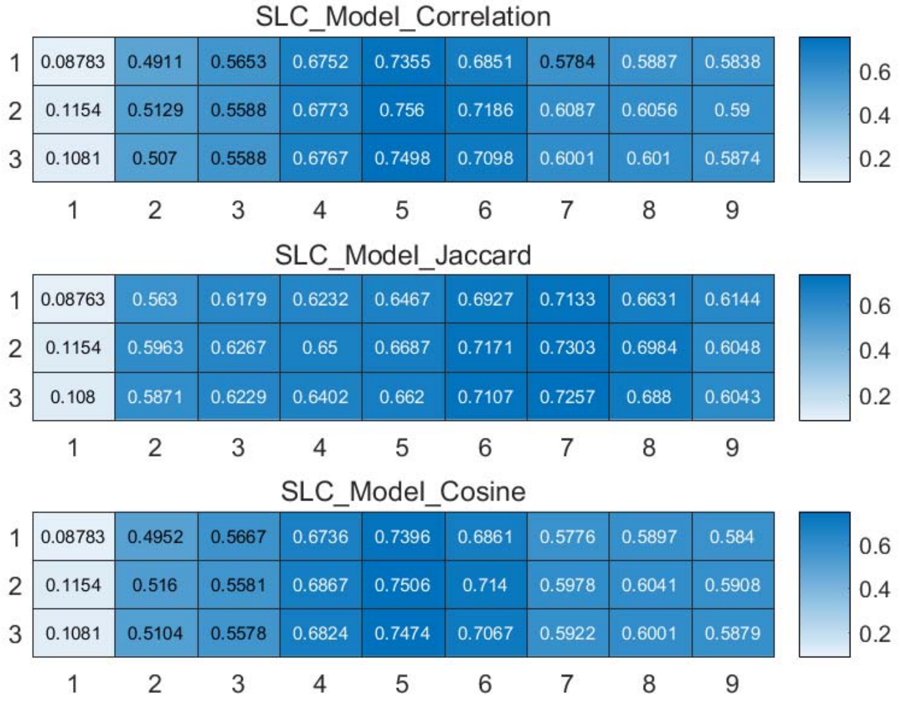


Figure S117: The weighted accuracy on every slice points for *Data_Macosko* dataset.

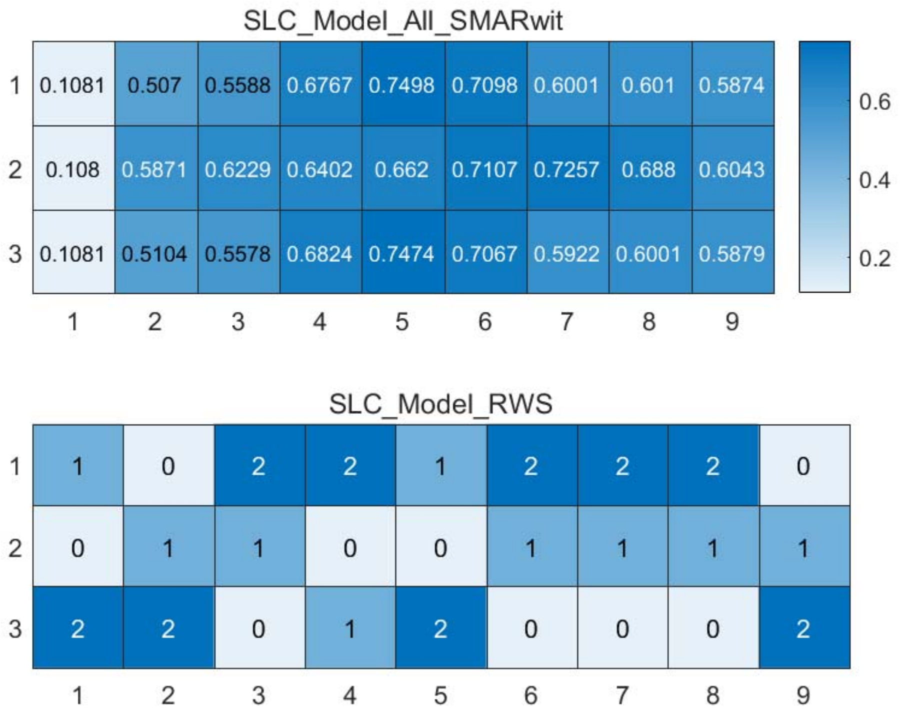


Figure S118: The RWS mode of meta classifiers for *Data_Macosko* dataset.

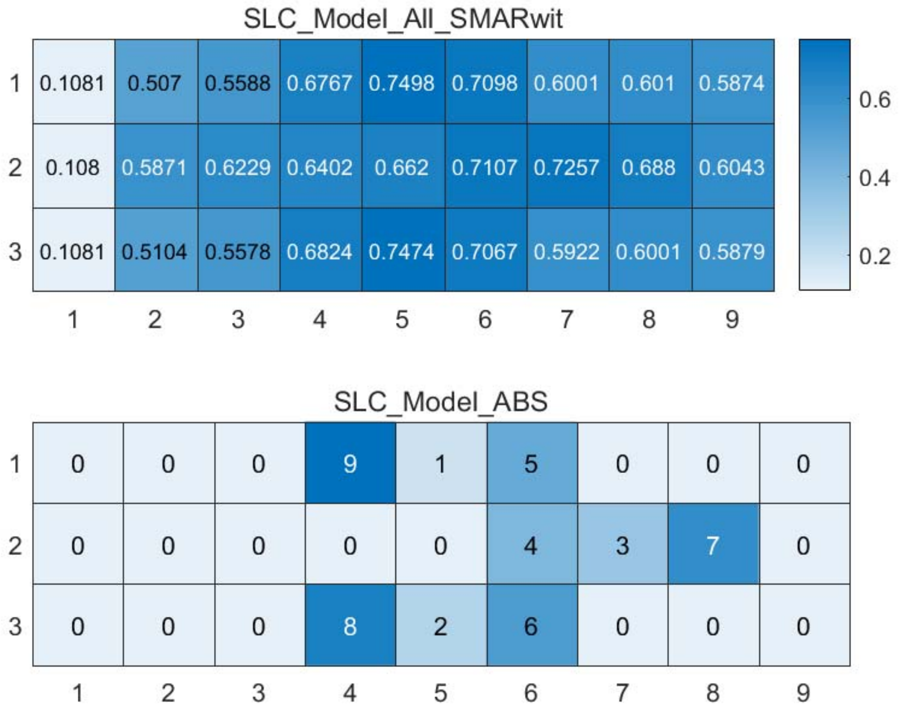


Figure S119: The ABS mode of meta classifiers for *Data_Macosko* dataset.

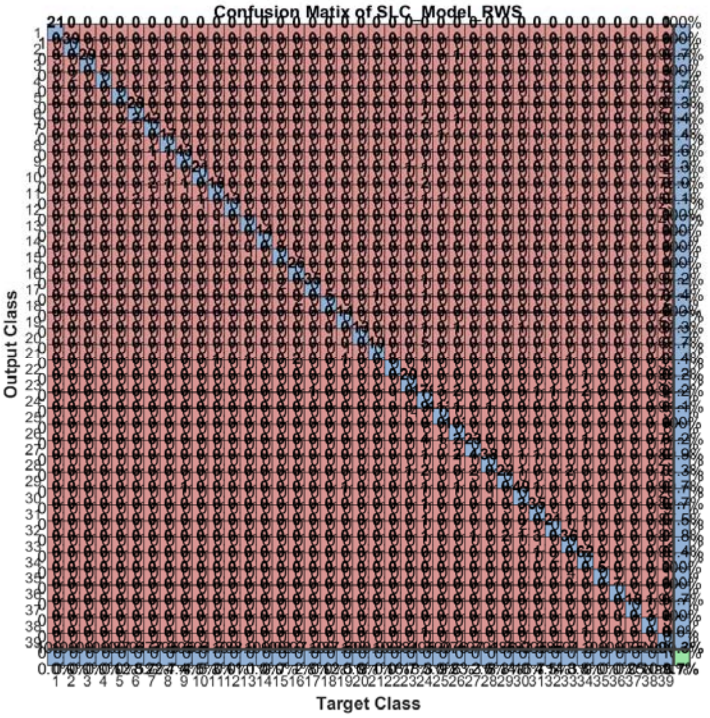


Figure S120: The confusion matrix of ensemble classifier with RWS mode for *Data_Macosko* dataset. (*Note*: the confusion matrix is too large to display completely in one figure)

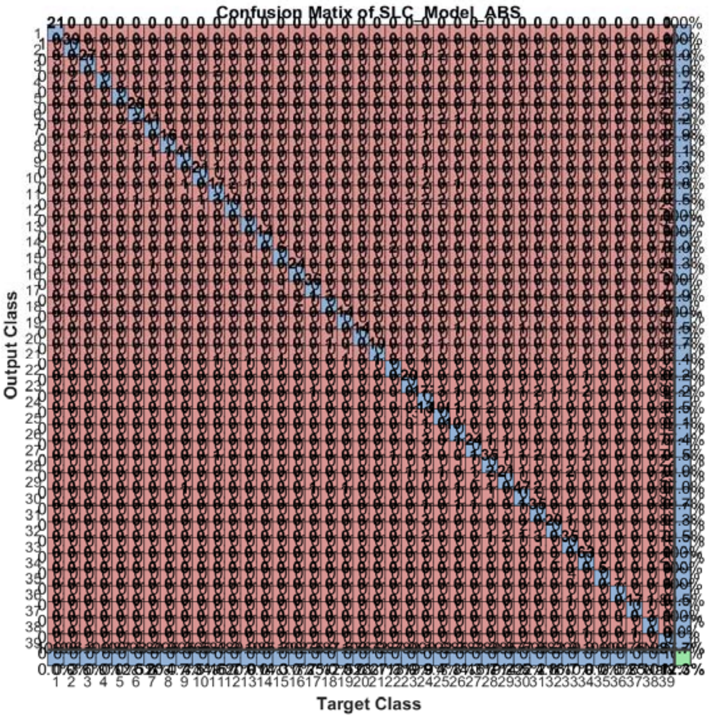


Figure S121: The confusion matrix of ensemble classifier with ABS mode for *Data_Macosko* dataset. (*Note*: the confusion matrix is too large to display completely in one figure)

#### 2.14 The running parameters and output figures for *Data_Pollen* dataset

Table S12: The running parameters for *Data_Pollen* dataset.

```
load('MPSSC\Data_Pollen.mat'); in_X = 0.3*in_X;
[in_X_SLC,class_num,slice_tik,binary_mod] = slicematrix(in_X,true_labs,linspace(0,1.3,60));
[in_X_SLC_diff,in_X_SLC_SRCIstd,slice_bst,slice_vle] = slicediffer(in_X_SLC,slice_tik,9);
[SLC_Model_DIS1,istrain,istest] = slicemethod(in_X_SLC,true_labs,slice_tik,slice_bst,1,'correlation','inverse',5);
[SLC_Model_DIS2,istrain,istest] = slicemethod(in_X_SLC,true_labs,slice_tik,slice_bst,1,'jaccard','inverse',5);
[SLC_Model_DIS3,istrain,istest] = slicemethod(in_X_SLC,true_labs,slice_tik,slice_bst,1,'cosine','inverse',5);
[SLC_Model_All,SLC_Model_All_SMARwit] = sliceweight(SLC_Model_DIS1,SLC_Model_DIS2,SLC_Model_DIS3);
[SLC_Model_RWS,SLC_Model_All_SMESrws] = sliceswitch(SLC_Model_All,SLC_Model_All_SMARwit,'rws',2);
[SLC_Model_ABS,SLC_Model_All_SMESabs] = sliceswitch(SLC_Model_All,SLC_Model_All_SMARwit,'abs',9);
[SLC_Model_FIT1,FIT1_accuracy] = sliceprerws(in_X(istest,:),true_labs(istest),binary_mod,class_num,slice_vle,SLC_Model_RWS);
[SLC_Model_FIT2,FIT2_accuracy] = slicepreabs(in_X(istest,:),true_labs(istest),binary_mod,class_num,slice_vle,SLC_Model_ABS);
```

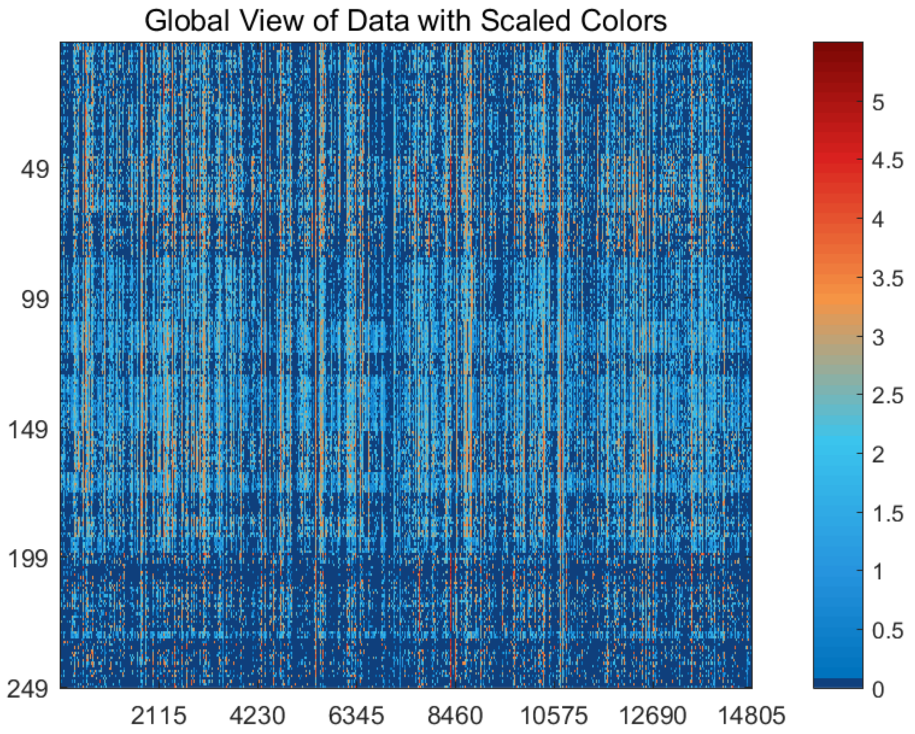


Figure S122: The global view of *Data_Pollen* dataset with scaled colors.

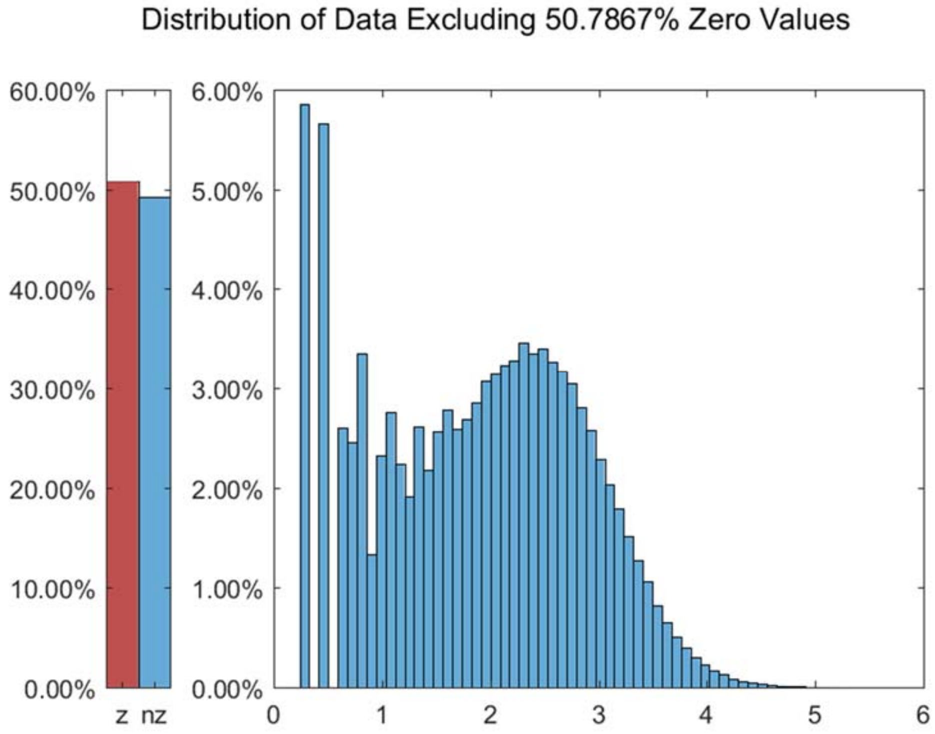


Figure S123: The distribution of *Data_Pollen* dataset excluding all zero values.

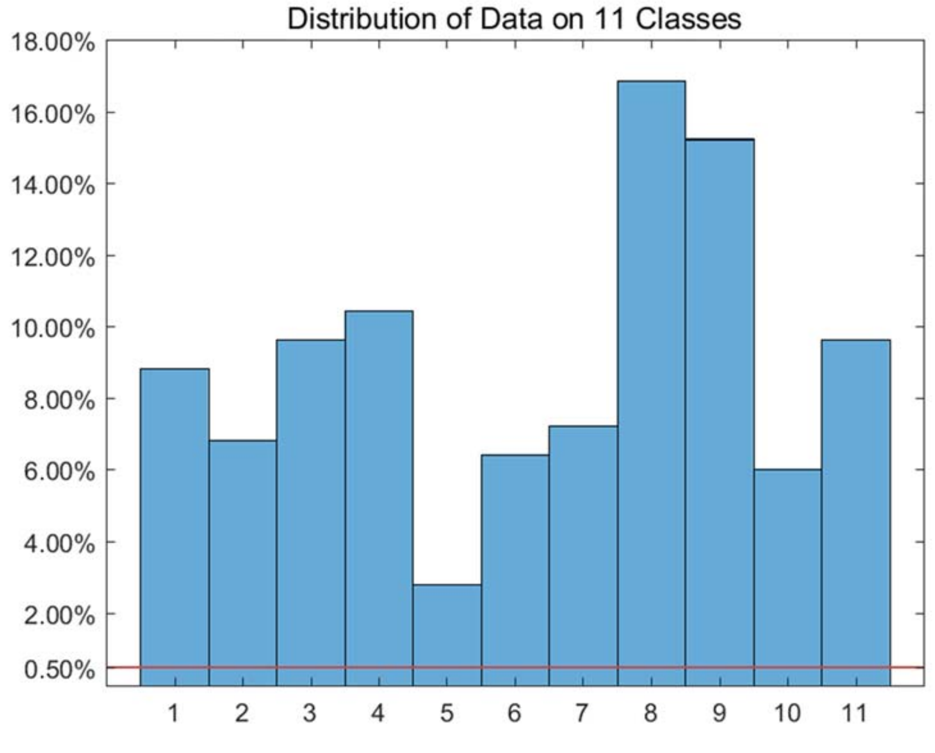


Figure S124: The distribution of *Data_Pollen* dataset on every classes.

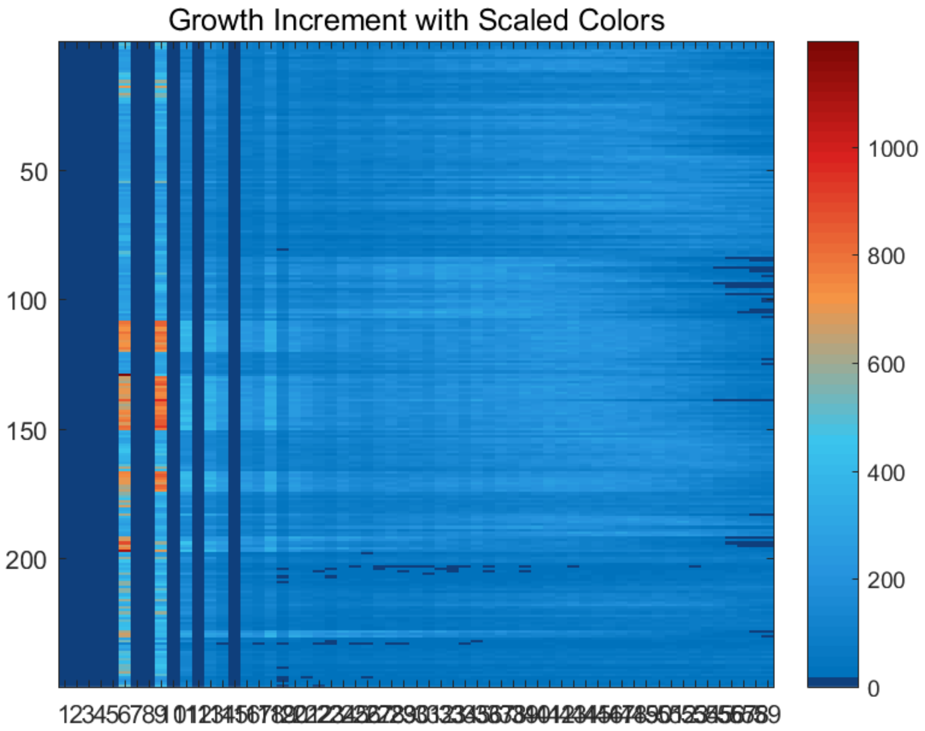


Figure S125: The growth increment with scaled colors for *Data_Pollen* dataset.

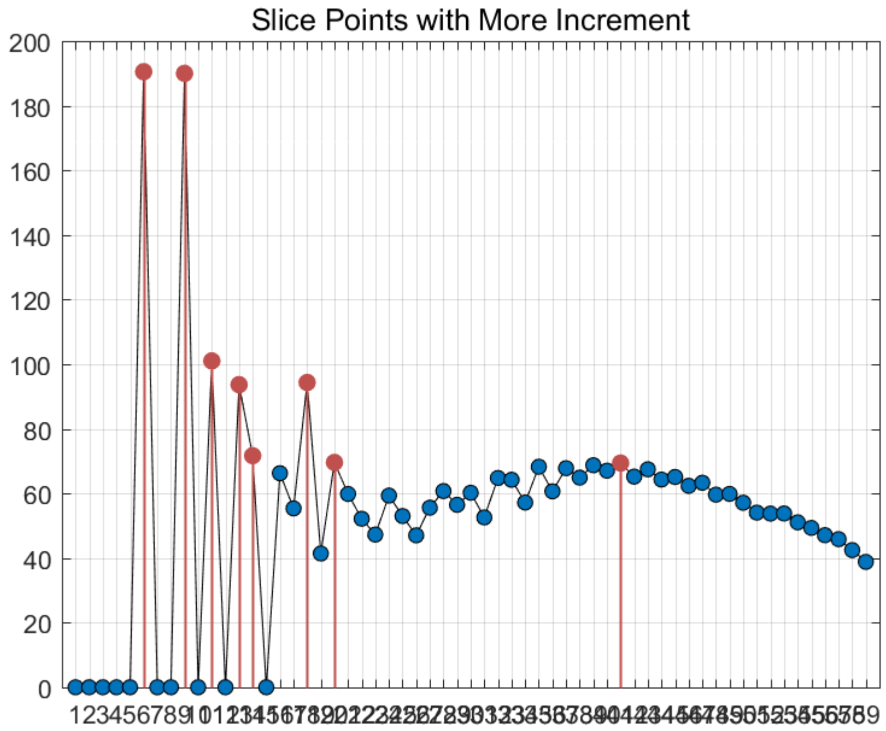


Figure S126: The slice points with more increment for *Data_Pollen* dataset.

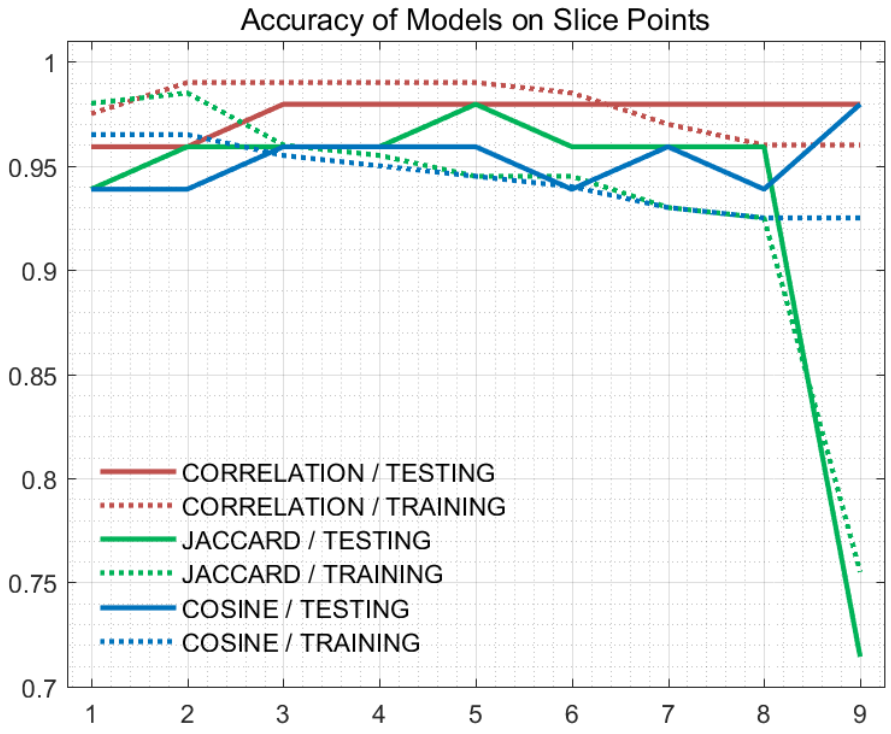


Figure S127: The accuracy of models on every slice points for *Data_Pollen* dataset.

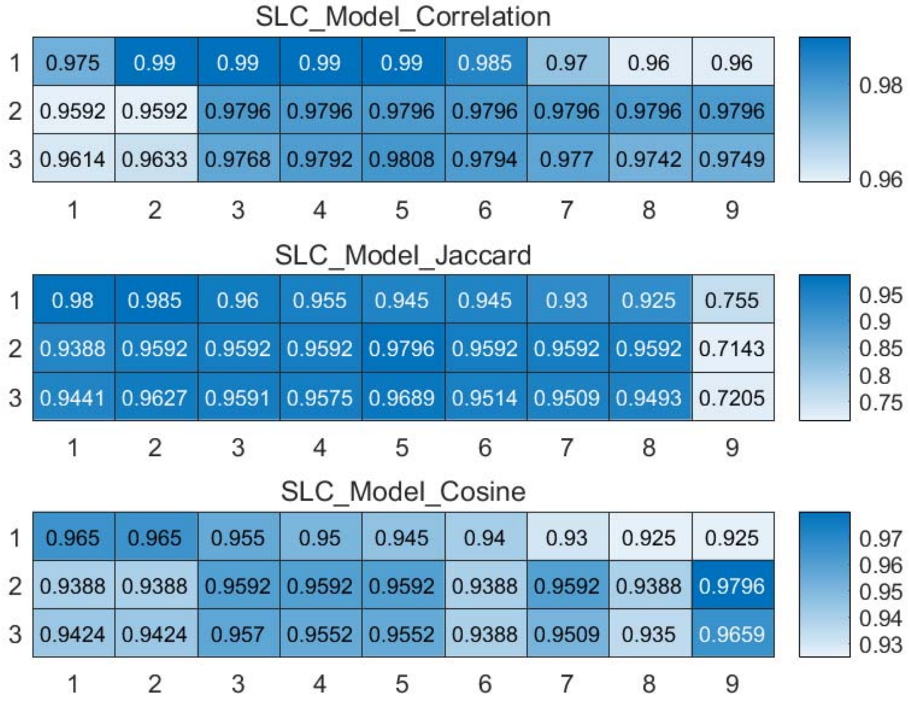


Figure S128: The weighted accuracy on every slice points for *Data_Pollen* dataset.

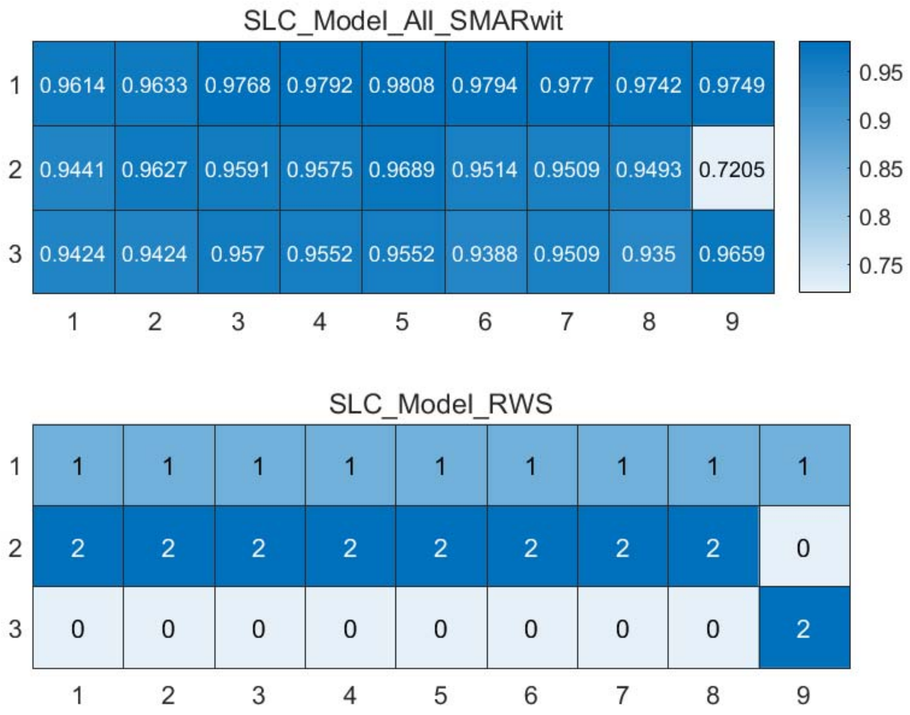


Figure S129: The RWS mode of meta classifiers for *Data_Pollen* dataset.

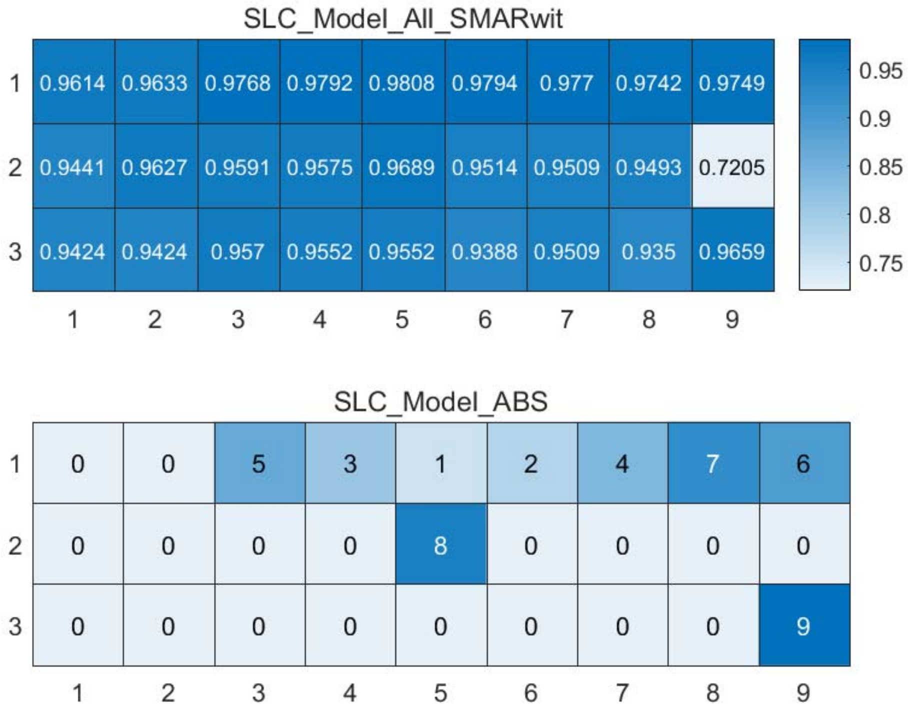


Figure S130: The ABS mode of meta classifiers for *Data_Pollen* dataset.

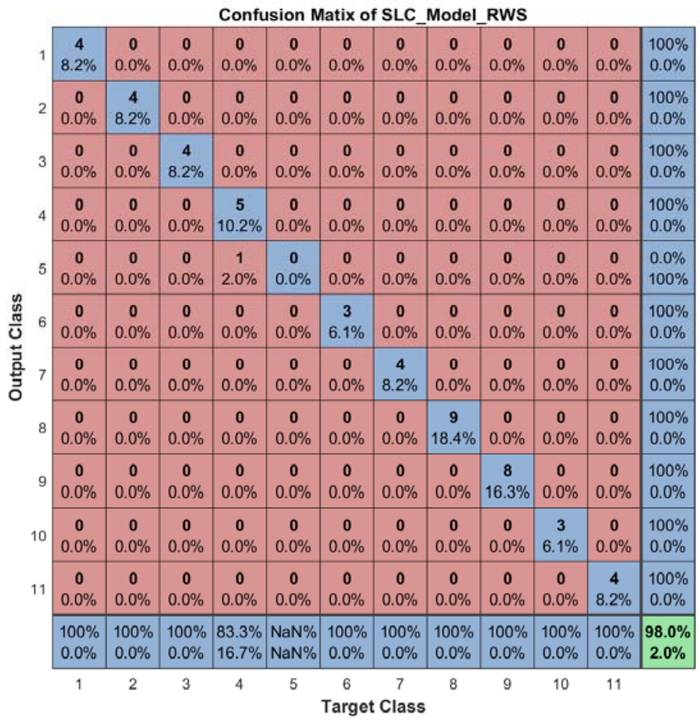


Figure S131: The confusion matrix of ensemble classifier with RWS mode for *Data_Pollen* dataset.

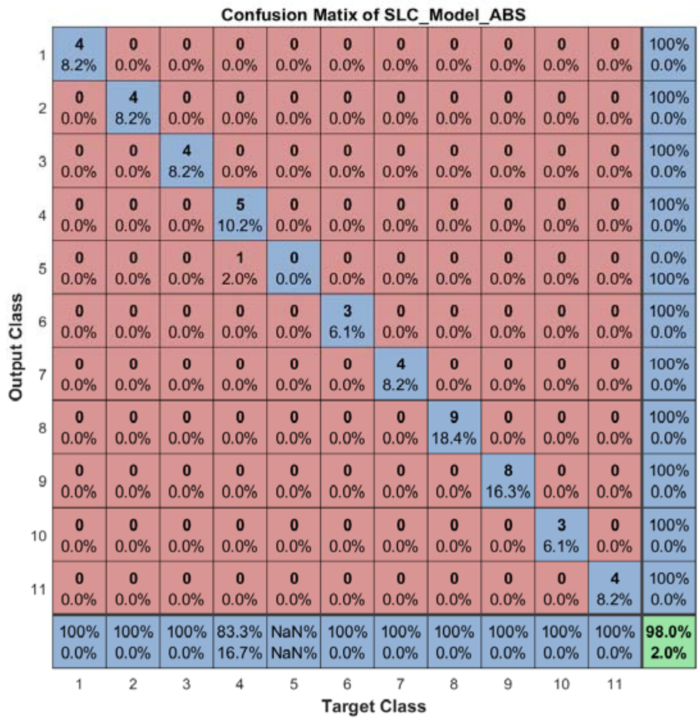


Figure S132: The confusion matrix of ensemble classifier with ABS mode for *Data_Pollen* dataset.

#### 2.15 The running parameters and output figures for *Data_Tasic* dataset

Table S13: The running parameters for *Data_Tasic* dataset.

```
load('MPSSC\Data_Tasic.mat'); true_labs = true_labs-double(true_labs>25);
[in_X_SLC,class_num,slice_tik,binary_mod] = slicematrix(in_X,true_labs,0:0.1:3);
[in_X_SLC_diff,in_X_SLC_SRCIstd,slice_bst,slice_vle] = slicediffer(in_X_SLC,slice_tik,6);
[SLC_Model_DIS1,istrain,istest] = slicemethod(in_X_SLC,true_labs,slice_tik,slice_bst,1,'correlation','inverse',3);
[SLC_Model_DIS2,istrain,istest] = slicemethod(in_X_SLC,true_labs,slice_tik,slice_bst,1,'jaccard','inverse',3);
[SLC_Model_DIS3,istrain,istest] = slicemethod(in_X_SLC,true_labs,slice_tik,slice_bst,1,'cosine','inverse',3);
[SLC_Model_All,SLC_Model_All_SMARwit] = sliceweight(SLC_Model_DIS1,SLC_Model_DIS2,SLC_Model_DIS3);
[SLC_Model_RWS,SLC_Model_All_SMESrws] = sliceswitch(SLC_Model_All,SLC_Model_All_SMARwit,'rws',2);
[SLC_Model_ABS,SLC_Model_All_SMESabs] = sliceswitch(SLC_Model_All,SLC_Model_All_SMARwit,'abs',9);
[SLC_Model_FIT1,FIT1_accuracy] = sliceprerws(in_X(istest,:),true_labs(istest),binary_mod,class_num,slice_vle,SLC_Model_RWS);
[SLC_Model_FIT2,FIT2_accuracy] = slicepreabs(in_X(istest,:),true_labs(istest),binary_mod,class_num,slice_vle,SLC_Model_ABS);
```

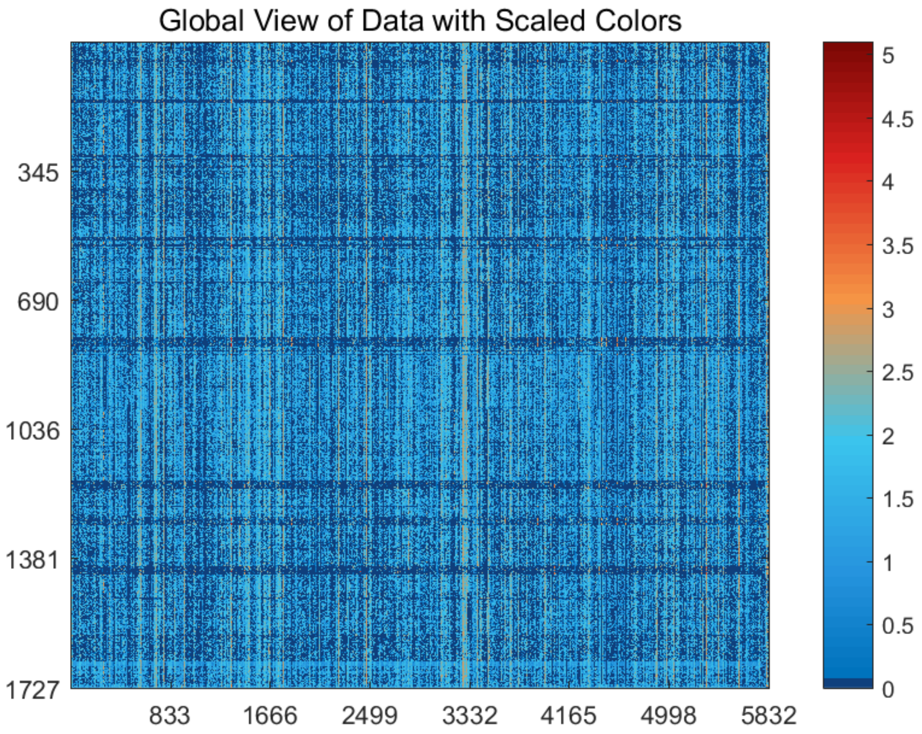


Figure S133: The global view of *Data_Tasic* dataset with scaled colors.

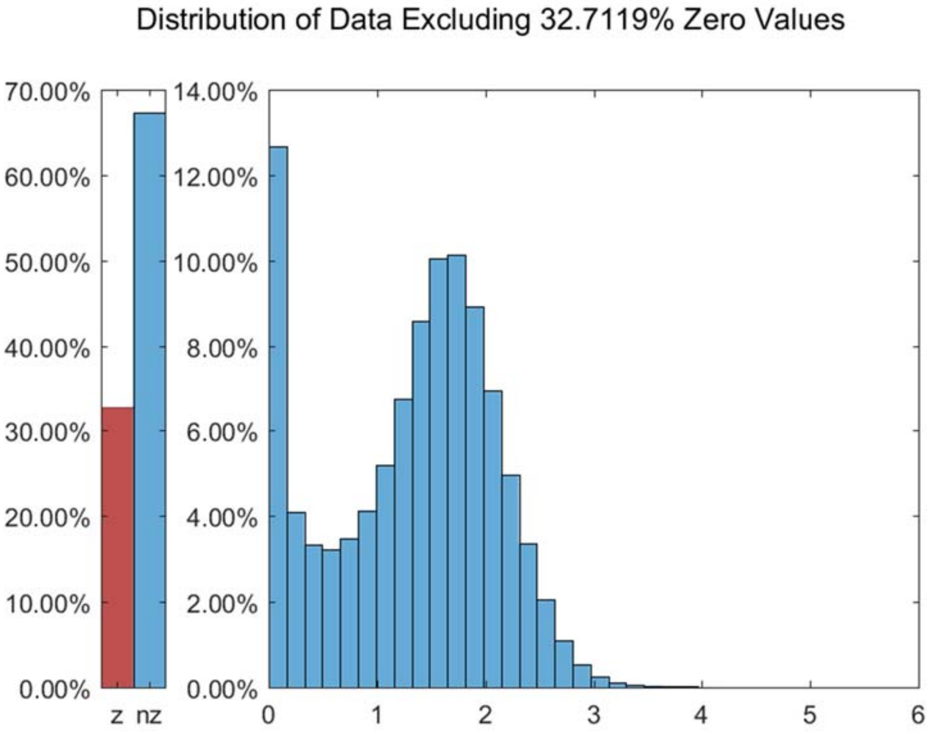


Figure S134: The distribution of *Data_Tasic* dataset excluding all zero values.

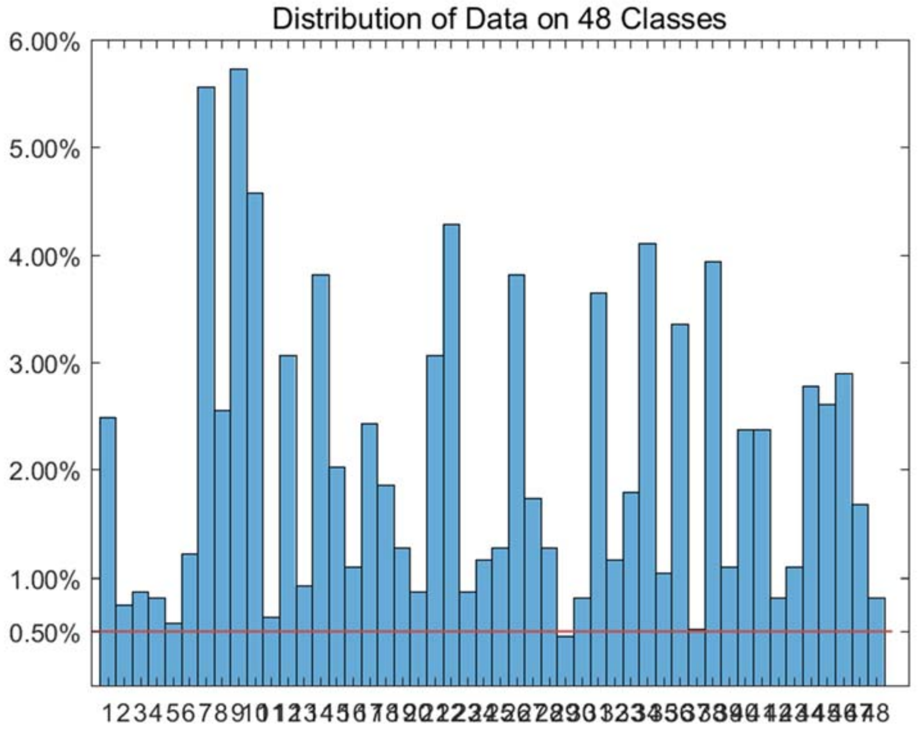


Figure S135: The distribution of *Data_Tasic* dataset on every classes.

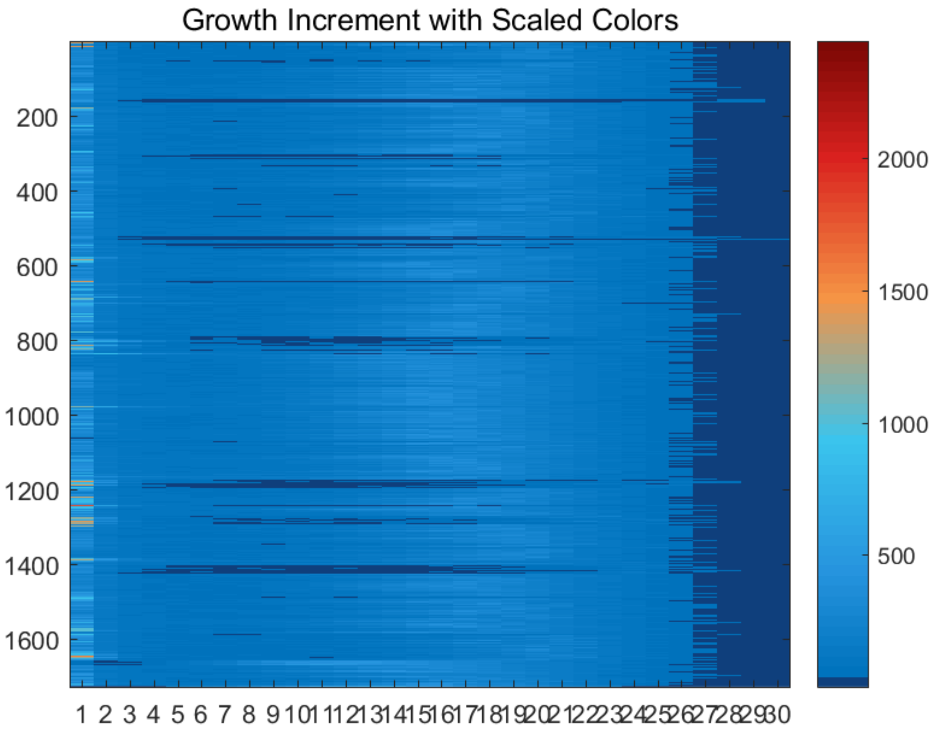


Figure S136: The growth increment with scaled colors for *Data_Tasic* dataset.

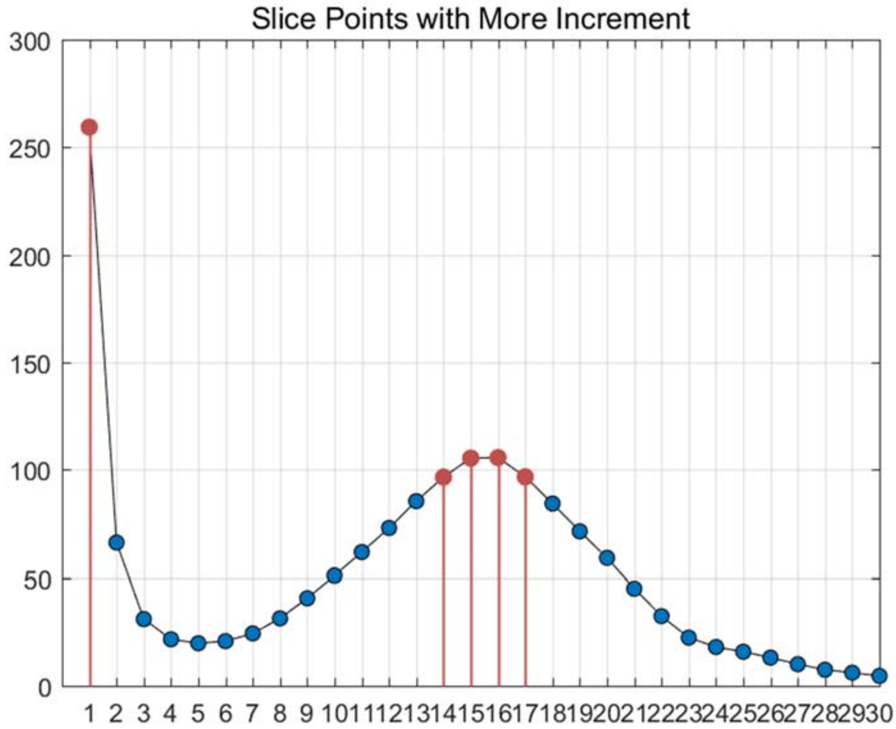


Figure S137: The slice points with more increment for *Data_Tasic* dataset.

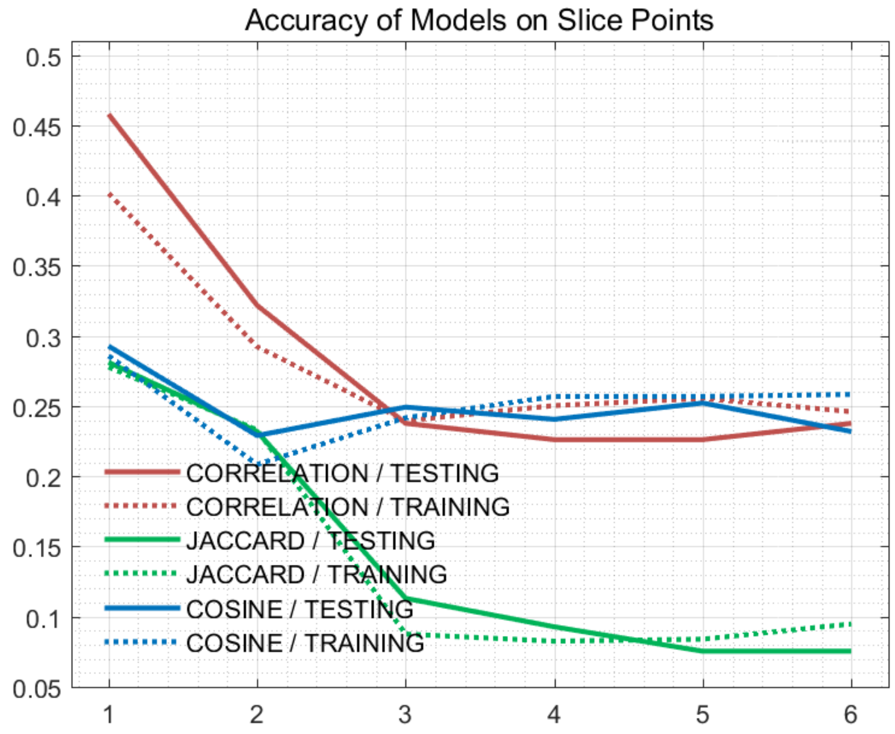


Figure S138: The accuracy of models on every slice points for *Data_Tasic* dataset.

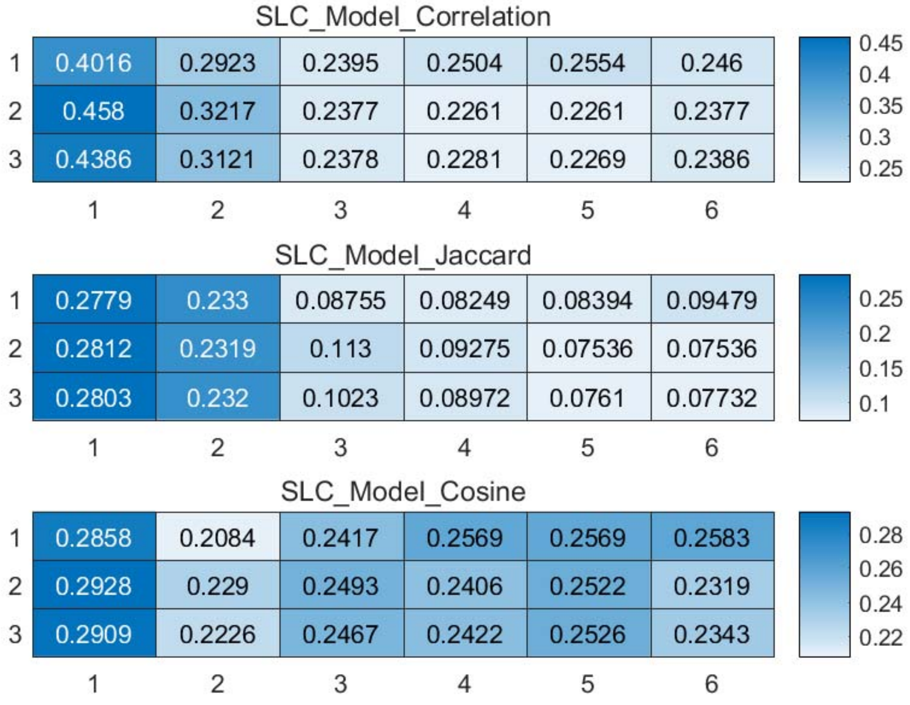


Figure S139: The weighted accuracy on every slice points for *Data_Tasic* dataset.

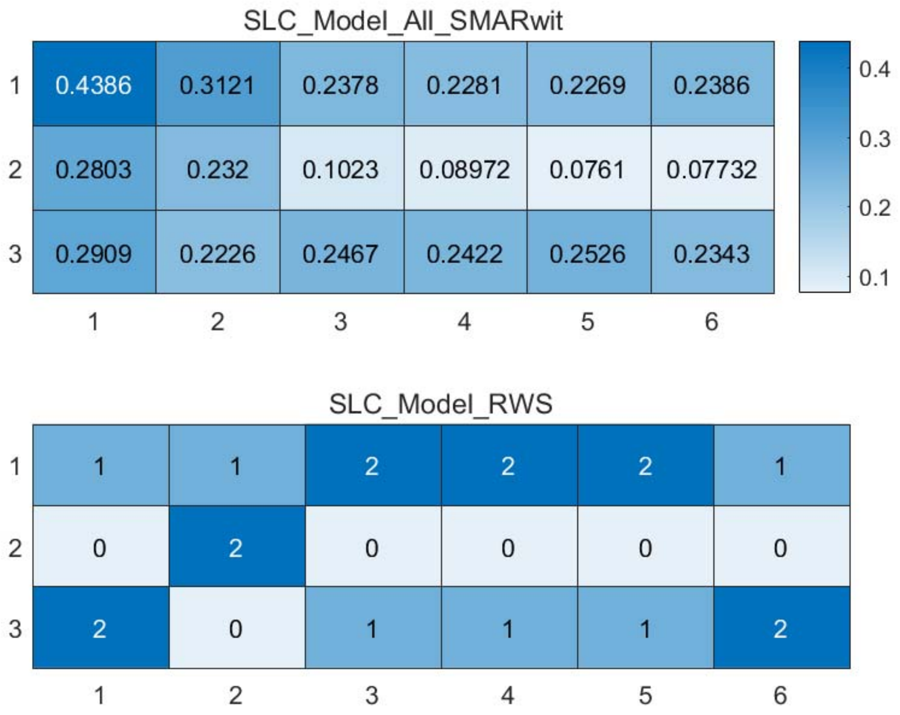


Figure S140: The RWS mode of meta classifiers for *Data_Tasic* dataset.

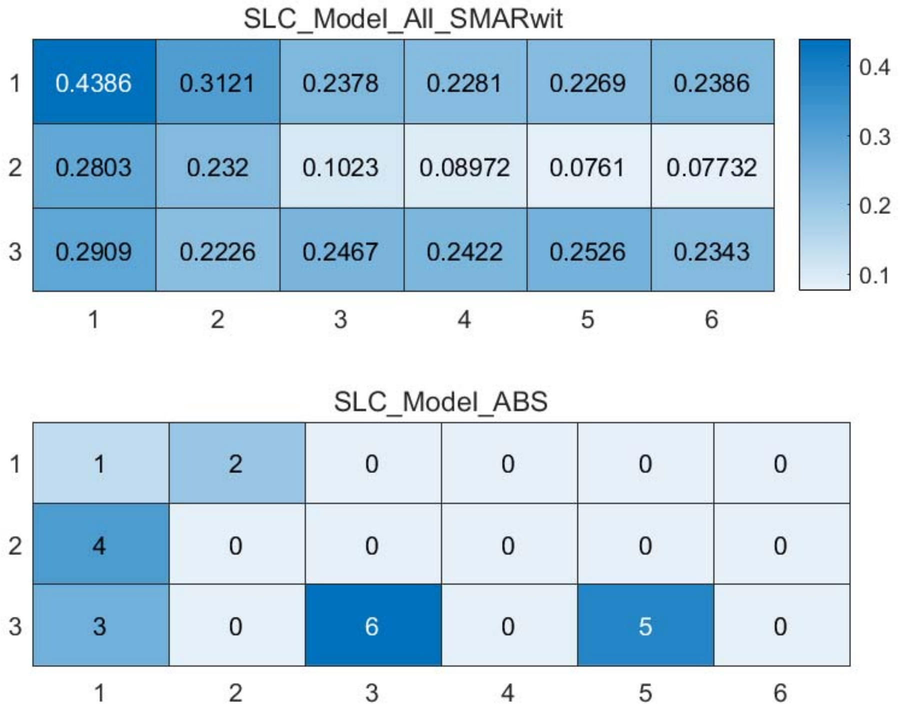


Figure S141: The ABS mode of meta classifiers for *Data_Tasic* dataset.

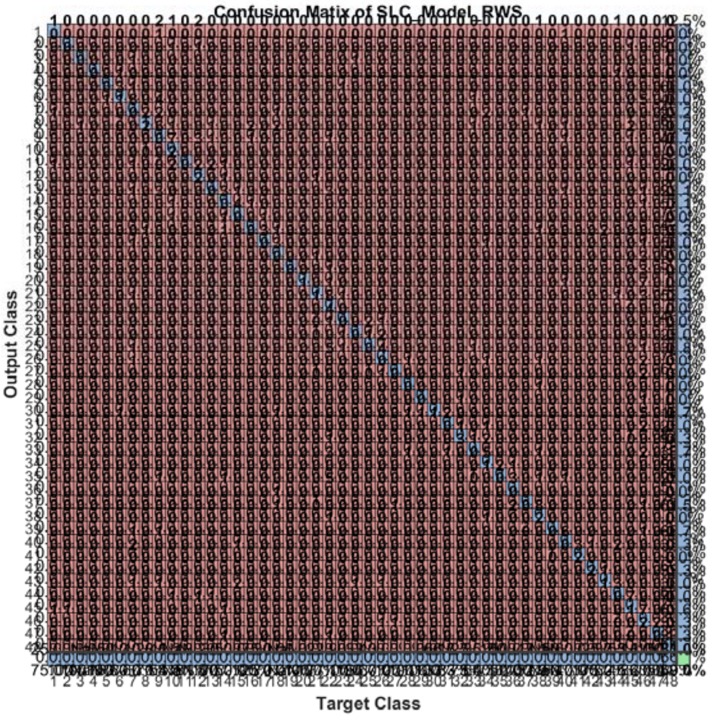


Figure S142: The confusion matrix of ensemble classifier with RWS mode for *Data_Tasic* dataset. (*Note*: the confusion matrix is too large to display completely in one figure)

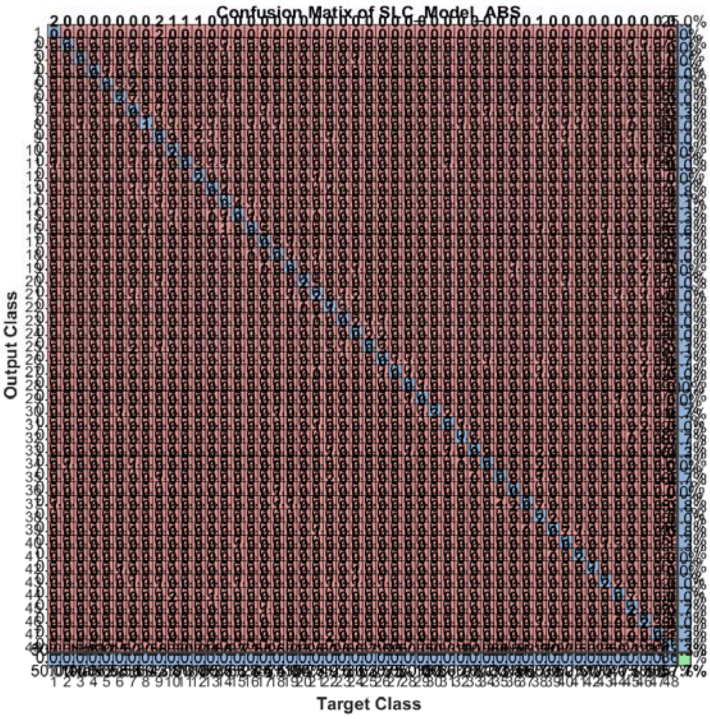


Figure S143: The confusion matrix of ensemble classifier with ABS mode for *Data_Tasic* dataset. (*Note*: the confusion matrix is too large to display completely in one figure)

#### 2.16 The running parameters and output figures for *Data_Tasic* dataset (all corr)

Table S14: The running parameters for *Data_Tasic* dataset.

```
load('MPSSC\Data_Tasic.mat'); true_labs = true_labs-double(true_labs>25);
[in_X_SLC,class_num,slice_tik,binary_mod] = slicematrix(in_X,true_labs,linspace(0,0.06,60));
[in_X_SLC_diff,in_X_SLC_SRCIstd,slice_bst,slice_vle] = slicediffer(in_X_SLC,slice_tik,8);
[SLC_Model_DIS1,istrain,istest] = slicemethod(in_X_SLC,true_labs,slice_tik,slice_bst,6,'correlation','inverse',3);
[SLC_Model_DIS2,istrain,istest] = slicemethod(in_X_SLC,true_labs,slice_tik,slice_bst,8,'correlation','inverse',3);
[SLC_Model_DIS3,istrain,istest] = slicemethod(in_X_SLC,true_labs,slice_tik,slice_bst,10,'correlation','inverse',3);
[SLC_Model_All,SLC_Model_All_SMARwit] = sliceweight(SLC_Model_DIS1,SLC_Model_DIS2,SLC_Model_DIS3);
[SLC_Model_RWS,SLC_Model_All_SMESrws] = sliceswitch(SLC_Model_All,SLC_Model_All_SMARwit,'rws',2);
[SLC_Model_ABS,SLC_Model_All_SMESabs] = sliceswitch(SLC_Model_All,SLC_Model_All_SMARwit,'abs',6);
[SLC_Model_FIT1,FIT1_accuracy] = sliceprerws(in_X(istest,:),true_labs(istest),binary_mod,class_num,slice_vle,SLC_Model_RWS);
[SLC_Model_FIT2,FIT2_accuracy] = slicepreabs(in_X(istest,:),true_labs(istest),binary_mod,class_num,slice_vle,SLC_Model_ABS);
```

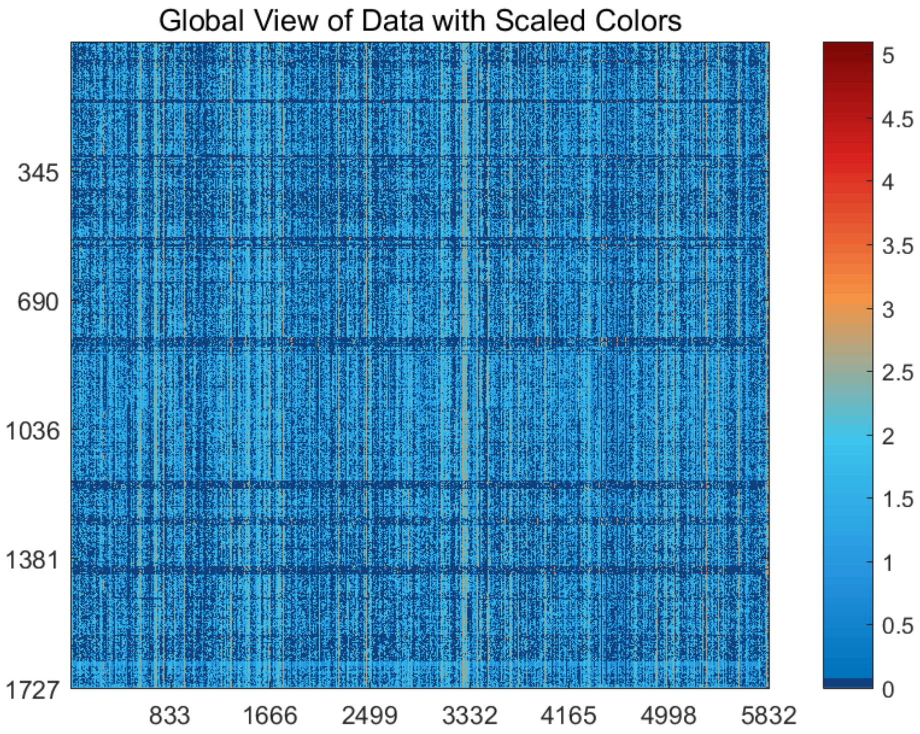


Figure S144: The global view of *Data_Tasic* dataset with scaled colors.

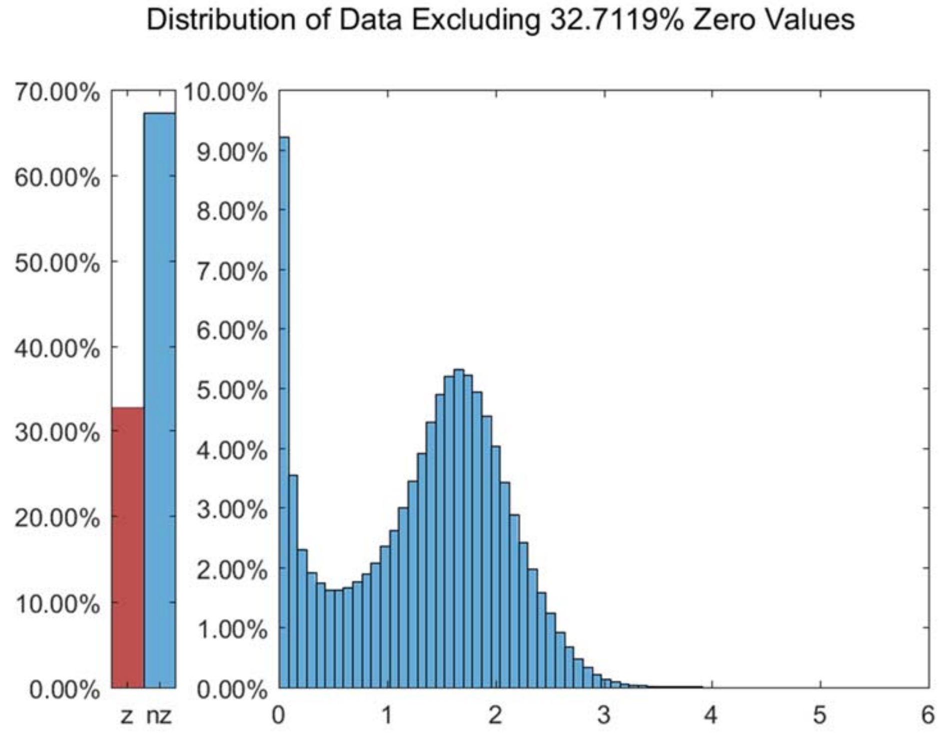


Figure S145: The distribution of *Data_Tasic* dataset excluding all zero values.

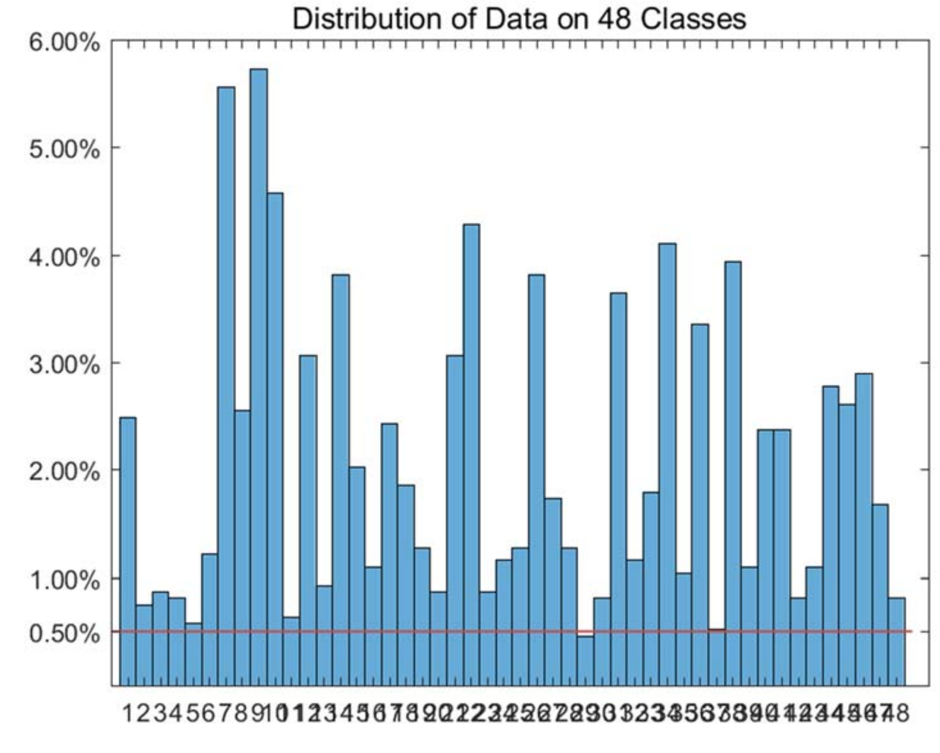


Figure S146: The distribution of *Data_Tasic* dataset on every classes.

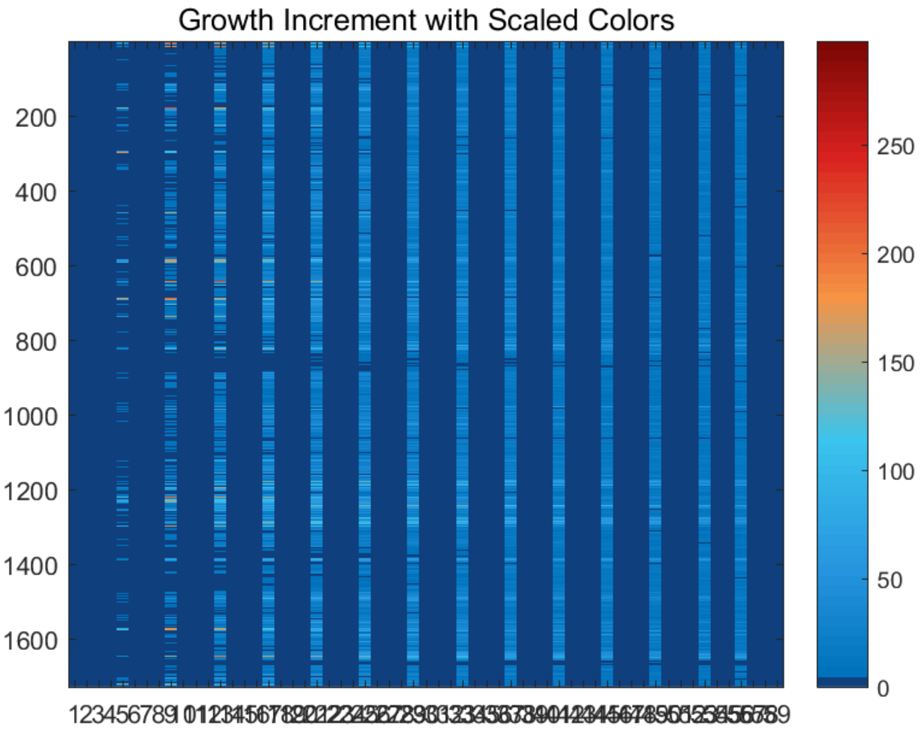


Figure S147: The growth increment with scaled colors for *Data_Tasic* dataset.

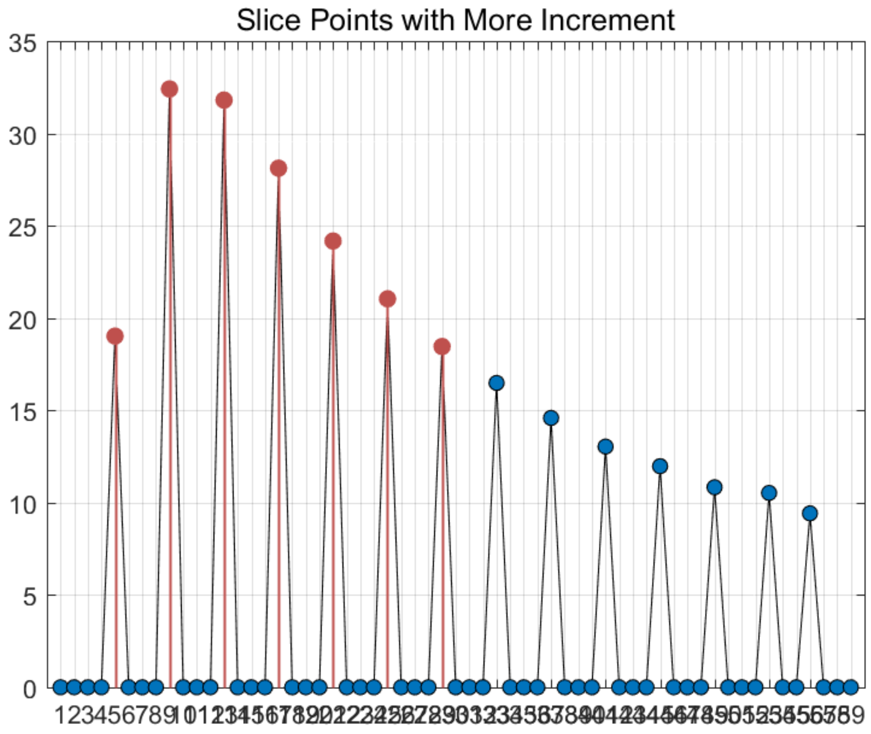


Figure S148: The slice points with more increment for *Data_Tasic* dataset.

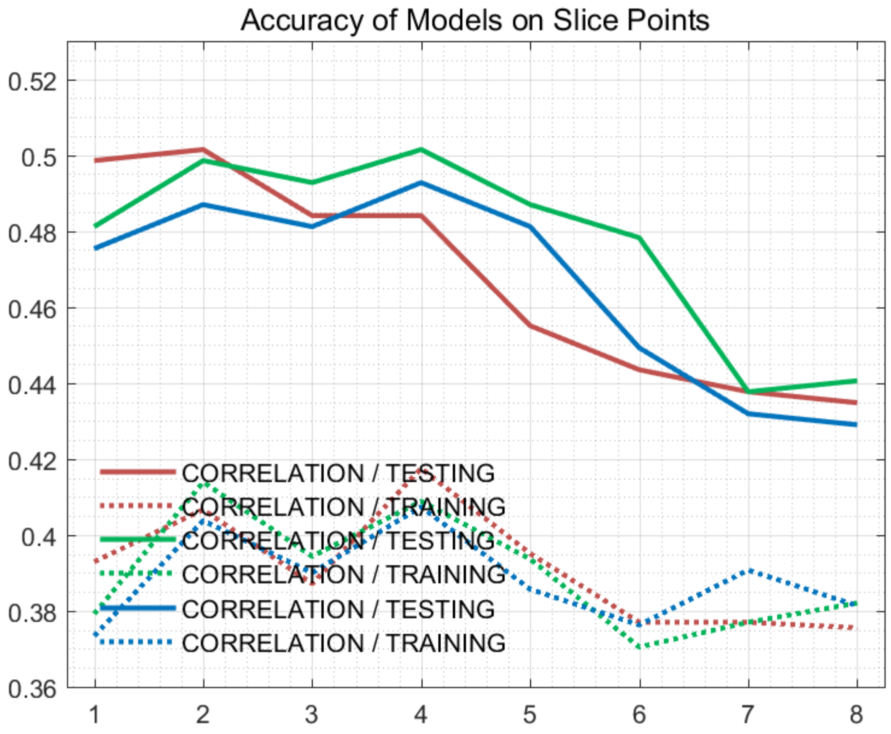


Figure S149: The accuracy of models on every slice points for *Data_Tasic* dataset.

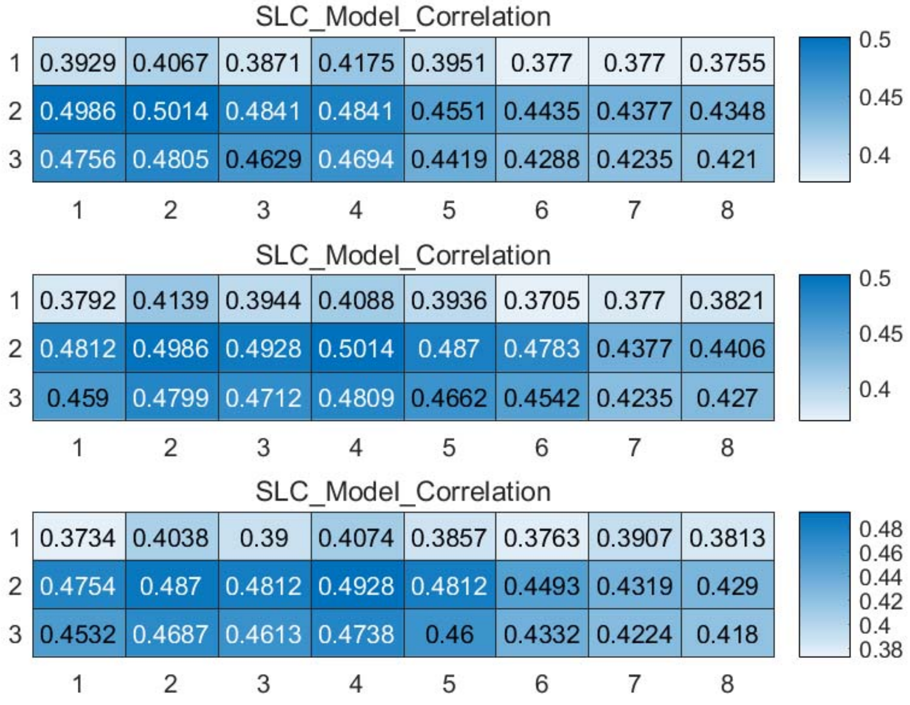


Figure S150: The weighted accuracy on every slice points for *Data_Tasic* dataset.

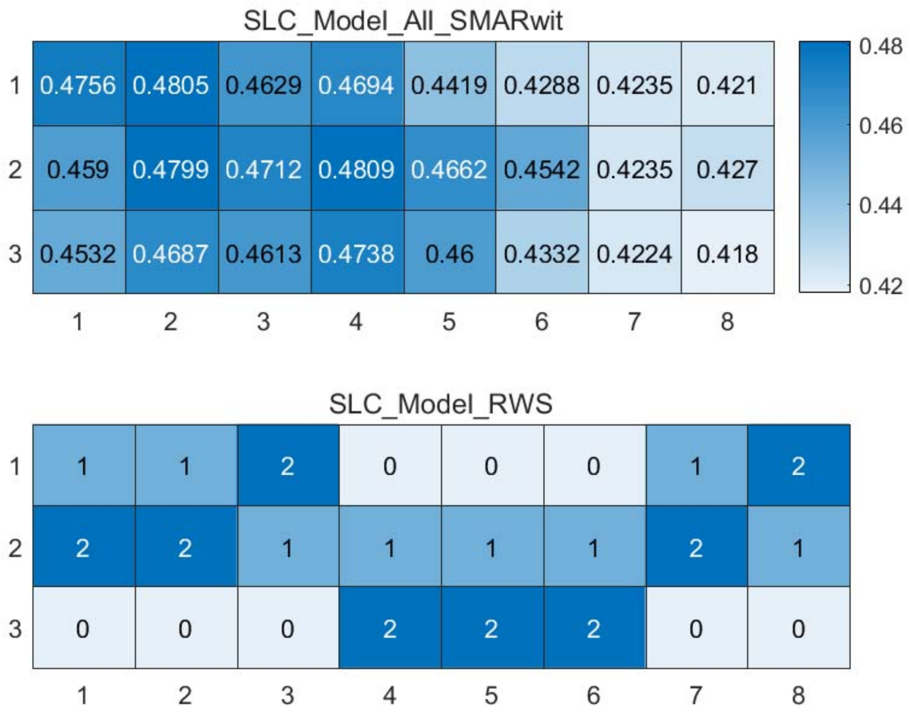


Figure S151: The RWS mode of meta classifiers for *Data_Tasic* dataset.

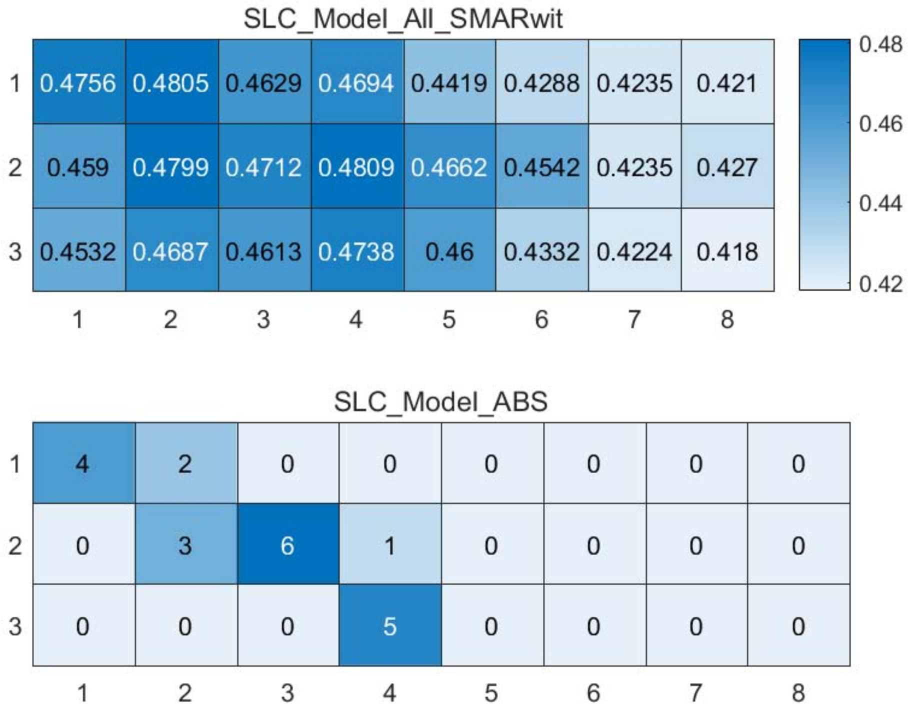


Figure S152: The ABS mode of meta classifiers for *Data_Tasic* dataset.

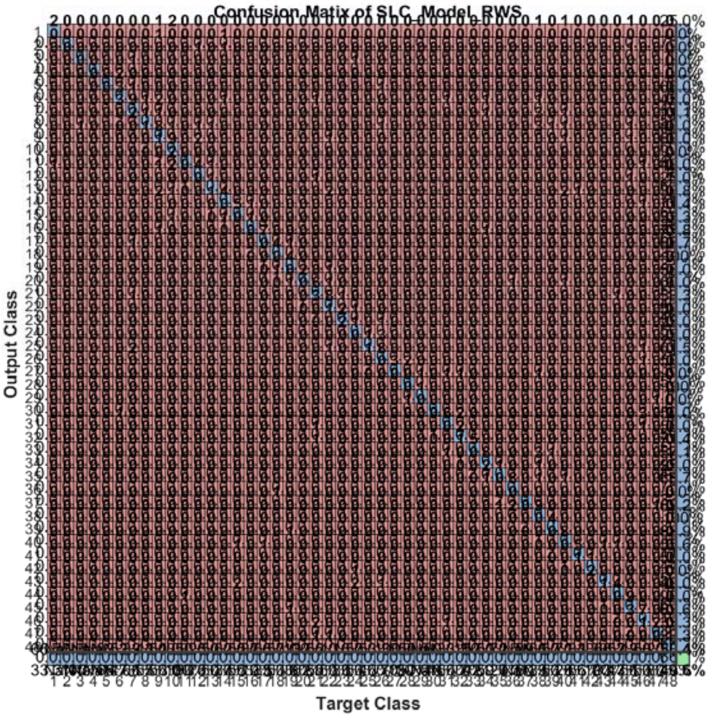


Figure S153: The confusion matrix of ensemble classifier with RWS mode for *Data_Tasic* dataset. (*Note*: the confusion matrix is too large to display completely in one figure)

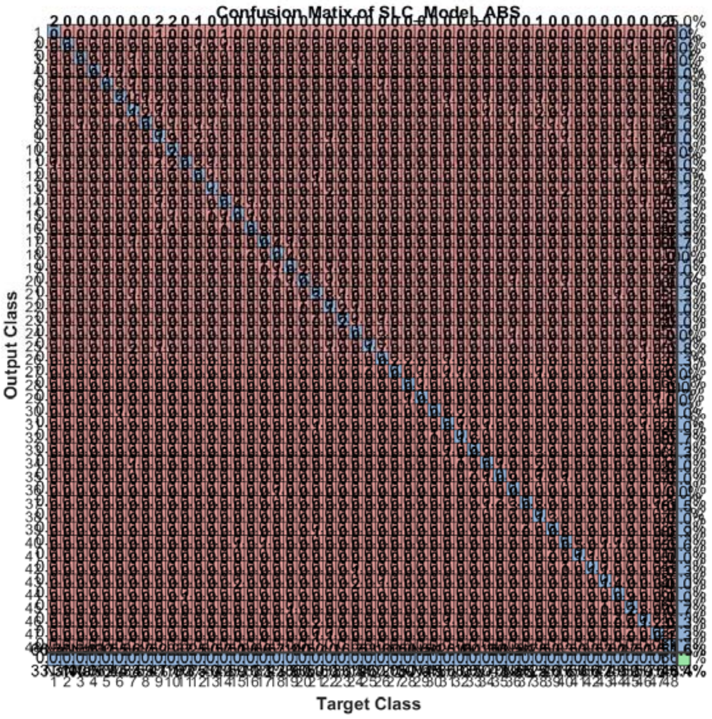


Figure S154: The confusion matrix of ensemble classifier with ABS mode for *Data_Tasic* dataset. (*Note*: the confusion matrix is too large to display completely in one figure)

#### 2.17 The running parameters and output figures for *Data_Ting* dataset

Table S15: The running parameters for *Data_Ting* dataset.

```
load('MPSSC\Data_Ting.mat'); true_labs = true_labs';
[in_X_SLC,class_num,slice_tik,binary_mod] = slicematrix(in_X,true_labs,linspace(0,0.4,60));
[in_X_SLC_diff,in_X_SLC_SRCIstd,slice_bst,slice_vle] = slicediffer(in_X_SLC,slice_tik,9);
[SLC_Model_DIS1,istrain,istest] = slicemethod(in_X_SLC,true_labs,slice_tik,slice_bst,1,'correlation','inverse',5);
[SLC_Model_DIS2,istrain,istest] = slicemethod(in_X_SLC,true_labs,slice_tik,slice_bst,1,'jaccard','inverse',5);
[SLC_Model_DIS3,istrain,istest] = slicemethod(in_X_SLC,true_labs,slice_tik,slice_bst,1,'cosine','inverse',5);
[SLC_Model_All,SLC_Model_All_SMARwit] = sliceweight(SLC_Model_DIS1,SLC_Model_DIS2,SLC_Model_DIS3);
[SLC_Model_RWS,SLC_Model_All_SMESrws] = sliceswitch(SLC_Model_All,SLC_Model_All_SMARwit,'rws',2);
[SLC_Model_ABS,SLC_Model_All_SMESabs] = sliceswitch(SLC_Model_All,SLC_Model_All_SMARwit,'abs',9);
[SLC_Model_FIT1,FIT1_accuracy] = sliceprerws(in_X(istest,:),true_labs(istest),binary_mod,class_num,slice_vle,SLC_Model_RWS);
[SLC_Model_FIT2,FIT2_accuracy] = slicepreabs(in_X(istest,:),true_labs(istest),binary_mod,class_num,slice_vle,SLC_Model_ABS);
```

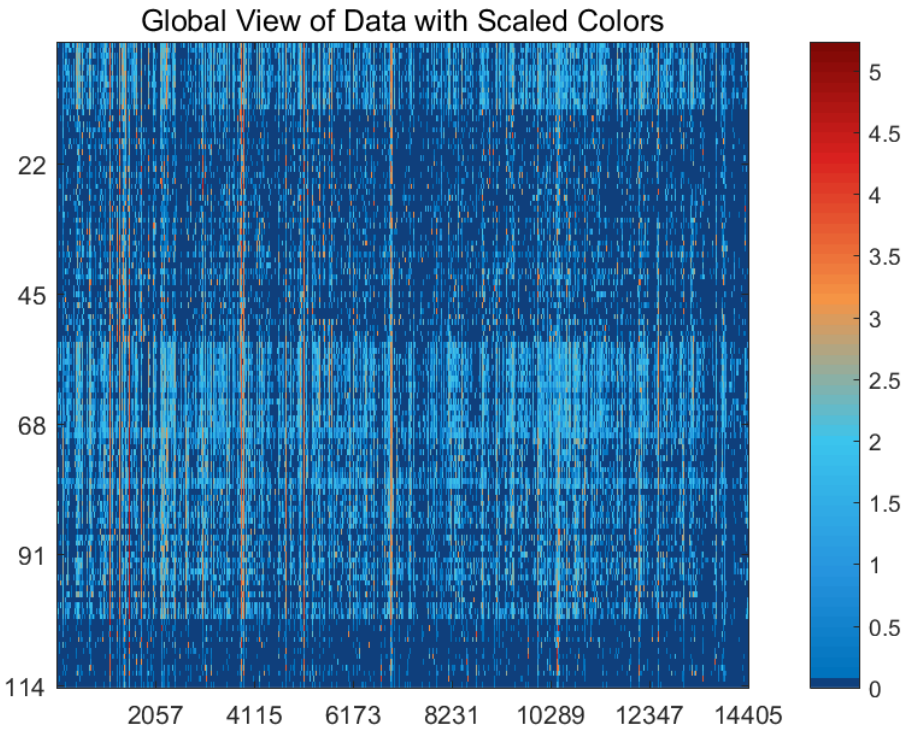


Figure S155: The global view of *Data_Ting* dataset with scaled colors.

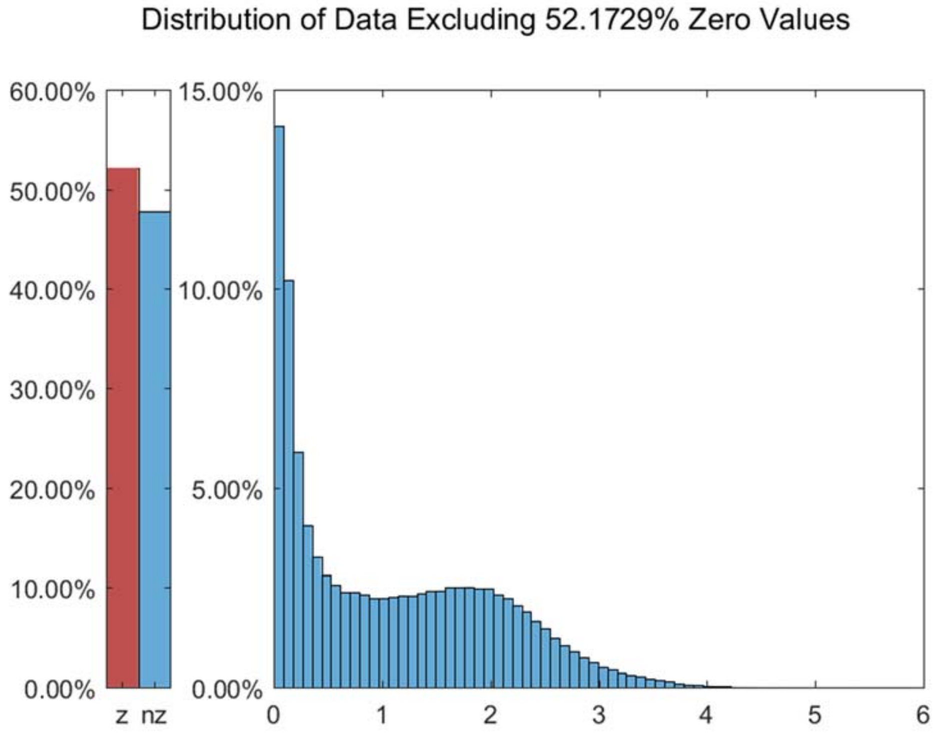


Figure S156: The distribution of *Data_Ting* dataset excluding all zero values.

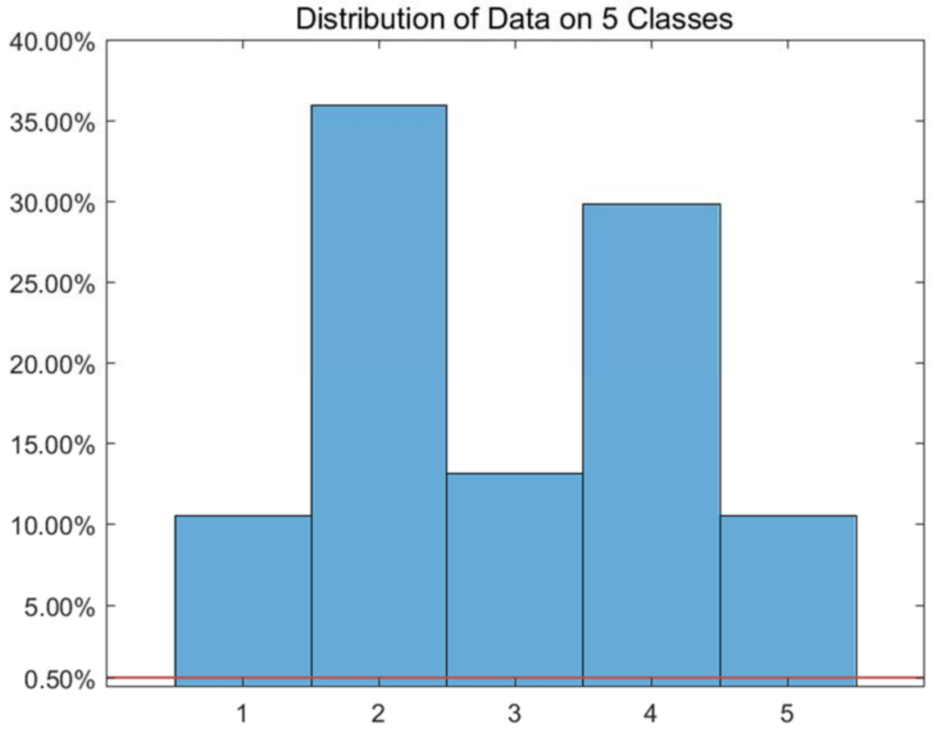


Figure S157: The distribution of *Data_Ting* dataset on every classes.

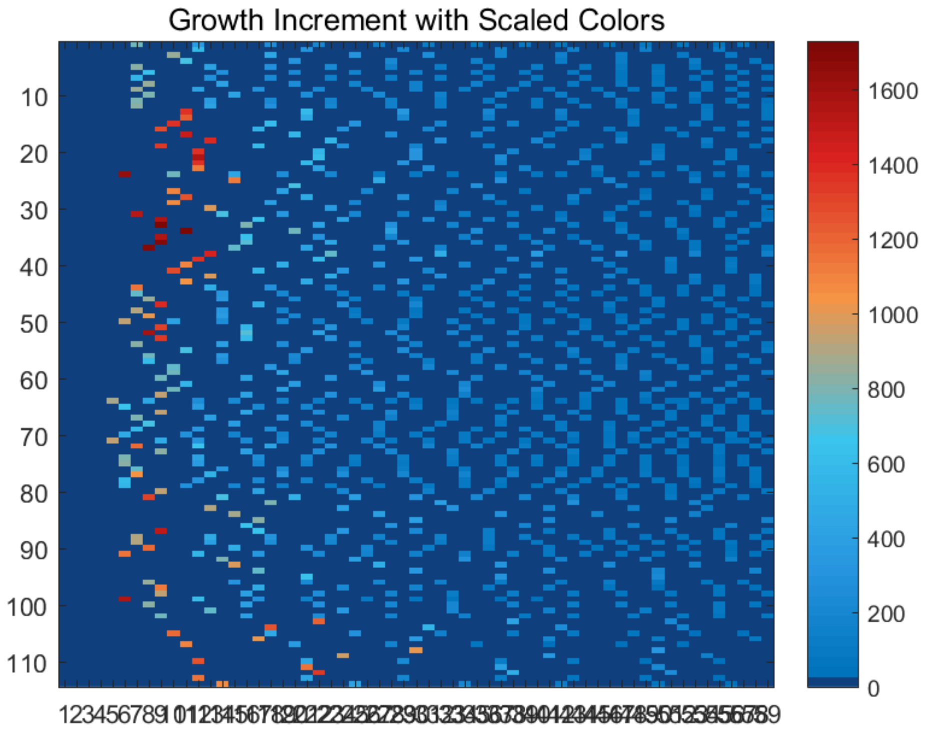


Figure S158: The growth increment with scaled colors for *Data_Ting* dataset.

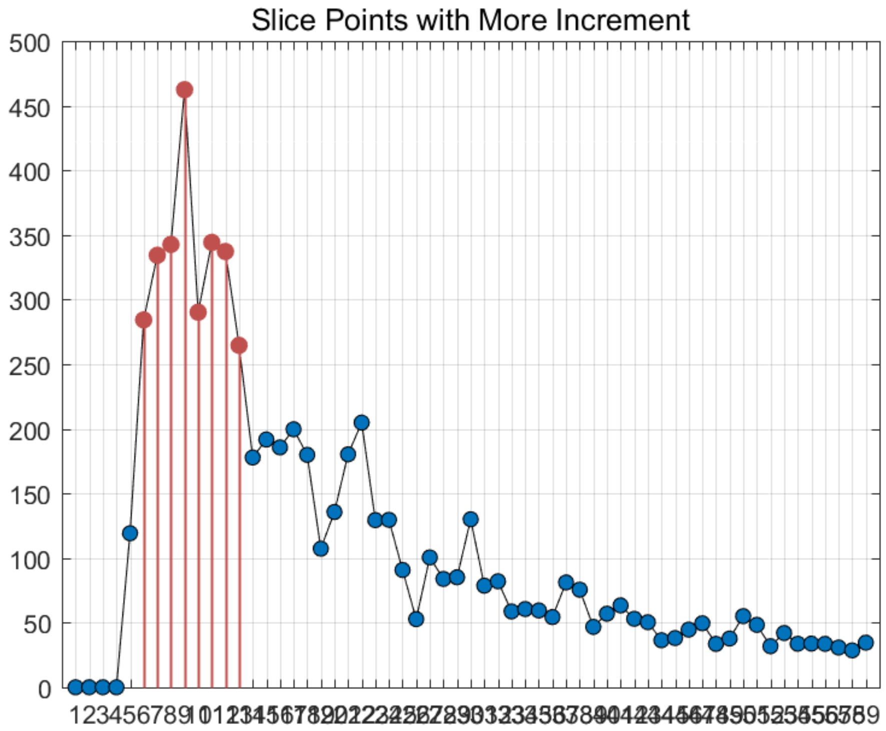


Figure S159: The slice points with more increment for *Data_Ting* dataset.

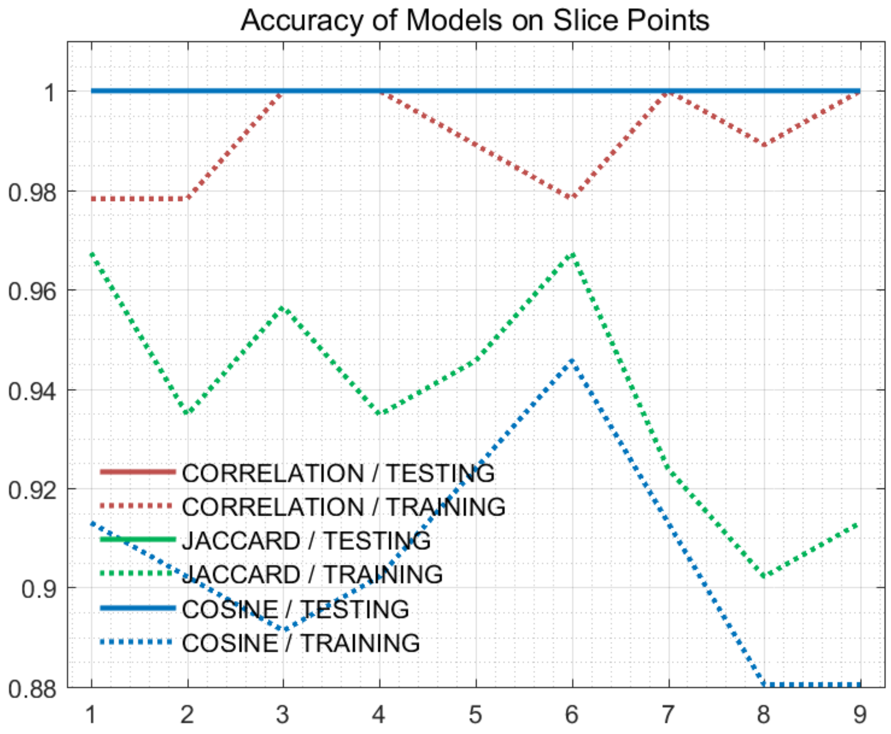


Figure S160: The accuracy of models on every slice points for *Data_Ting* dataset.

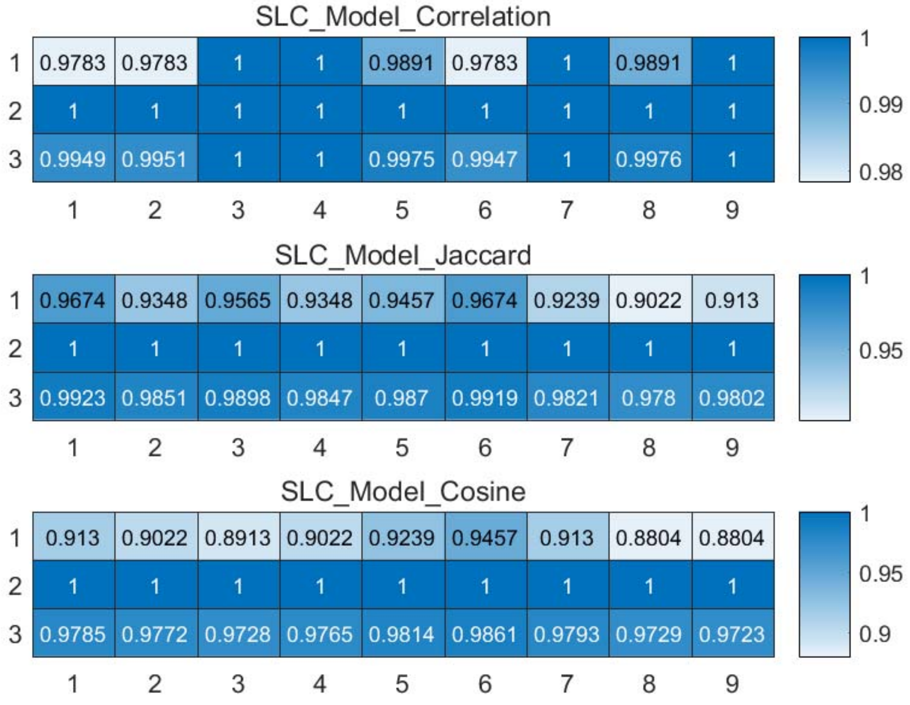


Figure S161: The weighted accuracy on every slice points for *Data_Ting* dataset.

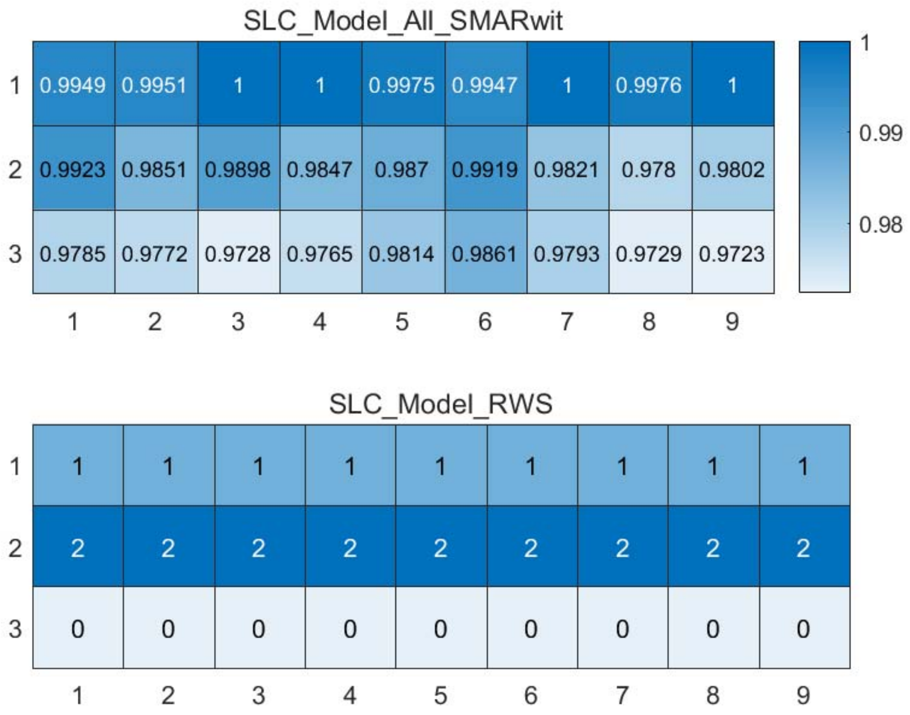


Figure S162: The RWS mode of meta classifiers for *Data_Ting* dataset.

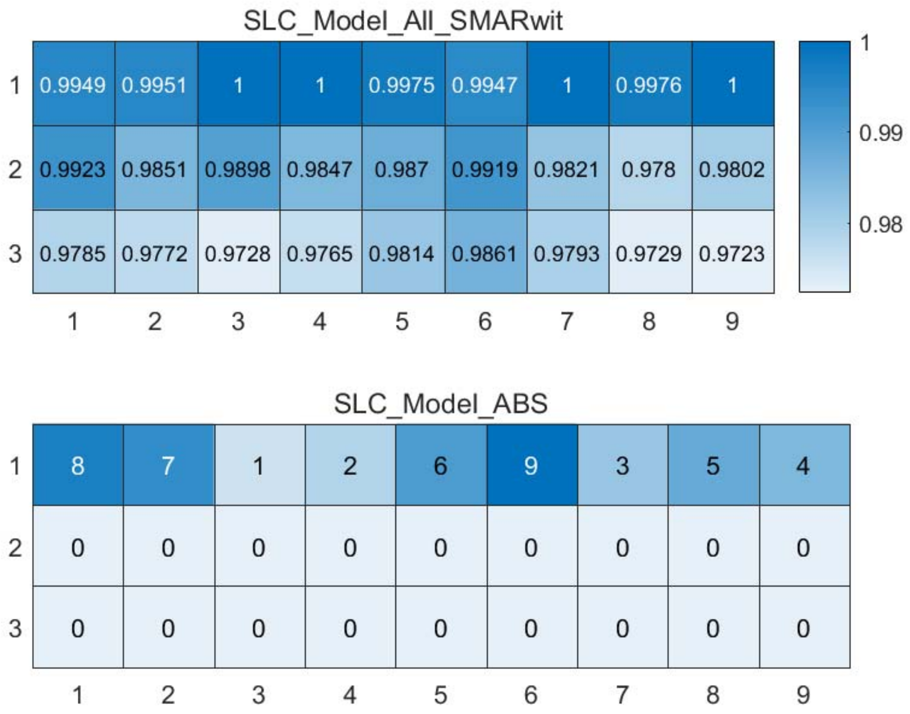


Figure S163: The ABS mode of meta classifiers for *Data_Ting* dataset.

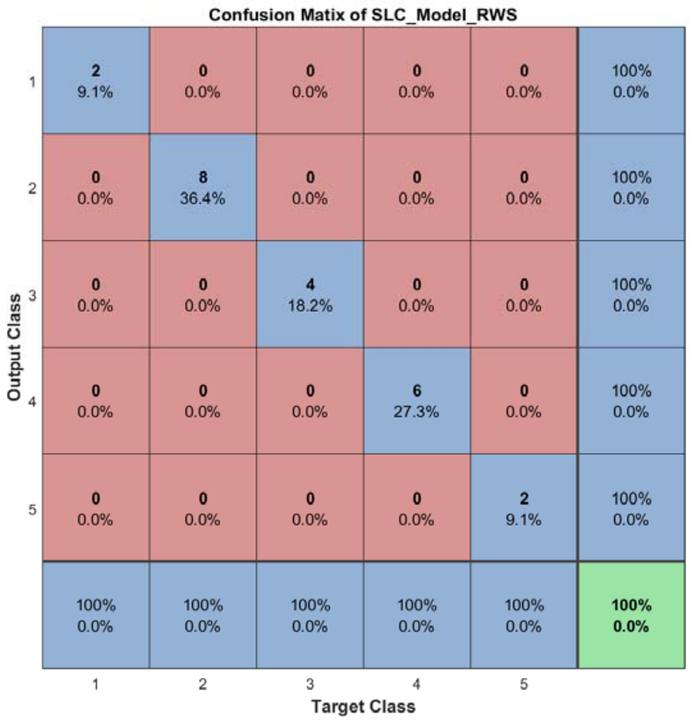


Figure S164: The confusion matrix of ensemble classifier with RWS mode for *Data_Ting* dataset.

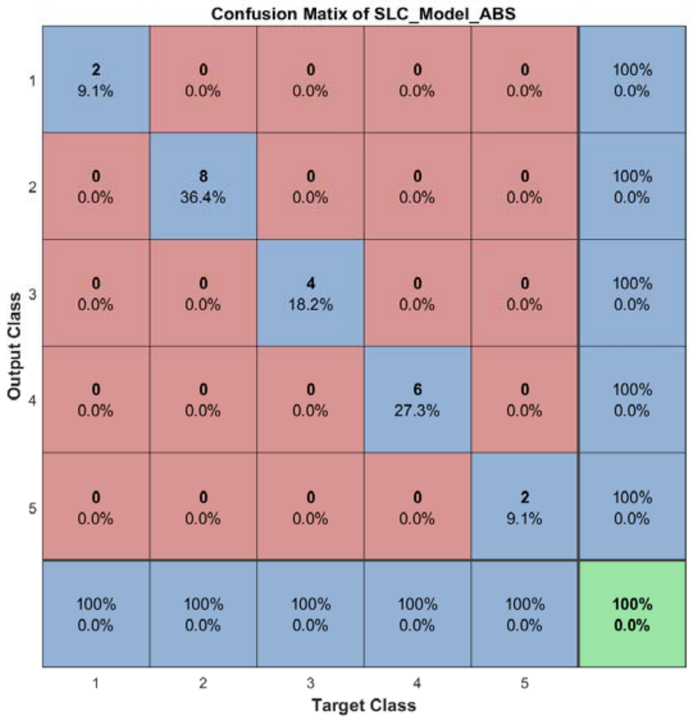


Figure S165: The confusion matrix of ensemble classifier with ABS mode for *Data_Ting* dataset.

#### 2.18 The running parameters and output figures for *Data_Treutlin* dataset

Table S16: The running parameters for *Data_Treutlin* dataset.

```
load('MPSSC\Data_Treutlin.mat'); in_X = 0.3*in_X;
[in_X_SLC,class_num,slice_tik,binary_mod] = slicematrix(in_X,true_labs,linspace(0,max(max(in_X)),90));
[in_X_SLC_diff,in_X_SLC_SRCIstd,slice_bst,slice_vle] = slicedabout(in_X_SLC,slice_tik,21);
[SLC_Model_DIS1,istrain,istest] = slicemethod(in_X_SLC,true_labs,slice_tik,slice_bst,1,'correlation','squaredinverse',2);
[SLC_Model_DIS2,istrain,istest] = slicemethod(in_X_SLC,true_labs,slice_tik,slice_bst,1,'jaccard','squaredinverse',2);
[SLC_Model_DIS3,istrain,istest] = slicemethod(in_X_SLC,true_labs,slice_tik,slice_bst,1,'cosine','squaredinverse',2);
[SLC_Model_All,SLC_Model_All_SMARwit] = sliceweight(SLC_Model_DIS1,SLC_Model_DIS2,SLC_Model_DIS3);
[SLC_Model_RWS,SLC_Model_All_SMESrws] = sliceswitch(SLC_Model_All,SLC_Model_All_SMARwit,'rws',2);
[SLC_Model_ABS,SLC_Model_All_SMESabs] = sliceswitch(SLC_Model_All,SLC_Model_All_SMARwit,'abs',9);
[SLC_Model_FIT1,FIT1_accuracy] = sliceprerws(in_X(istest,:),true_labs(istest),binary_mod,class_num,slice_vle,SLC_Model_RWS);
[SLC_Model_FIT2,FIT2_accuracy] = slicepreabs(in_X(istest,:),true_labs(istest),binary_mod,class_num,slice_vle,SLC_Model_ABS);
```

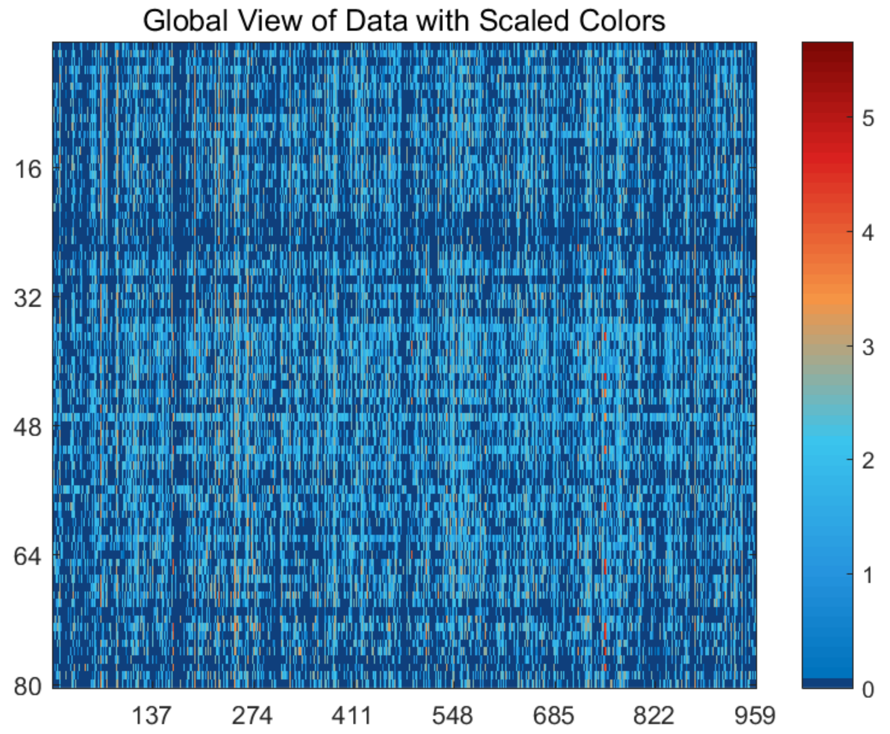


Figure S166: The global view of *Data_Treutlin* dataset with scaled colors.

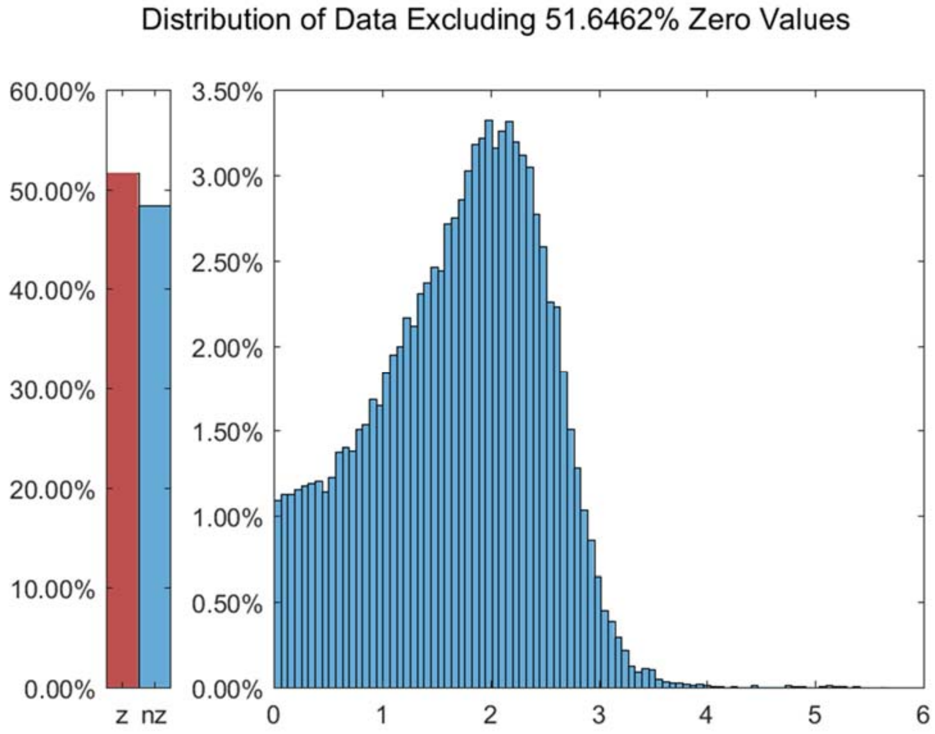


Figure S167: The distribution of *Data_Treutlin* dataset excluding all zero values.

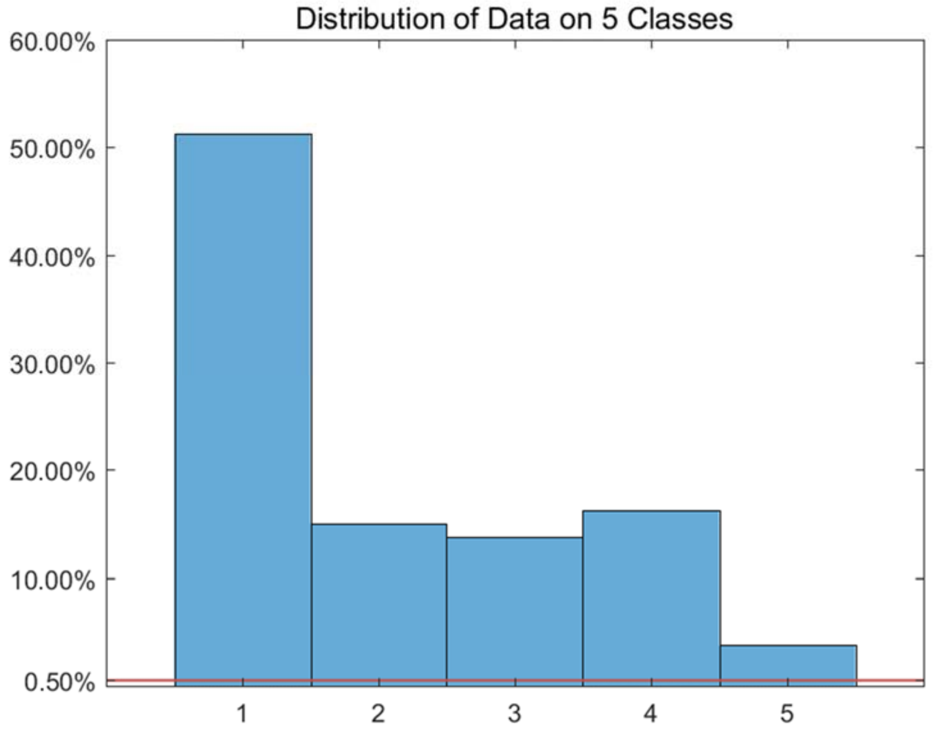


Figure S168: The distribution of *Data_Treutlin* dataset on every classes.

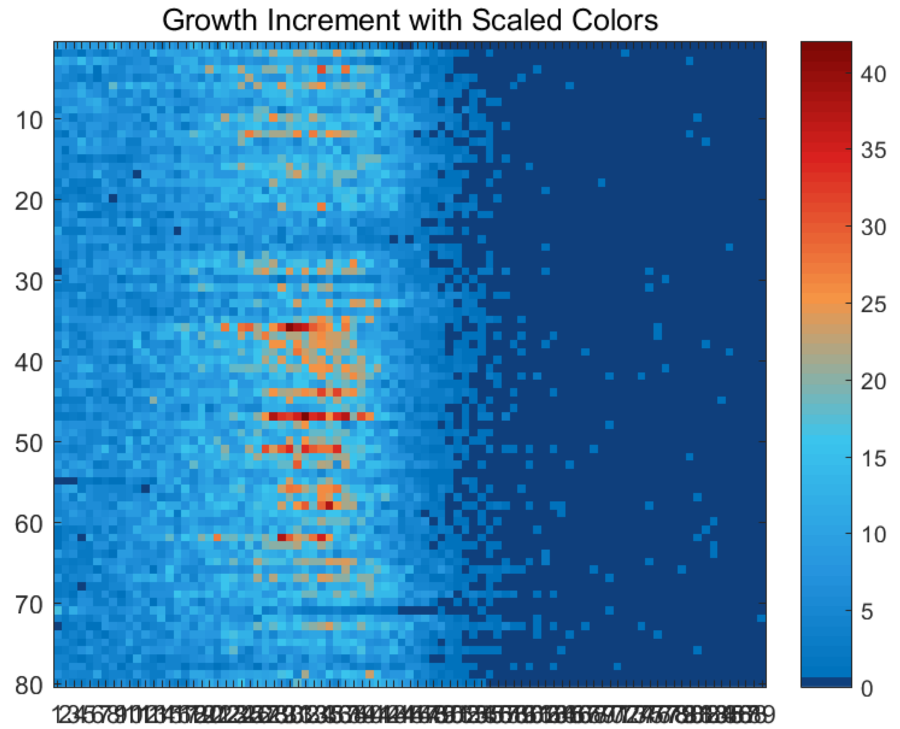


Figure S169: The growth increment with scaled colors for *Data_Treutlin* dataset.

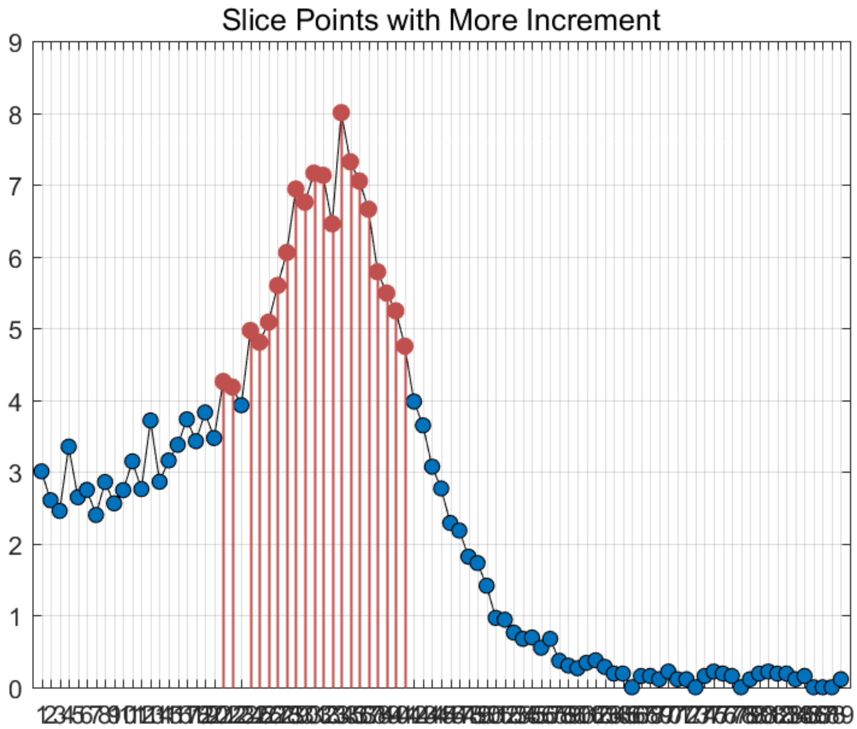


Figure S170: The slice points with more increment for *Data_Treutlin* dataset.

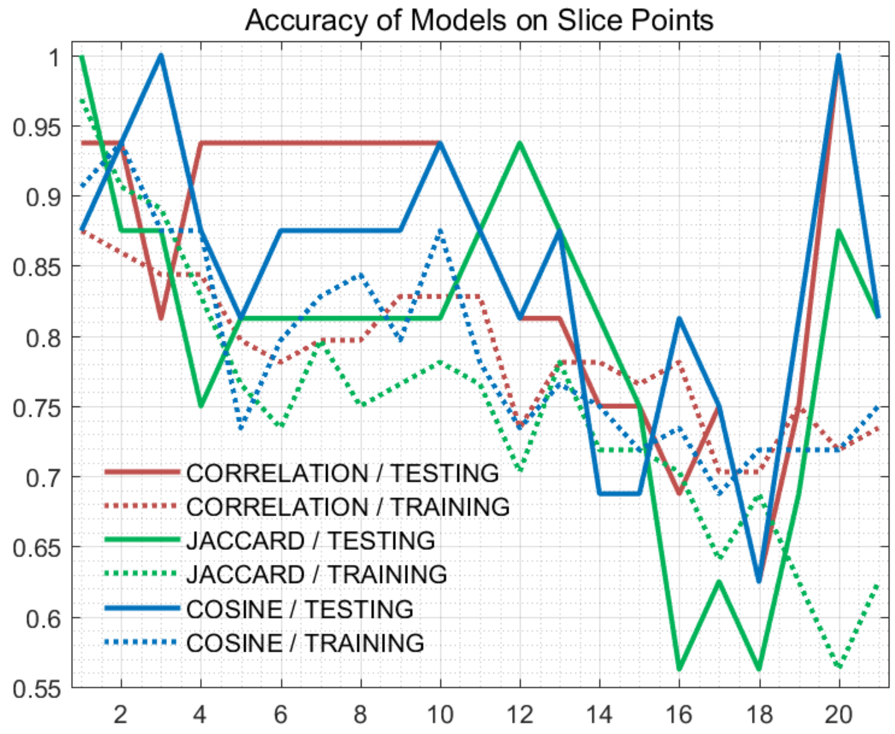


Figure S171: The accuracy of models on every slice points for *Data_Treutlin* dataset.

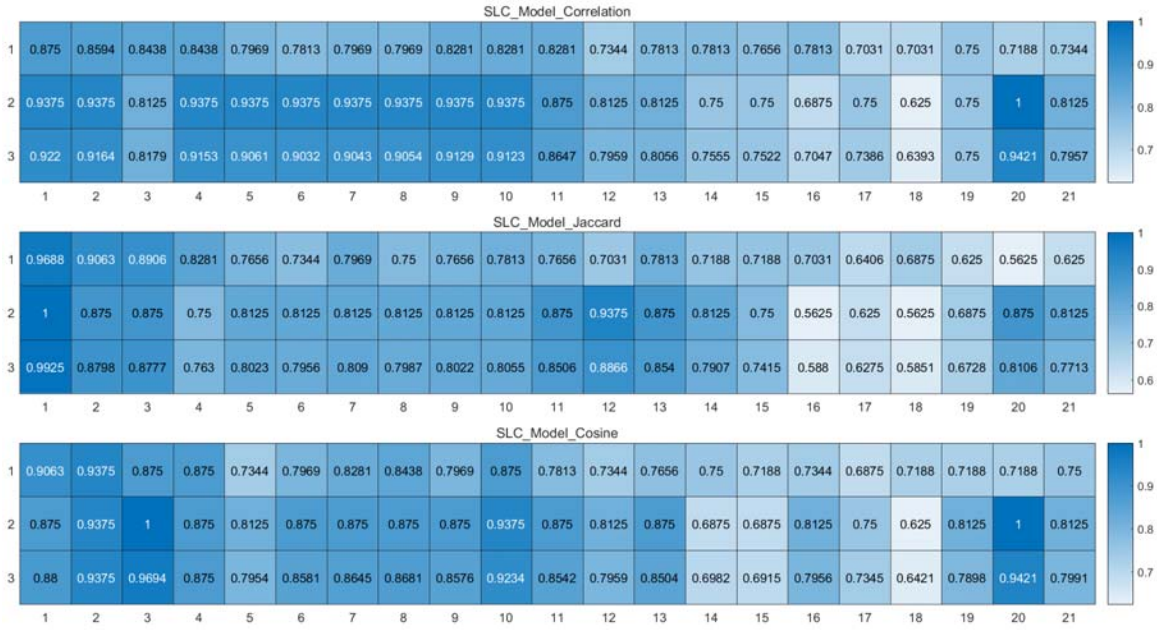


Figure S172: The weighted accuracy on every slice points for *Data_Treutlin* dataset.

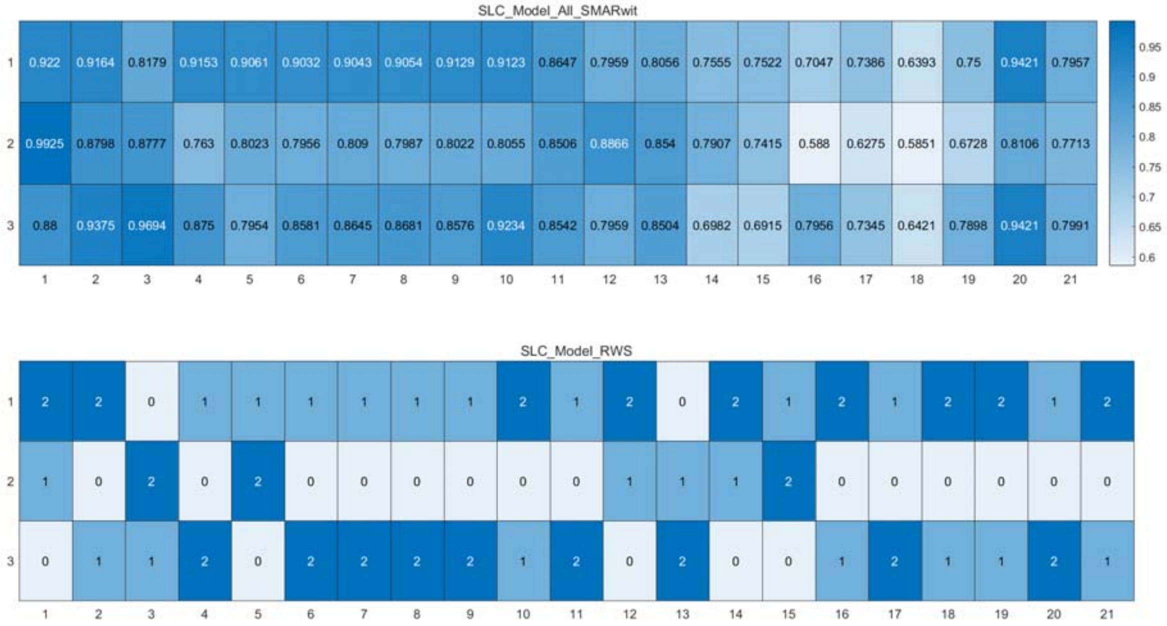


Figure S173: The RWS mode of meta classifiers for *Data_Treutlin* dataset.

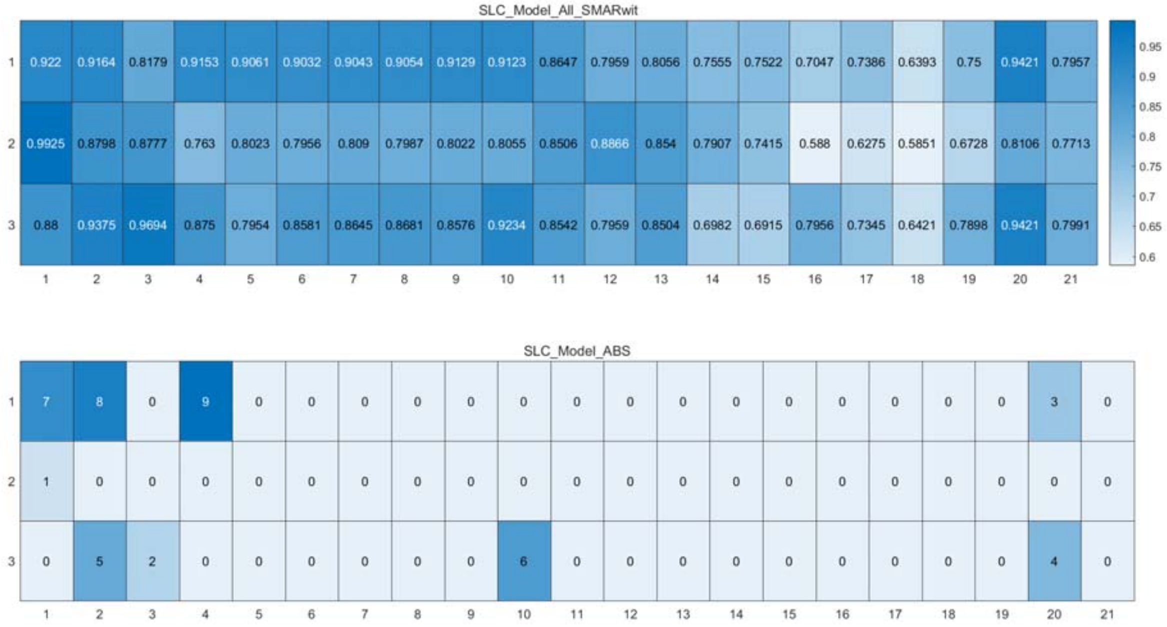


Figure S174: The ABS mode of meta classifiers for *Data_Treutlin* dataset.

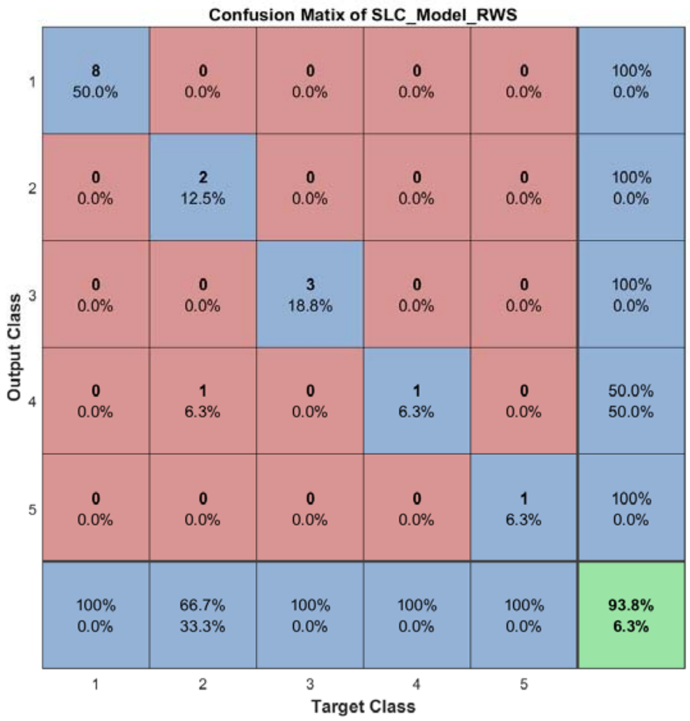


Figure S175: The confusion matrix of ensemble classifier with RWS mode for *Data_Treutlin* dataset.

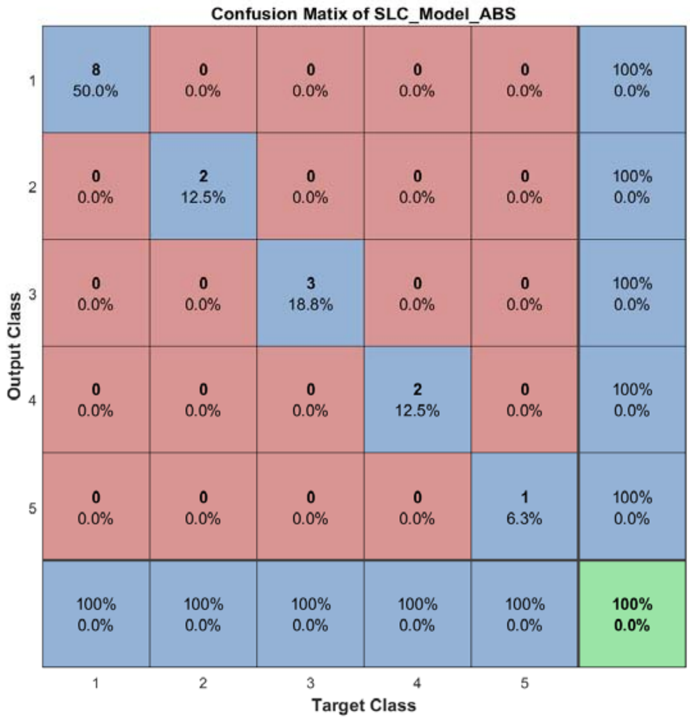


Figure S176: The confusion matrix of ensemble classifier with ABS mode for *Data_Treutlin* dataset.

#### 2.19 The running parameters and output figures for *Data_Zeisel* dataset

Table S17: The running parameters for *Data_Zeisel* dataset.

```
load('MPSSC\Data_Zeisel.mat');
[in_X_SLC,class_num,slice_tik,binary_mod] = slicematrix(in_X,true_labs,0:0.3:2.1);
[in_X_SLC_diff,in_X_SLC_SRCIstd,slice_bst,slice_vle] = slicediffer(in_X_SLC,slice_tik,6);
[SLC_Model_DIS1,istrain,istest] = slicemethod(in_X_SLC,true_labs,slice_tik,slice_bst,1,'correlation','inverse',3);
[SLC_Model_DIS2,istrain,istest] = slicemethod(in_X_SLC,true_labs,slice_tik,slice_bst,1,'jaccard','inverse',3);
[SLC_Model_DIS3,istrain,istest] = slicemethod(in_X_SLC,true_labs,slice_tik,slice_bst,1,'cosine','inverse',3);
[SLC_Model_All,SLC_Model_All_SMARwit] = sliceweight(SLC_Model_DIS1,SLC_Model_DIS2,SLC_Model_DIS3);
[SLC_Model_RWS,SLC_Model_All_SMESrws] = sliceswitch(SLC_Model_All,SLC_Model_All_SMARwit,'rws',2);
[SLC_Model_ABS,SLC_Model_All_SMESabs] = sliceswitch(SLC_Model_All,SLC_Model_All_SMARwit,'abs',6);
[SLC_Model_FIT1,FIT1_accuracy] = sliceprerws(in_X(istest,:),true_labs(istest),binary_mod,class_num,slice_vle,SLC_Model_RWS);
[SLC_Model_FIT2,FIT2_accuracy] = slicepreabs(in_X(istest,:),true_labs(istest),binary_mod,class_num,slice_vle,SLC_Model_ABS);
```

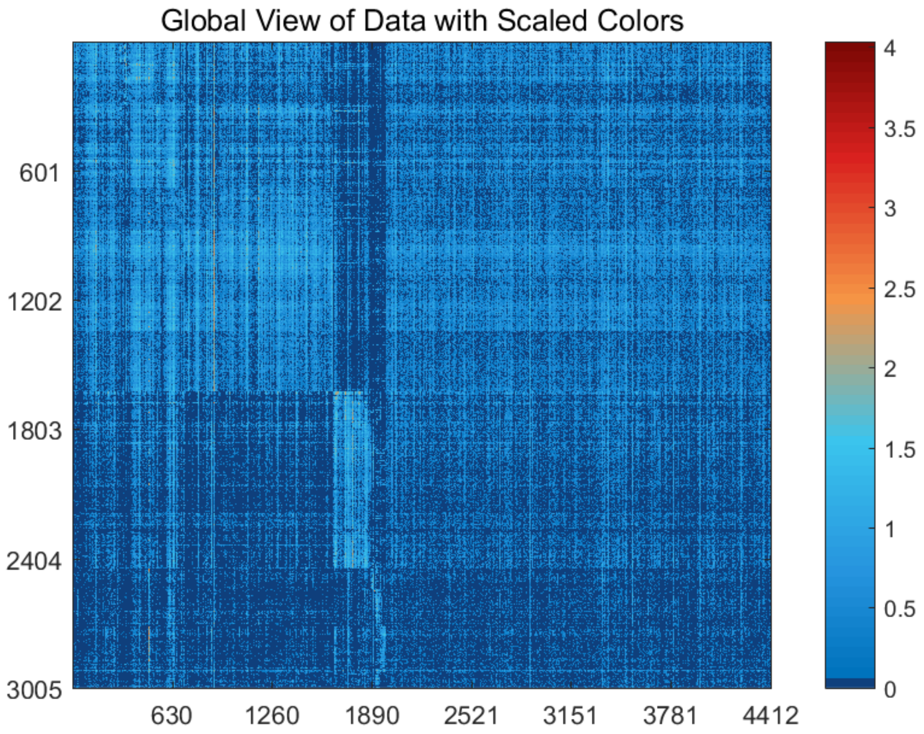


Figure S177: The global view of *Data_Zeisel* dataset with scaled colors.

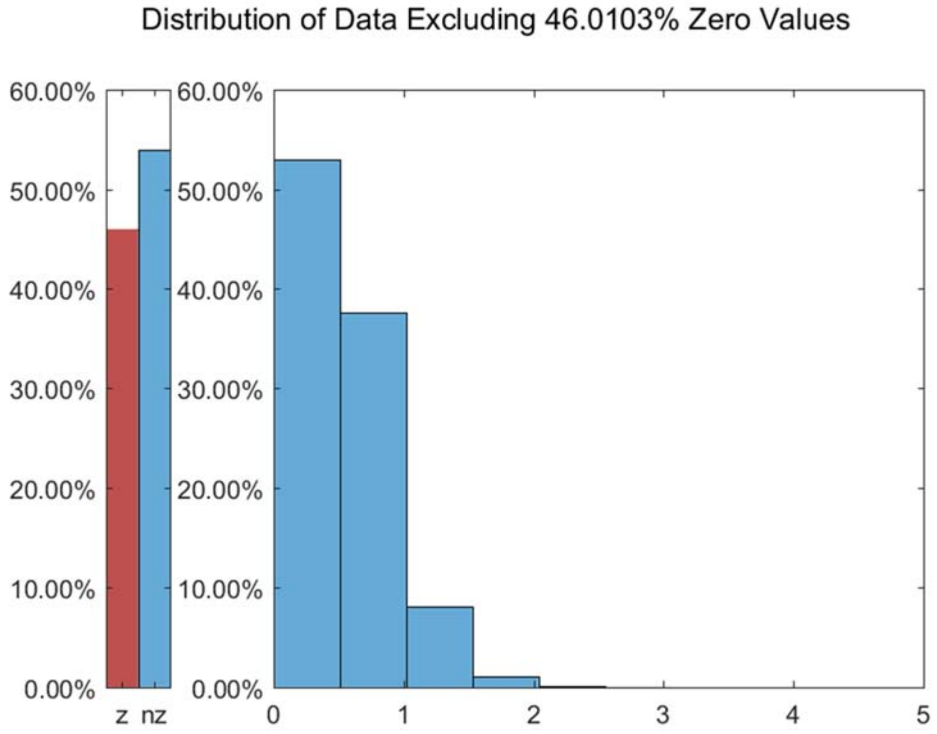


Figure S178: The distribution of *Data_Zeisel* dataset excluding all zero values.

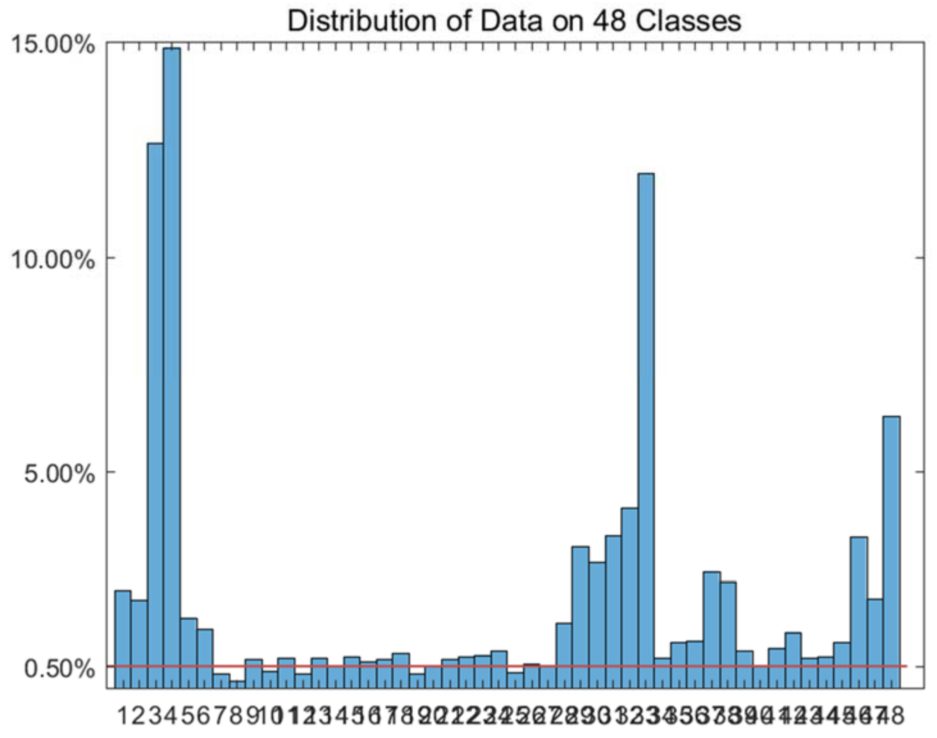


Figure S179: The distribution of *Data_Zeisel* dataset on every classes.

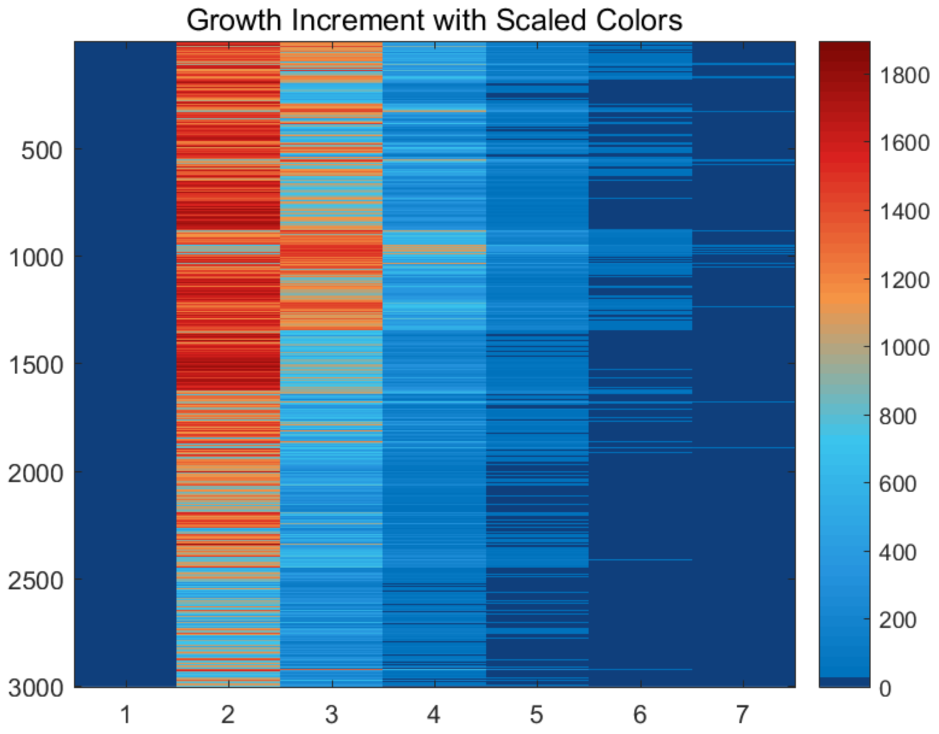


Figure S180: The growth increment with scaled colors for *Data_Zeisel* dataset.

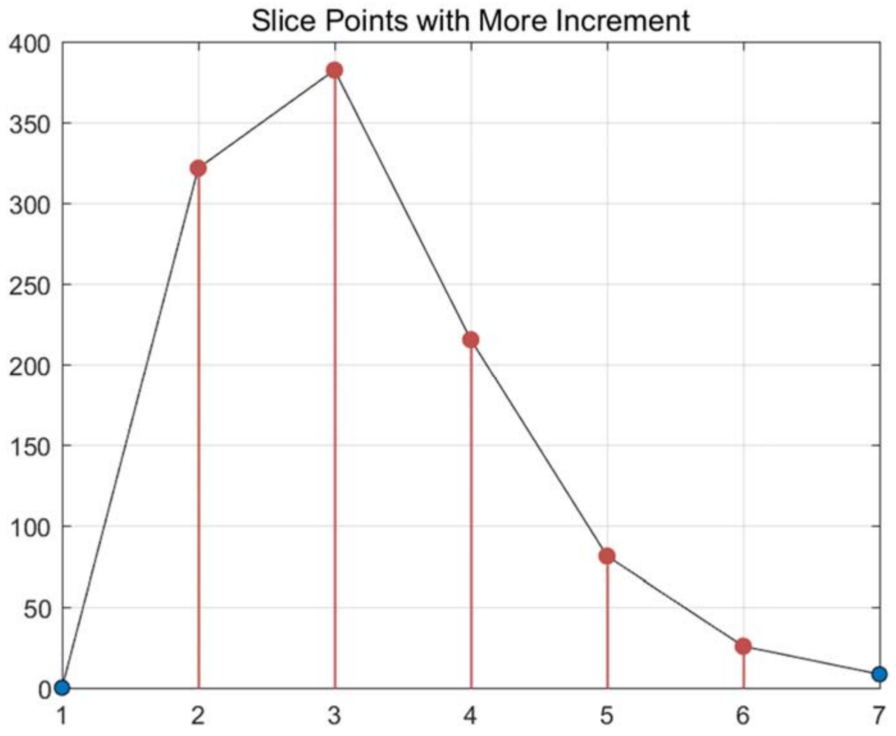


Figure S181: The slice points with more increment for *Data_Zeisel* dataset.

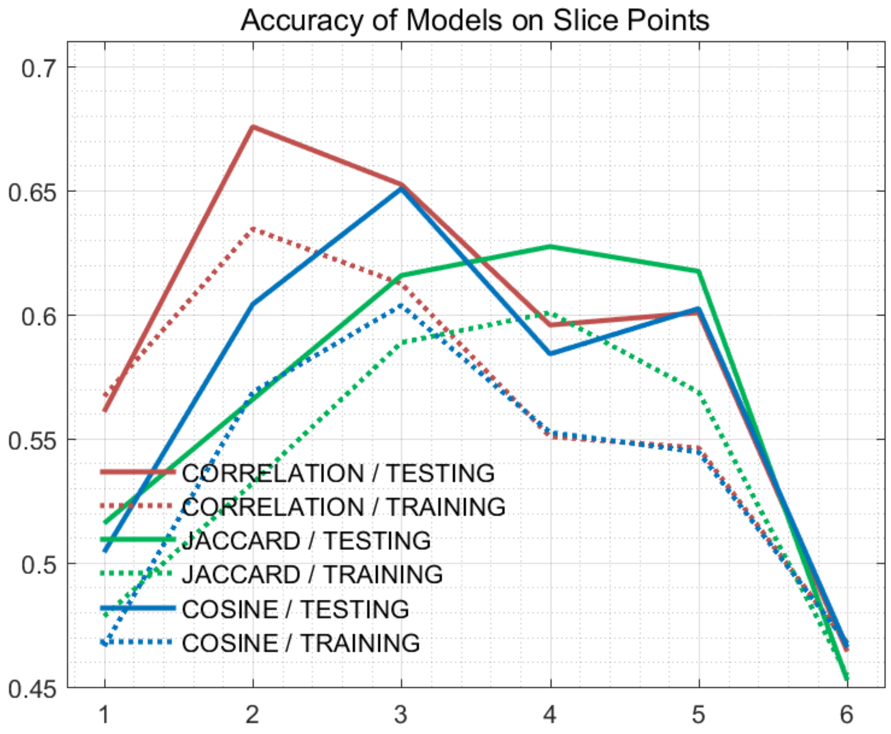


Figure S182: The accuracy of models on every slice points for *Data_Zeisel* dataset.

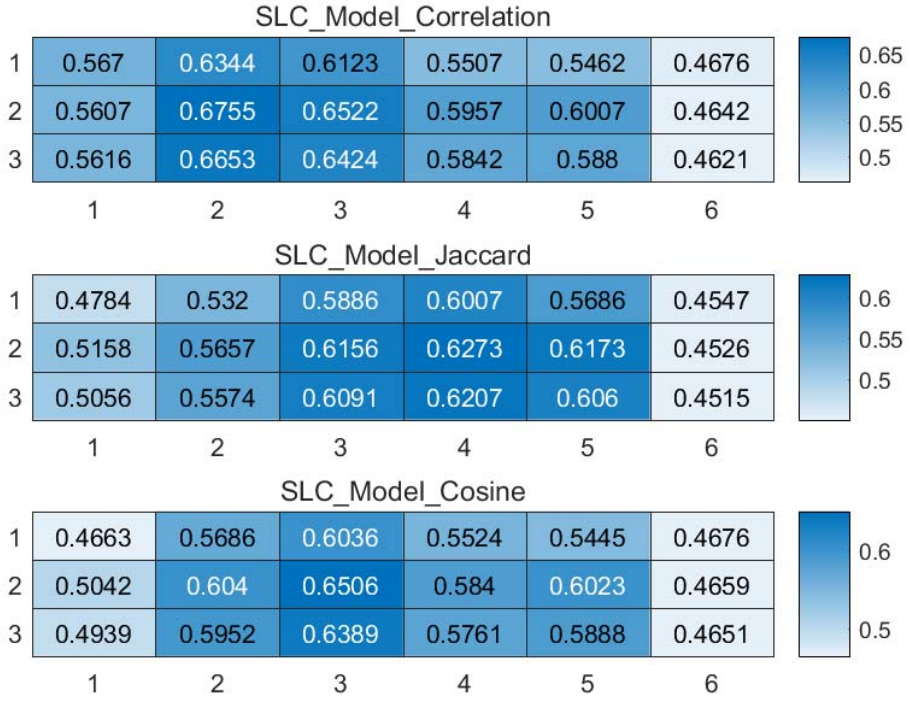


Figure S183: The weighted accuracy on every slice points for *Data_Zeisel* dataset.

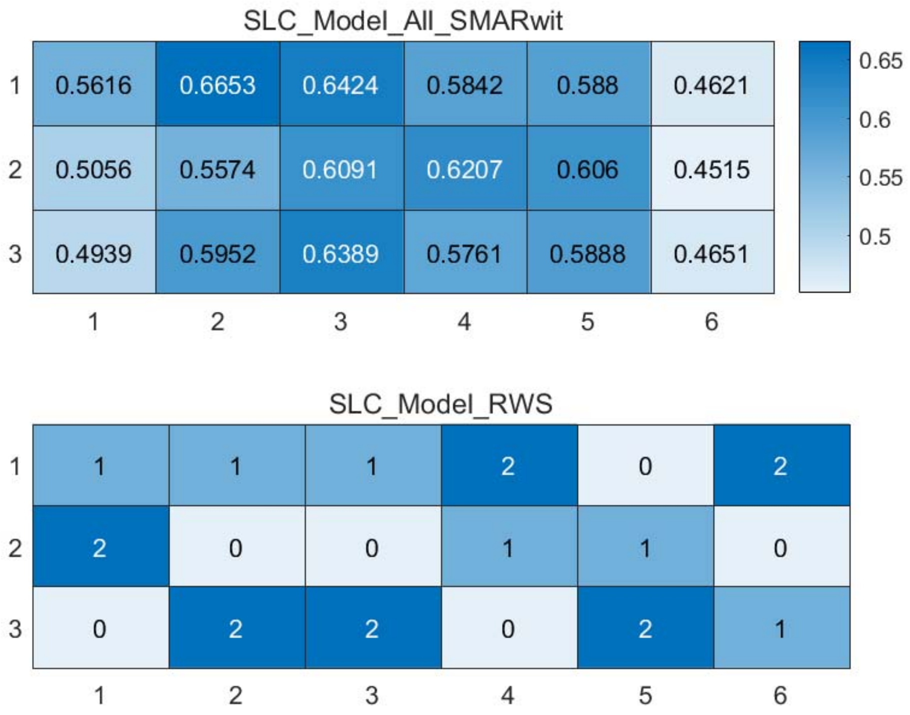


Figure S184: The RWS mode of meta classifiers for *Data_Zeisel* dataset.

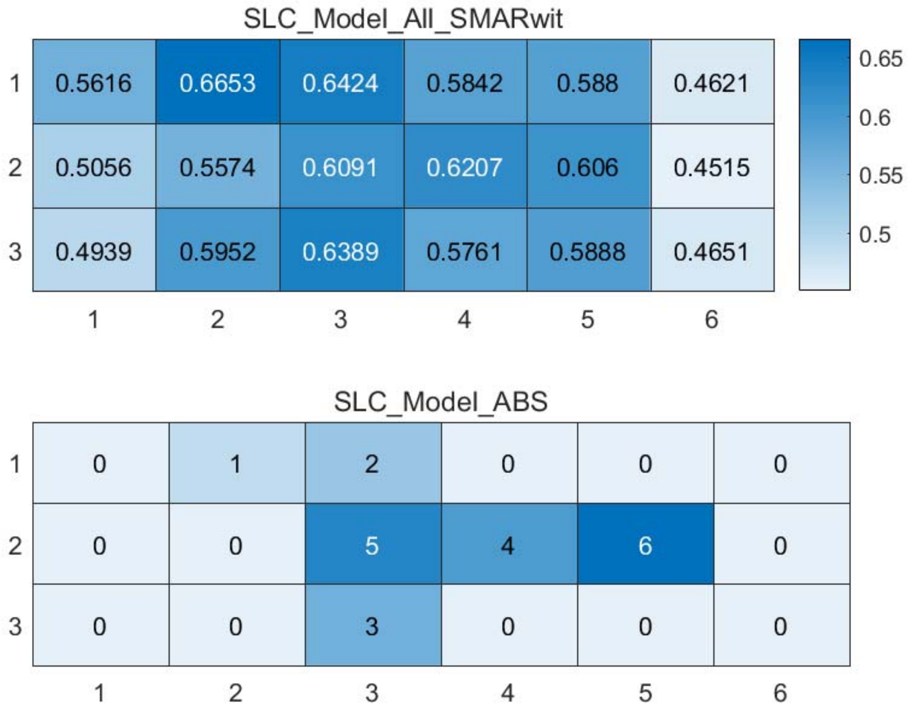


Figure S185: The ABS mode of meta classifiers for *Data_Zeisel* dataset.

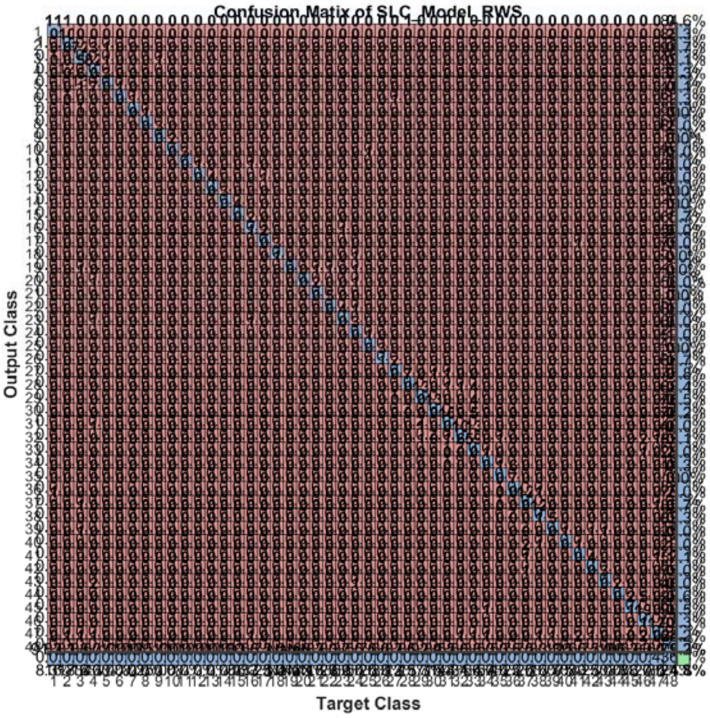


Figure S186: The confusion matrix of ensemble classifier with RWS mode for *Data_Zeisel* dataset. (*Note*: the confusion matrix is too large to display completely in one figure)

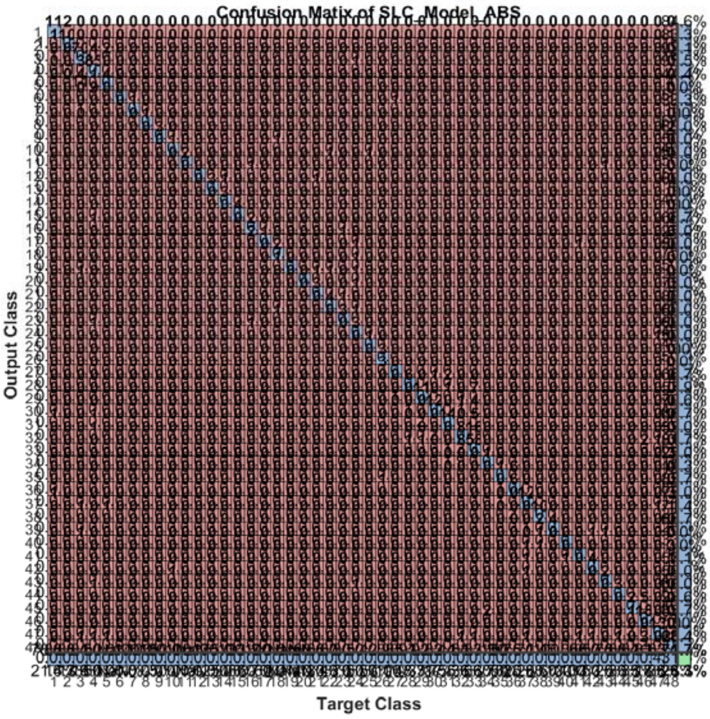


Figure S187: The confusion matrix of ensemble classifier with ABS mode for *Data_Zeisel* dataset. (*Note*: the confusion matrix is too large to display completely in one figure)

Dataset References:

Buettner.mat refers to http://www.ncbi.nlm.nih.gov/pubmed/25599176

Kolod.mat refers to http://www.ncbi.nlm.nih.gov/pmc/articles/PMC4595712/

Pollen.mat refers to http://www.ncbi.nlm.nih.gov/pubmed/25086649

Usoskin.mat refers to http://www.ncbi.nlm.nih.gov/pubmed/25420068

Zeisel.mat refers to https://www.ncbi.nlm.nih.gov/pubmed/25700174

Data_Buettner.mat refers to https://www.ncbi.nlm.nih.gov/pubmed/25599176

Data_Deng.mat refers to http://science.sciencemag.org/content/343/6167/193

Data_Ginhoux.mat refers to https://www.ncbi.nlm.nih.gov/pubmed/26054720

Data_Macosko.mat refers to https://www.ncbi.nlm.nih.gov/pubmed/26000488

Data_Pollen.mat refers to https://www.nature.com/articles/nbt.2967

Data_Tasic.mat refers to https://www.ncbi.nlm.nih.gov/pubmed/26727548

Data_Ting.mat refers to https://www.ncbi.nlm.nih.gov/pubmed/25242334

Data_Treutlin.mat refers to https://www.ncbi.nlm.nih.gov/pubmed/24739965

Data_Zeisel.mat refers to https://www.ncbi.nlm.nih.gov/pubmed/25700174

### Additional Notes:

scASK based on adaptive data slicing is a generic ensemble classification framework especially for classifying cell types based on scRNA-seq data, but also be competent for most existed datasets for classification. Such as UCI Machine Learning Repository: [http://archive.ics.uci.edu/ml/index.php](http://archive.ics.uci.edu/ml/index.php). The only requirement is that the input matrix named in_X should keep rows representing instances and columns representing attributes. The standard input matrix and input labels for scASK should have the format shown below:

| | | | | | | | | | | |
|---|---|---|---|---|---|---|---|---|---|---|
| 0.30103 | 2.0086 | 0 | 0 | 0 | 0 | 0 | 1.30103 | 0.30103 | ··· | 10 |
| 0 | 0 | 0 | 0 | 0 | 0 | 2.089905 | 0 | 1.041393 | ··· | 10 |
| 0 | 0 | 0 | 0 | 0 | 0 | 0 | 0 | 1.544068 | ··· | 10 |
| 0 | 0 | 0 | 0 | 0 | 2.523746 | 0 | 0 | 2.599883 | ··· | 9 |
| 0 | 0 | 1.30103 | 0 | 0 | 0 | 2.584331 | 0 | 2.457882 | ··· | 9 |
| 0 | 0 | 1.70757 | 0 | 0 | 2.523746 | 0.845098 | 0 | 1.78533 | ··· | 9 |
| 0 | 0 | 0 | 0 | 0 | 0 | 3.202761 | 0 | 2.664642 | ··· | 9 |
| 0 | 0 | 0 | 0 | 0 | 1.380211 | 0 | 0 | 1.869232 | ··· | 9 |
| 0 | 0 | 0 | 0 | 1.041393 | 0 | 0.30103 | 0 | 0 | ··· | 9 |
| 0 | 0 | 0.69897 | 0 | 0 | 2.093422 | 1 | 0 | 0 | ··· | 9 |
| 0 | 0 | 0 | 0 | 0 | 0 | 1.819544 | 0 | 1.556303 | ··· | 9 |
| 0 | 0 | 0 | 0 | 0 | 1.518514 | 1.278754 | 0 | 0 | ··· | 9 |
| 0 | 0 | 0 | 0 | 0 | 0 | 0 | 0 | 0 | ··· | 10 |
| 0 | 0 | 0 | 0 | 0 | 0 | 0 | 0 | 0 | ··· | 10 |
| 0 | 0 | 0 | 0 | 0 | 1.770852 | 0 | 0 | 0 | ··· | 10 |
| 0 | 1.380211 | 0 | 0 | 0.60206 | 1.544068 | 2.079181 | 0 | 0.60206 | ··· | 10 |
| 2.39794 | 1.875061 | 0 | 0 | 0 | 1.041393 | 0 | 0 | 0 | ··· | 10 |
| 1.568202 | 1.113943 | 0 | 0 | 0 | 2.255273 | 0 | 0 | 0 | ··· | 10 |
| 0 | 0 | 0 | 0 | 0 | 2.481443 | 2.311754 | 0 | 1 | ··· | 10 |
| 1.146128 | 0 | 0 | 0 | 0 | 0 | 0 | 0 | 2.568202 | ··· | 10 |
| ··· | ··· | ··· | ··· | ··· | ··· | ··· | ··· | ··· | ··· | ··· |
| 0 | 0 | 0 | 0 | 0 | 0 | 0 | 0 | 0 | ··· | 4 |

true_labs

in_X
